# Genotype-by-environment interactions shape ubiquitin-proteasome system activity

**DOI:** 10.1101/2024.11.21.624644

**Authors:** Randi R. Avery, Mahlon A. Collins, Frank W. Albert

## Abstract

In genotype-by-environment interactions (GxE), the effect of a genetic variant on a trait depends on the environment. GxE influences numerous organismal traits across eukaryotic life. However, we have a limited understanding of how GxE shapes the molecular processes that give rise to organismal traits. Here, we characterized how GxE shapes protein degradation, an essential molecular process that influences numerous aspects of cellular and organismal physiology. Using the yeast *Saccharomyces cerevisiae*, we characterized GxE in the activity of the ubiquitin-proteasome system (UPS), the primary protein degradation system in eukaryotes. By mapping genetic influences on the degradation of six substrates that engage multiple distinct UPS pathways across eight diverse environments, we discovered extensive GxE in the genetics of UPS activity. Hundreds of locus effects on UPS activity varied depending on the substrate, the environment, or both. Most of these cases corresponded to loci that were present in one environment but not another (“presence / absence” GxE), while a smaller number of loci had opposing effects in different environments (“sign change” GxE). The number of loci exhibiting GxE, their genomic location, and the type of GxE (presence / absence or sign change) varied across UPS substrates. Loci exhibiting GxE were clustered at genomic regions that contain core UPS genes and especially at regions containing variation that affects the expression of thousands of genes, suggesting indirect contributions to UPS activity. Our results reveal highly complex interactions at the level of substrates and environments in the genetics of protein degradation.

## Introduction

Genotype-by-environment interactions (GxE) occur when a genetic variant’s effect on a trait is environment-dependent. GxE can have profound effects on organismal physiology. For example, in the disease phenylketonuria individuals with two defective copies of the phenylalanine hydroxylase gene develop severe symptoms, including brain damage and intellectual disabilities, if they consume a diet with standard amounts of phenylalanine. However they can avoid most symptoms by consuming a diet with reduced phenylalanine (Bickel et al., 1953; Guthrie, 1961; Shostak, 2003; Widaman, 2009). Other prominent examples of GxE exist in pharmacogenetics, where genetic differences modulate drug efficacy (Pirmohamed, 2023). Thus, understanding the extent and genetic basis of GxE has been a longstanding goal in biomedical research.

Efforts to this end have revealed that GxE is widespread at the level of organismal traits. GxE has been observed for a variety of morphological (e.g., *Drosophila* bristle number; Gurganus et al., 1998) and developmental (e.g., flowering time in various temperatures; Sasaki et al., 2015) traits in numerous organisms. Recent work in humans has begun to explore the impact of environmental factors on traits related to health and disease. By using self-reported and demographic information to integrate environmental factors into genome-wide association studies, these efforts have revealed that GxE shapes the genetics of a variety of clinical syndromes, including depression (C. Li et al., 2022), cancer (Yang et al., 2020), and health-related traits, such as body mass index (Robinson et al., 2017).

However, to what extent GxE occurs at the level of the molecular processes that give rise to organismal phenotypes in humans and other species is poorly understood. A key challenge is that most traits are genetically complex, influenced by variation at loci throughout the genome. Profiling sufficiently large samples to achieve the statistical power needed to detect the effects of multiple loci and their interaction with environmental factors requires assays with high-throughput and quantitative precision. This has limited our ability to understand how environmental factors modulate genetic influences on all but a small number of molecular processes, in particular gene expression and cellular growth. Considerable work has focused on GxE in gene expression (Boye et al., 2024; Grishkevich & Yanai, 2013). For example, studies in flies (Huang et al., 2020), plants (Cubillos et al., 2014), roundworms (Y. Li et al., 2006), and mice (Ballinger et al., 2023) have assayed gene expression in genetically different individuals in different temperatures. In these studies, GxE predominantly occurred at loci that influence gene expression (“expression quantitative trait loci,” eQTLs) via *trans*-acting mechanisms. In humans, eQTLs identified in immune cells display considerable GxE from variation in genes in pathways that become activated upon exposure to various immunogenic stimuli, demonstrating that GxE can occur via direct effects on genes in a relevant pathway (Fairfax et al., 2014; Kim-Hellmuth et al., 2017; Nédélec et al., 2016; Quach et al., 2016). Profiling stimulated human immune cells has also revealed condition-specific *trans*-eQTLs, showing that GxE can also occur via indirect mechanisms (Fairfax et al., 2014; Lee et al., 2014). Context-specific eQTLs displaying GxE are enriched for GWAS signals for complex organismal traits (Kim-Hellmuth et al., 2017; Lea et al., 2022), highlighting the value of profiling molecular traits that give rise to organismal phenotypes.

The yeast *Saccharomyces cerevisiae* has served as a powerful model for dissecting GxE (Yadav & Sinha, 2018) because thousands of natural, genetically different isolates (Warringer et al., 2011), their cross progeny (Bloom et al., 2019; Nguyen Ba et al., 2022; Smith & Kruglyak, 2008), or strains harboring engineered natural variants (Chen et al., 2023) can be exposed to tightly controlled environments at high levels of replication. These approaches have revealed that GxE in cellular growth is widespread. A survey of natural yeast isolates showed that isolates from genetically different populations grew differently in nearly half of 200 assayed environments (Warringer et al., 2011). Later work using linkage mapping in crosses revealed considerable heterogeneity in the genetic architecture of yeast growth in dozens of environments, including loci that only affected growth in specific environments and loci whose direction of effect differed between environments (Bloom et al., 2013; Nguyen Ba et al., 2022). Recently, Chen et al. (2023) revealed that 93.7% of natural variants that had a significant effect on growth in at least one condition showed evidence of GxE (Chen et al., 2023). Additionally, in a cross between two yeast strains, Smith & Kruglyak (2008) found GxE at about 40% of loci that shape transcript abundance in media with glucose versus ethanol as the carbon source.

Despite these foundational insights from humans and model systems, our knowledge about GxE in molecular traits remains limited. Profiling additional molecular processes with known roles in organismal physiology would expand our understanding of how GxE shapes health, disease, and evolution. Protein degradation is an essential molecular process that influences numerous aspects of cellular and organismal physiology. In eukaryotes, most protein degradation (70-80%) is carried out by the ubiquitin-proteasome system (UPS) (Bachmair et al., 1986; G. A. Collins & Goldberg, 2017; Coux et al., 1996; Hershko & Ciechanover, 1998). By degrading substrate proteins, the UPS regulates protein abundance and removes misfolded and damaged proteins from cells (G. A. Collins & Goldberg, 2017; Hanna & Finley, 2007; Varshavsky, 2011). The central importance of UPS protein degradation is illustrated by the defects in this process that occur in numerous human diseases, including cancers, immune disorders, and neurodegenerative diseases (Dantuma & Bott, 2014; Schwartz & Ciechanover, 1999; Shringarpure & Davies, 2002; Zheng et al., 2014). The UPS comprises the ubiquitin system, a collection of enzymes that mark substrate proteins for degradation, and the proteasome, a multi-protein complex that degrades substrate proteins to small peptides. The ubiquitin system recognizes short signal sequences (termed “degrons”; Varshavsky, 1991) in proteins, then marks the substrate protein for degradation by covalently attaching the small protein ubiquitin. Ubiquitinated substrate proteins are bound by the proteasome’s 19S regulatory particle, unfolded, and degraded to short peptides by the proteasome’s 20S core particle (Bett, 2016; Finley et al., 2012; Hershko & Ciechanover, 1998). The proteasome can also bind and degrade certain substrates directly, independent of the ubiquitin system (Finley et al., 2012). UPS protein degradation is organized into multiple distinct pathways based on the ubiquitin system enzymes and proteasome receptors involved in targeting and binding substrates of a given pathway. This can result in highly pathway- and even substrate-specific UPS activity that is tailored to the physiological needs of the cell. For example, during proteotoxic stress, UPS activity towards misfolded proteins can selectively increase (Gardner et al., 2005; Ibarra et al., 2016; Rosenbaum et al., 2011).

We recently showed that UPS activity is a genetically complex trait (M. A. Collins et al., 2022, 2023). By measuring UPS activity towards multiple substrates that engage distinct UPS targeting and degradation pathways, we revealed that many variant effects are substrate-specific in that the magnitude and, in some cases, direction of their effects on UPS activity varied between substrates. Some loci exerting substrate-specific effects contained causal variants in UPS genes, while other loci did not contain any genes with known roles in UPS activity, suggesting indirect effects. However, these experiments were performed in a single environment. The extent of GxE in the genetics of UPS activity is unknown.

Protein degradation is highly environment-dependent, raising the possibility that the genetics of UPS activity is subject to GxE. For example, in environmental conditions that cause misfolded or oxidatively damaged proteins to accumulate, UPS activity increases to clear these molecules from the cell (Finley & Prado, 2020; Grimm et al., 2012; Sontag et al., 2014). In contrast, UPS protein degradation, an ATP-dependent process, decreases in nutrient-poor conditions, to conserve cellular resources (Bajorek et al., 2003; Laporte et al., 2008; Waite et al., 2016). This process is well-characterized in yeast cells, which decrease UPS activity in low glucose environments by sequestering proteasomes in inactive aggregates and in low nitrogen environments by autophagically degrading proteasomes (Laporte et al., 2008; J. Li et al., 2019). These environmental influences on UPS activity, combined with the fact that UPS activity is affected by complex natural genetic variation, suggests that genetic influences on UPS activity are influenced by GxE.

Here, we quantified and mapped GxE in the genetics of UPS activity. By measuring genetic influences on UPS activity towards six substrates that engage distinct UPS pathways in eight environments, we discovered extensive GxE in the genetics of protein degradation. We found hundreds of instances where a locus altered UPS activity in one environment but not another, as well as a smaller number of instances where a locus effect changed direction between environments. Patterns of loci exhibiting GxE were also highly specific to individual UPS substrates. Our results reveal a high degree of environment- and pathway-specific GxE in the genetic architecture of UPS activity and expand our understanding of how GxE shapes molecular traits.

## Results

### Experimental Design Overview

To study GxE in the UPS in a pathway-specific manner, we compared the UPS activity of two genetically divergent yeast strains for six UPS substrates that engage multiple distinct UPS pathways in eight environments comprising multiple starvation and chemical stressors (Fig. 1A). We compared the BY laboratory strain, a close relative of the S288C reference strain, to RM, a vineyard isolate. These strains differ on average one nucleotide per 200 base pairs (bp), providing abundant genetic variation that is known to affect molecular and cellular traits (Albert et al., 2018; Bloom et al., 2013; Brem et al., 2002; Brion et al., 2020; Nguyen Ba et al., 2022) that may be subject to GxE, including UPS activity (M. A. Collins et al., 2022, 2023).

**Fig. 1:**
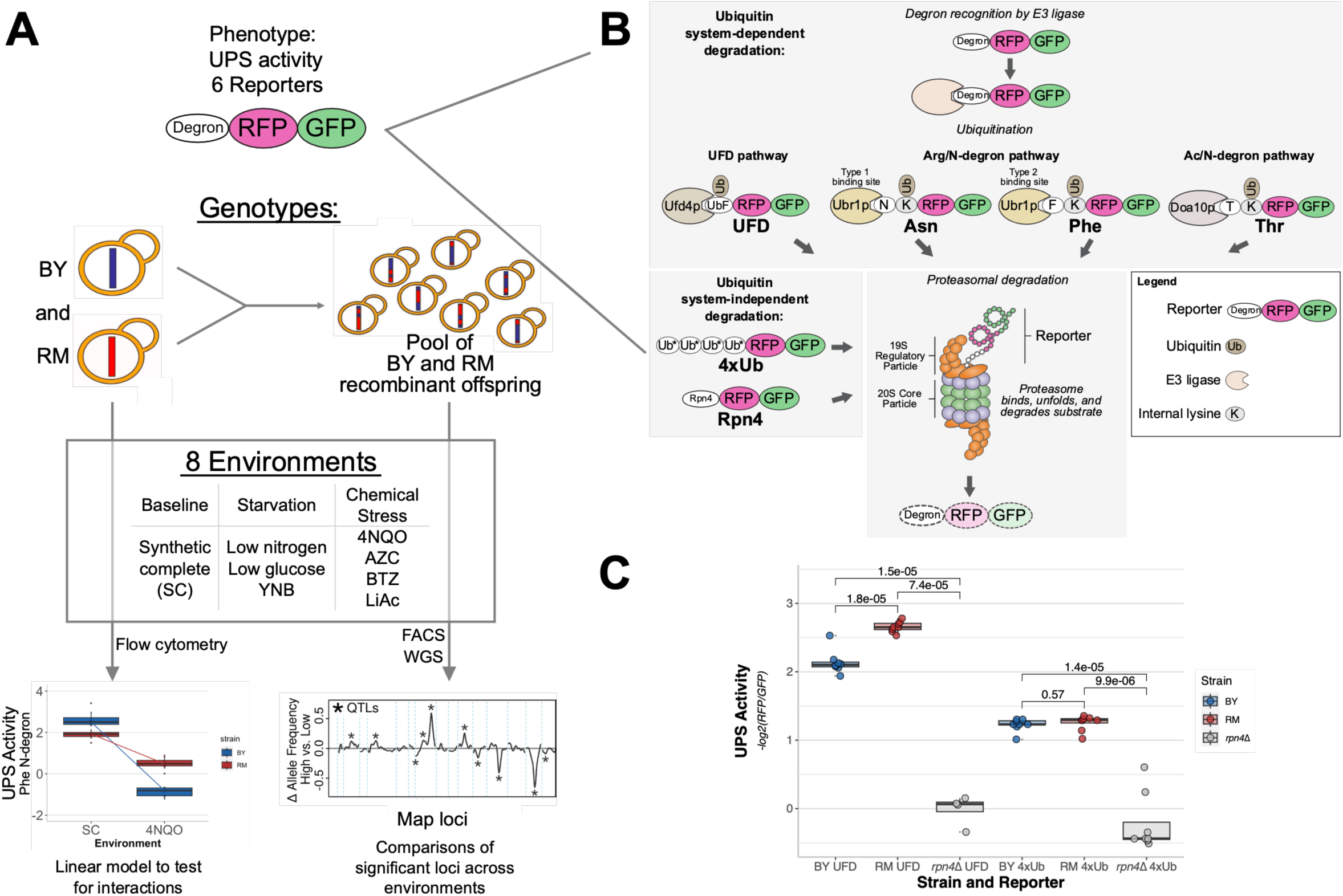
Study design. **A.** Experimental design overview. FACS: fluorescence activated cell sorting. WGS: whole-genome sequencing. **B.** Simplified schematics of the six reporters used in this study to measure ubiquitin system-dependent and -independent UPS pathways. UbF: ubiquitin with G76V substitution. Ub*: ubiquitin with G76V and K29/48/63R substitutions. Adapted from M. A. Collins et al., 2022, 2023. **C.** UPS activity from UFD and 4xUb reporters in BY, RM, and BY *rpn4*Δ. P-values from two-tailed T-tests are indicated.

We selected eight environments predicted to alter UPS activity. Throughout this paper we consider synthetic complete medium (“SC”), a nutrient-rich medium, as the baseline environment for normal growth. UPS activity in SC was compared to seven other environments intended to produce diverse impacts on cellular physiology, some of which have known effects on UPS activity (Supplementary File 1). The environments included three conditions with reduced nutrients compared to SC, such that these “starvation” conditions are predicted to decrease UPS activity: low glucose, low nitrogen, and yeast nitrogen base (YNB) without amino acids. Protein degradation via the UPS plays a critical role in the response to multiple forms of chemical stress. We assayed four chemical stress conditions by adding bortezomib (BTZ), L-azetidine-2-carboxylic acid (AZC), 4NQO, or lithium acetate (LiAc) to SC. Bortezomib (BTZ) inhibits proteasomal protein degradation by tightly and selectively binding the 20S proteasome’s catalytically active site, causing proteolytic stress (Nunes & Annunziata, 2017; Work & Brandman, 2021). The proline analog AZC causes misfolding of nascent proteins, leading to cellular “folding stress” (Rodgers & Shiozawa, 2008; Work & Brandman, 2021) wherein misfolded proteins accumulate in protein aggregates, resulting in increased UPS activity (Work & Brandman, 2021). The mutagen 4NQO has been shown to increase global protein degradation (Burgis & Samson, 2007). Finally, we chose LiAc, a chemical used in yeast transformations (Gietz & Schiestl, 2007). High salt concentrations cause the proteasome’s 19S regulatory particle to dissociate from the 20S core particle (Glickman et al., 1998; Saeki et al., 2000), and lithium chloride has been shown to inhibit purified 20S core particles (Holtz et al., 2003), suggesting that LiAc could decrease UPS activity.

To measure UPS activity, we used tandem fluorescent protein timers (TFTs). TFTs are two-color fluorescent protein constructs that provide high-throughput measurements of protein turnover (Khmelinskii et al., 2012; Khmelinskii & Knop, 2014) (Fig. 1B; Supplementary File 1). In the most common implementation, which is used here, the TFT consists of a linear fusion of a faster-maturing green fluorescent protein (GFP) and a more slowly-maturing red fluorescent protein (RFP). If the TFT’s degradation rate is faster than the RFP’s maturation rate, then the -log_2_ (RFP / GFP) ratio is directly proportional to the TFT’s degradation rate (Khmelinskii et al., 2012; Khmelinskii & Knop, 2014). Because the RFP and GFP are synthesized from the same mRNA transcript, the TFT ratio is independent of the expression level of the TFT (Kats et al., 2018; Khmelinskii et al., 2012; Khmelinskii & Knop, 2014; Kong et al., 2021). TFTs can thus be used to measure protein degradation in genetically distinct cell populations where reporter expression may vary (M. A. Collins et al., 2022, 2023; Khmelinskii et al., 2012). An additional key feature of the TFT system is its transferability. By fusing a TFT to a UPS substrate, the substrate’s degradation rate can be measured, provided its degradation rate is faster than the RFP’s maturation rate (Khmelinskii et al., 2012; Khmelinskii & Knop, 2014). Prior efforts have fused a variety of distinct UPS substrates to TFTs to serve as high-throughput, pathway- and substrate-specific reporters of UPS activity in live cells (M. A. Collins et al., 2022, 2023).

We attached six degron-containing substrate sequences that engage multiple distinct UPS pathways to our TFTs to serve as six reporters of UPS activity (Fig. 1B). The UPS reporters studied here included four substrates targeted by the ubiquitin system (“ubiquitin system-dependent” reporters) and two substrates that are directly bound and degraded by the proteasome (“ubiquitin system-independent” reporters). Three of the ubiquitin system-dependent reporters probe the three branches of the N-degron pathway, a UPS pathway in which a protein’s N-terminal amino acid functions as a degron (Supplementary File 1) (Varshavsky, 2011, 2019, 2024). These include the Type-1 Arg/N-degron pathway (using asparagine as the N-terminal amino acid; “Asn”), which targets basic N-terminal amino acids; the Type-2 Arg/N-degron pathway (phenylalanine; “Phe”), which targets bulky hydrophobic N-terminal amino acids; and the Ac/N-degron pathway (threonine; “Thr”), which targets acetylated uncharged N-terminal amino acids (Varshavsky, 2011, 2019, 2024). These N-degron pathways influence multiple aspects of cellular physiology by regulating protein abundance. To capture genetic influences on protein quality control-associated UPS activity, we constructed a TFT reporter that measures the activity of the ubiquitin fusion degradation (UFD) pathway (Johnson et al., 1995). In the UFD pathway, a non-cleavable ubiquitin moiety acts as a degron that is recognized by the Ufd4p E3 ligase, which targets misfolded proteins and is involved in the response to proteotoxic stress (Devarajan et al., 2020; Johnson et al., 1995; Theodoraki et al., 2012).

To measure genetic effects on proteasome activity separately from the ubiquitin system, we used two reporters containing degrons that are directly bound and degraded by the proteasome. First, the Rpn4 reporter (M. A. Collins et al., 2023) contains the first 80 amino acids of the Rpn4 protein, which are directly bound by the Rpn2p and Rpn5p receptors of the 19S regulatory particle of the proteasome (Ha et al., 2012; Ju & Xie, 2004; Prakash et al., 2004; Xie & Varshavsky, 2001). The second ubiquitin system-independent reporter contains a linear fusion of four ubiquitin molecules (Stack et al., 2000), which functions as a degron that is recognized by the proteasome receptor Rpn13p (Thrower et al., 2000). This “4xUb” reporter serves as a minimal degron that mimics the degradation of the majority of physiological UPS substrates (Inobe et al., 2011; Martinez-Fonts et al., 2020; Thrower et al., 2000; Zhao & Ulrich, 2010). Because the Rpn4 and 4xUb degrons have different sizes, sequence compositions, and structures, we reasoned that they may be influenced by distinct sets of loci, as in our prior studies of ubiquitin-independent substrates (M. A. Collins et al., 2023). Our selection of substrates thus allowed us to capture genetic influences on the activity of multiple UPS pathways involved in physiological protein abundance regulation and protein quality control.

The UFD and 4xUb reporters were developed for this study, and we characterized these reporters using flow cytometry in BY, RM, and in a BY strain with reduced UPS activity due to deletion of the *RPN4* gene (‘*rpn4*Δ’) (Xie & Varshavsky, 2001). As expected, both reporters showed significantly lower UPS activity in the BY *rpn4*Δ strain than in the BY and RM strains when grown in SC (Fig. 1C). RM showed higher UPS activity than BY for UFD (T-test: p-value = 1.8e-5), and there was no difference between BY and RM for 4xUb (p = 0.57). Thus, together with our previous work on N-degron pathways and the Rpn4 reporter (M. A. Collins et al., 2022, 2023), all six reporters provide quantitative, substrate-specific, *in vivo* measurements of UPS activity.

### Widespread GxE in UPS activity between two yeast strains

To estimate the extent of GxE in the UPS, we exposed BY and RM strains carrying one of the six reporters to the eight environments and used flow cytometry to assay UPS activity in 20,000 cells in each of eight biological replicates per combination of strain, reporter, and environment. Compared to the SC baseline, all seven environments affected UPS activity in at least one strain and for at least one reporter (Fig. 2; Supplementary File 1). UPS activity significantly decreased in 39 of 80 comparisons and increased in four comparisons (T-test; Bonferroni-corrected p < 0.05). Of the increases, three were seen for BY in AZC (Asn, Rpn4, and 4xUb), in line with the reported modest increases in *RPN4* expression caused by this treatment in a BY strain (Work & Brandman, 2021). Notably, the effects of environment on UPS activity were highly strain-dependent. For example, all four cases of environmentally-induced increases in UPS activity were seen in only one of the two strains (Fig. 2A-C). Together, these observations suggest widespread GxE in UPS activity.

**Fig. 2:**
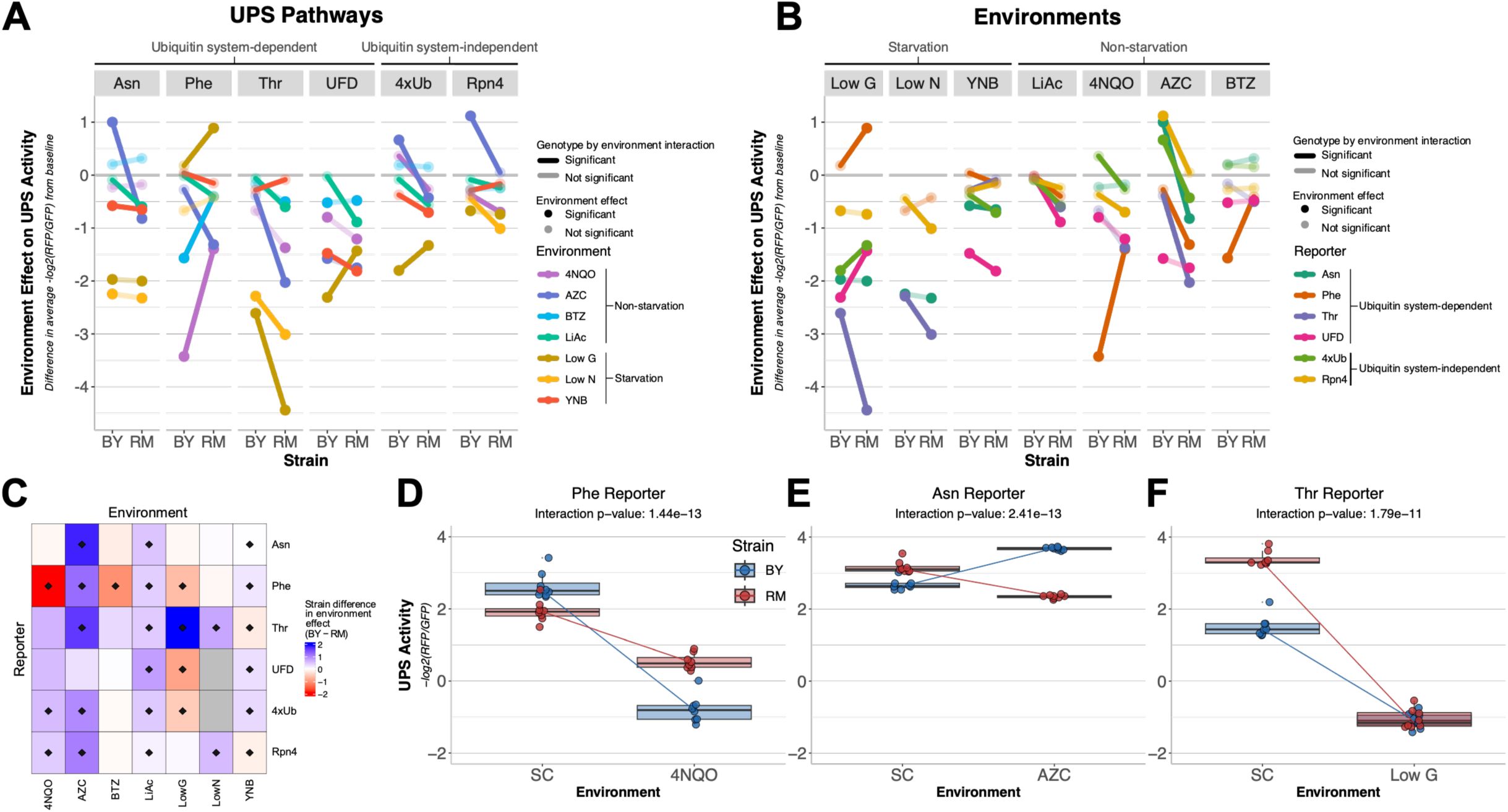
GxE in BY and RM across reporters and environments. **A & B**. Environment effects on UPS activity. Y-axis: the median UPS activity among replicates in SC was subtracted from that in the given environment to visualize environment effects. Negative values indicate that the environment caused a decrease in UPS activity compared to SC, and positive values indicate increased UPS activity. A value of zero means the environment did not affect UPS activity. Significant GxE terms (analysis of variance interaction term p-value < 0.05 after Bonferroni correction) are highlighted by opaque lines. Opaque points indicate a significant difference (T-test; Bonferroni-corrected p < 0.05) in UPS activity between the given environment and SC. **A**. Data organized by pathway. **B**. Data as in A, but reorganized by environment. **C.** Heatmap summarizing strain differences in environment effect. Diamonds indicate significant GxE (Bonferroni-corrected p < 0.05). **D-F.** Reporter / environment combinations that exhibited the most significant GxE, ranked by p-value of the interaction term in a linear model. Eight replicates were measured for each strain / reporter / environment combination. The center line of each box plot corresponds to the median of the eight replicates, with the lower and upper hinges showing the first and third quartiles, respectively. Whiskers extend to 1.5 times the interquartile range and lines connect the respective BY and RM medians. **D**. Phe N-degron reporter in 4NQO. **E**. Asn N-degron reporter in AZC. **F**. Thr N-degron reporter in low glucose.

To search for GxE more formally, we fit linear models that compared UPS activity for a given reporter between BY and RM and between SC and one of the other environments (Methods). GxE was detected in 27 (67.5%) of 40 tests (analysis of variance interaction term p-value < 0.05 after Bonferroni correction), revealing numerous cases in which an environmental effect on UPS activity depended on the strain (Fig. 2A-C; Supplementary File 1; Supplementary File 2).

The most significant interaction effect (p = 1e-13) was seen for the phenylalanine N-degron reporter in 4NQO (Fig. 2D). 4NQO reduced the UPS activity measured by this reporter in both BY and RM (environment main effect: p = 8e-19, T-tests: p ≤ 1e-7). However, the reduction was stronger in BY than in RM, such that BY had higher UPS activity than RM in SC, while it had lower activity than RM in 4NQO. The GxE term with the second most significant interaction effect was seen for the asparagine N-degron reporter in AZC (p = 2e-13) (Fig. 2E). In this case, AZC lowered UPS activity in RM but increased it in BY. This also resulted in a rank order change: in SC RM had higher UPS activity than BY (T-test, p = 3e-5), while in AZC RM had lower UPS activity (p = 2e-17). The GxE term with the third-smallest p-value was observed for the threonine N-degron reporter in low glucose (p = 2e-11) (Fig. 2F). Here, RM showed higher UPS activity than BY in SC (T-test, p = 4e-9). However, in low glucose, UPS activity was much lower for both BY and RM, dropping to the two lowest (out of 96) mean UPS activity values in this experiment and removing the strain difference (T-test, p = 0.72) (Supplementary Fig. 1A).

Each of the reporters and environments showed at least one significant case of GxE (Fig. 2A-C, Supplementary Fig. 1B & C). Among environments, LiAc and YNB showed GxE for all six reporters when compared to SC, while BTZ had the lowest number of significant interaction effects (1 / 6) (Fig. 2B & C, Supplementary Fig. 1C). The five strongest (ranked by the absolute difference in strain response to the given environment) and most statistically significant cases of GxE were all for N-degron pathway reporters (Fig. 2A & C, Supplemental File 1). N-degron substrates require ubiquitin system targeting and, for two of the three studied substrates (Asn and Thr), pre-processing to produce functional N-degrons. The complex cascade of molecular events required to degrade these substrates may result in stronger GxE in the genetics of UPS activity towards N-degrons relative to the other substrates tested here.

Some environments had consistent effects across reporters (Fig. 2B & C). For example, in LiAc, BY showed no significant change in UPS activity for any reporter, while RM showed at least nominally (T-test, p ≤ 0.04) significant reductions for all reporters. Following treatment with AZC, BY had higher UPS activity than RM for all reporters, either because AZC increased activity in BY while activity in RM was unchanged (Rpn4) or even reduced (Asn, 4xUb), or because RM experienced greater reductions in activity than BY (Phe, Thr). Other environmental effects were heterogeneous across UPS pathways. For example, glucose starvation led to a larger decrease in UPS activity in RM than in BY for the Thr N-degron reporter (p ≤ 5e-11, Fig. 2B), but showed the opposite pattern for UFD and 4xUb (p ≤ 4-e6), and even increased degradation of the Phe N-degron reporter in RM (p = 1e-5) with no change in BY (p = 0.25). Our results reveal previously-unappreciated complexities in the influence of strain background, substrate, and environment on UPS activity. Specifically, the effects of multiple environments commonly reported to consistently affect UPS activity were distinct, and in some cases discrepant, between strain backgrounds and UPS substrates (Fig. 2A-C).

### Heterogeneous genetic architectures of UPS activity across pathways and environments

To identify genetic loci affecting UPS activity between BY and RM, we used a genetic mapping approach based on bulk segregant analysis (Albert et al., 2014; Brion et al., 2020; Ehrenreich et al., 2010; Michelmore et al., 1991) (Methods). Briefly, UPS activity was measured in large, genetically diverse cell populations of haploid meiotic recombinant progeny (“segregants”) generated by mating RM with BY strains harboring the UPS activity reporters. We exposed two independent segregant populations derived from independent BY / RM matings to each of the eight environments and used fluorescence-activated cell sorting (FACS) to collect pools of segregants from the extreme tails of the UPS activity distribution. Sorted segregant pools were then whole-genome sequenced to determine BY and RM allele frequencies.

Genome regions where pools with high and low UPS activity differ in allele frequency indicate quantitative trait loci (QTLs) that influence UPS activity (Fig. 1A).

In the baseline SC condition, we identified 46 QTLs across the six UPS reporters. Four of these reporters (Rpn4, Asn, Phe, and Thr) were previously mapped in SC (M. A. Collins et al., 2022, 2023). To assess reproducibility, we compared QTLs identified here and in our previous studies. We observed high concordance with prior results in terms of the QTLs detected, the corresponding allele frequency differences, and the overall shape of the QTL traces (Supplementary Fig. 2). Of the 39 QTLs identified here for these four reporters, 30 were also seen in Collins et al. (2022 & 2023) (Supplementary Fig. 2E). The remaining nine QTLs had significantly lower LODs and effects sizes, as measured by the absolute allele frequency difference (T-test: p = 0.004, and 0.0001, respectively) (Supplementary Fig. 2F), suggesting that they may have been missed due to limited power. Thus, our approach represents a highly reproducible method for characterizing the genetics of UPS activity.

Across the six UPS reporters and eight environments, we identified a total of 416 QTLs (Fig. 3A-C, Supplementary Fig. 4A; Supplementary File 1). All 47 assayed reporter / environment combinations (Methods) had at least one QTL (Supplementary File 3). The number of QTLs across environments and reporters ranged from one (4xUb in LiAc) to 19 (Asn in low glucose and Thr in low nitrogen) (Fig. 3D, Supplementary File 3). Among reporters, the largest number of QTLs was found for the Thr reporter (n = 111) and the fewest for the 4xUb reporter (n = 30), when summing across all eight environments (Fig. 3B).

**Fig. 3:**
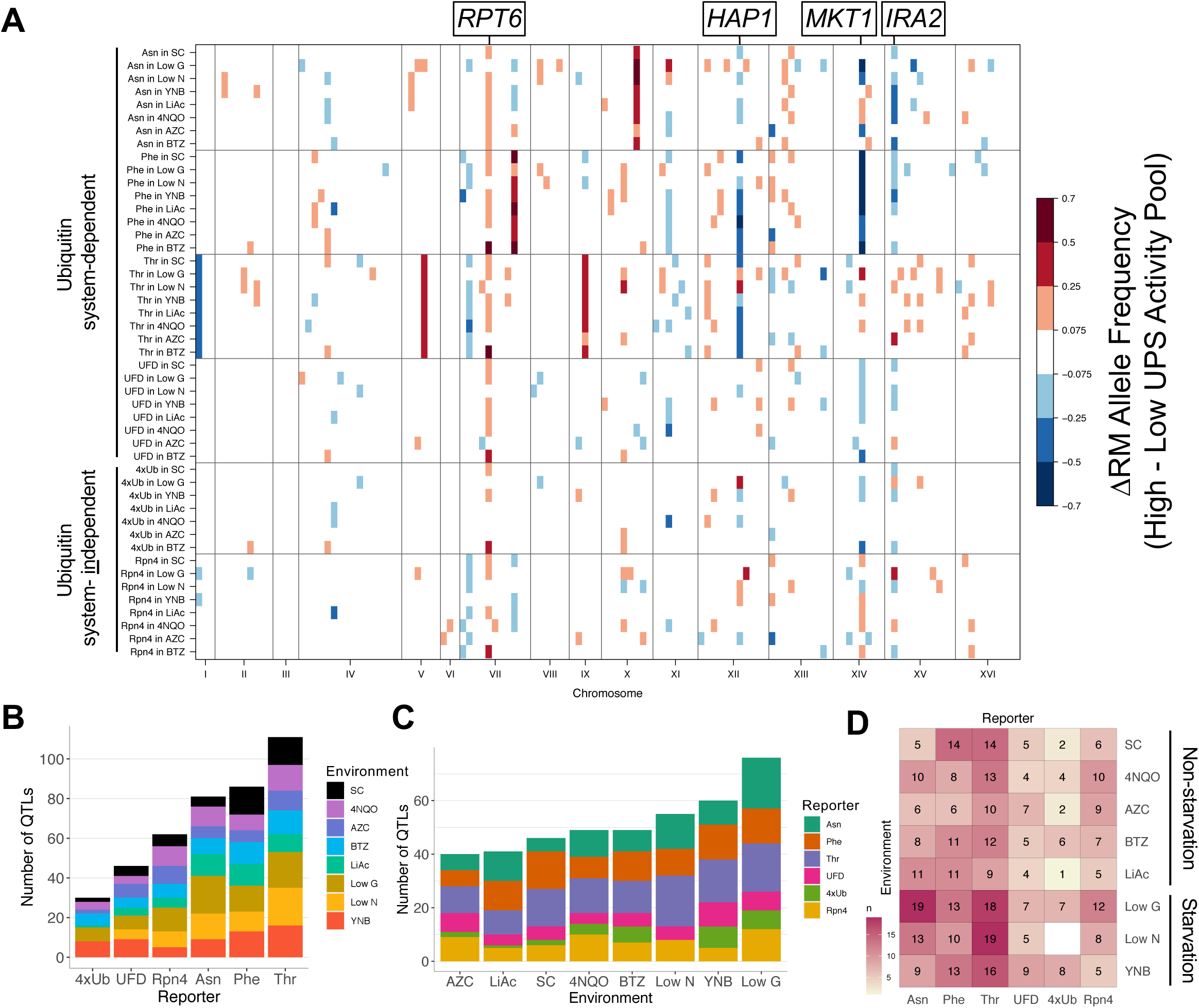
UPS activity QTLs across reporters and environments. **A.** QTL mapping results for the six reporters across eight environments. Colored blocks denote genome bins that contain QTLs detected in each of two independent biological replicates, colored according to the direction and magnitude of the effect size, expressed as the RM allele frequency difference between high and low UPS activity pools. Candidate causal genes discussed in the text are indicated. No data were collected for 4xUb in low nitrogen (Methods). **B.** Number of QTLs found per reporter and environment. **C.** Data as in B, but rearranged by environment. **D.** Heatmap showing the number of QTLs per reporter / environment combination.

The ubiquitin system-dependent pathways had 324 QTLs across the eight environments. Among these, the UFD pathway (Fig. 1B), which was not mapped in our previous studies, had five QTLs in the baseline SC condition (Fig. 3B, Supplementary Fig. 3A). The five QTLs for UFD in SC include a QTL on chromosome XII (peak position at 950,450) that was not seen for other reporters here or previously. In this region, the gene *RPN13* encodes a subunit of the 19S regulatory particle of the proteasome that acts as a ubiquitin receptor. There are multiple BY-RM promoter and missense variants at *RPN13*, along with a strong *cis*-eQTL for this gene (Albert et al., 2018), suggesting *RPN13* as a causal gene for this QTL.

The two ubiquitin system-independent reporters had 92 QTLs across environments (Fig. 3). The 4xUb reporter, which we had not assayed in previous work, had two QTLs in the baseline SC condition (n = 2), which is the fewest of all the reporters in SC (Fig. 3, Supplementary Fig. 3B). Both of these 4xUb QTLs were identified for other reporters (M. A. Collins et al., 2023). Specifically, the QTL on chromosome VII contains *RPT6*, which encodes an ATPase of the 19S regulatory proteasome particle. At a causal variant in the *RPT6* promoter, the derived RM allele broadly increases the activity of multiple ubiquitin system-dependent and -independent UPS pathways by increasing *RPT6* expression (M. A. Collins et al., 2023), suggesting that this variant also affects 4xUb. The QTL on chromosome XV was previously seen for the Rpn4 reporter (M. A. Collins et al., 2023). The causal gene in this QTL is likely *IRA2*, a gene that underlies a *trans*-eQTL hotspot that affects the expression of thousands of genes and numerous growth traits (Lutz et al., 2021; Smith & Kruglyak, 2008). While altered *RPT6* expression underlying the QTL on chromosome VII likely affects UPS activity directly, coding variants in *IRA2* (Lutz et al., 2021) that alter the activity of the Ira2 RAS signaling regulator likely affect UPS activity indirectly.

Collapsing combinations of pathways and environments into physiologically relevant categories revealed that ubiquitin system-dependent pathways had significantly more QTLs (median = 10) than ubiquitin system-independent pathways (median = 6; Wilcoxon test p-value = 0.004; Fig. 3B), indicating greater genetic complexity. The substrates of the ubiquitin system-dependent pathways must undergo binding and processing by various enzymes before being bound by the proteasome. The genes encoding this machinery provide additional targets for genetic variation, likely contributing to the higher number of QTLs observed for these pathways compared to ubiquitin system-independent pathways.

Among environments and across all six reporters, most QTLs were found in low glucose (n = 76) and the fewest for AZC (n = 40) (Fig. 3C). The three starvation environments (low glucose, YNB, and low nitrogen) had significantly more QTLs (median = 10) than the non-starvation environments (median = 7) (Wilcoxon test p-value = 0.012). One potential explanation for these results is that starvation environments may have more wide-reaching, systemic effects on cellular physiology than the chemical stressors tested here. As a result, they may create more opportunities for variant effects that alter UPS activity through potentially highly indirect mechanisms. We also note that wild strains, such as RM, commonly undergo periods of nutrient deprivation, akin to the starvation environments tested here (Hong & Gresham, 2014; Wenger et al., 2011). Consequently, they may harbor genetic variation reflecting physiological adaptation to nutrient-poor environments, such as increased protein turnover for amino acid recycling (Vabulas & Hartl, 2005). Consistent with this notion, the RM allele increased UPS activity more often than the BY allele overall (binomial test p = 0.04), and specifically for starvation environments (binomial test p = 0.04) (Supplementary Fig. 4B & C), congruous with Collins et al. (2022 & 2023).

Many QTLs mapped to the same genomic locations (Fig. 3A). To quantify the number of unique locations, we counted the number of QTL peaks within 128 genomic bins of 100 thousand base pairs (kb) (M. A. Collins et al., 2022). The top four bins accounted for 28% (116 / 416) of the QTLs, illustrating that variation at a few locations underlies many of the effects on UPS activity. The bin on chromosome VII at 400 – 500 kb, which contains *RPT6,* harbored the most QTL peaks (n = 32, Fig. 3A). For all of these, the RM allele increased UPS activity, consistent with *RPT6* as the causal gene in this bin. This region contains other genes involved in the UPS with sequence variation between BY and RM: *SCL1*, which encodes the alpha 1 subunit of the 20S proteasome, and *RPN14*, an assembly-chaperone for the 19S regulatory particle. Thus, this locus likely shapes UPS activity directly via *RPT6*, and potentially *SCL1* and *RPN14* as well.

Three bins on chromosomes XIV, XII, and XV contained 31, 26, and 26 QTLs, respectively (Fig. 3A). The direction of effect of the QTLs in these bins depended on the pathway and environment (Fig. 3A). These bins contain the genes *MKT1, HAP1,* and *IRA2,* respectively, which are all known to harbor variation that results in hotspots that affect the expression of thousands of genes in *trans* (Albert et al., 2018). None of these genes has obvious connections to UPS function: *MKT1* encodes a poorly characterized protein that appears to bind certain mRNAs for genes with mitochondrial functions (Dimitrov et al., 2009; Wickner, 1987); *HAP1* encodes a transcription factor that activates genes involved in osmotic stress (Gaisne et al., 1999); and *IRA2* encodes a regulator of Ras signaling (Tanaka et al., 1990). Therefore, the wide-reaching effects of these three genes likely alter UPS activity via indirect mechanisms.

Most QTLs were not specific to a single environment (Supplementary Fig. 4A), with two notable exceptions. First, is a locus on chromosome IV at 490 – 570 kb for LiAc (Supplementary Fig. 4A). Here, the BY allele was associated with higher UPS activity than the RM allele. This locus contains *ENA1, ENA2,* and *ENA5*, which all encode sodium pumps. *ENA1* and *ENA5* both contain multiple missense and frameshift variants between BY and RM, and the ENA locus harbors structural variation among yeast strains (Treusch et al., 2015; Warringer et al., 2011). Collectively, this variation likely leads to the QTLs seen here in the LiAc environment, where higher activity of BY ENA alleles may reduce LiAc concentrations in the cell compared to RM alleles, alleviating osmotic stress on the UPS. At the *RPT6* locus on chromosome VII with QTLs in multiple environments, BTZ stood out in that the QTLs in BTZ had by far the largest effects at this locus (Supplementary Fig. 4A). This locus also contains *PDR1*, which encodes a transcription factor that regulates genes involved in the yeast pleiotropic drug response (Moye-Rowley, 2003), perhaps affecting cellular BTZ concentrations. Taken together, these QTL mapping results show that in addition to being highly substrate-specific (M. A. Collins et al., 2022, 2023), genetic influences on UPS activity also depend on the environment.

### GxE arises from loci that affect the UPS through direct and indirect mechanisms

The distinct patterns of QTLs across pathways and environments indicated a high degree of GxE in UPS activity. To identify specific loci whose effects depend on the environment, we classified QTLs into three categories based on pairwise comparisons between the baseline SC condition and individual environments. While these comparisons do not account for GxE among non-SC conditions, the patterns described here are broadly representative of other environment comparisons (Fig. 3). In our data, GxE at individual loci can be detected in two forms. First, a QTL may be detected in one environment but not in the other, which we refer to as “presence / absence” GxE. We defined such QTLs as those detected in both replicates of one environment and with no QTLs within 100 kb in either replicate of the other environment (based on QTL peak positions as in prior studies (M. A. Collins et al., 2022, 2023; Methods; Fig. 4A & B).

**Fig. 4:**
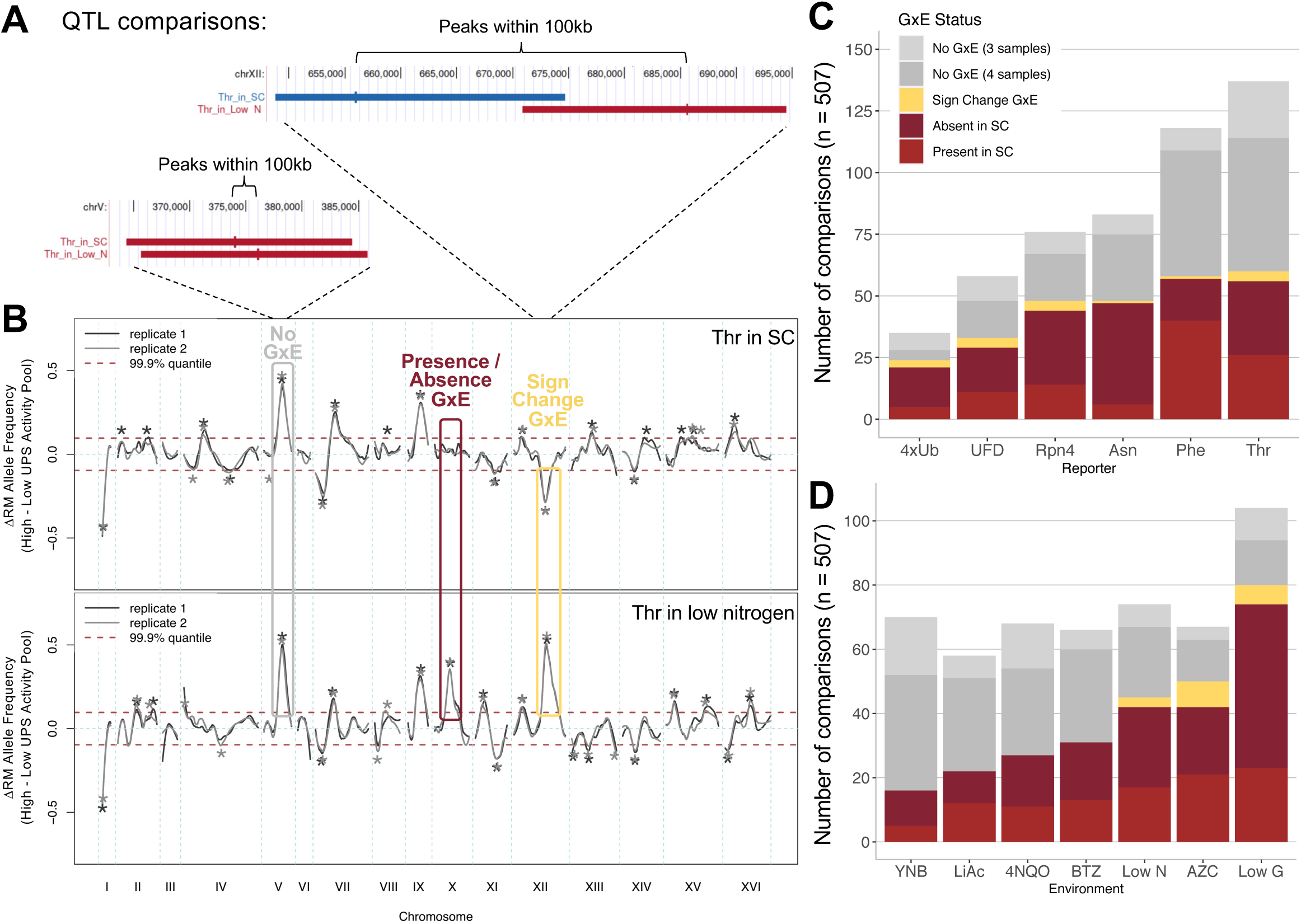
GxE at individual loci. **A & B.** Examples of pairs of QTLs that show no GxE, presence / absence GxE, and sign change GxE. **A.** Peaks (short vertical lines) of QTLs (horizontal bars; positions averaged across the two biological replicates) in the two compared conditions must be within 100 kb to be considered present in both environments. Color indicates direction of allelic effect as in Fig. 3A. Blue: BY allele increases UPS activity, red: RM allele increases UPS activity. **B.** QTL traces for the Thr N-degron reporter in SC (top) and in low nitrogen (bottom). Boxes highlight examples of the three categories of pairwise comparisons of loci. **C.** Pairwise comparisons of loci between SC and other environments, across reporters. Light gray indicates comparisons where a QTL was present in both replicates of one environment and only one replicate of the other environment, with the same direction of effect. **D.** Data as in C, but arranged by environment.

By requiring a locus to be absent or present in both biological replicates, we focus on the strongest cases of presence / absence GxE, reducing the chance that a locus is actually present in both environments but happened to escape detection in one environment due to insufficient statistical power. Second, a QTL may be detected in both environments but with opposing directions of effect, which we call “sign change” GxE. Sign change GxE was defined as QTLs detected in both replicates of one environment and at least one replicate of the other environment, with opposite effect direction (Fig. 4A & B). Fig. 4B displays QTL traces with examples of these two categories. Finally, QTLs within 100 kb of each other with the same direction of effect between environments were considered not to show GxE (Fig. 4B). We exclusively compared QTLs from the same reporter. A total of 507 comparisons between QTLs in different environments were categorized according to this scheme (Fig. 4C & D; Supplementary File 1).

All reporters and all environments had loci exhibiting GxE (Fig. 4C & D, Supplementary Fig. 5). Presence / absence GxE was seen for half (254) of the 507 comparisons (Fig. 4C & D). For the Asn N-degron reporter, presence / absence GxE mostly involved QTLs that were absent in SC but present in another environment (Fig. 4C). Conversely, for the Phe N-degron reporter, presence / absence GxE mostly involved QTLs that were present in SC but absent in the other environment (Fig. 4C). Sign change GxE was much rarer (17 / 507 comparisons, Fig. 4C & D), but was nonetheless seen for all reporters (Fig. 4C), and was observed in low nitrogen, AZC, and low glucose environments compared to SC (Fig. 4D).

Ubiquitin system-independent pathways had a slightly larger proportion of QTLs with GxE than ubiquitin system-dependent pathways (Wilcoxon p = 0.046, Supplementary Fig. 5). There was no difference in the proportion of loci with GxE between starvation and non-starvation environments (Wilcoxon p = 0.82, Supplementary Fig. 5), as illustrated by the fact that both the fewest (YNB), and the most (low glucose) QTLs exhibiting GxE were seen for starvation conditions (Fig. 4D). These results show that half of the loci that shape UPS activity are subject to GxE, primarily via presence / absence of a given locus in different conditions.

We examined the loci of QTL comparisons showing GxE using genomic bins as above, and observed a non-uniform distribution (Fig. 5A). Almost all (71/78, 91%) bins that contained any QTLs contained loci that exhibited GxE (Fig. 5A & B). The five bins with the most GxE cases contained 23% (63 / 271) of all GxE cases, showing that a small portion of the genome harbors much of the GxE seen at individual QTLs (Fig. 5A). Only two of these top five bins contained candidate genes clearly related to the UPS: the bin containing *RPT6*, with 10 GxE cases, and a bin at position 0 – 100 kb on chromosome XIII that contained 14 cases of GxE across all six reporters (Fig. 5A). The latter bin contains the *BUL2* gene, which encodes a component of the Rsp5p E3-ubiquitin ligase complex. Two of the top GxE bins correspond to the *trans*-eQTL hotspots at *IRA2* (15 GxE cases) and *HAP1* (11 GxE cases) (Fig. 5A). A second bin on chromosome XIII, at 300 – 400 kb, had 13 GxE cases spanning all six reporters and did not contain an obvious candidate gene. Overall, QTLs with cases of GxE tended to be located in bins that also contained a large number of QTLs of any type (Spearman correlation across 78 bins with any QTLs: rho = 0.73, p = 5e-14; Fig. 5B). These data show that loci subject to GxE occur throughout the genome but are clustered in regions with many QTLs, including those caused by variation in core UPS genes as well as indirect, pleiotropic regulators.

**Fig. 5:**
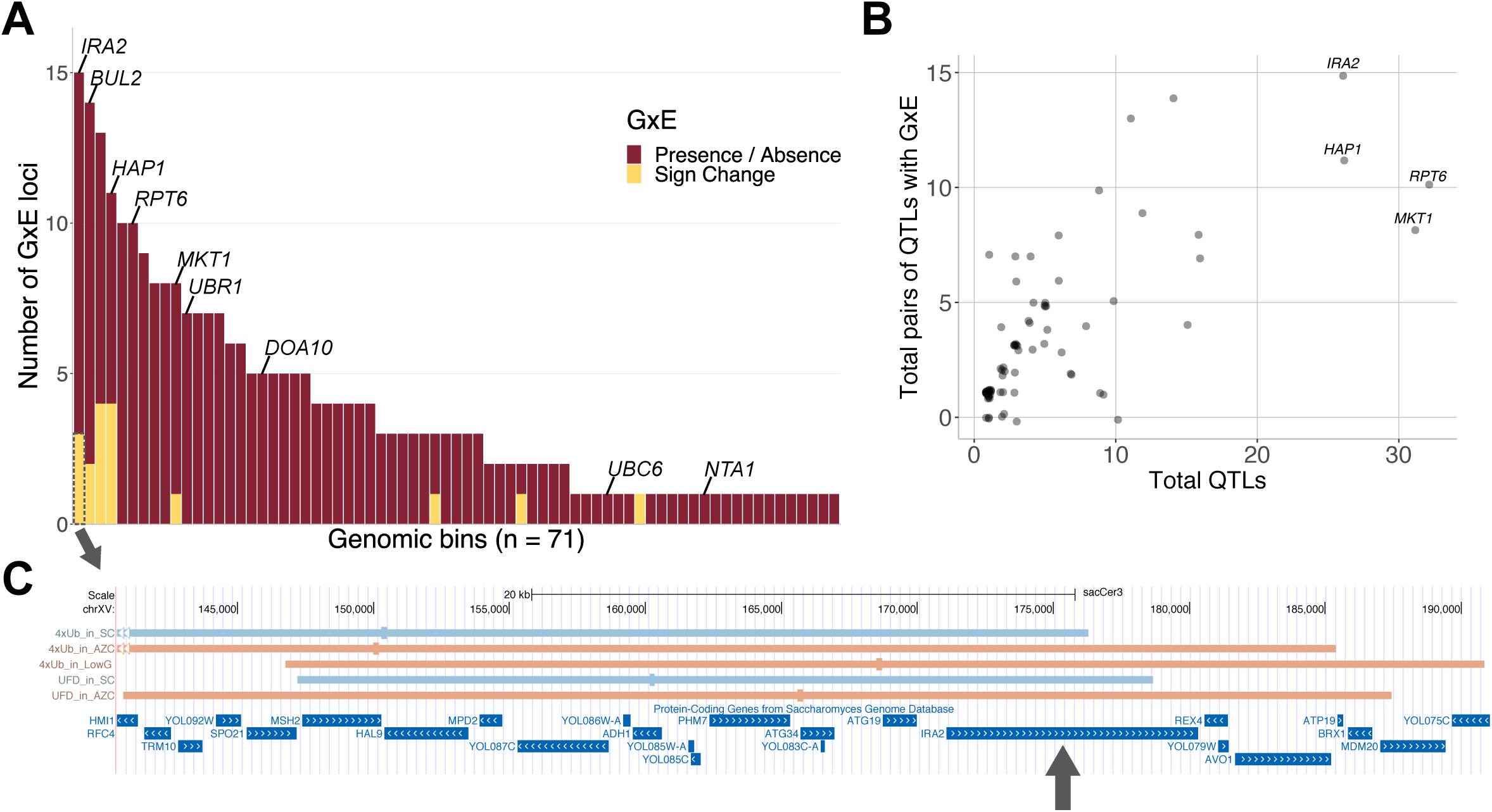
Patterns of QTLs with GxE across the genome. **A.** Distribution of QTL comparisons that exhibited GxE in 100 kb bins. Shown are 71 of 128 bins that contained QTL comparisons with GxE. Bins are sorted based on the number of GxE cases they contain, followed by genomic position to break ties. Candidate causal genes are indicated. **B.** A comparison of the number of total QTLs and of QTL comparisons exhibiting GxE for each of 78 genomic bins that contained at least one QTL. Spearman correlation: rho = 0.73, p = 5e-14. **C.** Locus plot showing five QTLs involved in three sign changes (one for UFD, two for 4xUb) in the bin with the most cases of GxE. Genes in this region are indicated, with *IRA2* highlighted by the arrow. QTL confidence intervals are shown as horizontal bars and peaks are indicated by the small rectangle within each QTL. QTL colors indicate direction and strength of effect as in Fig. 3A. The QTLs for 4xUb in SC and AZC extend leftwards to position 124,750 and 112,300, respectively.

Sign change is a particularly interesting form of GxE, in which the same locus results in opposite effects on UPS activity depending on the environment. The 17 cases of QTL comparisons with sign change GxE involved 29 unique QTLs (Supplementary File 1). Examination of the peaks and confidence intervals of these 29 QTLs revealed that they clustered at seven genomic regions (Fig. 5C and Supplementary Fig. 6A-F). Of these regions, three corresponded to the *trans*-eQTL hotspots at *HAP1* (7 QTLs involved in sign changes, Supplementary Fig. 6A), *IRA2* (5 QTLs, Fig. 5C), and *MKT1* (3 QTLs, Supplementary Fig. 6B). These genes have no obvious direct connections to the UPS. Two regions had no candidate genes with obvious UPS functions: a region from 280 – 470 kb on chromosome XIII with six sign change QTLs (Supplementary Fig. 6C), and a sign change pair (for Rpn4 in AZC), which was located ∼98 kb from *UBC6* (Supplementary Fig. 6D). While Collins et al. (2022) identified *UBC6* as a causal gene affecting degradation of the Thr N-degron reporter, this gene encodes the E2 ubiquitin-conjugating enzyme of the Ac/N-degron pathway (Varshavsky, 2024). Its function in the ubiquitin system and its fairly large distance from the sign change pair makes it unclear if *UBC6* is a causal gene for the ubiquitin system-independent Rpn4 reporter, with no other obvious candidate genes in this region. The remaining two regions with sign change QTLs did contain likely causal genes with direct UPS functions: one sign change pair (for the UFD pathway in AZC) at *RPT6* (Supplementary Fig. 6E), and four such QTLs at *BUL2* (Supplementary Fig. 6F). Notably, none of the remaining causal genes that we previously determined to shape UPS activity (*UBR1*, *UBC6, NTA1*, and *DOA10*; M. A. Collins et al., 2022) had sign changes between SC and other environments, even though they did show presence / absence GxE. All of these genes encode core UPS components. Thus, loci with genes that may affect UPS activity in a direct fashion (*RPT6* and *BUL2*) accounted for only 21% (6 / 29) of the QTLs involved in sign change GxE. Most of the sign change QTLs (52%, 15 / 29) appear to arise from genes that shape UPS activity indirectly.

In sum, these results suggest that GxE in the UPS, especially sign change GxE, is mostly caused by indirect mechanisms, such as widespread changes in gene expression due to *trans*-eQTL hotspots, rather than by variation in core genes directly involved in the UPS.

## Discussion

To characterize GxE in the genetics of protein degradation, we measured UPS activity towards six distinct substrates in single cells of two strains of *S. cerevisiae* and their progeny across eight environments. GxE was pervasive between the two strains. The activity of every measured pathway was modified by the environment, and all of the tested environments led to GxE.

The BY and RM strains differed greatly in how they responded to a given environment. Remarkably, UPS activity increased in one strain but decreased in another for some combinations of reporter and environment. Previous studies of the UPS using some of the environments studied here were based on lab strains, suggesting that some published treatment effects may not represent the *S. cerevisiae* species as a whole. Strain-dependency of treatment effects has been widely documented, including between closely related strains of mice (Simon et al., 2013) and yeast (Elserafy & El-Khamisy, 2018; Matheson et al., 2017), and our results reinforce the value of studying physiological effects in multiple genetic backgrounds including those that have evolved in different environments.

The interaction of genetics and environment was apparent for loci shaping UPS activity. All reporter / environment combinations had unique QTL patterns, and about half of the QTL comparisons we conducted revealed evidence of GxE. Presence of a given QTL in one but not another environment was by far the predominant form of GxE, making up 94% of the detected GxE cases. In spite of the large number of QTLs showing GxE, almost no QTLs were entirely specific to a particular environment, with the ENA locus in LiAc as the closest exception. Most QTLs were seen in several, but not all environments. Thus, the distinct QTL patterns for specific pathway / environment combinations were formed from subsets of the total set of QTLs identified across the entire study.

The QTLs we identified for different environments and pathways, as well as QTLs with GxE, tended to be clustered at certain genomic locations, as seen in our previous work on the UPS (M. A. Collins et al., 2022, 2023) and reflecting work on GxE in yeast gene expression (Smith & Kruglyak, 2008) and in other systems such as flowering time in *A. thaliana* (Sasaki et al., 2015). GxE tended to be seen in regions that had many QTLs overall (Fig. 5B). Thus, when genetic variation leads to changes in UPS activity, it is likely that this variation will be subject to GxE in some environments.

A genome region with numerous QTLs and cases of GxE contained *RPT6*, which we earlier showed to affect the Rpn4 reporter as well as the Ac/N-degron pathway (M. A. Collins et al., 2022, 2023). Here, the *RPT6* locus affected all six assayed reporters in at least one condition. Thus, the causal variant in the *RPT6* promoter appears to have wide-reaching effects on UPS activity. *RPT6* can be considered a “core” gene under an omnigenic model of complex traits (Boyle et al., 2017), in which core genes encode proteins that are directly related to the given trait, while “peripheral” genes act as indirect regulators that shape complex traits through *trans*-acting effects on core genes. All core UPS genes we previously showed to cause UPS activity variation (*UBR1*, *UBC6*, *NTA1*, *DOA10*) had QTLs and presence / absence GxE in this study.

Many individual QTLs, QTLs with GxE, and in particular sign change QTLs, occurred at genes known to cause *trans*-eQTL hotspots, where variation at a single gene affects the expression of hundreds or thousands of genes throughout the genome (Albert et al., 2018; Brem et al., 2002; Smith & Kruglyak, 2008; Zhu et al., 2008). These loci exercise their broad effects through indirect mechanisms in which they alter cellular states that in turn affect many downstream traits, including growth in diverse conditions (Bloom et al., 2013; Renganaath & Albert, 2023). Their many pleiotropic effects include UPS activity (M. A. Collins et al., 2022), for which they can be interpreted as “peripheral” genes under an omnigenic model.

Our results here show that these hotspots are also focal points for GxE in UPS activity. Their indirect mode of action likely presents numerous molecular steps before reaching the UPS, offering opportunities for a given environment to change how their effects are propagated. This stands in contrast to variation at core UPS genes, which was less prone to GxE, in particular sign changes. The direct effects of core genes on the UPS may be less responsive to environmental influences compared to the more indirect, pleiotropic hotspot modulators.

Our study had several limitations. Because the bulk-segregant mapping design we employed makes it difficult to rigorously detect differences in magnitude of locus effect in the same direction, we did not search for such loci even though magnitude GxE could be prevalent (Cubillos et al., 2014; Smith & Kruglyak, 2008). As such, our GxE QTL results are conservative in that they only search for extreme cases in which a QTL is present in only one condition or switches sign. Due to linkage in the segregant population, a sign change QTL could reflect two presence / absence loci in close proximity. Only experimental determination of causal genes can ultimately rule out this possibility, although we note that GxE in gene expression has previously been shown to arise from variation in the single *IRA2* gene (Smith & Kruglyak, 2008). Future work will elucidate the causal genes in the loci identified here and reveal how they cause GxE.

In conclusion, our results show that the genetic architecture affecting UPS activity is complex, pathway-specific, and subject to a considerable degree of modulation by the environment. Different UPS pathways affect the degradation of distinct substrates, different environments challenge proteostasis in different ways, and genetic variants differ in how they affect a given pathway in a specific environment. The extent of GxE shown here is similar to the amount of GxE seen previously in yeast for transcript abundance and growth in various environments. Thus, GxE is an important factor in yeast complex traits, with important implications for predicting phenotype from genotype.

## Methods

### UPS Activity Reporters

To measure ubiquitin-proteasome system activity, we used tandem fluorescent protein timers (TFTs; (Khmelinskii et al., 2012), a two-color fluorescent reporter system that provides high-throughput measurements of protein turnover. TFTs are linear fusions of two fluorescent proteins. In the most common implementation, the TFT consists of a faster-maturing green fluorescent protein (GFP) and a slower-maturing red fluorescent protein (RFP). Because the two fluorophores fold and emit fluorescence over distinct time scales, the ratio of RFP to GFP changes over time. If the TFT’s degradation rate is more rapid than the RFP’s maturation rate, the TFT ratio can be used to measure the construct’s degradation rate (Khmelinskii et al., 2012; Khmelinskii & Knop, 2014). We used superfolder green fluorescent protein (sfGFP) as the GFP in all TFTs (Supplementary File 1) and mCherry or mRuby as the RFP as indicated below. As in prior studies, we inserted an unstructured 35 amino acid sequence (GSGSREARHKQKIVAPVKQTLNFDLLKLAGDVESN) between the fluorophores to minimize fluorescence resonance energy transfer (M. A. Collins et al., 2022, 2023; Khmelinskii et al., 2012).

To relate the output of our TFT reporters to UPS activity, we utilized degrons that engage distinct UPS pathways to our TFTs. The resulting constructs provide quantitative, high-throughput, pathway-specific readouts of UPS activity in live, single cells that are sensitive to genetic and chemical perturbations that alter UPS activity (M. A. Collins et al., 2022, 2023). We measured the activity of 4 ubiquitin system-dependent UPS pathways. Three of these were N-terminal amino acids that engage distinct branches of UPS N-degron pathways, in which a protein’s N-terminal amino acid functions as a degron (hereafter, an “N-degron”; Varshavsky, 2019). Our N-degron TFT reporters include the Thr N-degron of the Ac/N-degron pathway, the Asn N-degron of the type I Arg/N-degron pathway, and the Phe degron of the type II Arg/N-degron pathway. We also added a non-cleavable ubiquitin moiety (ubiquitin G76V) to the N-terminus of the sfGFP / mCherry TFT. The resulting construct measures the activity of the ubiquitin-fusion domain (UFD) pathway, a UPS pathway involved in protein quality control (Devarajan et al., 2020; Johnson et al., 1995; Theodoraki et al., 2012). We also created two reporters that measure the activity of ubiquitin-system independent UPS pathways. One reporter contains the ubiquitin-independent degron encoded in the N-terminal 80 amino acids of the Rpn4 protein (hereafter, the “Rpn4 degron”). The second reporter contains a linear chain of 4 ubiquitin molecules (hereafter, the “4xUb reporter”). Each of the ubiquitins in 4xUb have the G76V substitution so they cannot be cleaved, and K29/48/63R substitutions to prevent further ubiquitination. Both of the Rpn4 degron and 4xUb reporters are directly bound and degraded by the proteasome without targeting by the ubiquitin system. Their activities thus provide a readout of proteasome activity independent of the ubiquitin system. However, they are each bound by distinct proteasome receptors. Based on the half-lives of the N-degrons (Bachmair et al., 1986; Varshavsky, 2011), we used mRuby, which has a longer half-life than mCherry (168 vs. 40 minutes; Kredel et al., 2009; Shaner et al., 2004) for the Thr N-degron reporter to improve its dynamic range. For all other reporters, mCherry was used as the RFP. The 4xUb and UFD reporters were engineered for this study using procedures described in (M. A. Collins et al., 2022, 2023).

We packaged each reporter into plasmid backbone BFA0190 (M. A. Collins et al., 2022, 2023) containing common sequence elements for reporter integration, selection, and expression. Each reporter contains the *TDH3* promoter to drive strong, constitutive TFT expression, the *ADH1* terminator, codon-optimized sfGFP and mCherry or mRuby, and a KanMX cassette to select for presence of the reporter via resistance to the antibiotic G418. These elements are flanked by sequences homologous to the genomic regions immediately up- and downstream of *LYP1*. Transformation of the reporter containing these flanking sequences results in integration at the *LYP1* locus, which can be selected for using the toxic amino acid analogue thialysine. DNA fragments of the reporters used for transformations were made by PCR amplifying the sequence on the plasmid carrying the reporter sequence (Supplementary File 1). The PCR fragments were purified using Monarch® PCR & DNA Cleanup Kit (5 μg) from New England Biolabs (NEB) Cat#T1030L, or ran on an electrophoresis gel and purified using the Monarch DNA Gel Extraction Kit (NEB) Cat#T1020L, according to the manufacturer’s protocol.

### Yeast Strains and Handling

All experiments used yeast strains derived from two genetically divergent *Saccharomyces cerevisiae* strains. The haploid BY strain (genotype: *MAT**a** his3Δ hoΔ*) is closely related to the S288C laboratory strain. The haploid RM strain (genotype: *MAT! can1Δ::STE2*pr*-SpHIS5 his3Δ::*NatMX *AMN1-* BY *hoΔ::*HphMX *URA3-*FY) is derived from a wild strain that was originally isolated from a California vineyard. To characterize UPS reporters, we also used a previously characterized BY strain lacking the *RPN4* gene (genotype: *MAT**a** his3Δ hoΔ rpn4*Δ::NatMX) (M. A. Collins et al., 2022). All strains used in the present study are listed in Supplementary File 1.

We built strains harboring our UPS activity reporters using the following procedures.

Transformations using the reporter sequence fragments were performed using the Zymo Frozen-EZ Yeast Transformation II™ Kit Cat#T2001 according to the manufacturer’s protocol or the lithium acetate / single-stranded carrier DNA / poly-ethylene glycol (PEG) method (Gietz & Schiestl, 2007) as described in (M. A. Collins et al., 2022, 2023). We verified the presence of the reporters in the desired locus with colony PCR. Eight confirmed transformants for each reporter in each strain were collected as independent biological replicates.

### Yeast mating and segregant populations

To create large, genetically diverse cell populations for genetic mapping, we used a previously-described approach (Albert et al., 2014; Brion et al., 2020; Ehrenreich et al., 2010). We created segregant populations containing each UPS activity reporter using a modified synthetic genetic array methodology (Baryshnikova et al., 2010; Kuzmin et al., 2016). Briefly, BY strains (*MAT**a***) containing each reporter were mixed with a wild-type RM strain (*MAT!*, without a reporter: YFA0039) on solid YPD medium (all media are described in Supplementary File 1) and grown overnight at 30°C. This was done independently for two biological replicates based on two different colony PCR-confirmed transformants for each reporter. Diploids from the mating were selected on YPD plates containing G418 and CloNAT to select for the UPS reporter in the BY strain and *his3Δ::NatMX* in the RM strain, respectively. Five mL of liquid YPD was inoculated with the diploids and grown overnight to saturation at 30°C with rolling. The diploids were then spun down in 15 mL tubes in a tabletop centrifuge at 3000 rpm for two minutes. The cell pellet was resuspended in five mL of sporulation medium and transferred to glass tubes and incubated at room temperature for 10 days on a turning wheel. We evaluated the extent of sporulation in each culture using brightfield microscopy. When the cultures reached approximately 80% sporulation, we harvested the spores. To separate the spores from their asci, we spun the spores for 1.5 minutes at 5000 rpm in a tabletop centrifuge, discarded the supernatant, resuspended in water with 1 mg / mL Zymolyase lytic enzyme (United States Biological, Salem, MA, USA), and incubated for two hours, vortexing every half hour. We again washed the cells and plated them onto solid haploid selection medium with G418 and thialysine. We used this medium to select for recombined haploid cells (“segregants”) that contain the reporter via G418, the *MAT**a*** mating type locus via the *Schizosaccharomyces pombe HIS5* gene under the control of the *STE2* promoter (which is only active in *MAT**a*** cells), and replacement of the *LYP1* gene by the reporter via resistance to thialysine. We grew the resulting segregant populations for two days on haploid selection plates at 30°C, harvested the cells from the plates, and stored each population as a separate glycerol stock. We saved two biological replicates for each reporter as separate, individual stocks.

### Environments

To characterize how genetic influences on the UPS are shaped by environmental factors, we measured UPS activity in eight distinct media formulations, which we term “environments.” As a baseline environment, we used synthetic complete (SC), a nutrient-rich medium with glucose, nitrogen, and amino acids. G418 (200 mg/mL) was added to all environments to maintain the reporter sequence in the genome. We compared UPS activity in SC (SC -His -Lys + YNB + 0.1% MSG + 2% glucose) to that in seven different environments: low glucose (SC -His -Lys + YNB + 0.1% MSG + 0.025% glucose), low nitrogen (YNB + 2% glucose), yeast nitrogen base (YNB + 0.1% MSG + 2% glucose), SC + 4NQO (4NQO; 2 µg/mL), SC +L-azetidine-2-carboxylic acid (AZC; 4mM), SC + bortezomib (BTZ; 40µM), and SC + lithium acetate (LiAc; 20 mM). The media and chemical formulations and concentrations used for all experiments are described in Supplementary File 1.

### Growth and environmental exposures prior to flow cytometry

Eight biological replicates of each of the BY, RM, and *rpn4*Δ strains containing the reporters (Supplementary File 1) were grown overnight to saturation in SC medium. From a common saturated sample for each replicate, 4 µL was used to inoculate 400 µL media in 96 well plates for each environment. G418 (200 mg / mL) was added to all media except for the negative controls. The cultures were incubated at 30°C on a MixMate (Eppendorf, Hamburg, Germany) at 1100 rpm and incubation times in each medium were determined based on a combination of previous literature (Burgis & Samson, 2007; Laporte et al., 2008; J. Li et al., 2019; Marshall et al., 2016; Waite et al., 2016; Work & Brandman, 2021) and preliminary growth rate measurements in a plate reader to ensure that all cultures had similar optical density (O.D.) for flow cytometry and FACS. If cultures showed no growth in a given environment in preliminary experiments, we grew the cells in SC before exposing the cells to that environment (see below for details). For SC, LiAc, and YNB conditions, 400 µL of medium was inoculated with 4 µL of the overnight growth and incubated for 3 hours prior to flow cytometry. For the BTZ samples, 400 µL of SC + BTZ medium was inoculated with 4 µL of the overnight growth culture, incubated for 4 hours until flow cytometry measurements.

For samples that were to be exposed to low nitrogen, low glucose, 4NQO, and AZC, 4 µL of the overnight culture was added to 400 µL of SC and grown for 3 hours. After those 3 hours, samples to be exposed to low nitrogen and low glucose were spun down for 5 mins at 3000 rpm in 96 well plates on a tabletop centrifuge. The SC medium was replaced with either the low nitrogen or low glucose media and incubated for 24 hours until measured via flow cytometry. If the low nitrogen or low glucose samples were too dense for flow cytometry, they were diluted in a 1:3 ratio with the same media type. For the 4NQO samples, after the 3 hours of growth in SC, the cells were spun down for 5 mins at 3000 rpm in 96 well plates on a tabletop centrifuge. The SC media was replaced with SC + 4NQO medium and incubated for one hour until measured via flow cytometry. For the AZC samples, after the 3 hours of growth in SC, 3.2 µL of AZC stock was added to each well. After 5 hours of incubation, samples were measured via flow cytometry.

For all flow cytometry and cell sorting experiments, we used measurements of the wild-type BY strain (YFA0040) without a TFT reporter grown in SC only to determine background fluorescence levels. When the GFP signal of a TFT did not exceed that of YFA0040 in a given environment, we concluded that the UPS activity measured by the associated reporter could not be accurately measured in that environment. A 400 µL sample of SC -lys was inoculated with 4 µL of the overnight growth of YFA0040 and incubated for 3 hours prior to flow cytometry. Four of the UFD *rpn4Δ* replicates did not produce GFP fluorescence above the negative control (BY without a reporter). We therefore excluded those four samples without detectable GFP from analysis.

### Flow cytometry

All flow cytometry experiments were performed on the BD FACSymphony^TM^ A3 flow cytometer (BD, Franklin Lakes, NJ, USA) at the University of Minnesota University Flow Cytometry Resource. The cytometer is equipped with a 20 mW 488 nm laser with a 488 / 10 filter to measure forward scatter (FSC) and side scatter (SSC) and a 525 / 50 filter to measure GFP fluorescence, and a 40 mW 561 nm laser with a 610 / 20 filter to measure RFP fluorescence. The voltages for each parameter are listed in Supplementary File 1. We altered voltages for samples containing AZC compared to the other conditions so that GFP did not saturate at the upper end of detection, as GFP fluorescence was higher in those samples, as follows: FSC, 450; GFP, 400; RFP, 600.

We used flow cytometry to characterize the two new reporters, 4xUb and UFD, in SC medium. We recorded data for 10,000 cells from each of the eight biological replicates of each strain (BY, RM, *rpn4*Δ) containing these reporters (Strains listed in Supplementary File 1). We used flow cytometry to test for genotype-by-environment interactions (GxE) between the BY and RM strains for all six reporters. For these experiments, we recorded data for 20,000 cells from the eight biological replicates of BY and RM strains containing the six reporters in each of eight environments as described above.

### Analysis of flow cytometry data

Analyses were conducted using code adapted from (M. A. Collins et al., 2022). Briefly, we analyzed flow cytometry data using the R (R Foundation for Statistical Computing, Vienna Austria) package flowCore (Hahne et al., 2009). We first filtered each replicate to include only cells within 10% + / - the FSC median (proxy for cell size). This removed cellular debris, aggregates of multiple cells, and restricted our analyses to cells of the same approximate size. The low nitrogen samples showed two FSC peaks, perhaps due to incomplete budding of daughter cells. In order to only analyze single cells, we selected the smaller of the two peaks for analysis of the low nitrogen samples. The median FSC values of the smaller low nitrogen peaks were similar in size to the FSC medians of all other samples.

As in prior studies (Brion et al., 2020; M. A. Collins et al., 2023), we observed that the -log2(RFP / GFP) ratio changed over time within some replicates of the same strain, reporter, and environment. To correct for this, we used the residuals of a loess regression of the -log2(RFP / GFP) ratio, as in (Brion et al., 2020; M. A. Collins et al., 2023). We refer to the time-corrected -log2(RFP / GFP) ratio as “UPS activity” throughout. P-values for differences in UPS activity between the BY, RM, and *rpn4*Δ strains were calculated using a two-tailed T-test.

The GxE effect of each reporter / environment combination was determined using the following linear mixed model:

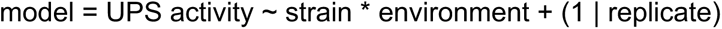

Here, the random effect term “(1 | replicate)” accounts for inter-individual variation among independent biological replicates. We conducted pairwise analyses, in which we compared the effect of each of the seven environments to the baseline SC environment, for one reporter at a time. If one of the eight replicates for a given reporter in a given environment had a median GFP level below that of the negative control (a BY strain with no reporter), all eight replicates for that reporter / environment combination were excluded. This exclusion applied to the 4xUb and UFD reporters in low nitrogen, resulting in 40 tests for GxE. Statistical significance of main effects of strain, environment, and the GxE interaction term between strain and environment was assessed using an ANOVA. We used Bonferroni-corrected p-value thresholds of 0.05 / 40 = 0.00125. The magnitude of the environmental effect on UPS activity (Fig. 2A & B) was determined by subtracting the median UPS activity in the given environment from the median UPS activity in SC. To describe GxE patterns revealed by the linear models, we also used T-tests comparing UPS activity for each strain between SC and another condition. For the T-tests, we used Bonferroni-corrected p-value thresholds of 0.05 / 80 = 0.000625.

### Growth and environmental exposures prior to fluorescence-activated cell sorting (FACS)

All incubations were performed at 30°C in glass test tubes with rolling. Two independent biological replicate segregant stocks were thawed and used to inoculate our baseline medium (SC -his -lys + G418) and grown overnight to saturation. This common culture was used to inoculate media for each environment as follows.

For SC, LiAc, and YNB, 4.2 mL of media was inoculated with 800 µL of the overnight growth culture and incubated for 3 hours until FACS. For the BTZ samples, 4.5 mL of SC + BTZ medium was inoculated with 500 µL of the overnight growth culture and incubated for 4 hours until FACS.

For samples that were exposed to low nitrogen, low glucose, 4NQO, and AZC, 500 µL of the overnight culture was added to 4.5 mL of SC and grown for 3 hours. Samples exposed to low nitrogen and low glucose were then spun down for 2 mins at 3000 rpm in 15 mL tubes on a tabletop centrifuge. The SC medium was replaced with 5 mL of the low nitrogen or low glucose media and incubated for 24 hours until cell sorting. For the 4NQO samples, after the 3 hours of growth in SC, 0.4 µL of 4NQO stock was added to the cultures. The 4NQO cultures were immediately vortexed and incubated for one hour until cell sorting. For the AZC samples, after the 3 hours of growth in SC, 40 µL of AZC stock was added to each sample and vortexed. AZC samples were incubated for 5 hours until FACS. These volumes and incubation times led to the samples being at approximately the same O.D. when sorted.

A BY strain without a UPS reporter (YFA0040) was grown overnight to saturation in SC -lys to be used as a negative fluorescence control. A 4.5 mL volume of SC -lys was inoculated with 500 µL of the overnight growth and incubated for 3 hours until FACS.

### FACS

We used FACS to isolate phenotypically extreme cell populations as part of a bulk segregant analysis genetic mapping approach (Albert et al., 2014; Brion et al., 2020). All cell sorting was performed on a FACSAria II cell sorter (BD) by the University of Minnesota Flow Cytometry Resource. To remove doublets from each sample, we used plots of SSC height by width and FSC height by width. We kept cells within the peak of FSC area + / - 7.5%, which maintained our primary haploid cell population and excluded cellular debris and aggregates (M. A. Collins et al., 2022, 2023). We restricted our sorts to populations of cells with GFP fluorescence above that of the negative control BY strain YFA0040, which does not express any fluorescent proteins. We collected populations of cells from the 2% high and low tails of the RFP / GFP ratio distribution. We aimed to collect pools of 20,000 cells for each of these populations. When cultures did not contain a sufficient amount of GFP-positive cells to collect 20,000 cells, we collected fewer cells. We empirically determined reporter / environment combinations for which the cell pools did not grow well after sorting in a preliminary experiment. We therefore collected more cells for those samples when we sorted the cells used in the downstream analyses, up to 100,000 cells. The final numbers of cells collected for both replicates of the high and low pools for each reporter and each environment are reported in Supplementary File 1. The segregants with the 4xUb reporter in the low nitrogen environment did not produce fluorescence above the negative control, and therefore no cells were collected for that combination, and all downstream analyses do not include 4xUb in low nitrogen.

Cells were collected into sterile 1.5 mL polypropylene tubes with 1 mL of SC -his -lys medium and grown at 30°C with rolling at least 26 hours or until saturation. We added 1 mL of each culture to a 96-well plate with 600 µL of 40% glycerol and stored at -80°C for subsequent genomic DNA extraction.

### DNA extraction and library preparation

We isolated genomic DNA from thawed glycerol stocks of the sorted segregant pools for whole-genome sequencing. We centrifuged 800 µL of each pool at 3700 rpm for 10 minutes to pellet the cells and discarded the supernatant. To digest cell walls we resuspended the cells in 800 µL of 1 M sorbitol, 0.1 M EDTA, 14.3 mM *β*-mercaptoethanol, and 500 U of Zymolyase lytic enzyme (United States Biological) and incubated for 2 hours at 37°C on a MixMate at 1100 rpm. We re-pelleted the cells, removed the supernatant, and extracted DNA from the cells using the Quick-DNA 96 Plus kit (Zymo Research, Irvine, CA, USA), according to the manufacturer’s instructions, including an overnight protein digestion in 20 mg / mL of proteinase K solution. We eluted the DNA using 35 µL of DNA Elution Buffer (10 mM Tris-HCl [pH 8.5], 0.1 mM EDTA) and determined DNA concentration on a Synergy H1 plate reader (BioTek Instruments, Winooski, VT, USA) in 96-well plates using the Qubit dsDNA BR assay kit (Thermo Fisher Scientific, Waltham, MA, USA).

We prepared the genomic DNA for short-read whole-genome sequencing on the Illumina NovaSeq platform using a previously established approach (Albert et al., 2014; Brion et al., 2020; M. A. Collins et al., 2022, 2023). We used the Nextera DNA library kit (Illumina, San Diego, CA, USA) in 96 well plates according to the manufacturer’s instructions, except that we used a 1:20 dilution of the Tagment DNA enzyme in Tagment DNA buffer. After library generation, we quantified the DNA concentration of each sample using the Qubit dsDNA BR assay kit (Thermo Fisher Scientific). For each 96 well plate, 10 µL of each sample was pooled and 1 mL of that pool was run in a large well on a 2% agarose gel. We extracted and purified the DNA in the 400 to 600 bp region using the Monarch Gel Extraction Kit (NEB) according to the manufacturer’s instructions. The DNA from each of the gel extractions was quantified and pooled in equimolar amounts, and submitted for sequencing.

The University of Minnesota Genomics Center (UMGC) staff performed quality control assays as described in (M. A. Collins et al., 2022, 2023) on the pooled library before sequencing. Briefly, library concentration was determined using PicoGreen dsDNA quantification reagent (Thermo Fisher Scientific), library size was determined using the Tapestation electrophoresis system (Agilent Technologies, Santa Clara, CA, USA), and library functionality was determined using the KAPA DNA Library Quantification kit (Roche, Penzberg, Germany). The submitted pooled library passed each quality control assay. The pooled library was sequenced on the Illumina NovaSeq with 150 bp paired end reads. Of the samples used for analysis, the median reads produced per sample was 1,692,391, with the minimum being 304,256 reads and the maximum being 5,874,865 reads. UMGC performed sequence data de-multiplexing.

### Genetic mapping

QTLs were determined using an established approach for bulk segregant analysis (Albert et al., 2014; Brion et al., 2020; Ehrenreich et al., 2010; Michelmore et al., 1991). Code from (M. A. Collins et al., 2022, 2023) was used to calculate allele frequencies via the following pipeline. From our whole-genome sequencing reads, we aligned reads to the *S. cerevisiae* reference genome (version sacCer3) using the BWA "mem" command (H. Li & Durbin, 2009) and retained alignments with a mapping quality score above 30. Using samtools (H. Li et al., 2009), we retained uniquely aligned reads and removed PCR duplicates (command: "samtools markdup -S"). VCF files with allelic read counts at 18,871 high-confidence, reliable SNPs (Bloom et al., 2013; Ehrenreich et al., 2010) were produced using the command: samtools mpileup - vu -t INFO / AD -l.

We used adapted code from (M. A. Collins et al., 2022, 2023) to calculate allele counts from the VCF files. Briefly, we excluded variants with allele frequencies lower than 0.1 or higher than 0.9 as in (Albert et al., 2014; Brion et al., 2020). We used MULTIPOOL (Edwards & Gifford, 2012) to estimate logarithm of the odds (LOD) scores comparing a model in which the high and low degradation activity pools come from one population to a model in which these pools come from two different populations with different allele frequencies. As in (M. A. Collins et al., 2022, 2023), we used the following MULTIPOOL settings: bp per centiMorgan = 2,200, bin size = 100 bp, effective pool size = 1,000. We called QTLs as loci with a LOD ≥ 4.5. Previous work has shown that this threshold produces a 0.5% false discovery rate for genetic mapping by bulk segregant analysis using TFT reporters (M. A. Collins et al., 2022). Confidence intervals (CI) for each significant QTL were determined using MULTIPOOL and defined as a 2-LOD drop from the position in the QTL interval with the highest LOD score, which we defined as the QTL peak position. We calculated the RM allele frequency difference (ΔAF) between the high and low degradation activity pools using a smoothed allele frequency via a loess regression to account for random counting noise at individual sequence variants. In our scheme, a positive ΔAF indicates that the RM allele of a QTL is associated with higher UPS activity. The loess smoothed values were used for plotting and determining QTL effect sizes.

QTLs were called separately for each biological replicate. To determine QTLs that were detected in both biological replicates, we used a previously described approach (M. A. Collins et al., 2022, 2023).

QTLs present in both replicates were defined as QTLs on the same chromosome with peaks within 100 kb and with the same effect direction (ΔAF sign). From our set of 694 QTLs across replicates, 416 QTLs (60%) were present in both replicates, with the remaining 278 QTLs detected in only one replicate. For QTLs present in both replicates, the left and right CI positions, peak position, LOD, and ΔAF were averaged and used for downstream analyses. For QTLs found in only one replicate, the left and right CI positions, peak position, LOD, and ΔAF were used without alteration.

The high and low populations of the second biological replicate of Phe in 4NQO did not have sufficient sequencing coverage to call QTLs. To replace these populations, we used additional populations we had collected during FACS of Phe in 4NQO from the first biological replicate. These additional populations consisted of cells with RFP / GFP ratios in the 3% to 5% area of the distribution. Therefore, both replicates used in data analysis of Phe in 4NQO came from the same original segregant pool.

### Comparison of QTLs from previous studies

QTLs for the Asn, Phe, Rpn4, and Thr reporters had been mapped previously in the same standard SC medium (M. A. Collins et al., 2022, 2023). For these four reporters, we analyzed 39 QTLs that were present in both replicates in SC in the present study and asked whether they were also present in at least one of the two replicates from the previous studies. If a QTL from the present study had a QTL whose peak was within 100 kb from (M. A. Collins et al., 2022, 2023) in at least one replicate, we determined that this QTL was present in both studies. All QTLs present in both studies had the same sign of ΔAF. A two-sample T-test was used to determine if there was a significant difference of LOD scores and absolute ΔAF between QTLs that were present in both studies compared to those found only in the present study.

### GxE in the QTLs

To determine GxE at individual QTLs, we compared loci between SC and each additional, distinct environment for each reporter. GxE at individual QTLs was classified as either 1) presence / absence or 2) sign change. Presence / absence GxE QTLs were defined as loci detected in both replicates of one environment, but where no QTL peak was found within 100 kb in a separate environment. Because a QTL might be absent due to insufficient power, we only considered QTLs that were present in both replicates of one environment (and therefore are likely to be relatively strong in that environment) and absent in both replicates of the other environment. Sign change GxE QTLs were defined as QTLs that were present in SC and a given environment but had ΔAF of a different sign. We considered QTLs to be present in both environments when their peak position occurred within 100 kb. We included QTLs in the sign change pairs that were found in both replicates of one environment and in one or both replicates of the other environment.

If a pair of QTLs whose peaks were within 100 kb between SC and a given environment had the same sign of ΔAF, the pair was considered not to exhibit GxE. We included pairs of QTLs where a QTL was present in both replicates of one environment and present in either one or both replicates of the other environment.

## Acknowledgements

We thank members of the Albert laboratory for experimental guidance and feedback on the manuscript. The authors acknowledge the University of Minnesota Genomics Center (UMGC) for providing resources that contributed to the research results reported in this paper. We thank Rashi Arora and the University of Minnesota Flow Cytometry Resource staff for technical assistance with flow cytometry and FACS.

## Funding

This work was supported by NIH grant R35GM124676 to FWA.

---

**Supplementary Fig. 1:**
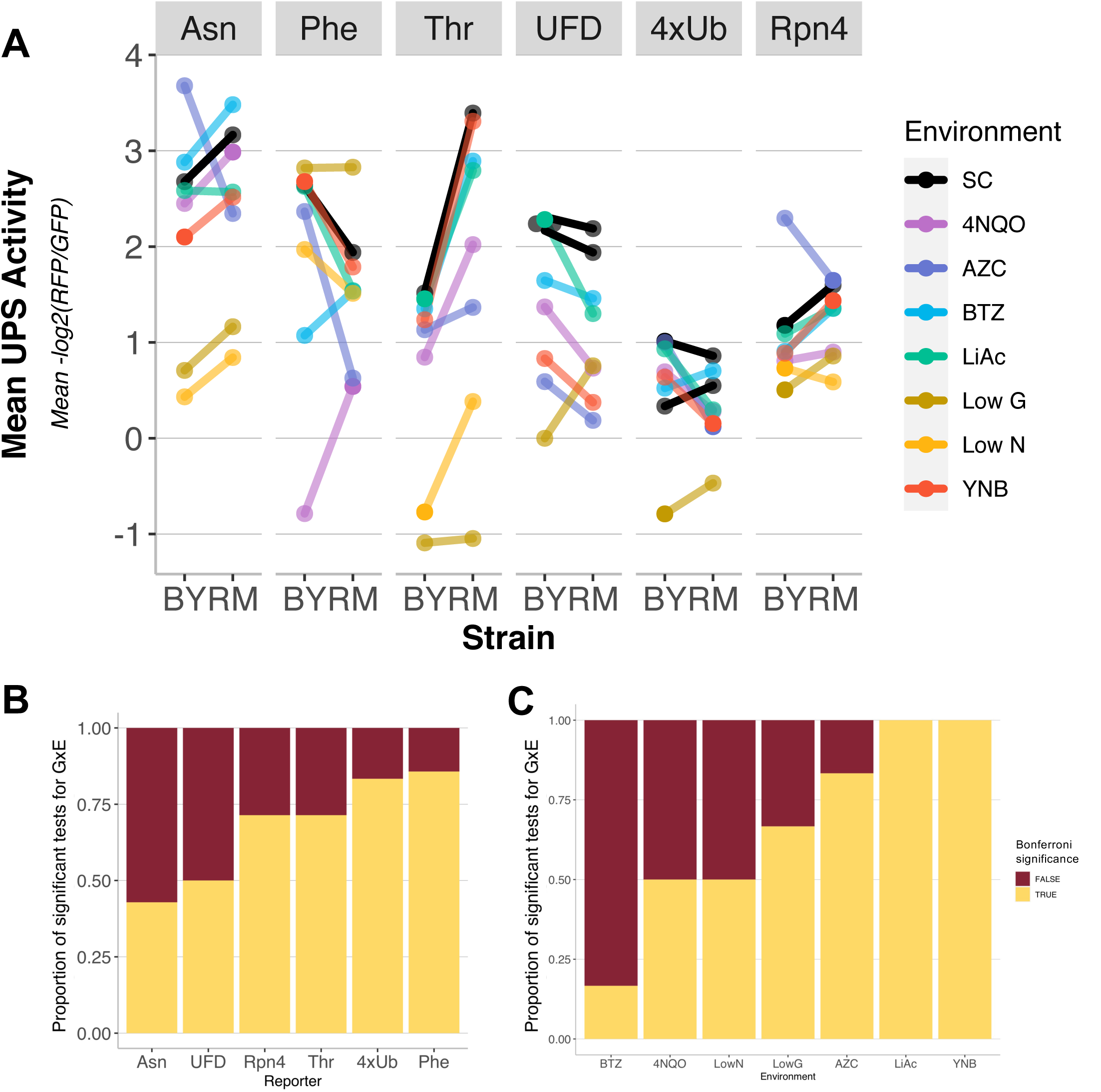
**A.** Mean UPS activity across eight replicates for each environment and reporter. **B.** Proportion of tests with significant (gold) and non-significant (maroon) GxE, by reporter. **C.** Data as in C, but rearranged by environment.

**Supplementary Fig. 2:**
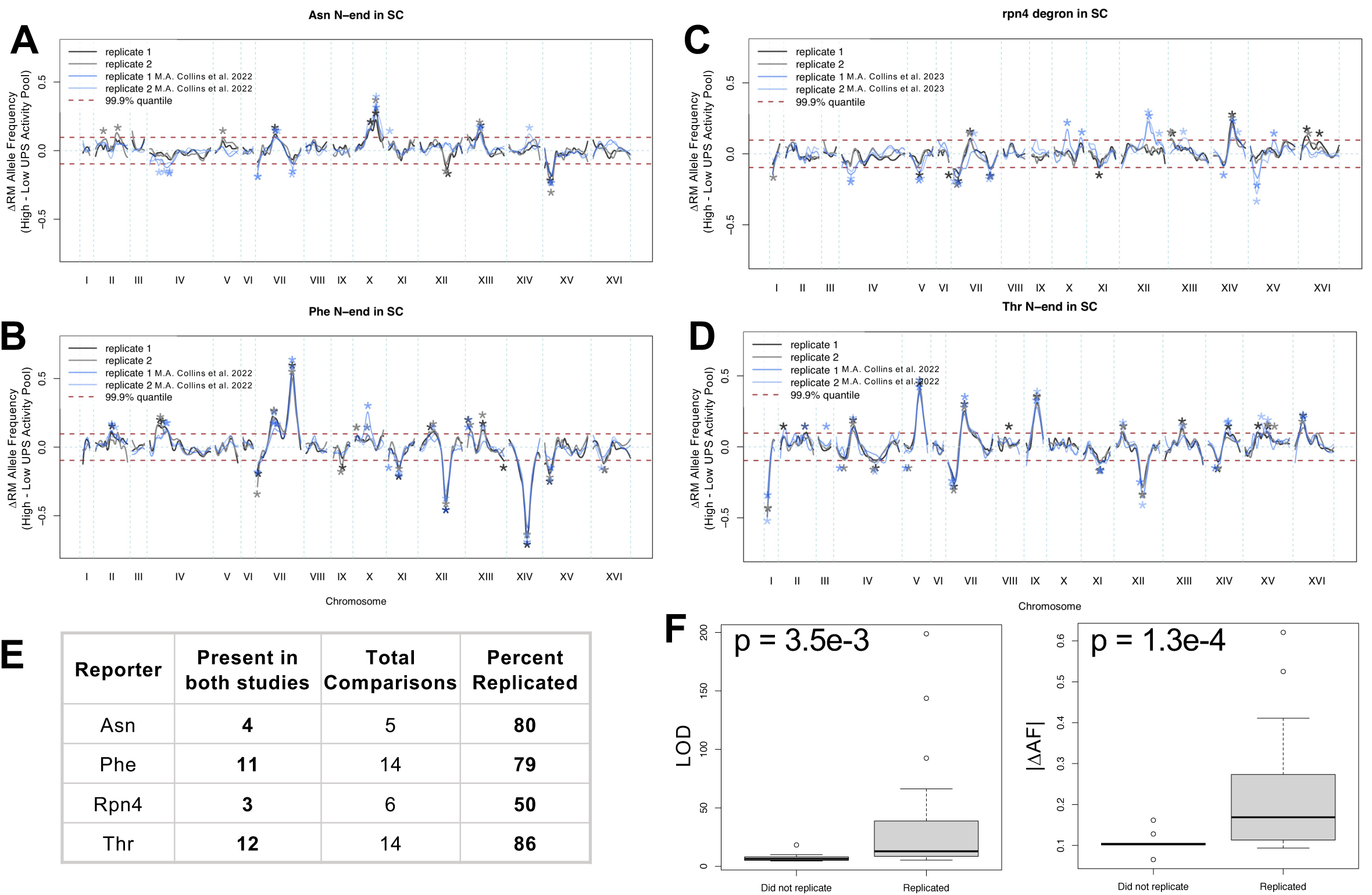
QTL reproducibility. **A-D.** QTL traces for the four reporters measured in SC in this study and M. A. Collins et al., 2022, 2023. **E.** Table summarizing the number of QTLs that replicated between studies. **F.** QTLs that did not replicate between studies had significantly lower LOD scores absolute delta allele frequencies.

**Supplementary Fig. 3:**
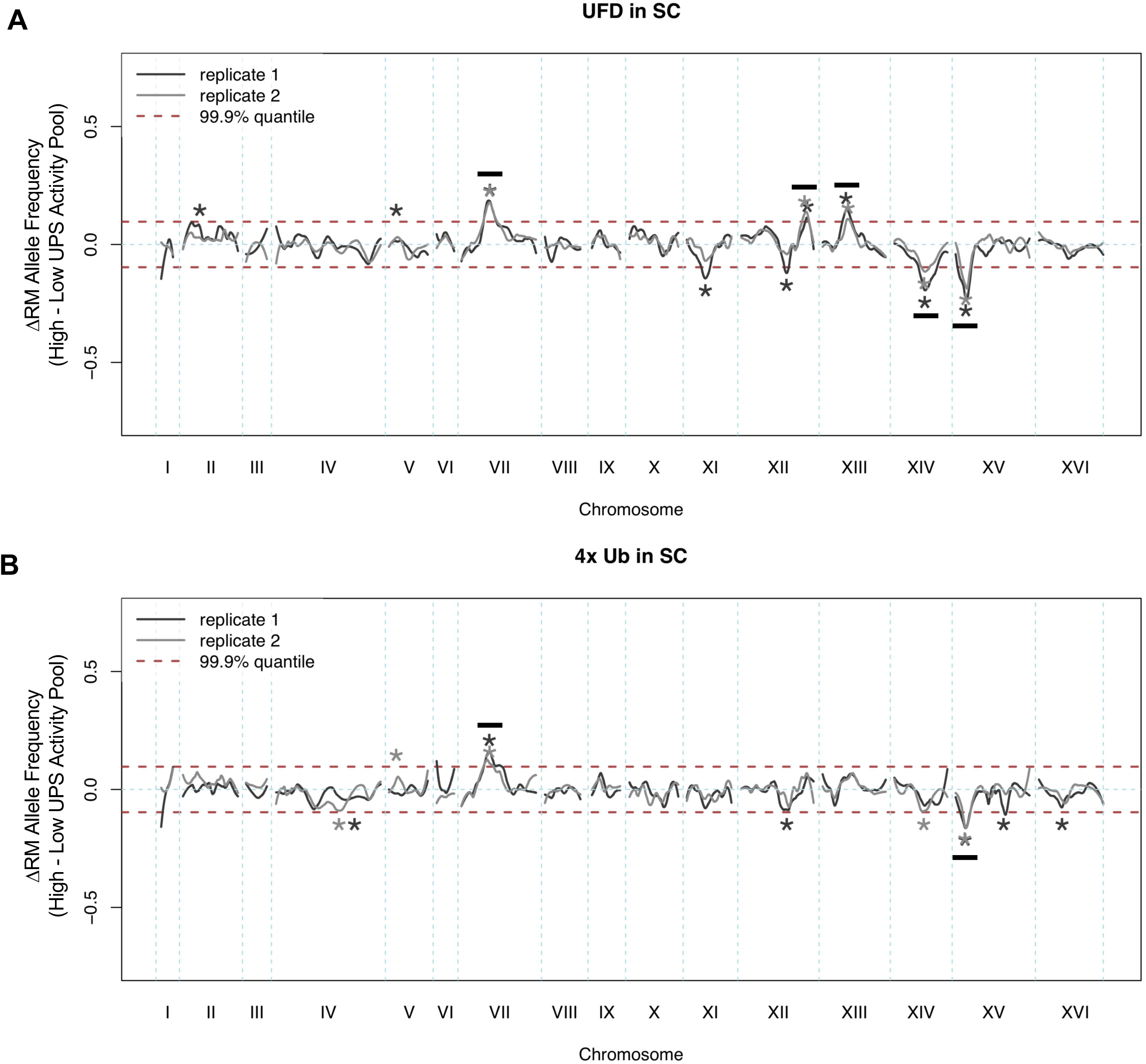
QTLs for 4xUb and UFD in SC. The plots show the loess-smoothed allele frequency difference between the high and low UPS activity pools across the genome for each of two independent biological replicates. Asterisks denote QTLs, defined by allele frequency differences that exceed an empirically-derived LOD score significance threshold in the given replicate. Horizontal black lines indicate QTLs that were present in both replicates. The dashed red horizontal lines denote an empirically-derived 99.9% quantile of the allele frequency difference. **A.** UFD reporter in SC. Five QTLs were present in both replicates. **B.** 4xUb reporter in SC. Two QTLs were present in both replicates.

**Supplementary Fig. 4:**
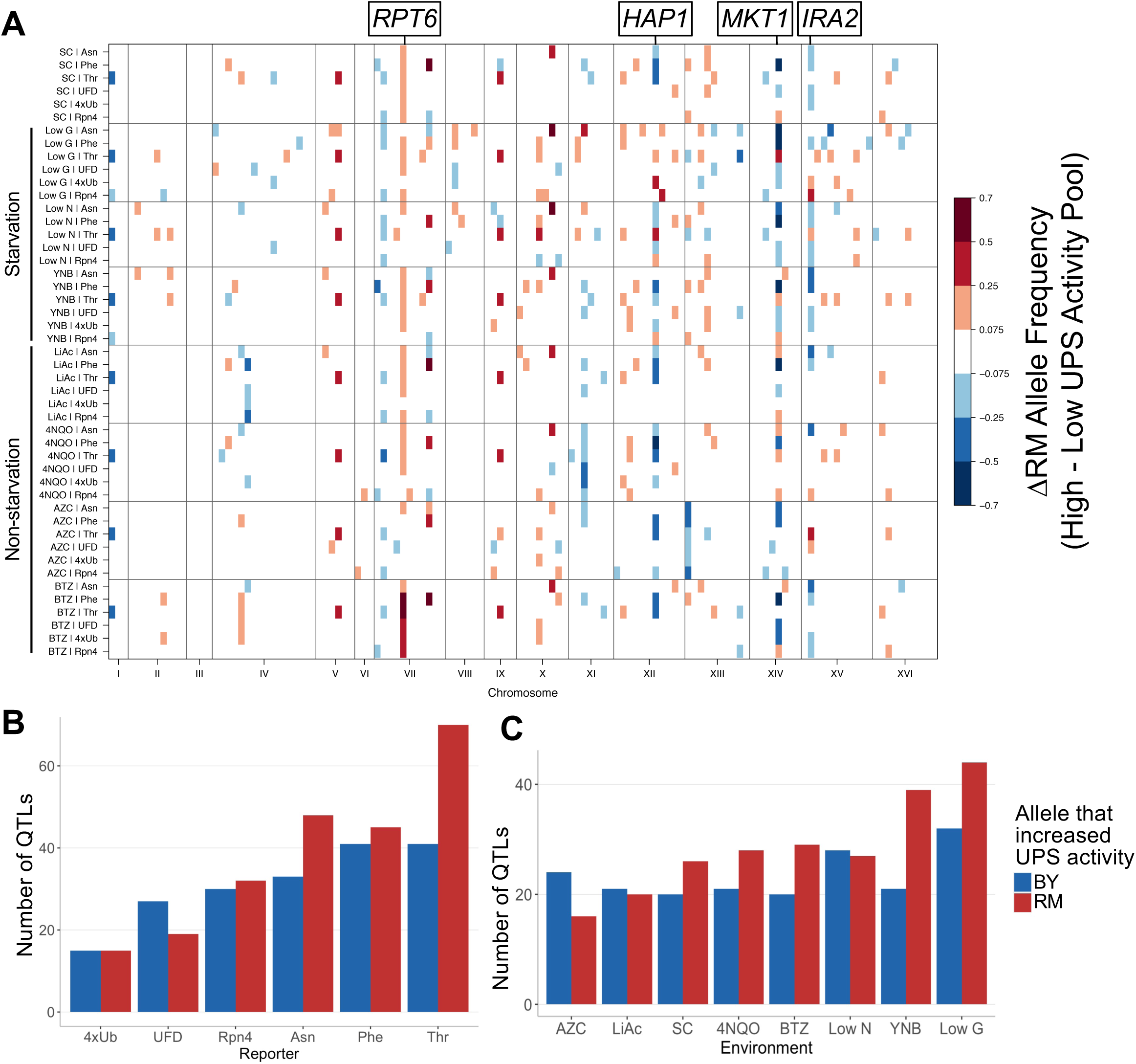
UPS activity QTLs across environments and reporters. **A.** QTLs for each reporter / environment combination. Data as in Fig. 3A, but reorganized according to environment. Colored blocks denote genome bins that contain QTLs detected in each of two independent biological replicates, colored according to the direction and magnitude of the effect size, expressed as the RM allele frequency difference between high and low UPS activity pools. Candidate causal genes discussed in the text are indicated. No data was collected for 4xUb in low nitrogen. **B.** Barplot showing the number of times the BY or RM allele increased degradation in the 416 QTLs, by reporter. **C.** Data as in B, but reorganized by environment.

**Supplementary Fig. 5:**
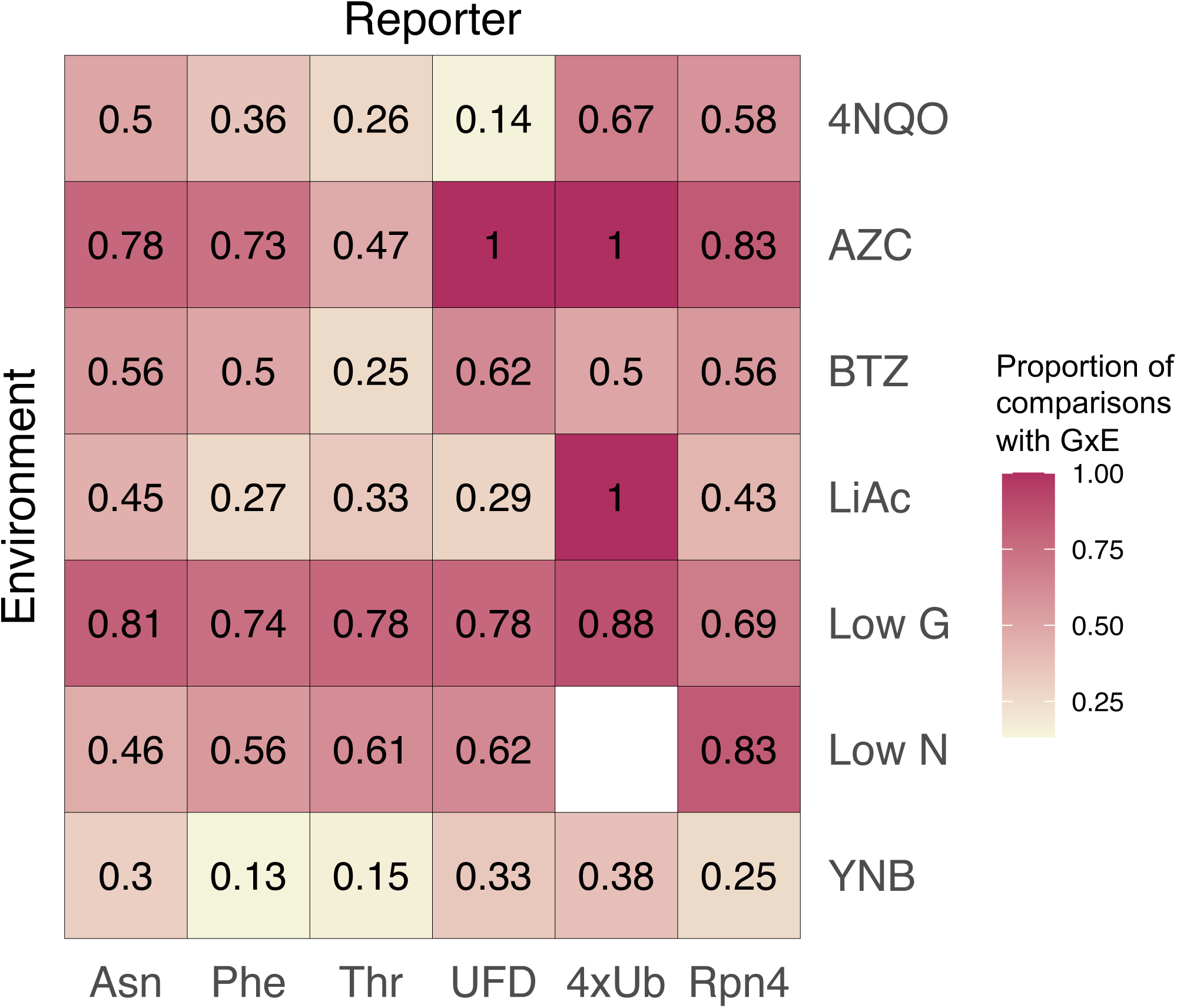
A heatmap showing the proportion of locus comparisons that showed GxE out of all comparisons, for combinations of reporters and environments. No data was collected for 4xUb in low nitrogen.

**Supplementary Fig. 6:**
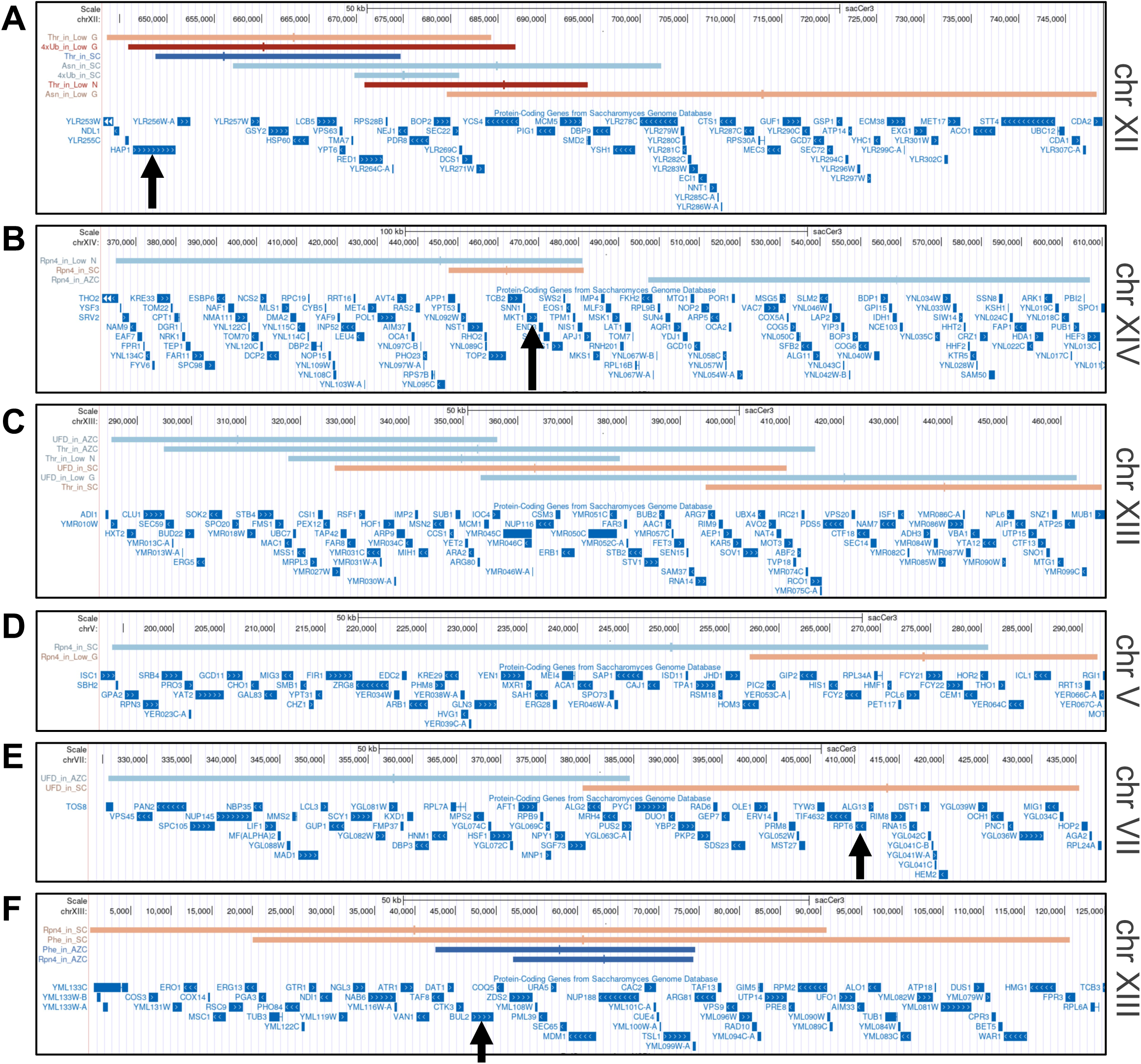
Locations of QTLs within sign change pairs: Locus plots of the 17 cases of sign change GxE (along with Fig. 5C). The confidence interval (horizontal bars) and peak position (short vertical lines) of the 29 unique QTLs involved in sign changes are shown along the *S. cerevisiae* genome (sacCer3) using UCSC Genome Browser (Nassar et al., 2023). For the QTLs, colors indicate direction and strength of effect as in Fig. 3A. Candidate causal genes are denoted with black arrows.

## Notes

### Competing Interest Statement

The authors have declared no competing interest.

## References

Albert, F. W., Bloom, J. S., Siegel, J., Day, L., & Kruglyak, L. (2018). Genetics of trans-regulatory variation in gene expression. eLife, 7, e35471. 10.7554/eLife.35471

Albert, F. W., Treusch, S., Shockley, A. H., Bloom, J. S., & Kruglyak, L. (2014). Genetics of single-cell protein abundance variation in large yeast populations. Nature, 506(7489), 494–497. 10.1038/nature12904

Bachmair, A., Finley, D., & Varshavsky, A. (1986). In vivo half-life of a protein is a function of its amino-terminal residue. *Science (New York*, N.Y.), 234(4773), 179–186. 10.1126/science.3018930

Bajorek, M., Finley, D., & Glickman, M. H. (2003). Proteasome disassembly and downregulation is correlated with viability during stationary phase. Current Biology: CB, 13(13), 1140–1144. 10.1016/s0960-9822(03)00417-2

Ballinger, M. A., Mack, K. L., Durkin, S. M., Riddell, E. A., & Nachman, M. W. (2023). Environmentally robust *cis* -regulatory changes underlie rapid climatic adaptation. Proceedings of the National Academy of Sciences, 120(39), e2214614120. 10.1073/pnas.2214614120

Baryshnikova, A., Costanzo, M., Dixon, S., Vizeacoumar, F. J., Myers, C. L., Andrews, B., & Boone, C. (2010). Synthetic genetic array (SGA) analysis in Saccharomyces cerevisiae and Schizosaccharomyces pombe. Methods in Enzymology, 470, 145–179. 10.1016/S0076-6879(10)70007-0

Bett, J. S. (2016). Proteostasis regulation by the ubiquitin system. Essays in Biochemistry, 60(2), 143–151. 10.1042/EBC20160001

Bickel, H., Gerrard, J., & Hickmans, E. M. (1953). Influence of phenylalanine intake on phenylketonuria. Lancet (London, England), 265(6790), 812–813. 10.1016/s0140-6736(53)90473-5

Bloom, J. S., Boocock, J., Treusch, S., Sadhu, M. J., Day, L., Oates-Barker, H., & Kruglyak, L. (2019). Rare variants contribute disproportionately to quantitative trait variation in yeast. eLife, *8*, e49212. 10.7554/eLife.49212

Bloom, J. S., Ehrenreich, I. M., Loo, W. T., Lite, T.-L. V., & Kruglyak, L. (2013). Finding the sources of missing heritability in a yeast cross. Nature, 494(7436), 234–237. 10.1038/nature11867

Boye, C., Nirmalan, S., Ranjbaran, A., & Luca, F. (2024). Genotype × environment interactions in gene regulation and complex traits. Nature Genetics, 56(6), 1057–1068. 10.1038/s41588-024-01776-w

Boyle, E. A., Li, Y. I., & Pritchard, J. K. (2017). An Expanded View of Complex Traits: From Polygenic to Omnigenic. Cell, 169(7), 1177–1186. 10.1016/j.cell.2017.05.038

Brem, R. B., Yvert, G., Clinton, R., & Kruglyak, L. (2002). Genetic Dissection of Transcriptional Regulation in Budding Yeast. Science, 296(5568), 752–755. 10.1126/science.1069516

Brion, C., Lutz, S. M., & Albert, F. W. (2020). Simultaneous quantification of mRNA and protein in single cells reveals post-transcriptional effects of genetic variation. eLife, 9, e60645. 10.7554/eLife.60645

Burgis, N. E., & Samson, L. D. (2007). The Protein Degradation Response of *Saccharomyces cerevisiae* to Classical DNA-Damaging Agents. Chemical Research in Toxicology, 20(12), 1843–1853. 10.1021/tx700126e

Chen, S.-A. A., Kern, A. F., Ang, R. M. L., Xie, Y., & Fraser, H. B. (2023). Gene-by-environment interactions are pervasive among natural genetic variants. Cell Genomics, 3(4). 10.1016/j.xgen.2023.100273

Collins, G. A., & Goldberg, A. L. (2017). The Logic of the 26S Proteasome. Cell, 169(5), 792–806. 10.1016/j.cell.2017.04.023

Collins, M. A., Avery, R., & Albert, F. W. (2023). Substrate-specific effects of natural genetic variation on proteasome activity. PLOS Genetics, 19(5), e1010734. 10.1371/journal.pgen.1010734

Collins, M. A., Mekonnen, G., & Albert, F. W. (2022). Variation in ubiquitin system genes creates substrate- specific effects on proteasomal protein degradation. eLife, 11, e79570. 10.7554/eLife.79570

Coux, O., Tanaka, K., & Goldberg, A. L. (1996). Structure and functions of the 20S and 26S proteasomes. Annual Review of Biochemistry, 65, 801–847. 10.1146/annurev.bi.65.070196.004101

Cubillos, F. A., Stegle, O., Grondin, C., Canut, M., Tisné, S., Gy, I., & Loudet, O. (2014). Extensive *cis* - Regulatory Variation Robust to Environmental Perturbation in *Arabidopsis*. The Plant Cell, 26(11), 4298–4310. 10.1105/tpc.114.130310

Dantuma, N. P., & Bott, L. C. (2014). The ubiquitin-proteasome system in neurodegenerative diseases: Precipitating factor, yet part of the solution. Frontiers in Molecular Neuroscience, 7, 70. 10.3389/fnmol.2014.00070

Devarajan, S., Meurer, M., van Roermund, C. W. T., Chen, X., Hettema, E. H., Kemp, S., Knop, M., & Williams, C. (2020). Proteasome-dependent protein quality control of the peroxisomal membrane protein Pxa1p. Biochimica Et Biophysica Acta. Biomembranes, 1862(9), 183342. 10.1016/j.bbamem.2020.183342

Dimitrov, L. N., Brem, R. B., Kruglyak, L., & Gottschling, D. E. (2009). Polymorphisms in multiple genes contribute to the spontaneous mitochondrial genome instability of Saccharomyces cerevisiae S288C strains. Genetics, 183(1), 365–383. 10.1534/genetics.109.104497

Edwards, M. D., & Gifford, D. K. (2012). High-resolution genetic mapping with pooled sequencing. BMC Bioinformatics, 13 Suppl 6(Suppl 6), S8. 10.1186/1471-2105-13-S6-S8

Ehrenreich, I. M., Torabi, N., Jia, Y., Kent, J., Martis, S., Shapiro, J. A., Gresham, D., Caudy, A. A., & Kruglyak, L. (2010). Dissection of genetically complex traits with extremely large pools of yeast segregants. Nature, 464(7291), 1039–1042. 10.1038/nature08923

Elserafy, M., & El-Khamisy, S. F. (2018). Choose your yeast strain carefully: The RAD5 gene matters. Nature Reviews Molecular Cell Biology, 19(6), 343–344. 10.1038/s41580-018-0005-2

Fairfax, B. P., Humburg, P., Makino, S., Naranbhai, V., Wong, D., Lau, E., Jostins, L., Plant, K., Andrews, R., McGee, C., & Knight, J. C. (2014). Innate Immune Activity Conditions the Effect of Regulatory Variants upon Monocyte Gene Expression. Science, 343(6175), 1246949. 10.1126/science.1246949

Finley, D., & Prado, M. A. (2020). The Proteasome and Its Network: Engineering for Adaptability. Cold Spring Harbor Perspectives in Biology, 12(1), a033985. 10.1101/cshperspect.a033985

Finley, D., Ulrich, H. D., Sommer, T., & Kaiser, P. (2012). The Ubiquitin–Proteasome System of *Saccharomyces cerevisiae*. Genetics, 192(2), 319–360. 10.1534/genetics.112.140467

Gaisne, M., Bécam, A. M., Verdière, J., & Herbert, C. J. (1999). A “natural” mutation in Saccharomyces cerevisiae strains derived from S288c affects the complex regulatory gene HAP1 (CYP1). Current Genetics, 36(4), 195–200. 10.1007/s002940050490

Gardner, R. G., Nelson, Z. W., & Gottschling, D. E. (2005). Degradation-mediated protein quality control in the nucleus. Cell, 120(6), 803–815. 10.1016/j.cell.2005.01.016

Gietz, R. D., & Schiestl, R. H. (2007). High-efficiency yeast transformation using the LiAc/SS carrier DNA/PEG method. Nature Protocols, 2(1), 31–34. 10.1038/nprot.2007.13

Glickman, M. H., Rubin, D. M., Coux, O., Wefes, I., Pfeifer, G., Cjeka, Z., Baumeister, W., Fried, V. A., & Finley, D. (1998). A Subcomplex of the Proteasome Regulatory Particle Required for Ubiquitin- Conjugate Degradation and Related to the COP9-Signalosome and eIF3. Cell, 94(5), 615–623. 10.1016/S0092-8674(00)81603-7

Grimm, S., Höhn, A., & Grune, T. (2012). Oxidative protein damage and the proteasome. Amino Acids, 42(1), 23–38. 10.1007/s00726-010-0646-8

Grishkevich, V., & Yanai, I. (2013). The genomic determinants of genotype × environment interactions in gene expression. Trends in Genetics, 29(8), 479–487. 10.1016/j.tig.2013.05.006

Gurganus, M. C., Fry, J. D., Nuzhdin, S. V., Pasyukova, E. G., Lyman, R. F., & Mackay, T. F. C. (1998). Genotype-Environment Interaction at Quantitative Trait Loci Affecting Sensory Bristle Number in Drosophila melanogaster. Genetics, 149(4), 1883–1898. 10.1093/genetics/149.4.1883

Guthrie, R. (1961). Blood Screening for Phenylketonuria. JAMA, 178(8), 863. 10.1001/jama.1961.03040470079019

Ha, S.-W., Ju, D., & Xie, Y. (2012). The N-terminal domain of Rpn4 serves as a portable ubiquitin- independent degron and is recognized by specific 19S RP subunits. Biochemical and Biophysical Research Communications, 419(2), 226–231. 10.1016/j.bbrc.2012.01.152

Hahne, F., LeMeur, N., Brinkman, R. R., Ellis, B., Haaland, P., Sarkar, D., Spidlen, J., Strain, E., & Gentleman, R. (2009). flowCore: A Bioconductor package for high throughput flow cytometry. BMC Bioinformatics, 10, 106. 10.1186/1471-2105-10-106

Hanna, J., & Finley, D. (2007). A proteasome for all occasions. FEBS Letters, 581(15), 2854–2861. 10.1016/j.febslet.2007.03.053

Hershko, A., & Ciechanover, A. (1998). The ubiquitin system. Annual Review of Biochemistry, 67, 425–479. 10.1146/annurev.biochem.67.1.425

Holtz, K. M., Rice, A. M., & Sartorelli, A. C. (2003). Lithium chloride inactivates the 20S proteasome from WEHI-3B D+ leukemia cells. Biochemical and Biophysical Research Communications, 303(4), 1058–1064. 10.1016/S0006-291X(03)00473-X

Hong, J., & Gresham, D. (2014). Molecular specificity, convergence and constraint shape adaptive evolution in nutrient-poor environments. PLoS Genetics, 10(1), e1004041. 10.1371/journal.pgen.1004041

Huang, W., Carbone, M. A., Lyman, R. F., Anholt, R. R. H., & Mackay, T. F. C. (2020). Genotype by environment interaction for gene expression in Drosophila melanogaster. Nature Communications, 11(1), 5451. 10.1038/s41467-020-19131-y

Ibarra, R., Sandoval, D., Fredrickson, E. K., Gardner, R. G., & Kleiger, G. (2016). The San1 Ubiquitin Ligase Functions Preferentially with Ubiquitin-conjugating Enzyme Ubc1 during Protein Quality Control. The Journal of Biological Chemistry, 291(36), 18778–18790. 10.1074/jbc.M116.737619

Inobe, T., Fishbain, S., Prakash, S., & Matouschek, A. (2011). Defining the geometry of the two-component proteasome degron. Nature Chemical Biology, 7(3), 161–167. 10.1038/nchembio.521

Johnson, E. S., Ma, P. C. M., Ota, I. M., & Varshavsky, A. (1995). A Proteolytic Pathway That Recognizes Ubiquitin as a Degradation Signal. Journal of Biological Chemistry, 270(29), 17442–17456. 10.1074/jbc.270.29.17442

Ju, D., & Xie, Y. (2004). Proteasomal degradation of RPN4 via two distinct mechanisms, ubiquitin- dependent and -independent. The Journal of Biological Chemistry, 279(23), 23851–23854. 10.1074/jbc.C400111200

Kats, I., Khmelinskii, A., Kschonsak, M., Huber, F., Knieß, R. A., Bartosik, A., & Knop, M. (2018). Mapping Degradation Signals and Pathways in a Eukaryotic N-terminome. Molecular Cell, 70(3), 488–501.e5. 10.1016/j.molcel.2018.03.033

Khmelinskii, A., Keller, P. J., Bartosik, A., Meurer, M., Barry, J. D., Mardin, B. R., Kaufmann, A., Trautmann, S., Wachsmuth, M., Pereira, G., Huber, W., Schiebel, E., & Knop, M. (2012). Tandem fluorescent protein timers for in vivo analysis of protein dynamics. Nature Biotechnology, 30(7), 708–714. 10.1038/nbt.2281

Khmelinskii, A., & Knop, M. (2014). Analysis of Protein Dynamics with Tandem Fluorescent Protein Timers. In A. I. Ivanov (Ed.), Exocytosis and Endocytosis (Vol. 1174, pp. 195–210). Springer New York. 10.1007/978-1-4939-0944-5_13

Kim-Hellmuth, S., Bechheim, M., Pütz, B., Mohammadi, P., Nédélec, Y., Giangreco, N., Becker, J., Kaiser, V., Fricker, N., Beier, E., Boor, P., Castel, S. E., Nöthen, M. M., Barreiro, L. B., Pickrell, J. K., Müller-Myhsok, B., Lappalainen, T., Schumacher, J., & Hornung, V. (2017). Genetic regulatory effects modified by immune activation contribute to autoimmune disease associations. Nature Communications, 8(1), 266. 10.1038/s41467-017-00366-1

Kong, K.-Y. E., Fischer, B., Meurer, M., Kats, I., Li, Z., Rühle, F., Barry, J. D., Kirrmaier, D., Chevyreva, V., San Luis, B.-J., Costanzo, M., Huber, W., Andrews, B. J., Boone, C., Knop, M., & Khmelinskii, A. (2021). Timer-based proteomic profiling of the ubiquitin-proteasome system reveals a substrate receptor of the GID ubiquitin ligase. Molecular Cell, 81(11), 2460–2476.e11. 10.1016/j.molcel.2021.04.018

Kredel, S., Oswald, F., Nienhaus, K., Deuschle, K., Röcker, C., Wolff, M., Heilker, R., Nienhaus, G. U., & Wiedenmann, J. (2009). mRuby, a bright monomeric red fluorescent protein for labeling of subcellular structures. PloS One, 4(2), e4391. 10.1371/journal.pone.0004391

Kuzmin, E., Costanzo, M., Andrews, B., & Boone, C. (2016). Synthetic Genetic Array Analysis. Cold Spring Harbor Protocols, 2016(4), pdb.prot088807. 10.1101/pdb.prot088807

Laporte, D., Salin, B., Daignan-Fornier, B., & Sagot, I. (2008). Reversible cytoplasmic localization of the proteasome in quiescent yeast cells. The Journal of Cell Biology, 181(5), 737–745. 10.1083/jcb.200711154

Lea, A. J., Peng, J., & Ayroles, J. F. (2022). Diverse environmental perturbations reveal the evolution and context-dependency of genetic effects on gene expression levels. Genome Research, genome;gr.276430.121v1. 10.1101/gr.276430.121

Lee, M. N., Ye, C., Villani, A.-C., Raj, T., Li, W., Eisenhaure, T. M., Imboywa, S. H., Chipendo, P. I., Ran, F. A., Slowikowski, K., Ward, L. D., Raddassi, K., McCabe, C., Lee, M. H., Frohlich, I. Y., Hafler, D. A., Kellis, M., Raychaudhuri, S., Zhang, F., … Hacohen, N. (2014). Common Genetic Variants Modulate Pathogen-Sensing Responses in Human Dendritic Cells. Science, 343(6175), 1246980. 10.1126/science.1246980

Li, C., Liang, X., Cheng, S., Wen, Y., Pan, C., Zhang, H., Chen, Y., Zhang, J., Zhang, Z., Yang, X., Meng, P., & Zhang, F. (2022). A multi-environments-gene interaction study of anxiety, depression and self- harm in the UK Biobank cohort. Journal of Psychiatric Research, 147, 59–66. 10.1016/j.jpsychires.2022.01.009

Li, H., & Durbin, R. (2009). Fast and accurate short read alignment with Burrows-Wheeler transform. Bioinformatics (Oxford, England), 25(14), 1754–1760. 10.1093/bioinformatics/btp324

Li, H., Handsaker, B., Wysoker, A., Fennell, T., Ruan, J., Homer, N., Marth, G., Abecasis, G., Durbin, R., & 1000 Genome Project Data Processing Subgroup. (2009). The Sequence Alignment/Map format and SAMtools. Bioinformatics (Oxford, England), *25*(16), 2078–2079. 10.1093/bioinformatics/btp352

Li, J., Breker, M., Graham, M., Schuldiner, M., & Hochstrasser, M. (2019). AMPK regulates ESCRT- dependent microautophagy of proteasomes concomitant with proteasome storage granule assembly during glucose starvation. PLOS Genetics, 15(11), e1008387. 10.1371/journal.pgen.1008387

Li, Y., Álvarez, O. A., Gutteling, E. W., Tijsterman, M., Fu, J., Riksen, J. A. G., Hazendonk, E., Prins, P., Plasterk, R. H. A., Jansen, R. C., Breitling, R., & Kammenga, J. E. (2006). Mapping Determinants of Gene Expression Plasticity by Genetical Genomics in C. elegans. PLoS Genetics, 2(12), e222. 10.1371/journal.pgen.0020222

Lutz, S., Van Dyke, K., Feraru, M. A., & Albert, F. W. (2021). Multiple epistatic DNA variants in a single gene affect gene expression in *trans*. *Genetics*, iyab208. 10.1093/genetics/iyab208

Marshall, R. S., McLoughlin, F., & Vierstra, R. D. (2016). Autophagic Turnover of Inactive 26S Proteasomes in Yeast Is Directed by the Ubiquitin Receptor Cue5 and the Hsp42 Chaperone. Cell Reports, 16(6), 1717–1732. 10.1016/j.celrep.2016.07.015

Martinez-Fonts, K., Davis, C., Tomita, T., Elsasser, S., Nager, A. R., Shi, Y., Finley, D., & Matouschek, A. (2020). The proteasome 19S cap and its ubiquitin receptors provide a versatile recognition platform for substrates. Nature Communications, 11(1), 477. 10.1038/s41467-019-13906-8

Matheson, K., Parsons, L., & Gammie, A. (2017). Whole-Genome Sequence and Variant Analysis of W303, a Widely-Used Strain of *Saccharomyces cerevisiae*. G3 Genes|Genomes|Genetics, *7*(7), 2219–2226. 10.1534/g3.117.040022

Michelmore, R. W., Paran, I., & Kesseli, R. V. (1991). Identification of markers linked to disease-resistance genes by bulked segregant analysis: A rapid method to detect markers in specific genomic regions by using segregating populations. Proceedings of the National Academy of Sciences of the United States of America, 88(21), 9828–9832. 10.1073/pnas.88.21.9828

Moye-Rowley, W. S. (2003). Transcriptional control of multidrug resistance in the yeast Saccharomyces. Progress in Nucleic Acid Research and Molecular Biology, 73, 251–279. 10.1016/s0079-6603(03)01008-0

Nassar, L. R., Barber, G. P., Benet-Pagès, A., Casper, J., Clawson, H., Diekhans, M., Fischer, C., Gonzalez, J. N., Hinrichs, A. S., Lee, B. T., Lee, C. M., Muthuraman, P., Nguy, B., Pereira, T., Nejad, P., Perez, G., Raney, B. J., Schmelter, D., Speir, M. L., … Kent, W. J. (2023). The UCSC Genome Browser database: 2023 update. Nucleic Acids Research, 51(D1), D1188–D1195. 10.1093/nar/gkac1072

Nédélec, Y., Sanz, J., Baharian, G., Szpiech, Z. A., Pacis, A., Dumaine, A., Grenier, J.-C., Freiman, A., Sams, A. J., Hebert, S., Pagé Sabourin, A., Luca, F., Blekhman, R., Hernandez, R. D., Pique-Regi, R., Tung, J., Yotova, V., & Barreiro, L. B. (2016). Genetic Ancestry and Natural Selection Drive Population Differences in Immune Responses to Pathogens. Cell, 167(3), 657–669.e21. 10.1016/j.cell.2016.09.025

Nguyen Ba, A. N., Lawrence, K. R., Rego-Costa, A., Gopalakrishnan, S., Temko, D., Michor, F., & Desai, M. M. (2022). Barcoded bulk QTL mapping reveals highly polygenic and epistatic architecture of complex traits in yeast. eLife, 11, e73983. 10.7554/eLife.73983

Nunes, A. T., & Annunziata, C. M. (2017). Proteasome inhibitors: Structure and function. Seminars in Oncology, 44(6), 377–380. 10.1053/j.seminoncol.2018.01.004

Pirmohamed, M. (2023). Pharmacogenomics: Current status and future perspectives. Nature Reviews Genetics, 24(6), 350–362. 10.1038/s41576-022-00572-8

Prakash, S., Tian, L., Ratliff, K. S., Lehotzky, R. E., & Matouschek, A. (2004). An unstructured initiation site is required for efficient proteasome-mediated degradation. Nature Structural & Molecular Biology, 11(9), 830–837. 10.1038/nsmb814

Quach, H., Rotival, M., Pothlichet, J., Loh, Y.-H. E., Dannemann, M., Zidane, N., Laval, G., Patin, E., Harmant, C., Lopez, M., Deschamps, M., Naffakh, N., Duffy, D., Coen, A., Leroux-Roels, G., Clément, F., Boland, A., Deleuze, J.-F., Kelso, J., … Quintana-Murci, L. (2016). Genetic Adaptation and Neandertal Admixture Shaped the Immune System of Human Populations. Cell, 167(3), 643–656.e17. 10.1016/j.cell.2016.09.024

Renganaath, K., & Albert, F. W. (2023). Trans *-eQTL hotspots shape complex traits by modulating cellular states*. Genetics. 10.1101/2023.11.14.567054

Robinson, M. R., English, G., Moser, G., Lloyd-Jones, L. R., Triplett, M. A., Zhu, Z., Nolte, I. M., Van Vliet- Ostaptchouk, J. V., Snieder, H., Esko, T., Milani, L., Mägi, R., Metspalu, A., Magnusson, P. K. E., Pedersen, N. L., Ingelsson, E., Johannesson, M., Yang, J., Cesarini, D., & Visscher, P. M. (2017). Genotype–covariate interaction effects and the heritability of adult body mass index. Nature Genetics, 49(8), 1174–1181. 10.1038/ng.3912

Rodgers, K. J., & Shiozawa, N. (2008). Misincorporation of amino acid analogues into proteins by biosynthesis. The International Journal of Biochemistry & Cell Biology, 40(8), 1452–1466. 10.1016/j.biocel.2008.01.009

Rosenbaum, J. C., Fredrickson, E. K., Oeser, M. L., Garrett-Engele, C. M., Locke, M. N., Richardson, L. A., Nelson, Z. W., Hetrick, E. D., Milac, T. I., Gottschling, D. E., & Gardner, R. G. (2011). Disorder targets misorder in nuclear quality control degradation: A disordered ubiquitin ligase directly recognizes its misfolded substrates. Molecular Cell, 41(1), 93–106. 10.1016/j.molcel.2010.12.004

Saeki, Y., Toh-e, A., & Yokosawa, H. (2000). Rapid Isolation and Characterization of the Yeast Proteasome Regulatory Complex. Biochemical and Biophysical Research Communications, 273(2), 509–515. 10.1006/bbrc.2000.2980

Sasaki, E., Zhang, P., Atwell, S., Meng, D., & Nordborg, M. (2015). “Missing” G x E Variation Controls Flowering Time in Arabidopsis thaliana. PLOS Genetics, 11(10), e1005597. 10.1371/journal.pgen.1005597

Schwartz, A. L., & Ciechanover, A. (1999). The ubiquitin-proteasome pathway and pathogenesis of human diseases. Annual Review of Medicine, 50, 57–74. 10.1146/annurev.med.50.1.57

Shaner, N. C., Campbell, R. E., Steinbach, P. A., Giepmans, B. N. G., Palmer, A. E., & Tsien, R. Y. (2004). Improved monomeric red, orange and yellow fluorescent proteins derived from Discosoma sp. Red fluorescent protein. Nature Biotechnology, 22(12), 1567–1572. 10.1038/nbt1037

Shostak, S. (2003). Locating gene–environment interaction: At the intersections of genetics and public health. Social Science & Medicine, 56(11), 2327–2342. 10.1016/S0277-9536(02)00231-9

Shringarpure, R., & Davies, K. J. A. (2002). Protein turnover by the proteasome in aging and disease. Free Radical Biology & Medicine, 32(11), 1084–1089. 10.1016/s0891-5849(02)00824-9

Simon, M. M., Greenaway, S., White, J. K., Fuchs, H., Gailus-Durner, V., Wells, S., Sorg, T., Wong, K., Bedu, E., Cartwright, E. J., Dacquin, R., Djebali, S., Estabel, J., Graw, J., Ingham, N. J., Jackson, I. J., Lengeling, A., Mandillo, S., Marvel, J., … Brown, S. D. (2013). A comparative phenotypic and genomic analysis of C57BL/6J and C57BL/6N mouse strains. Genome Biology, 14(7), R82. 10.1186/gb-2013-14-7-r82

Smith, E. N., & Kruglyak, L. (2008). Gene–Environment Interaction in Yeast Gene Expression. PLoS Biology, 6(4), e83. 10.1371/journal.pbio.0060083

Sontag, E. M., Vonk, W. I. M., & Frydman, J. (2014). Sorting out the trash: The spatial nature of eukaryotic protein quality control. Current Opinion in Cell Biology, 26, 139–146. 10.1016/j.ceb.2013.12.006

Stack, J. H., Whitney, M., Rodems, S. M., & Pollok, B. A. (2000). A ubiquitin-based tagging system for controlled modulation of protein stability. Nature Biotechnology, 18(12), 1298–1302. 10.1038/82422

Tanaka, K., Nakafuku, M., Tamanoi, F., Kaziro, Y., Matsumoto, K., & Toh-e, A. (1990). IRA2, a second gene of Saccharomyces cerevisiae that encodes a protein with a domain homologous to mammalian ras GTPase-activating protein. Molecular and Cellular Biology, 10(8), 4303–4313. 10.1128/mcb.10.8.4303-4313.1990

Theodoraki, M. A., Nillegoda, N. B., Saini, J., & Caplan, A. J. (2012). A network of ubiquitin ligases is important for the dynamics of misfolded protein aggregates in yeast. The Journal of Biological Chemistry, 287(28), 23911–23922. 10.1074/jbc.M112.341164

Thrower, J. S., Hoffman, L., Rechsteiner, M, & Pickart, C. M. (2000). Recognition of the polyubiquitin proteolytic signal. The EMBO Journal, 19(1), 94–102. 10.1093/emboj/19.1.94

Treusch, S., Albert, F. W., Bloom, J. S., Kotenko, I. E., & Kruglyak, L. (2015). Genetic Mapping of MAPK-Mediated Complex Traits Across S. cerevisiae. PLoS Genetics, 11(1), e1004913. 10.1371/journal.pgen.1004913

Vabulas, R. M., & Hartl, F. U. (2005). Protein synthesis upon acute nutrient restriction relies on proteasome function. *Science (New York*, N.Y.), 310(5756), 1960–1963. 10.1126/science.1121925

Varshavsky, A. (1991). Naming a targeting signal. Cell, 64(1), 13–15. 10.1016/0092-8674(91)90202-a

Varshavsky, A. (2011). The N-end rule pathway and regulation by proteolysis. Protein Science: A Publication of the Protein Society, 20(8), 1298–1345. 10.1002/pro.666

Varshavsky, A. (2019). N-degron and C-degron pathways of protein degradation. Proceedings of the National Academy of Sciences, 116(2), 358–366. 10.1073/pnas.1816596116

Varshavsky, A. (2024). N-degron pathways. Proceedings of the National Academy of Sciences, 121(39). 10.1073/pnas.2408697121

Waite, K. A., Mota-Peynado, A. D.-L., Vontz, G., & Roelofs, J. (2016). Starvation Induces Proteasome Autophagy with Different Pathways for Core and Regulatory Particles. Journal of Biological Chemistry, 291(7), 3239–3253. 10.1074/jbc.M115.699124

Warringer, J., Zörgö, E., Cubillos, F. A., Zia, A., Gjuvsland, A., Simpson, J. T., Forsmark, A., Durbin, R., Omholt, S. W., Louis, E. J., Liti, G., Moses, A., & Blomberg, A. (2011). Trait Variation in Yeast Is Defined by Population History. PLoS Genetics, 7(6), e1002111. 10.1371/journal.pgen.1002111

Wenger, J. W., Piotrowski, J., Nagarajan, S., Chiotti, K., Sherlock, G., & Rosenzweig, F. (2011). Hunger artists: Yeast adapted to carbon limitation show trade-offs under carbon sufficiency. PLoS Genetics, 7(8), e1002202. 10.1371/journal.pgen.1002202

Wickner, R. B. (1987). MKT1, a nonessential Saccharomyces cerevisiae gene with a temperature- dependent effect on replication of M2 double-stranded RNA. Journal of Bacteriology, 169(11), 4941–4945. 10.1128/jb.169.11.4941-4945.1987

Widaman, K. F. (2009). Phenylketonuria in Children and Mothers: Genes, Environments, Behavior. Current Directions in Psychological Science, 18(1), 48–52. 10.1111/j.1467-8721.2009.01604.x

Work, J. J., & Brandman, O. (2021). Adaptability of the ubiquitin-proteasome system to proteolytic and folding stressors. Journal of Cell Biology, 220(3), e201912041. 10.1083/jcb.201912041

Xie, Y., & Varshavsky, A. (2001). RPN4 is a ligand, substrate, and transcriptional regulator of the 26S proteasome: A negative feedback circuit. Proceedings of the National Academy of Sciences of the United States of America, 98(6), 3056–3061. 10.1073/pnas.071022298

Yadav, A., & Sinha, H. (2018). Gene–gene and gene–environment interactions in complex traits in yeast. Yeast, 35(6), 403–416. 10.1002/yea.3304

Yang, T., Tang, H., Risch, H. A., Olson, S. H., Peterson, G., Bracci, P. M., Gallinger, S., Hung, R. J., Neale, R. E., Scelo, G., Duell, E. J., Kurtz, R. C., Khaw, K.-T., Severi, G., Sund, M., Wareham, N., Amos, C. I., Li, D., & Wei, P. (2020). Incorporating multiple sets of eQTL weights into gene-by-environment interaction analysis identifies novel susceptibility loci for pancreatic cancer. Genetic Epidemiology, 44(8), 880–892. 10.1002/gepi.22348

Zhao, S., & Ulrich, H. D. (2010). Distinct consequences of posttranslational modification by linear versus K63-linked polyubiquitin chains. Proceedings of the National Academy of Sciences, 107(17), 7704– 7709. 10.1073/pnas.0908764107

Zheng, C., Geetha, T., & Babu, J. R. (2014). Failure of ubiquitin proteasome system: Risk for neurodegenerative diseases. Neuro-Degenerative Diseases, 14(4), 161–175. 10.1159/000367694

Zhu, J., Zhang, B., Smith, E. N., Drees, B., Brem, R. B., Kruglyak, L., Bumgarner, R. E., & Schadt, E. E. (2008). Integrating large-scale functional genomic data to dissect the complexity of yeast regulatory networks. Nature Genetics, 40(7), 854–861. 10.1038/ng.167

