## Supplementary File 2 for "Genotype-by-environment interactions shape ubiquitin-proteasome system activity"

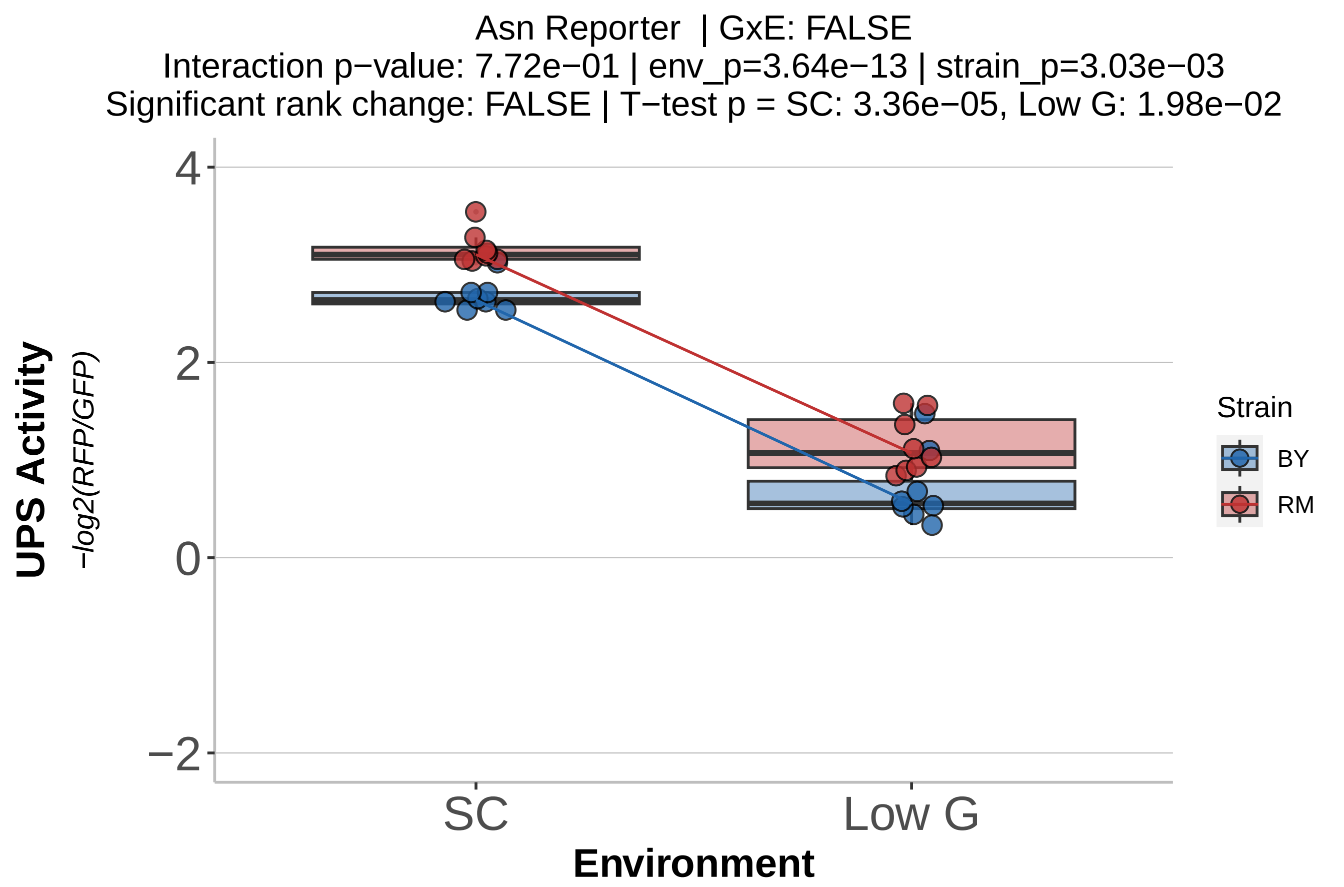
Asn Reporter | GxE: FALSE
Interaction p−value: 7.72e−01 | env_p=3.64e−13 | strain_p=3.03e−03
Significant rank change: FALSE | T−test p = SC: 3.36e−05, Low G: 1.98e−02
UPS Activity
−log2(RFP/GFP)
4
2
0
−2
SC
Low G
Environment
Strain
BY
RM

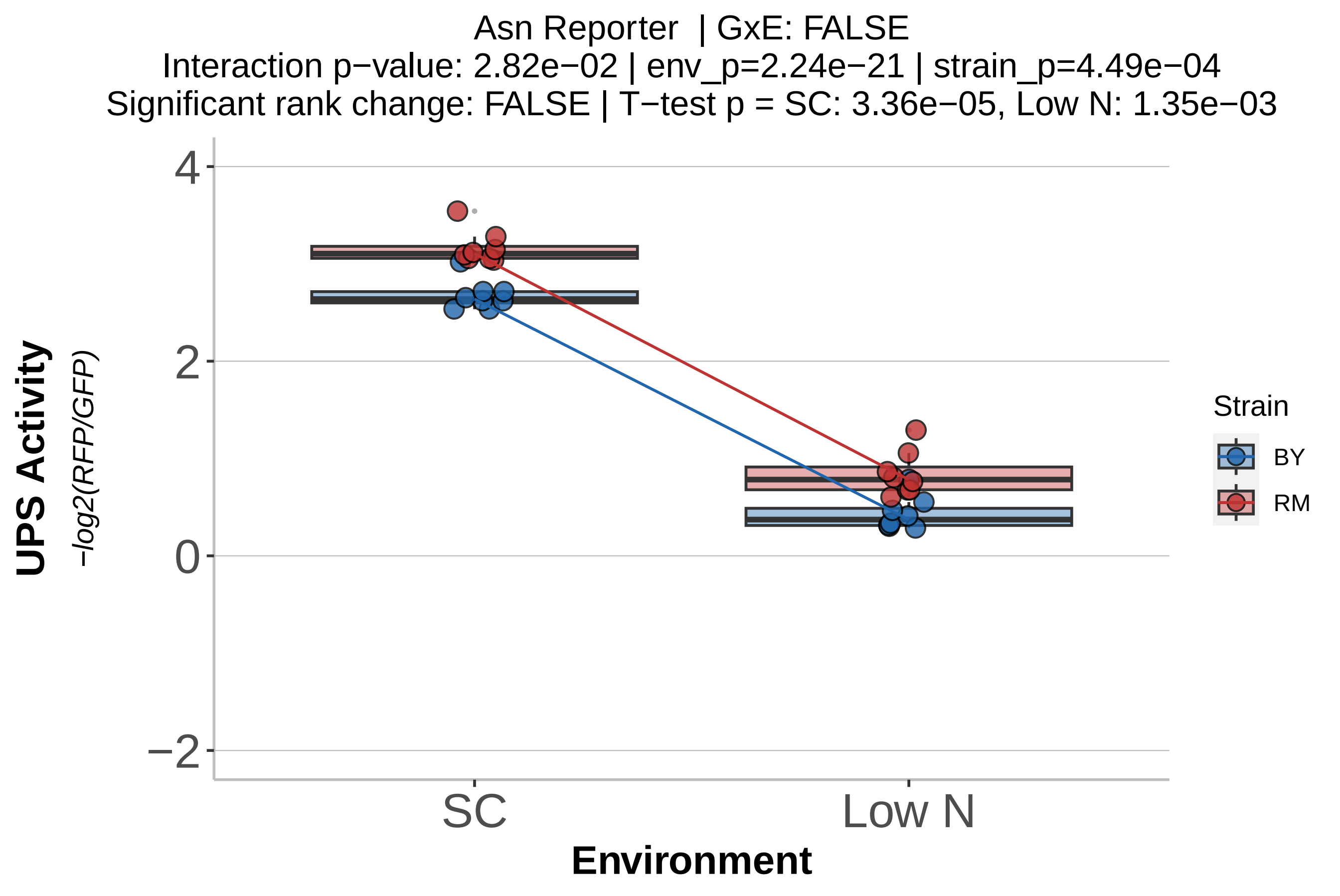
Asn Reporter | GxE: FALSE
Interaction p−value: 2.82e−02 | env_p=2.24e−21 | strain_p=4.49e−04
Significant rank change: FALSE | T−test p = SC: 3.36e−05, Low N: 1.35e−03
UPS Activity
−log2(RFP/GFP)
4
2
0
−2
SC
Low N
Environment
Strain
BY
RM

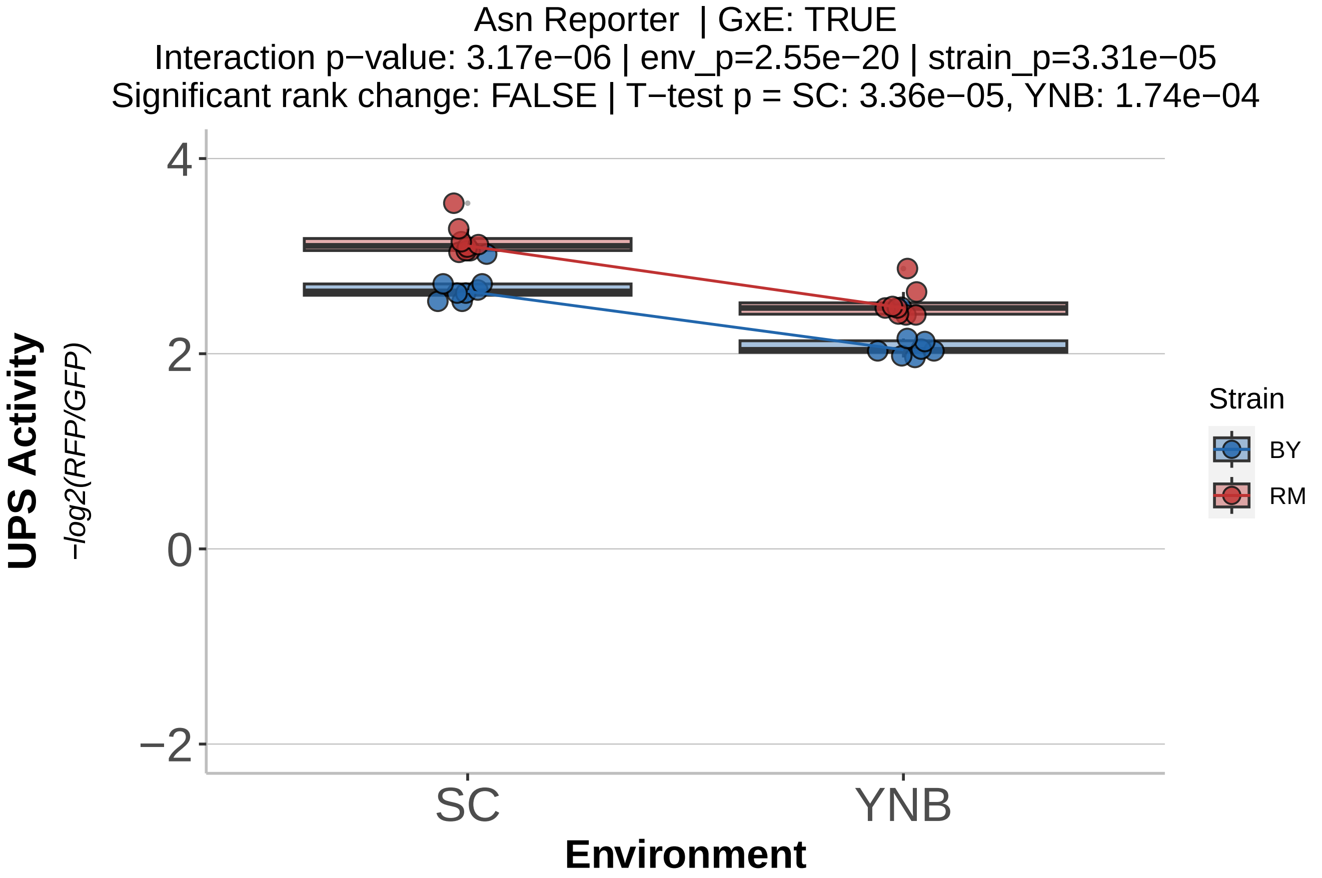
Asn Reporter | GxE: TRUE
Interaction p−value: 3.17e−06 | env_p=2.55e−20 | strain_p=3.31e−05
Significant rank change: FALSE | T−test p = SC: 3.36e−05, YNB: 1.74e−04
UPS Activity
−log2(RFP/GFP)
4
2
0
−2
SC
YNB
Environment
Strain
BY
RM

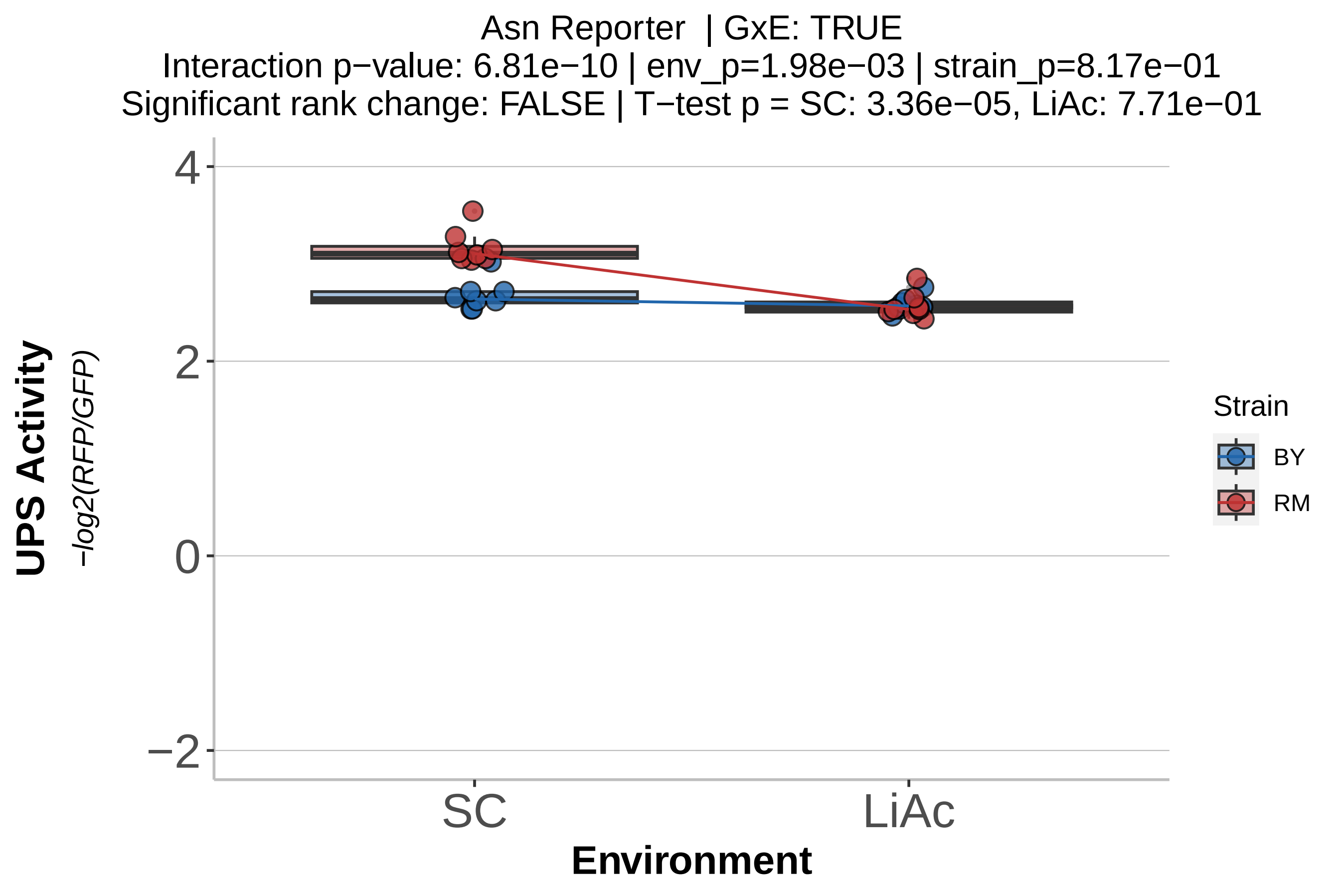

Asn Reporter | GxE: TRUE
Interaction p−value: 6.81e−10 | env_p=1.98e−03 | strain_p=8.17e−01
Significant rank change: FALSE | T−test p = SC: 3.36e−05, LiAc: 7.71e−01
UPS Activity
−log2(RFP/GFP)
4
2
0
−2
SC
LiAc
Environment
Strain
BY
RM

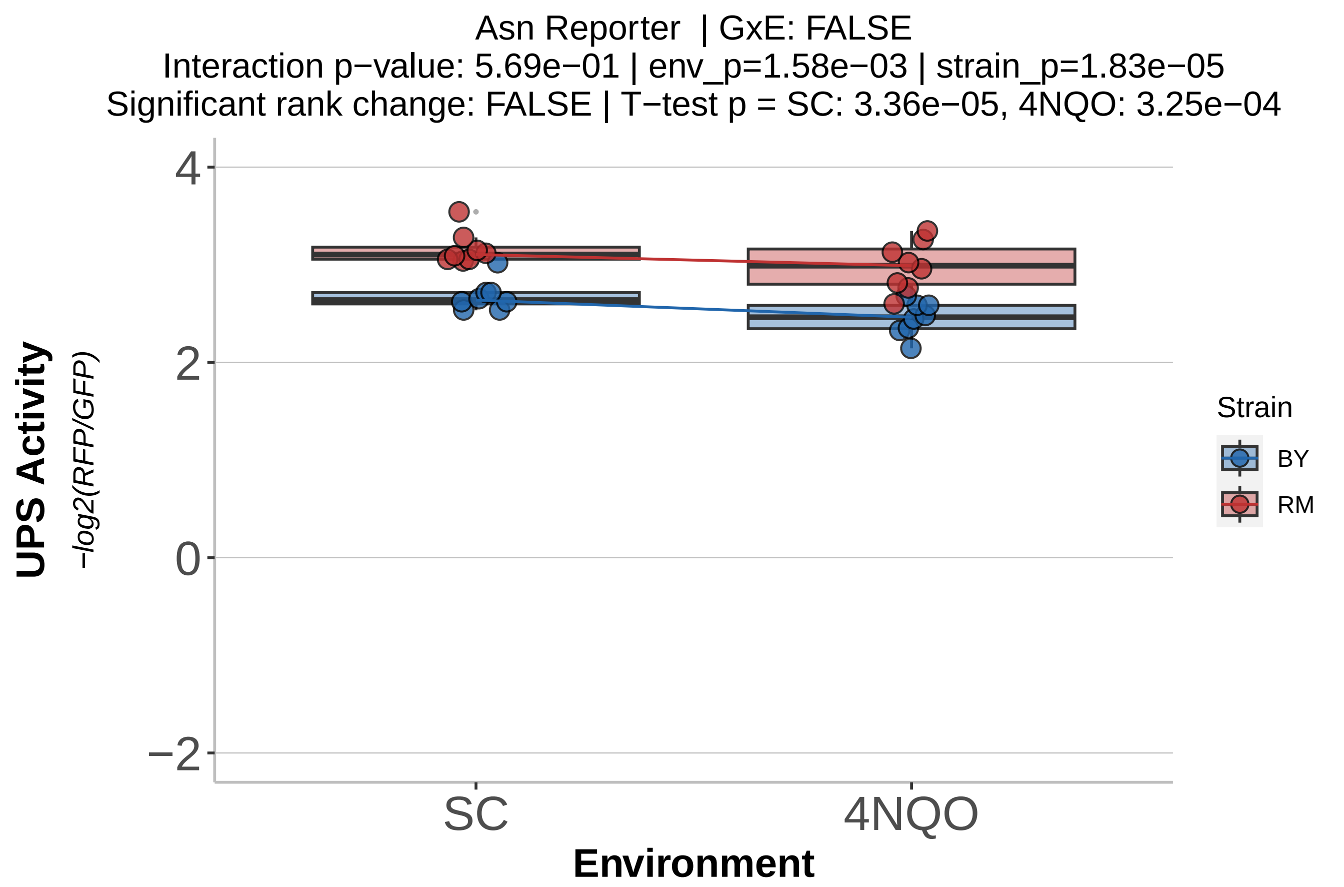

Asn Reporter | GxE: FALSE
Interaction p−value: 5.69e−01 | env_p=1.58e−03 | strain_p=1.83e−05
Significant rank change: FALSE | T−test p = SC: 3.36e−05, 4NQO: 3.25e−04
UPS Activity
−log2(RFP/GFP)
4
2
0
−2
SC
4NQO
Environment
Strain
BY
RM

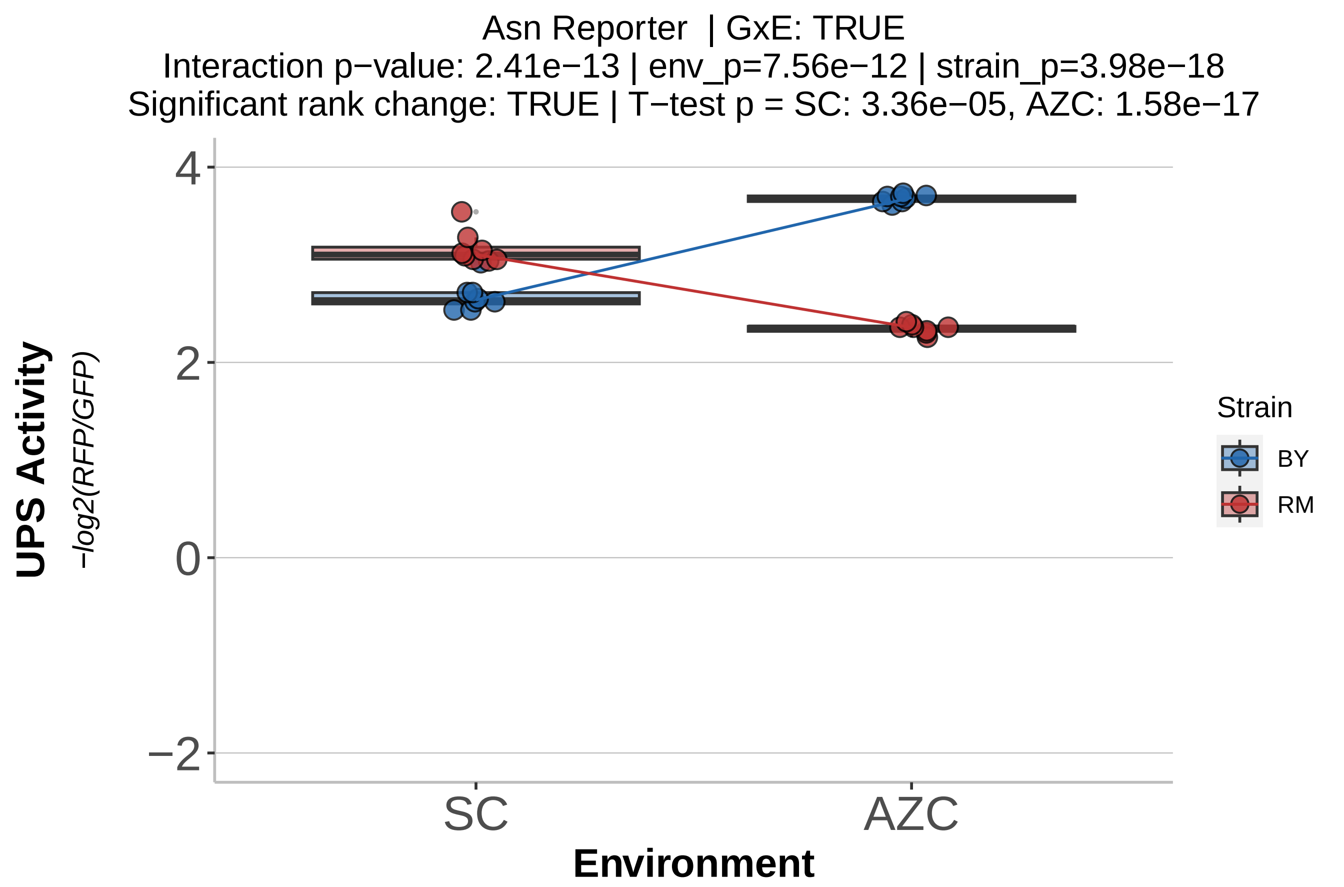

Asn Reporter | GxE: TRUE
Interaction p−value: 2.41e−13 | env_p=7.56e−12 | strain_p=3.98e−18
Significant rank change: TRUE | T−test p = SC: 3.36e−05, AZC: 1.58e−17
UPS Activity
−log2(RFP/GFP)
4
2
0
−2
SC
AZC
Environment
Strain
BY
RM

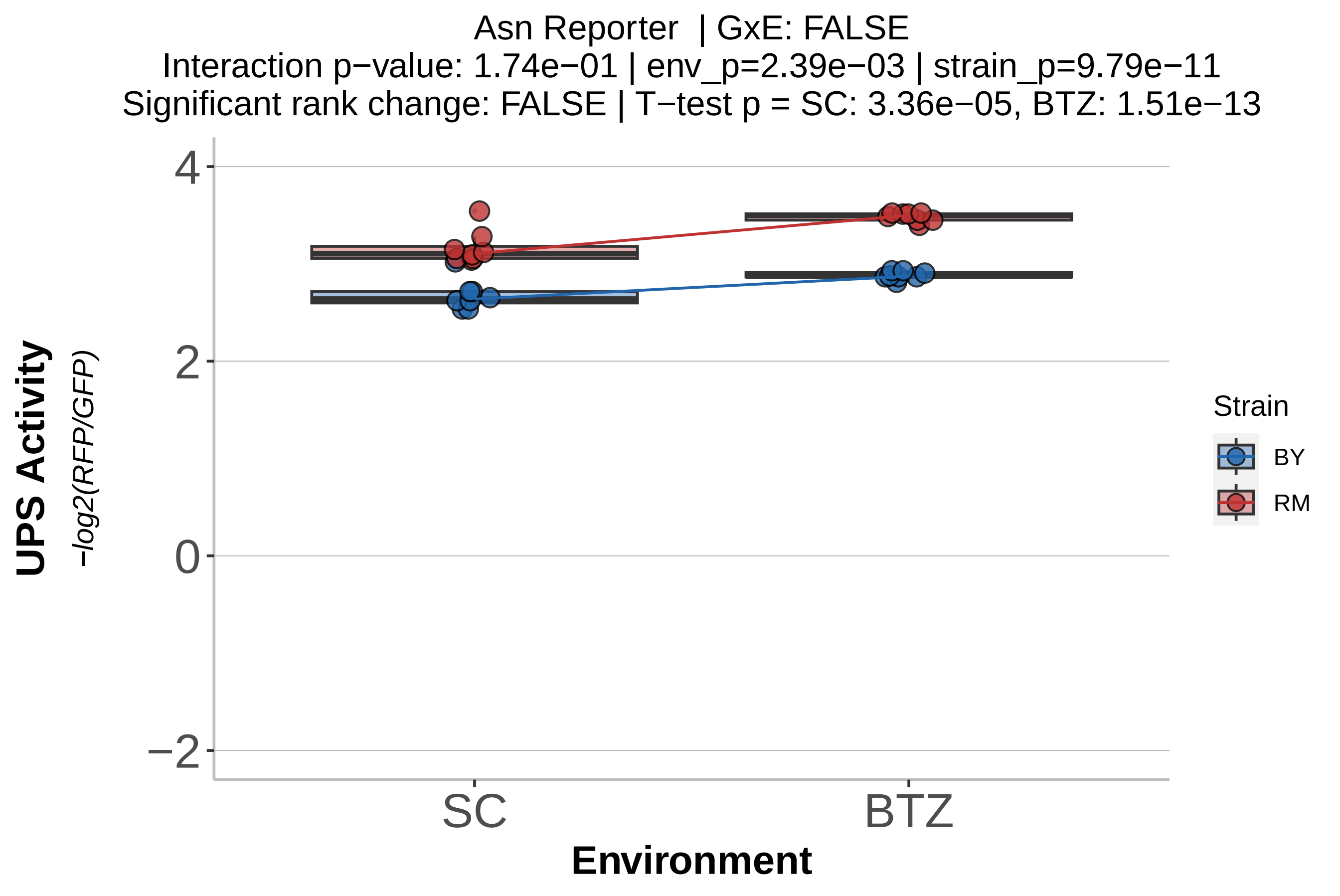
Asn Reporter | GxE: FALSE
Interaction p−value: 1.74e−01 | env_p=2.39e−03 | strain_p=9.79e−11
Significant rank change: FALSE | T−test p = SC: 3.36e−05, BTZ: 1.51e−13
UPS Activity
−log2(RFP/GFP)
4
2
0
−2
SC
BTZ
Environment
Strain
BY
RM

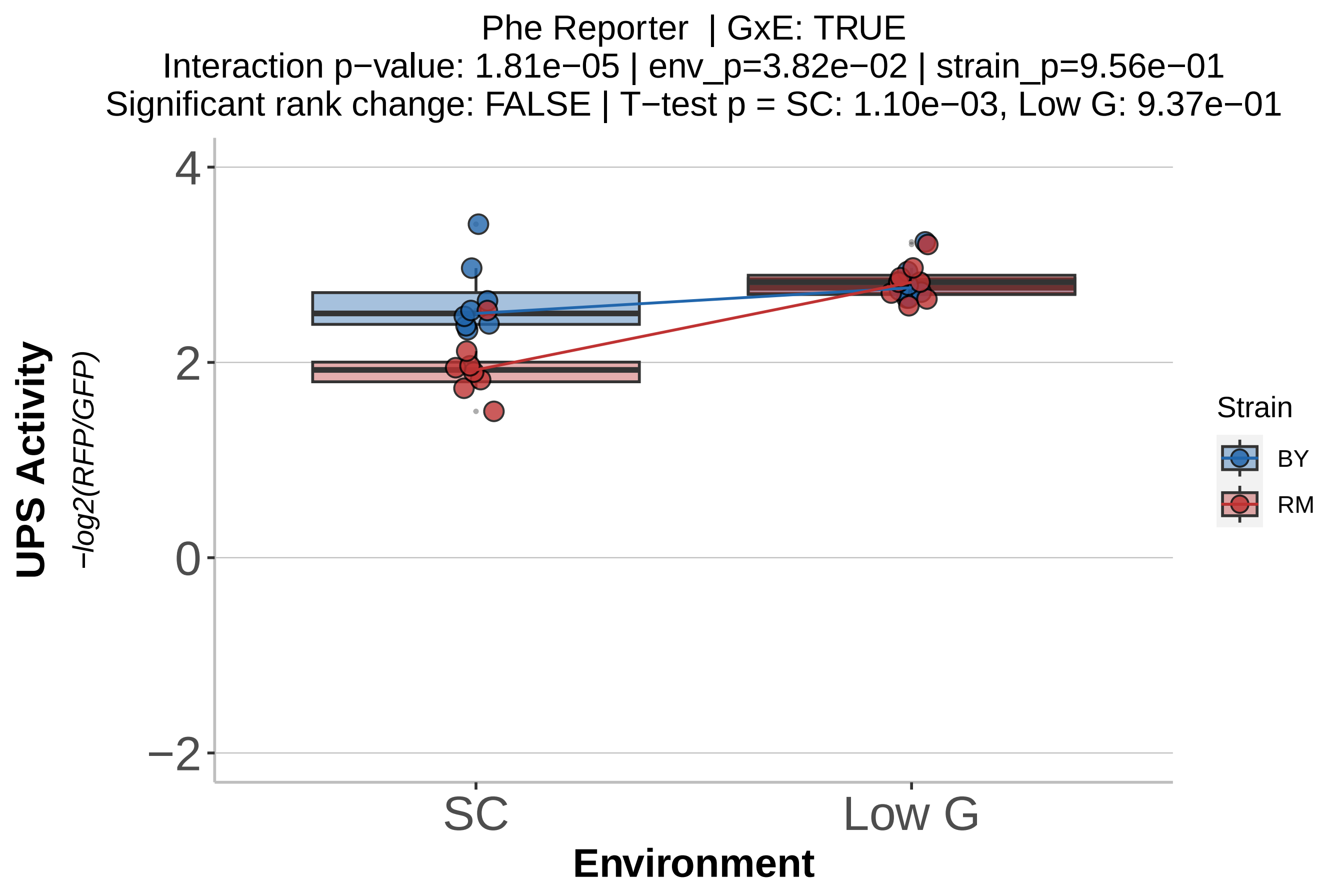
Phe Reporter | GxE: TRUE
Interaction p−value: 1.81e−05 | env_p=3.82e−02 | strain_p=9.56e−01
Significant rank change: FALSE | T−test p = SC: 1.10e−03, Low G: 9.37e−01
UPS Activity
−log2(RFP/GFP)
4
2
0
−2
SC
Low G
Environment
Strain
BY
RM

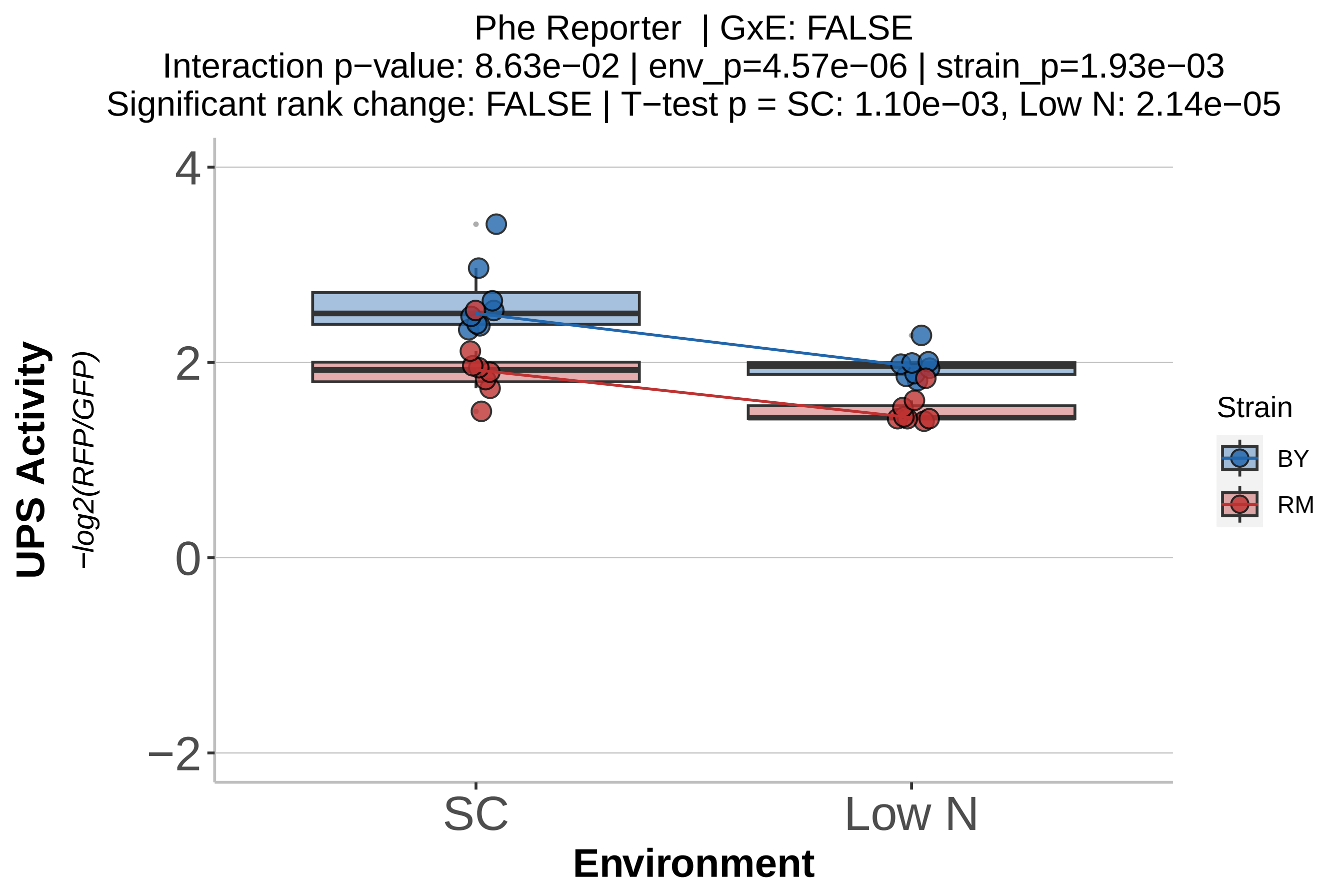

Phe Reporter | GxE: FALSE
Interaction p−value: 8.63e−02 | env_p=4.57e−06 | strain_p=1.93e−03
Significant rank change: FALSE | T−test p = SC: 1.10e−03, Low N: 2.14e−05
UPS Activity
−log2(RFP/GFP)
4
2
0
−2
SC
Low N
Environment
Strain
BY
RM

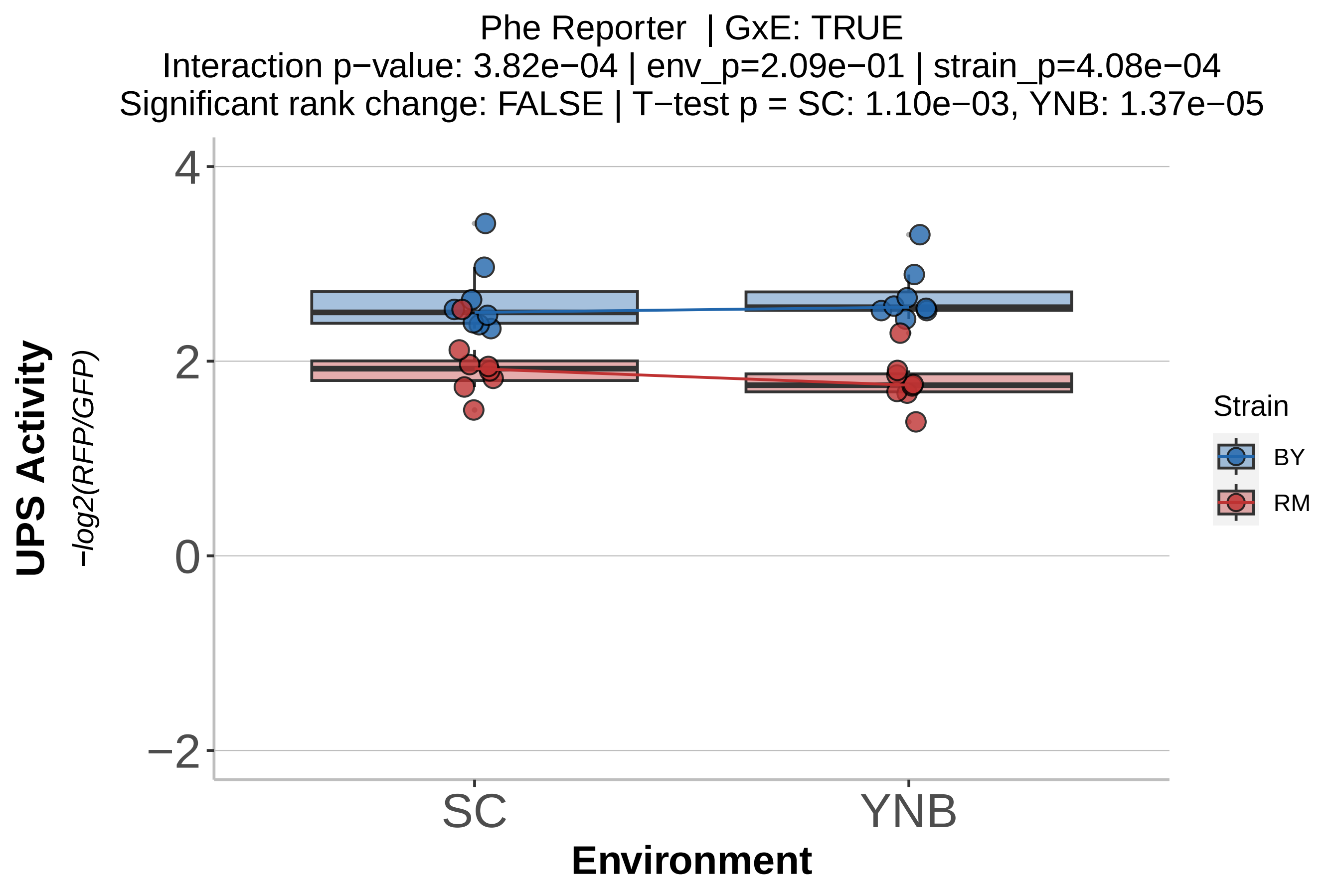
Phe Reporter | GxE: TRUE
Interaction p−value: 3.82e−04 | env_p=2.09e−01 | strain_p=4.08e−04
Significant rank change: FALSE | T−test p = SC: 1.10e−03, YNB: 1.37e−05
UPS Activity
−log2(RFP/GFP)
4
2
0
−2
SC
YNB
Environment
Strain
BY
RM

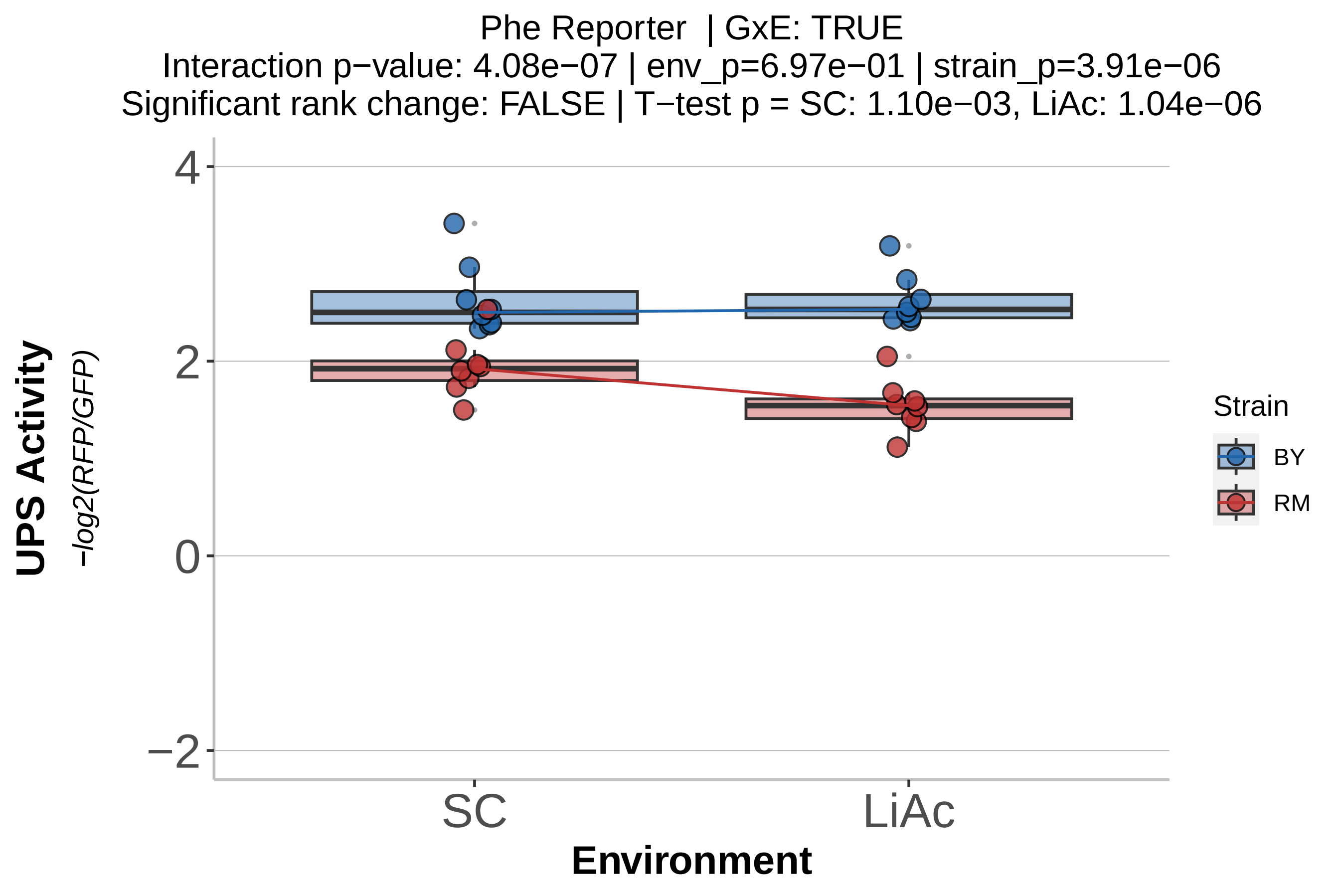
Phe Reporter | GxE: TRUE
Interaction p−value: 4.08e−07 | env_p=6.97e−01 | strain_p=3.91e−06
Significant rank change: FALSE | T−test p = SC: 1.10e−03, LiAc: 1.04e−06
UPS Activity
−log2(RFP/GFP)
4
2
0
−2
SC
LiAc
Environment
Strain
BY
RM

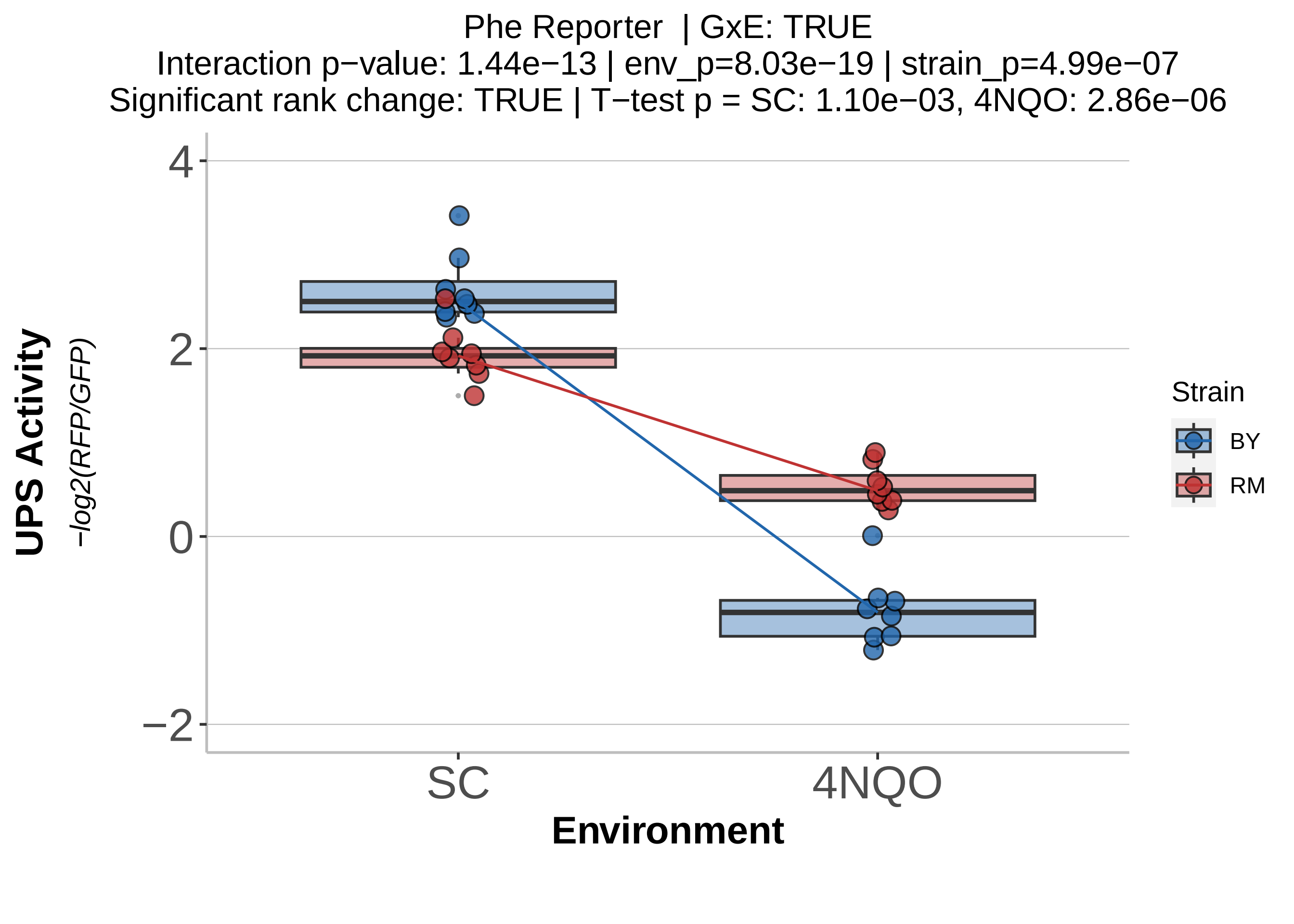
Phe Reporter | GxE: TRUE
Interaction p−value: 1.44e−13 | env_p=8.03e−19 | strain_p=4.99e−07
Significant rank change: TRUE | T−test p = SC: 1.10e−03, 4NQO: 2.86e−06
UPS Activity
−log2(RFP/GFP)
4
2
0
−2
SC
4NQO
Environment
Strain
BY
RM

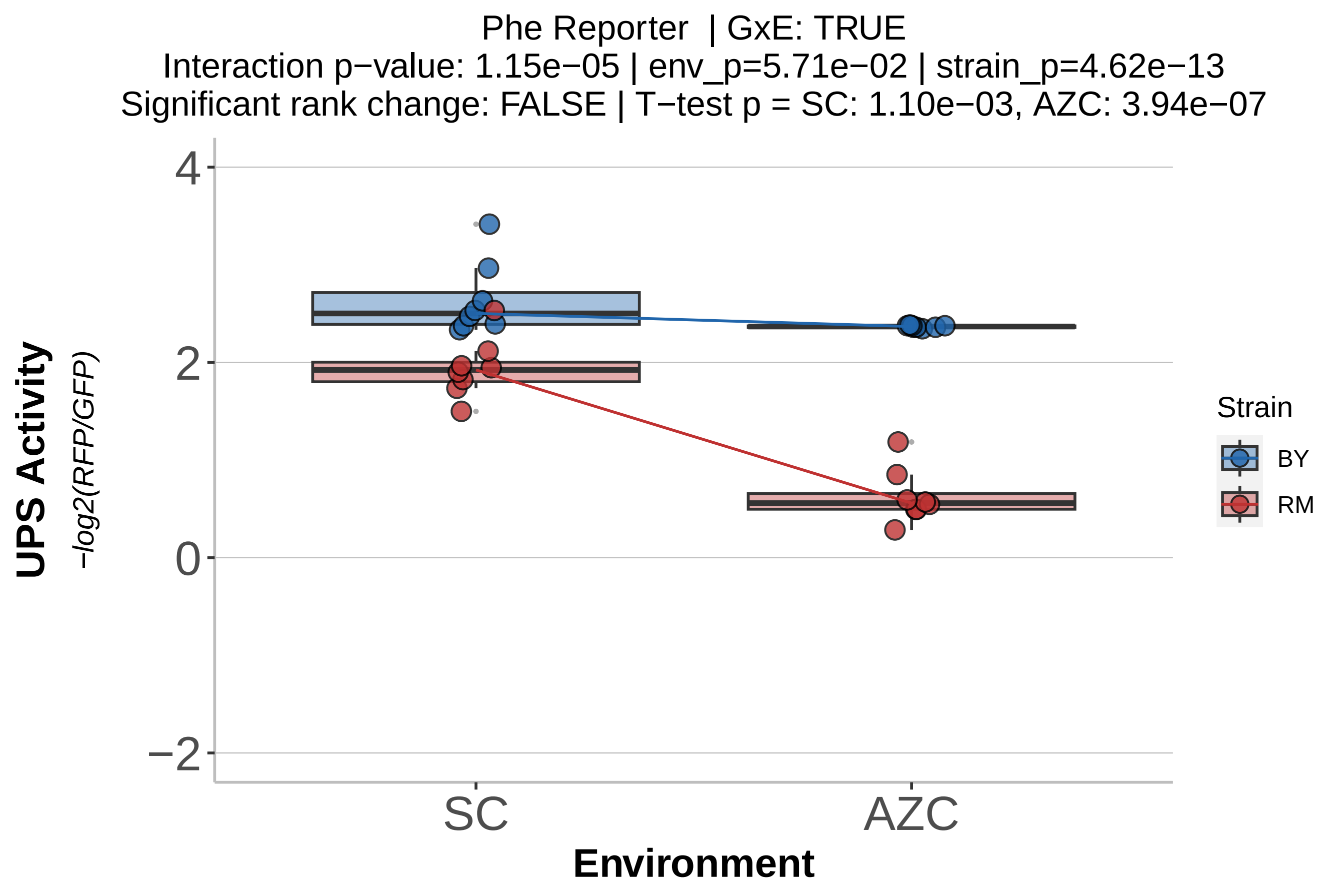

Phe Reporter | GxE: TRUE
Interaction p−value: 1.15e−05 | env_p=5.71e−02 | strain_p=4.62e−13
Significant rank change: FALSE | T−test p = SC: 1.10e−03, AZC: 3.94e−07
UPS Activity
−log2(RFP/GFP)
4
2
0
−2
SC
AZC
Environment
Strain
BY
RM

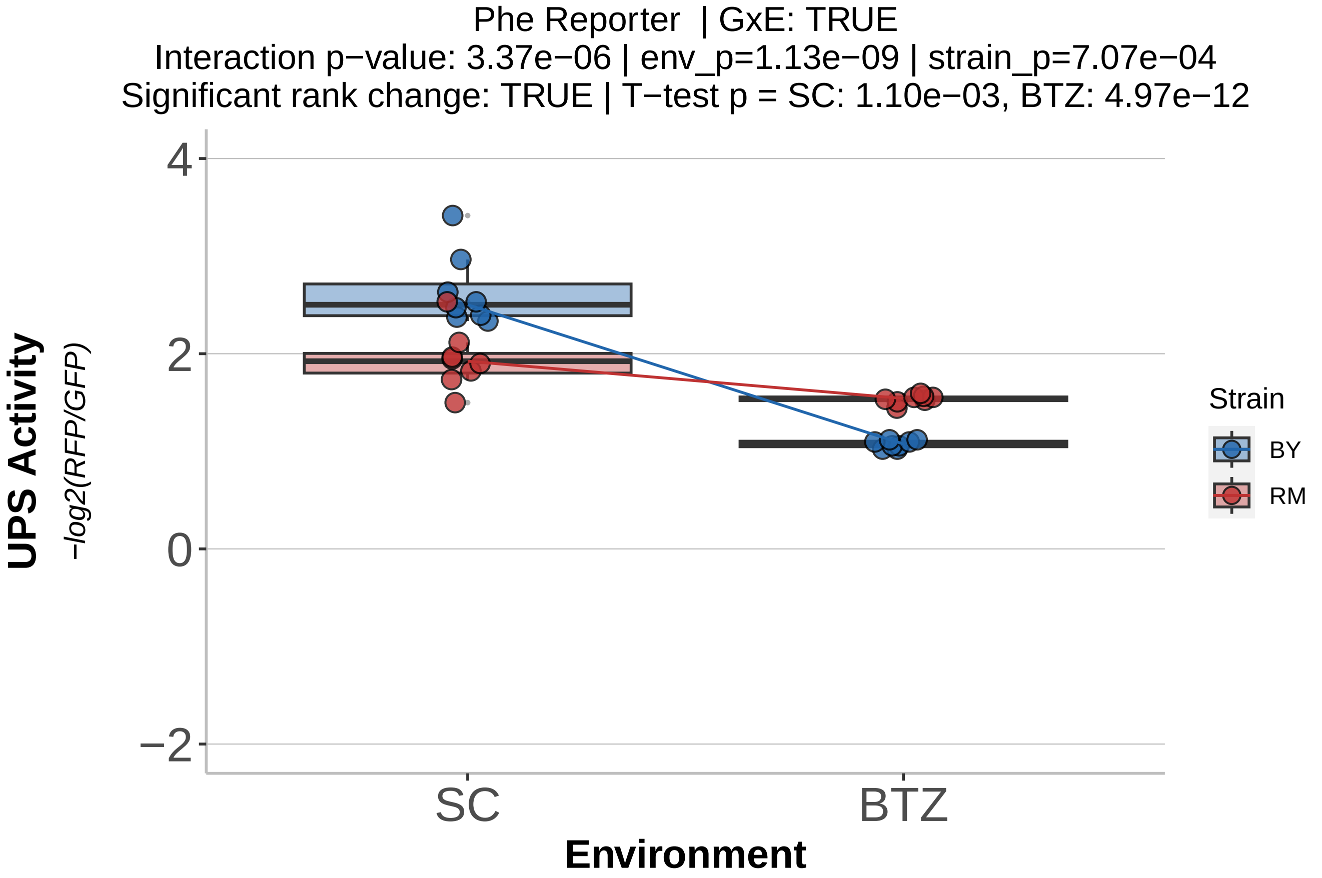
Phe Reporter | GxE: TRUE
Interaction p−value: 3.37e−06 | env_p=1.13e−09 | strain_p=7.07e−04
Significant rank change: TRUE | T−test p = SC: 1.10e−03, BTZ: 4.97e−12
UPS Activity
−log2(RFP/GFP)
4
2
0
−2
SC
BTZ
Environment
Strain
BY
RM

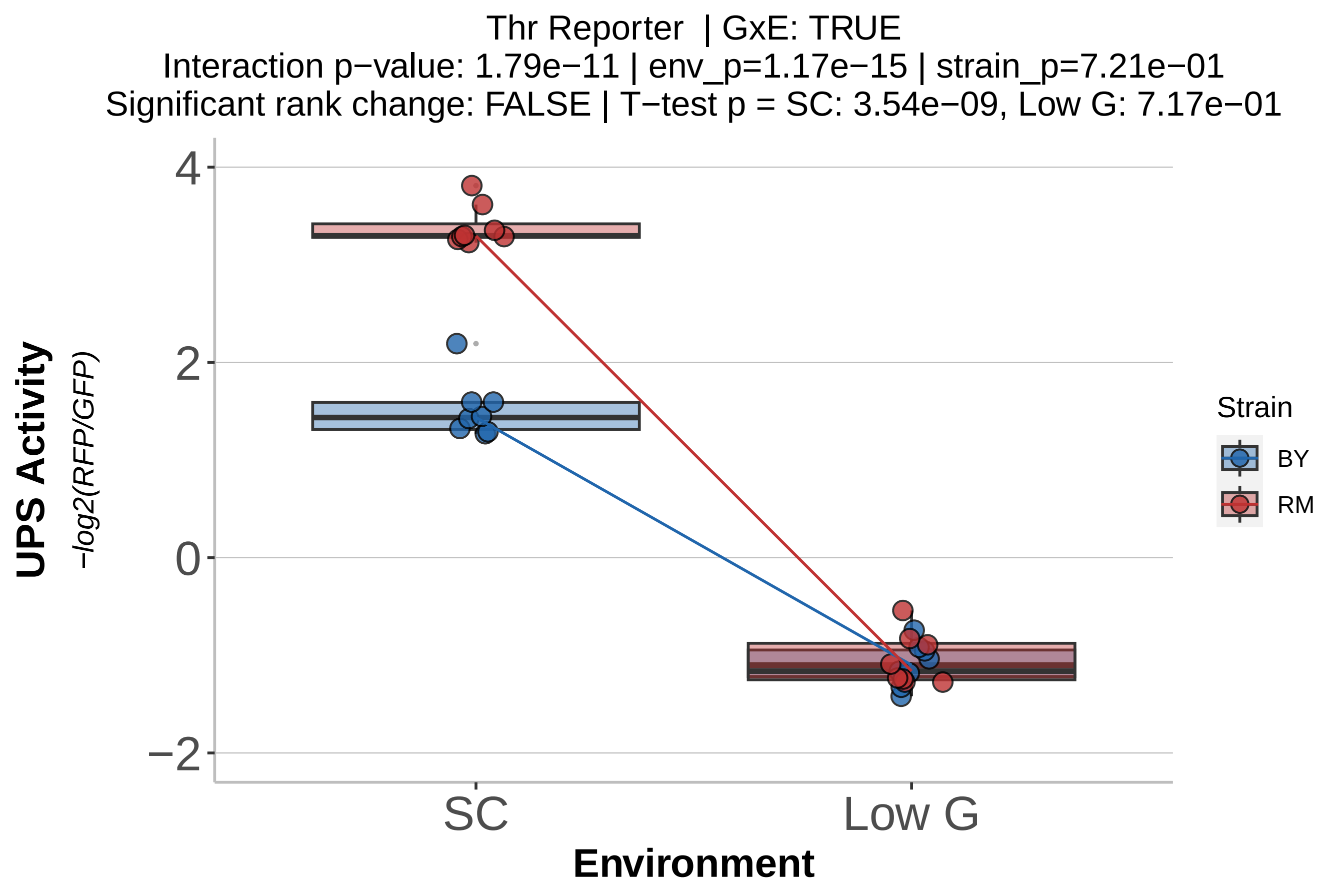
Thr Reporter | GxE: TRUE
Interaction p−value: 1.79e−11 | env_p=1.17e−15 | strain_p=7.21e−01
Significant rank change: FALSE | T−test p = SC: 3.54e−09, Low G: 7.17e−01
UPS Activity
−log2(RFP/GFP)
4
2
0
−2
SC
Low G
Environment
Strain
BY
RM

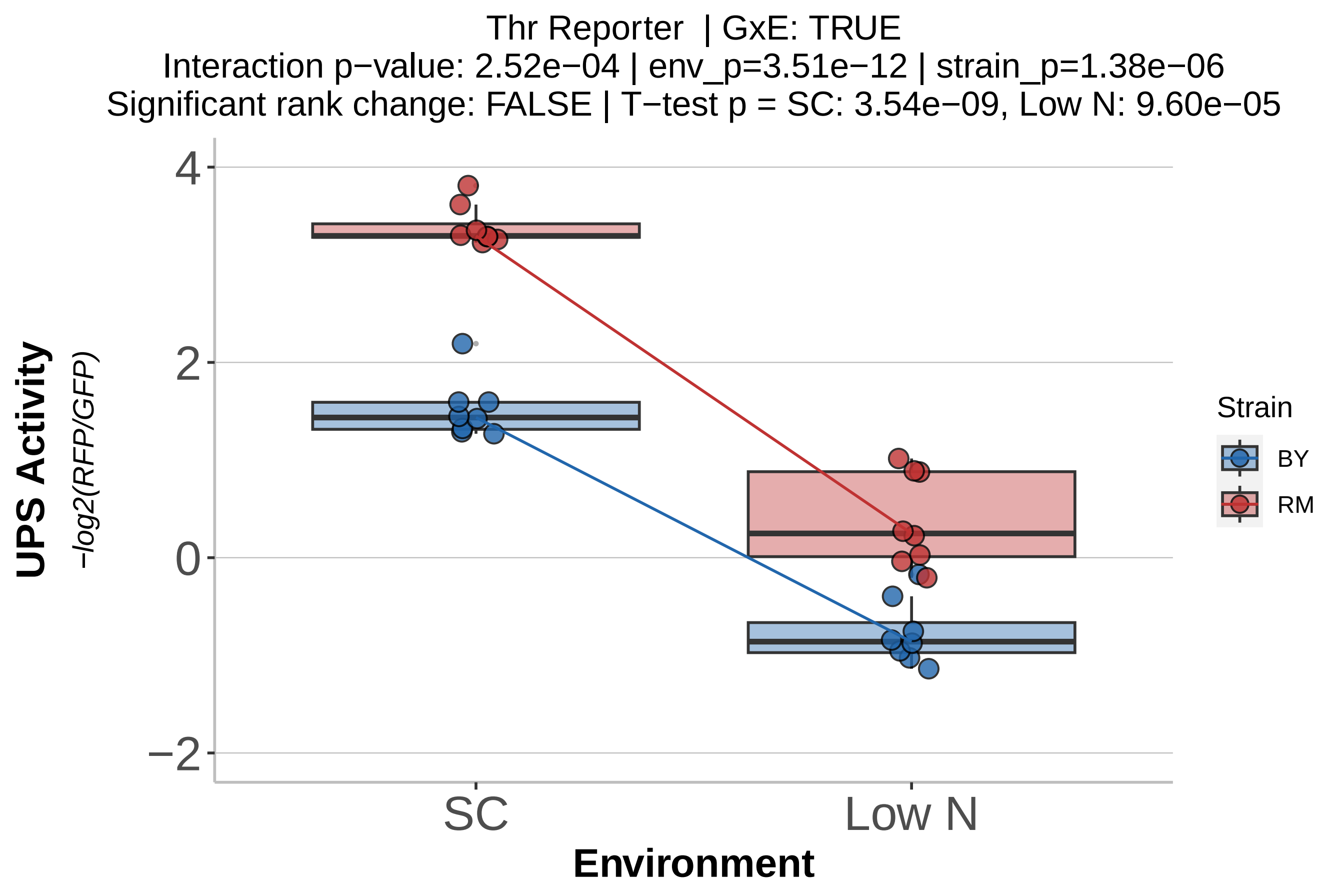
Thr Reporter | GxE: TRUE
Interaction p−value: 2.52e−04 | env_p=3.51e−12 | strain_p=1.38e−06
Significant rank change: FALSE | T−test p = SC: 3.54e−09, Low N: 9.60e−05
UPS Activity
−log2(RFP/GFP)
4
2
0
−2
SC
Low N
Environment
Strain
BY
RM

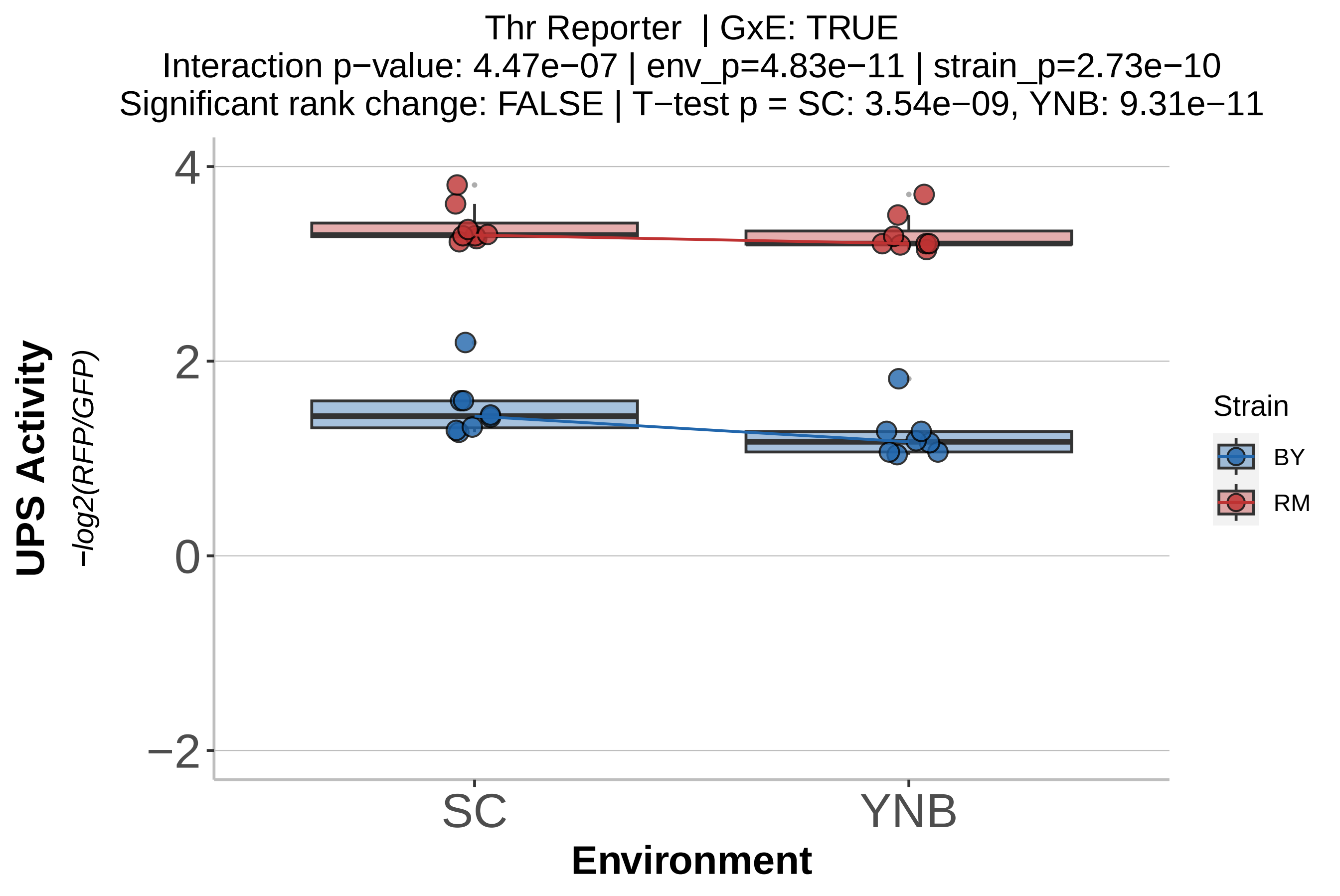
Thr Reporter | GxE: TRUE
Interaction p−value: 4.47e−07 | env_p=4.83e−11 | strain_p=2.73e−10
Significant rank change: FALSE | T−test p = SC: 3.54e−09, YNB: 9.31e−11
UPS Activity
−log2(RFP/GFP)
4
2
0
−2
SC
YNB
Environment
Strain
BY
RM

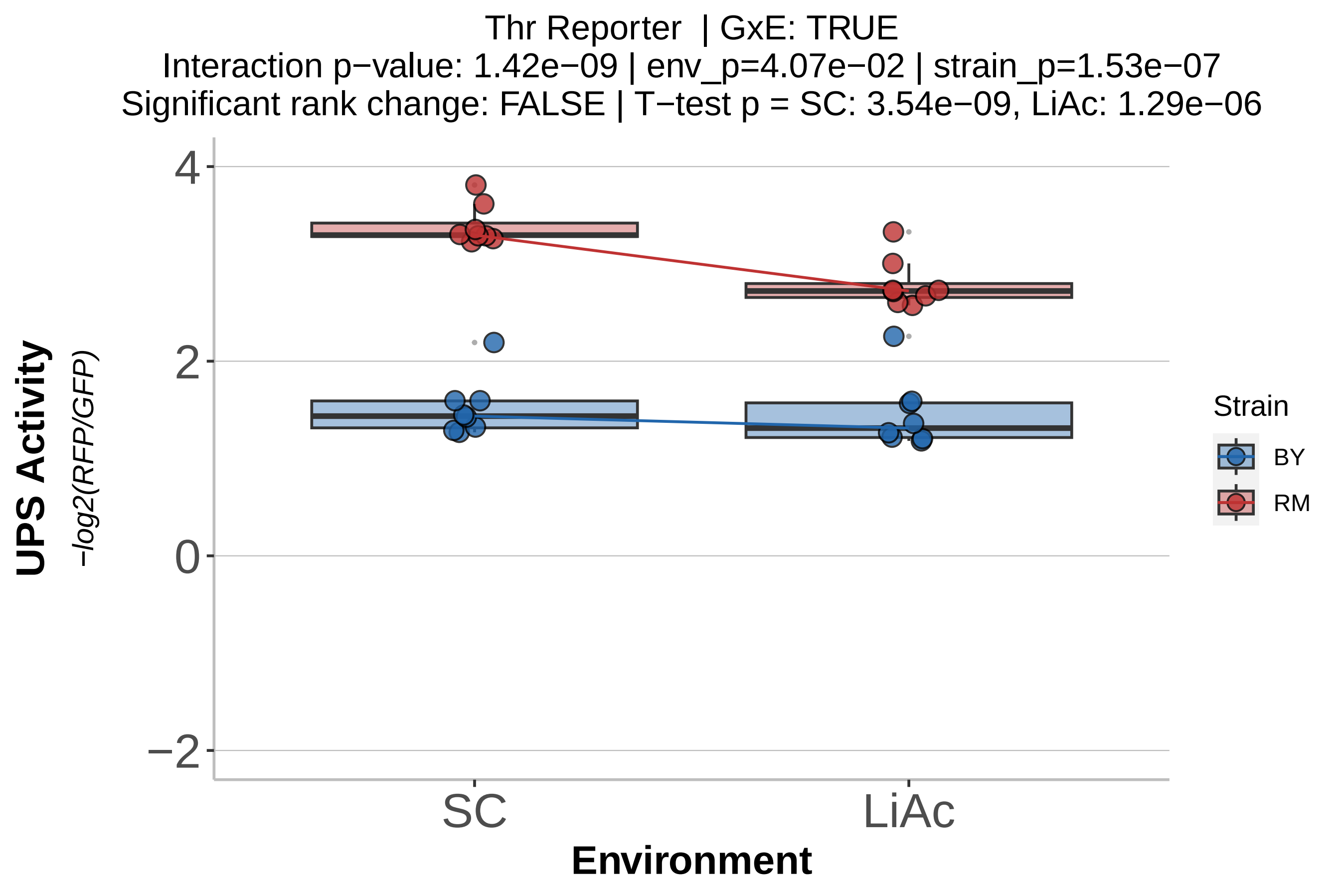
Thr Reporter | GxE: TRUE
Interaction p−value: 1.42e−09 | env_p=4.07e−02 | strain_p=1.53e−07
Significant rank change: FALSE | T−test p = SC: 3.54e−09, LiAc: 1.29e−06
UPS Activity
−log2(RFP/GFP)
4
2
0
−2
SC
LiAc
Environment
Strain
BY
RM

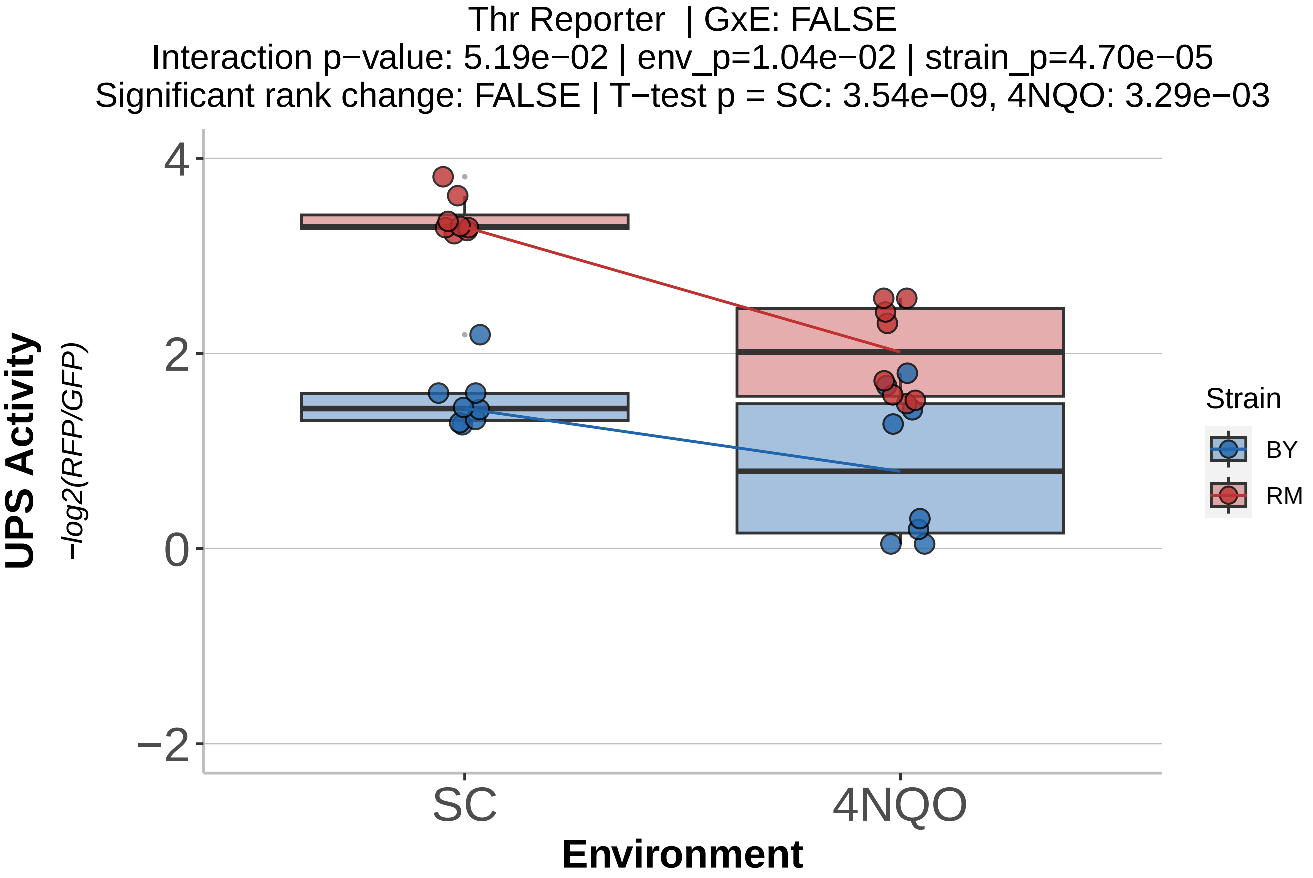
Thr Reporter | GxE: FALSE
Interaction p−value: 5.19e−02 | env_p=1.04e−02 | strain_p=4.70e−05
Significant rank change: FALSE | T−test p = SC: 3.54e−09, 4NQO: 3.29e−03
UPS Activity
−log2(RFP/GFP)
4
2
0
−2
SC
4NQO
Environment
Strain
BY
RM

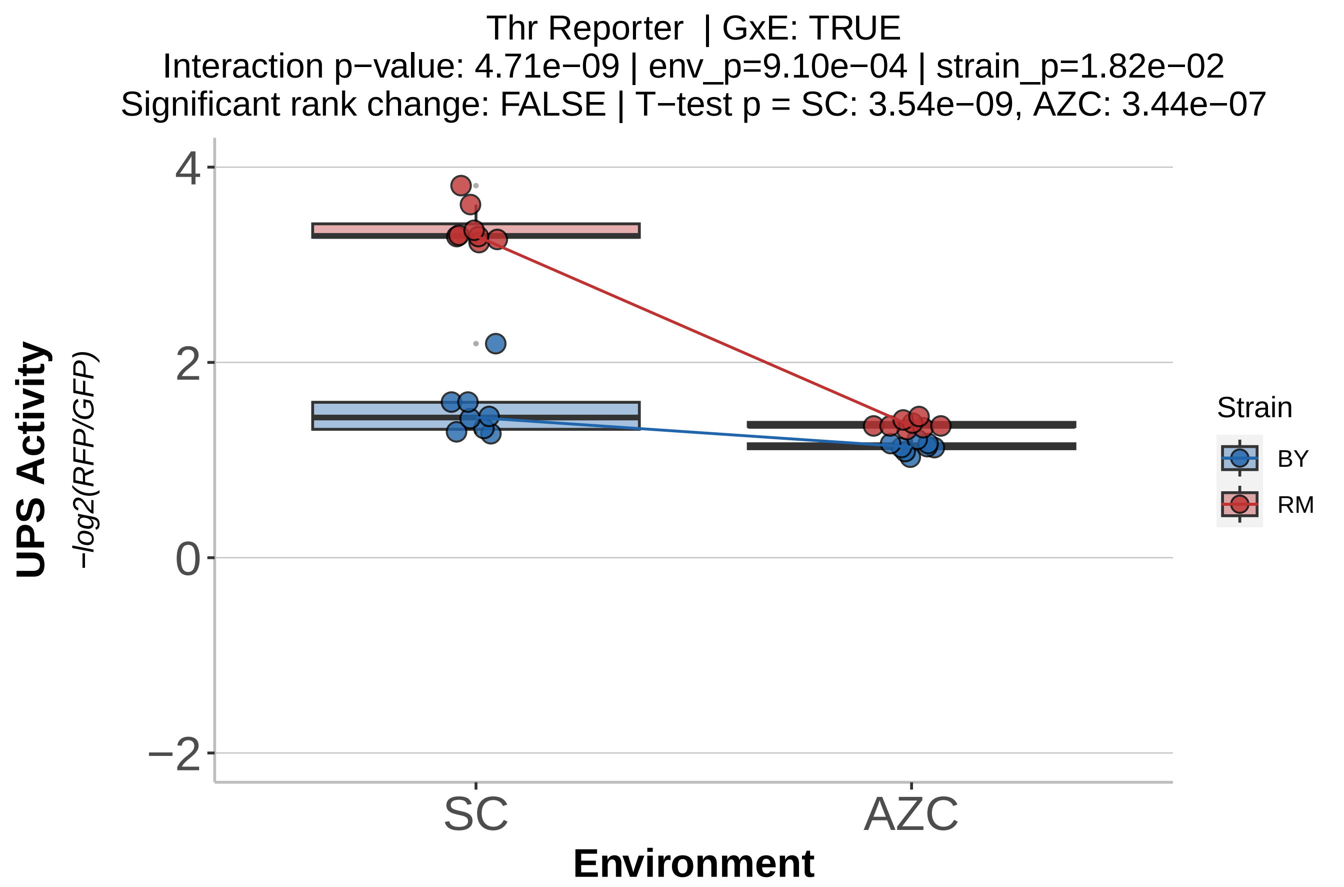
Thr Reporter | GxE: TRUE
Interaction p−value: 4.71e−09 | env_p=9.10e−04 | strain_p=1.82e−02
Significant rank change: FALSE | T−test p = SC: 3.54e−09, AZC: 3.44e−07
UPS Activity
−log2(RFP/GFP)
4
2
0
−2
SC
AZC
Environment
Strain
BY
RM

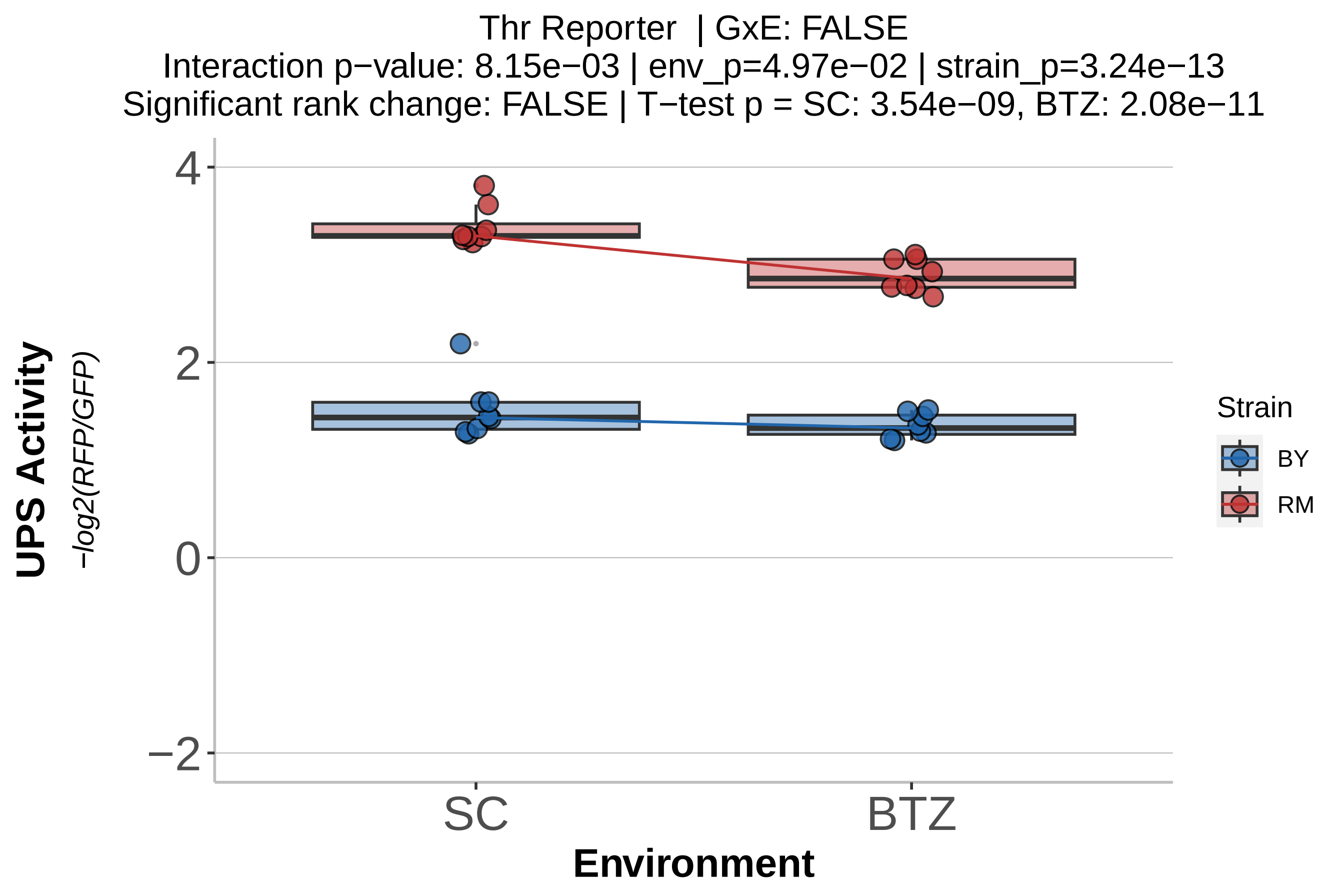
Thr Reporter | GxE: FALSE
Interaction p−value: 8.15e−03 | env_p=4.97e−02 | strain_p=3.24e−13
Significant rank change: FALSE | T−test p = SC: 3.54e−09, BTZ: 2.08e−11
UPS Activity
−log2(RFP/GFP)
4
2
0
−2
SC
BTZ
Environment
Strain
BY
RM

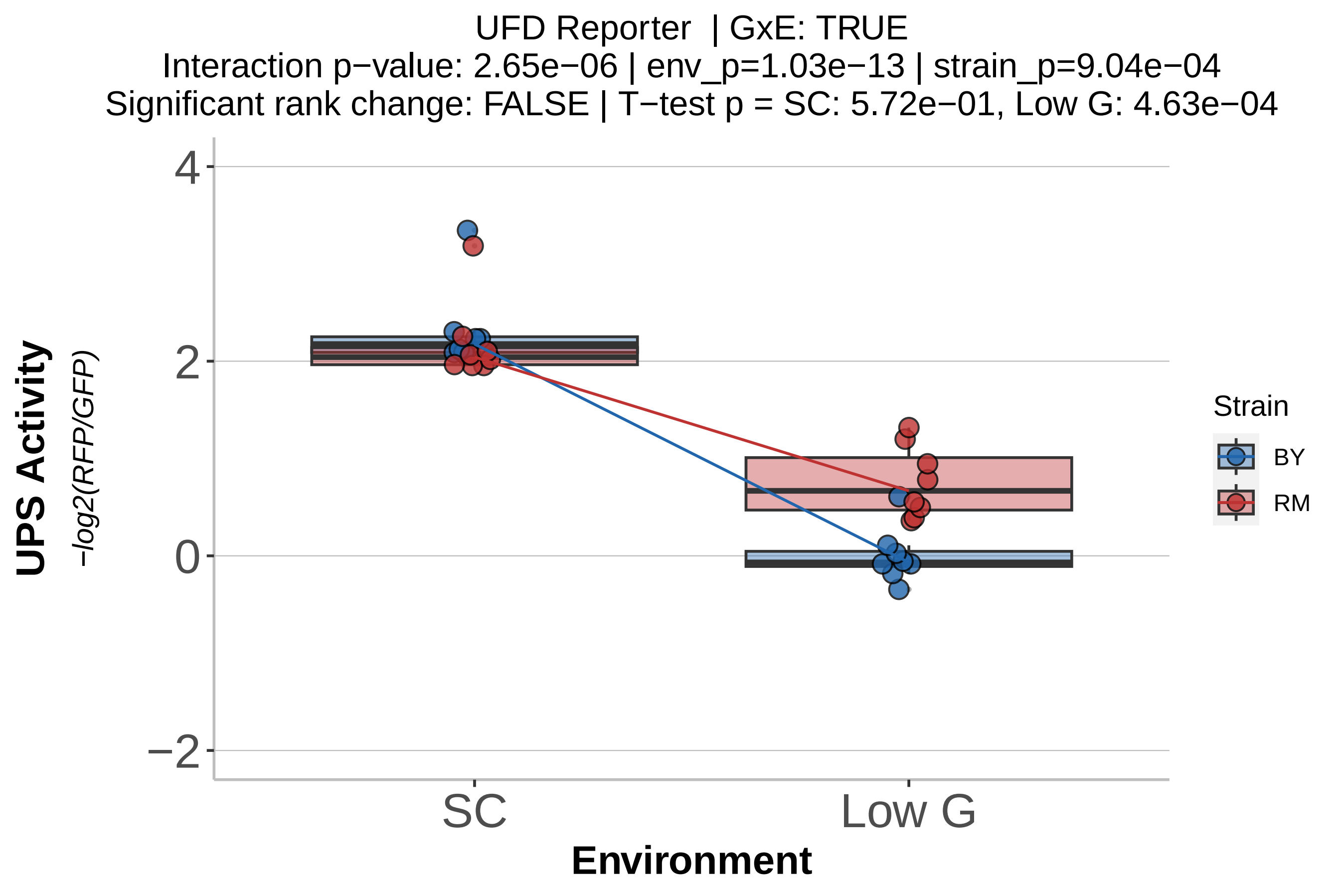
UFD Reporter | GxE: TRUE
Interaction p−value: 2.65e−06 | env_p=1.03e−13 | strain_p=9.04e−04
Significant rank change: FALSE | T−test p = SC: 5.72e−01, Low G: 4.63e−04
UPS Activity
−log2(RFP/GFP)
4
2
0
−2
SC
Low G
Environment
Strain
BY
RM

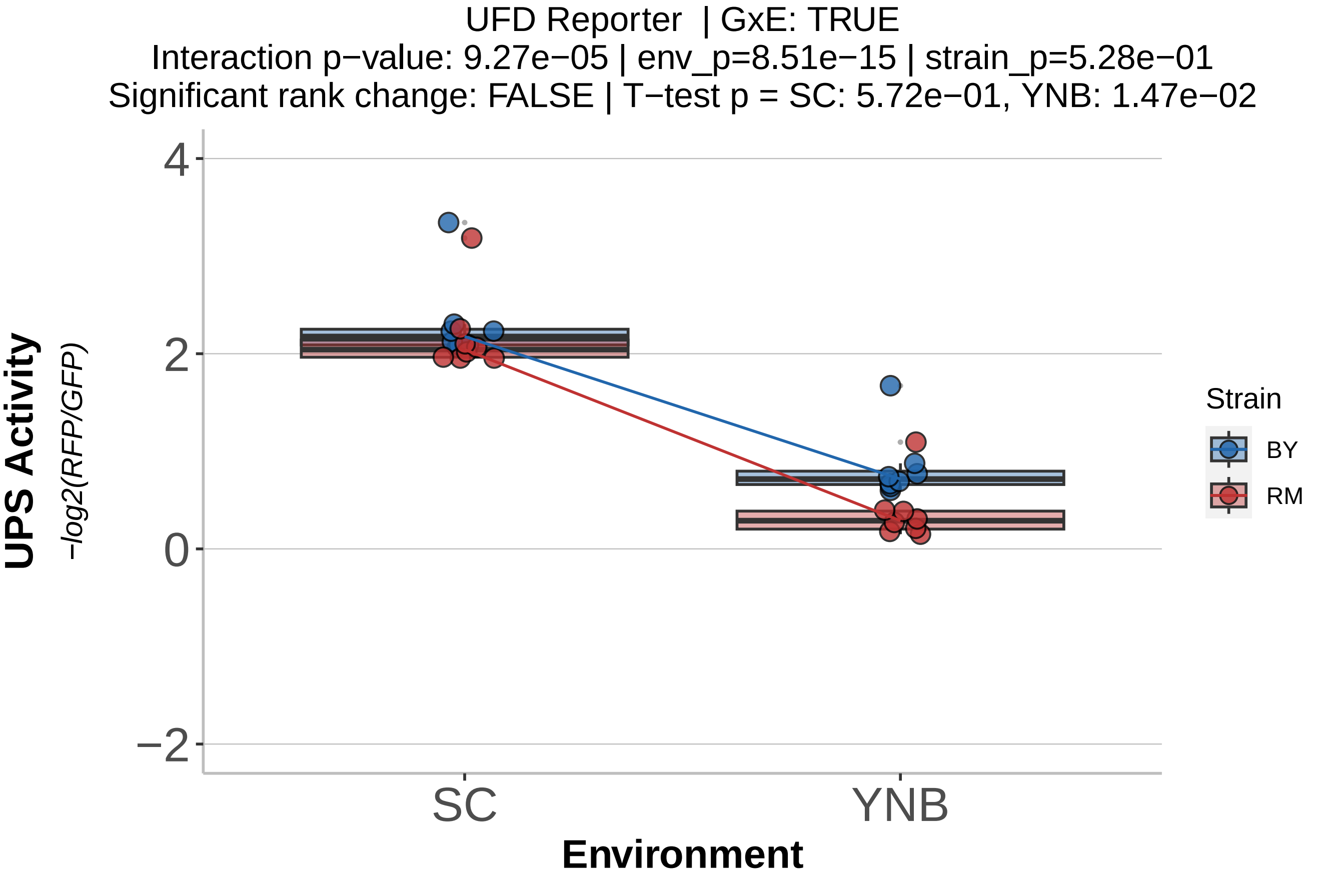

UFD Reporter | GxE: TRUE
Interaction p−value: 9.27e−05 | env_p=8.51e−15 | strain_p=5.28e−01
Significant rank change: FALSE | T−test p = SC: 5.72e−01, YNB: 1.47e−02
UPS Activity
−log2(RFP/GFP)
4
2
0
−2
SC
YNB
Environment
Strain
BY
RM

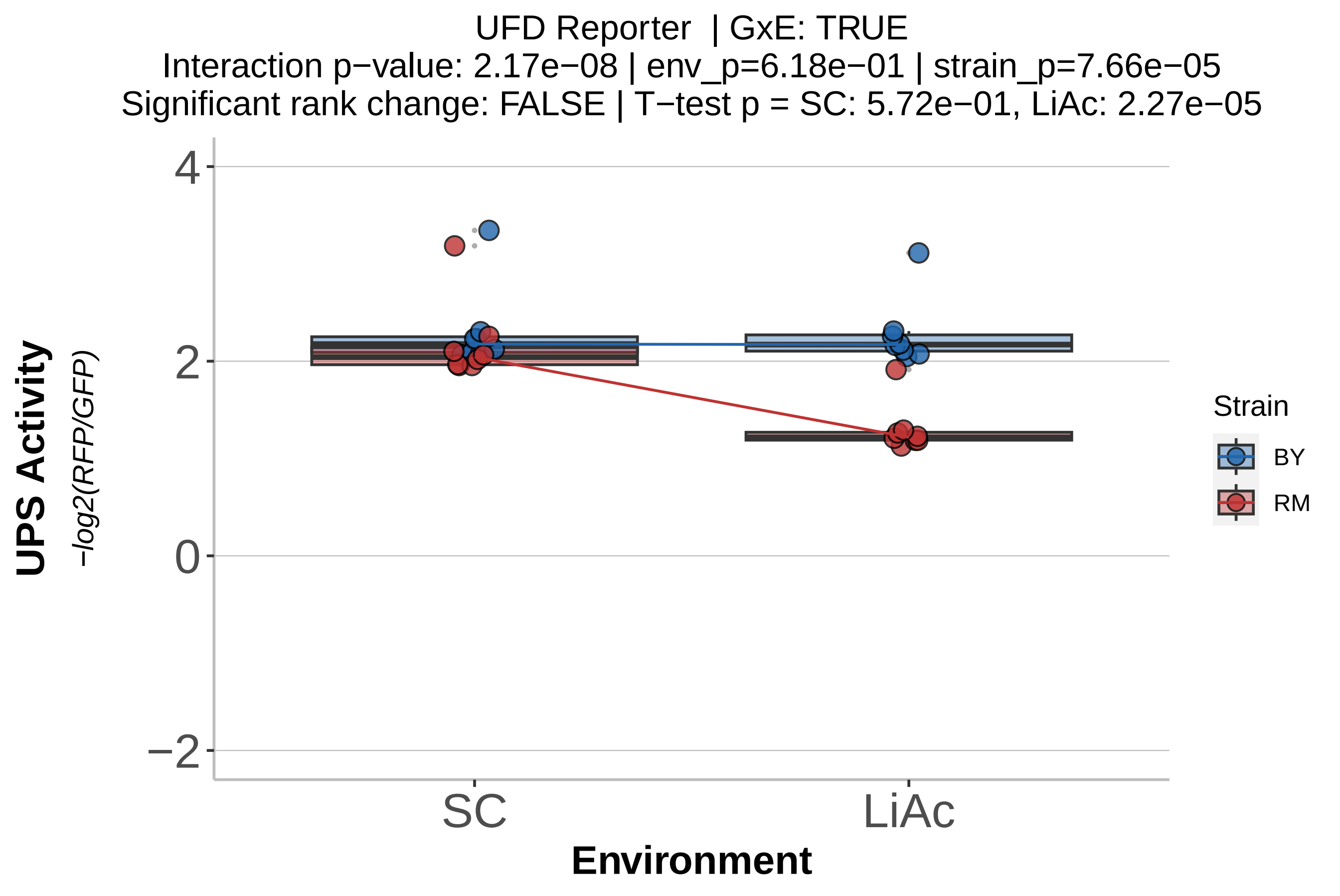

UFD Reporter | GxE: TRUE
Interaction p−value: 2.17e−08 | env_p=6.18e−01 | strain_p=7.66e−05
Significant rank change: FALSE | T−test p = SC: 5.72e−01, LiAc: 2.27e−05
UPS Activity
−log2(RFP/GFP)
4
2
0
−2
SC
LiAc
Environment
Strain
BY
RM

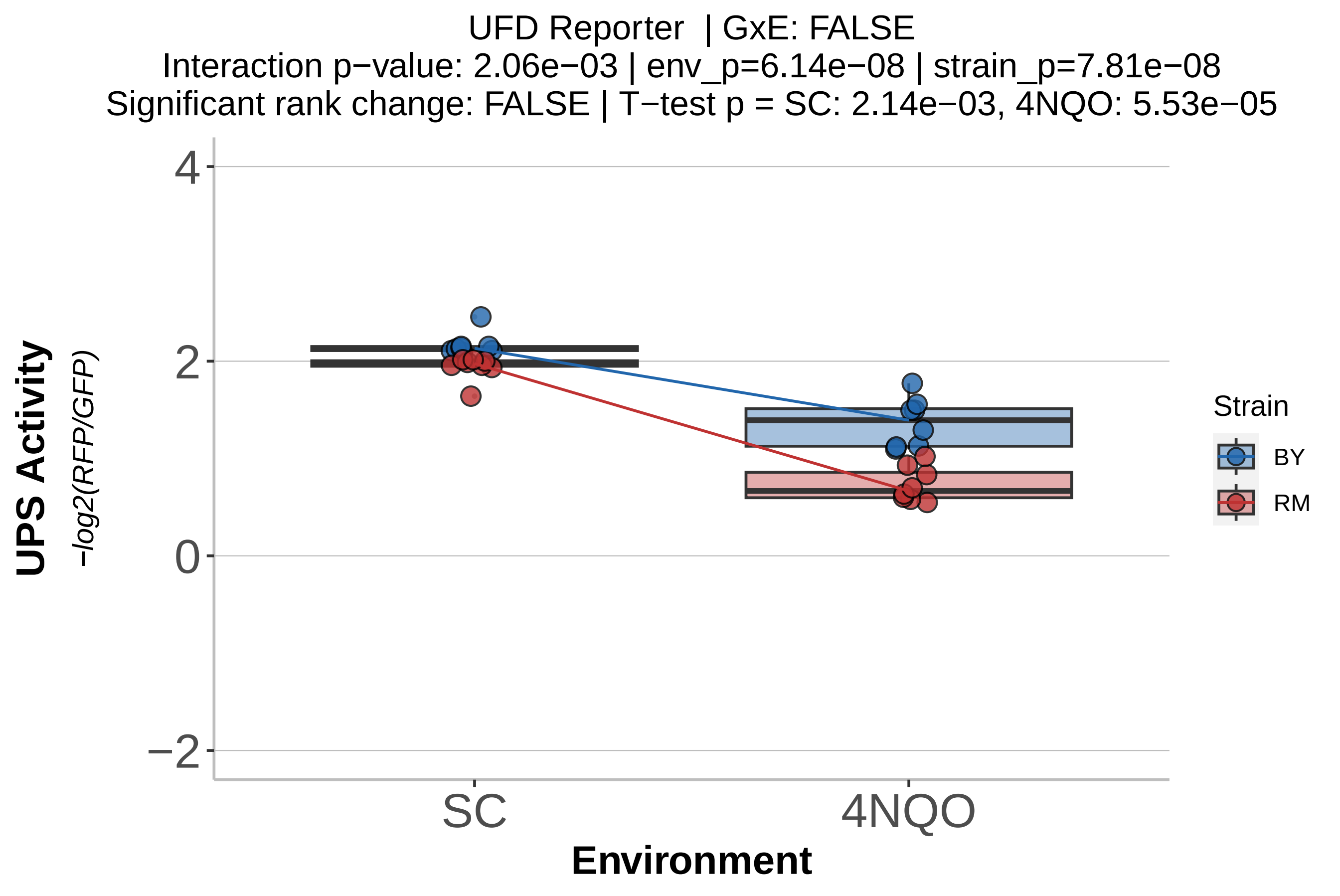

UFD Reporter | GxE: FALSE
Interaction p−value: 2.06e−03 | env_p=6.14e−08 | strain_p=7.81e−08
Significant rank change: FALSE | T−test p = SC: 2.14e−03, 4NQO: 5.53e−05
UPS Activity
−log2(RFP/GFP)
4
2
0
−2
SC
4NQO
Environment
Strain
BY
RM

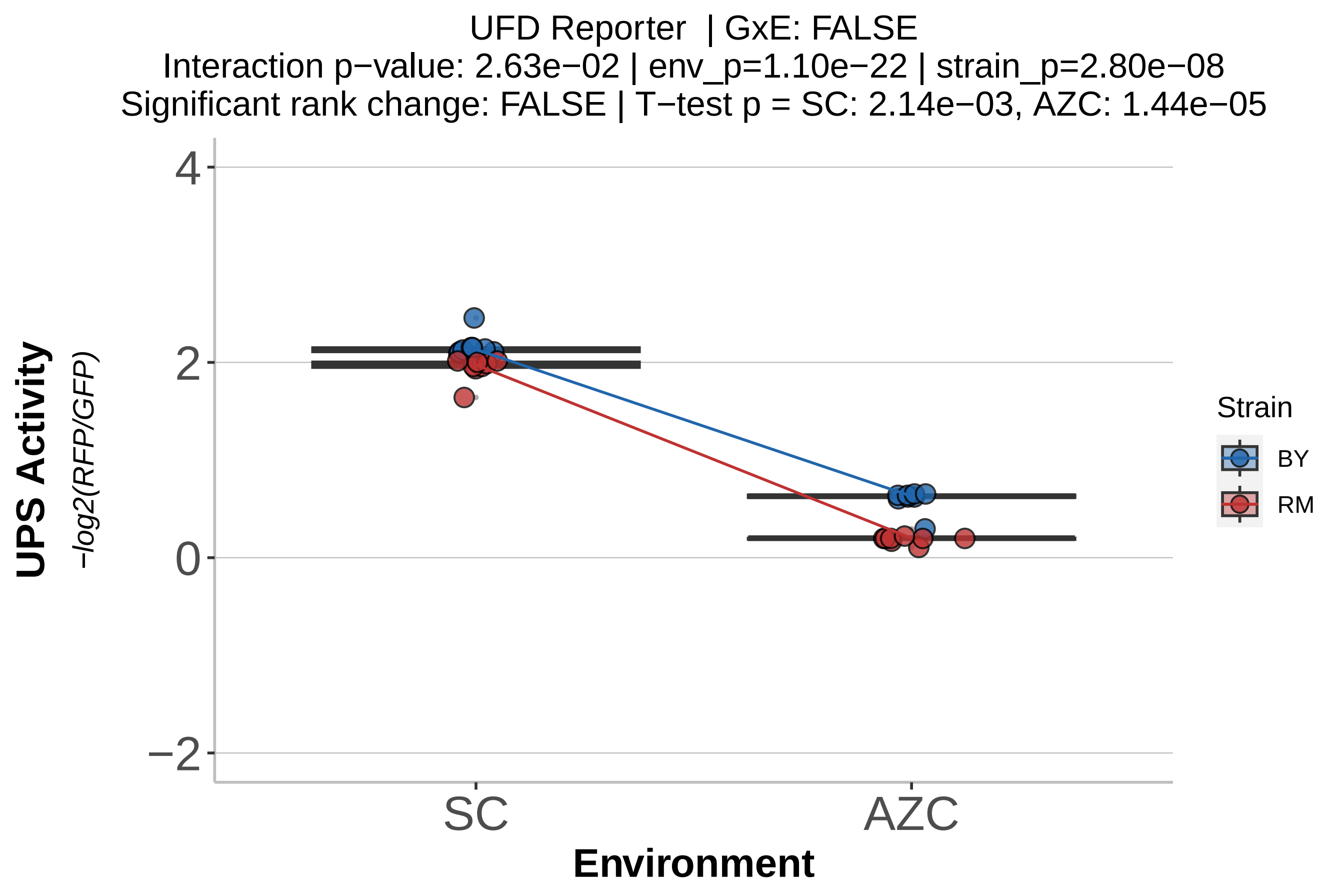

UFD Reporter | GxE: FALSE
Interaction p−value: 2.63e−02 | env_p=1.10e−22 | strain_p=2.80e−08
Significant rank change: FALSE | T−test p = SC: 2.14e−03, AZC: 1.44e−05
UPS Activity
−log2(RFP/GFP)
4
2
0
−2
SC
AZC
Environment
Strain
BY
RM

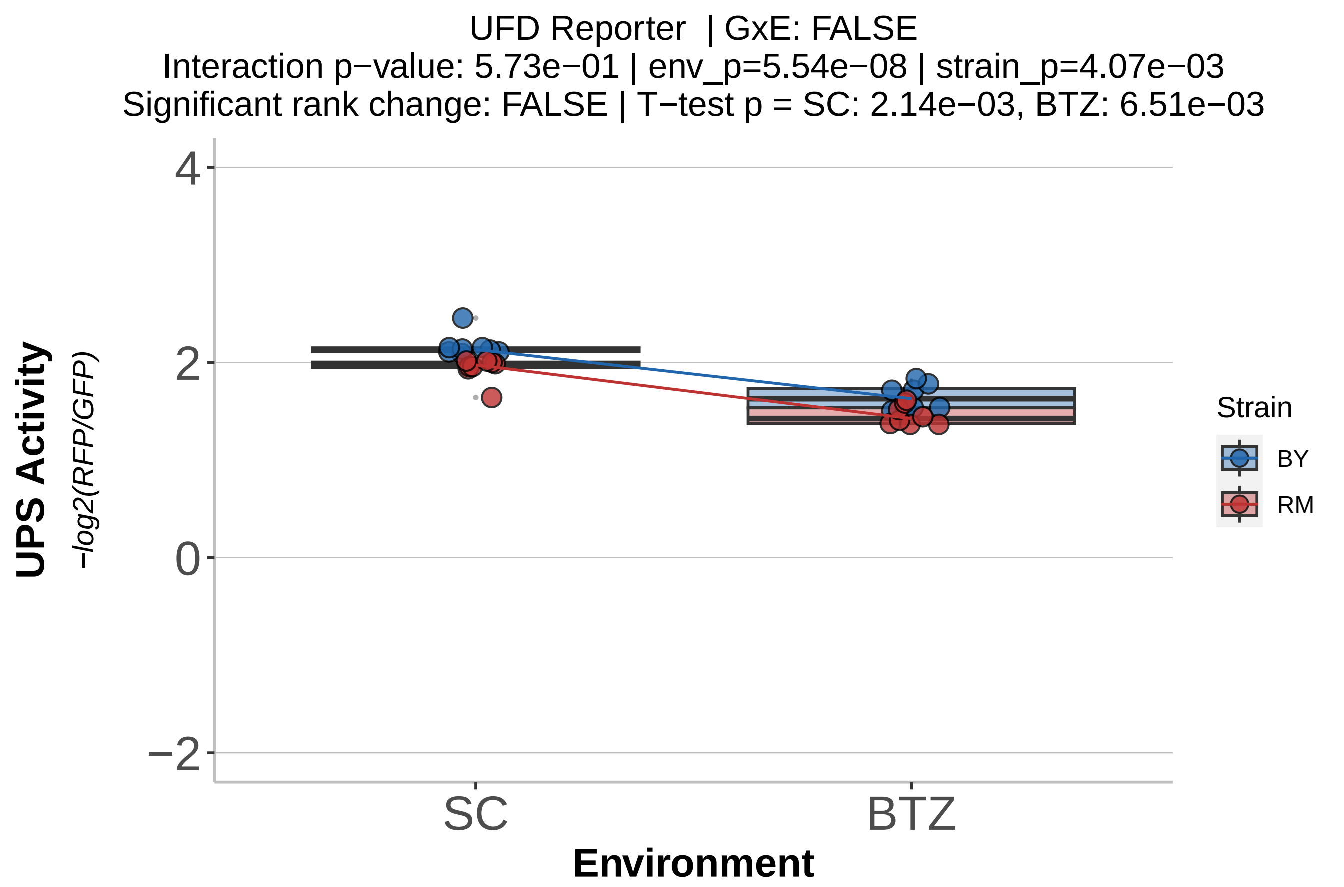
UFD Reporter | GxE: FALSE
Interaction p−value: 5.73e−01 | env_p=5.54e−08 | strain_p=4.07e−03
Significant rank change: FALSE | T−test p = SC: 2.14e−03, BTZ: 6.51e−03
UPS Activity
−log2(RFP/GFP)
4
2
0
−2
SC
BTZ
Environment
Strain
BY
RM

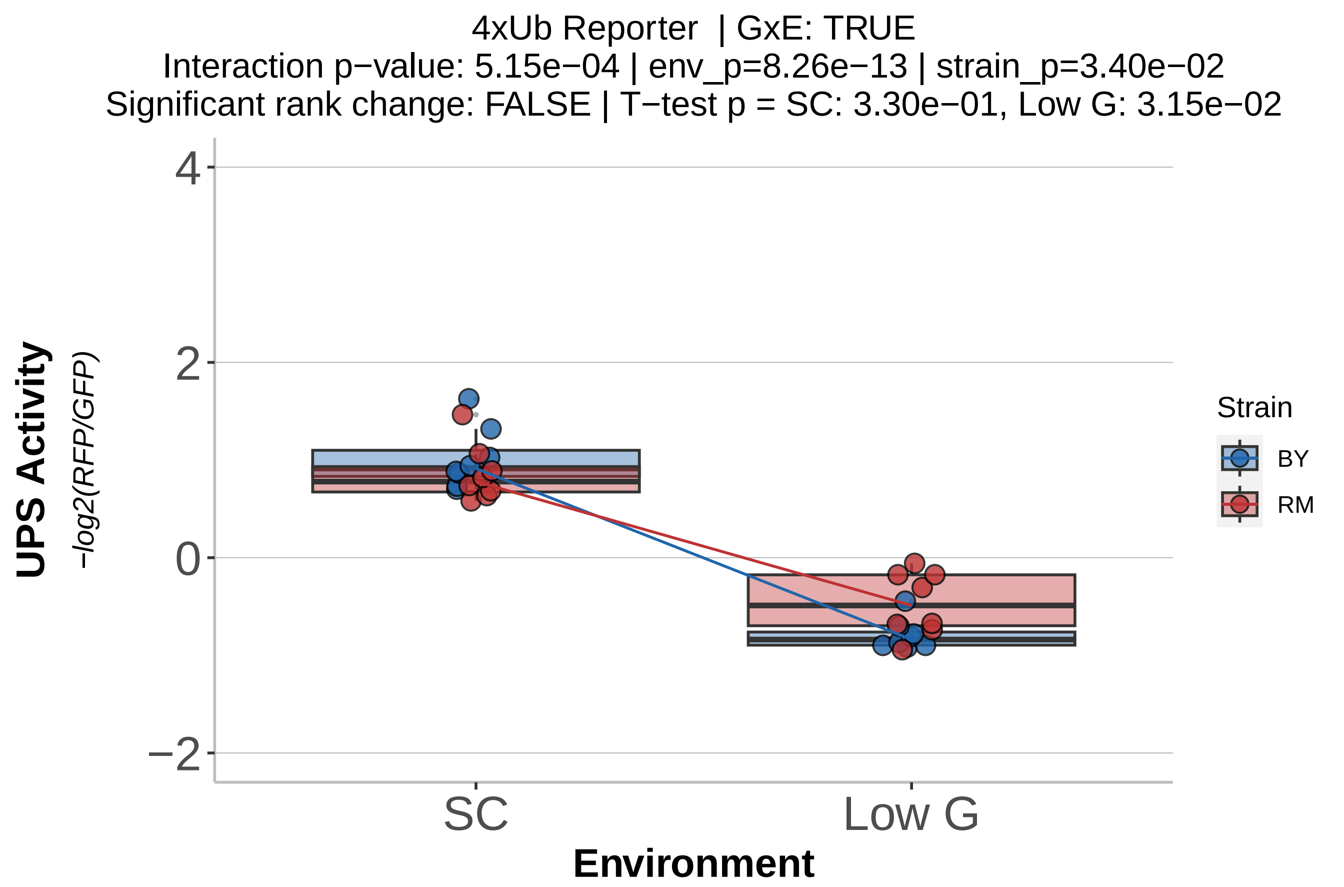
4xUb Reporter | GxE: TRUE
Interaction p−value: 5.15e−04 | env_p=8.26e−13 | strain_p=3.40e−02
Significant rank change: FALSE | T−test p = SC: 3.30e−01, Low G: 3.15e−02
UPS Activity
−log2(RFP/GFP)
4
2
0
−2
SC
Low G
Environment
Strain
BY
RM

4xUb Reporter | GxE: TRUE
Interaction p−value: 1.52e−07 | env_p=3.68e−10 | strain_p=2.87e−01
Significant rank change: FALSE | T−test p = SC: 3.30e−01, YNB: 1.65e−03

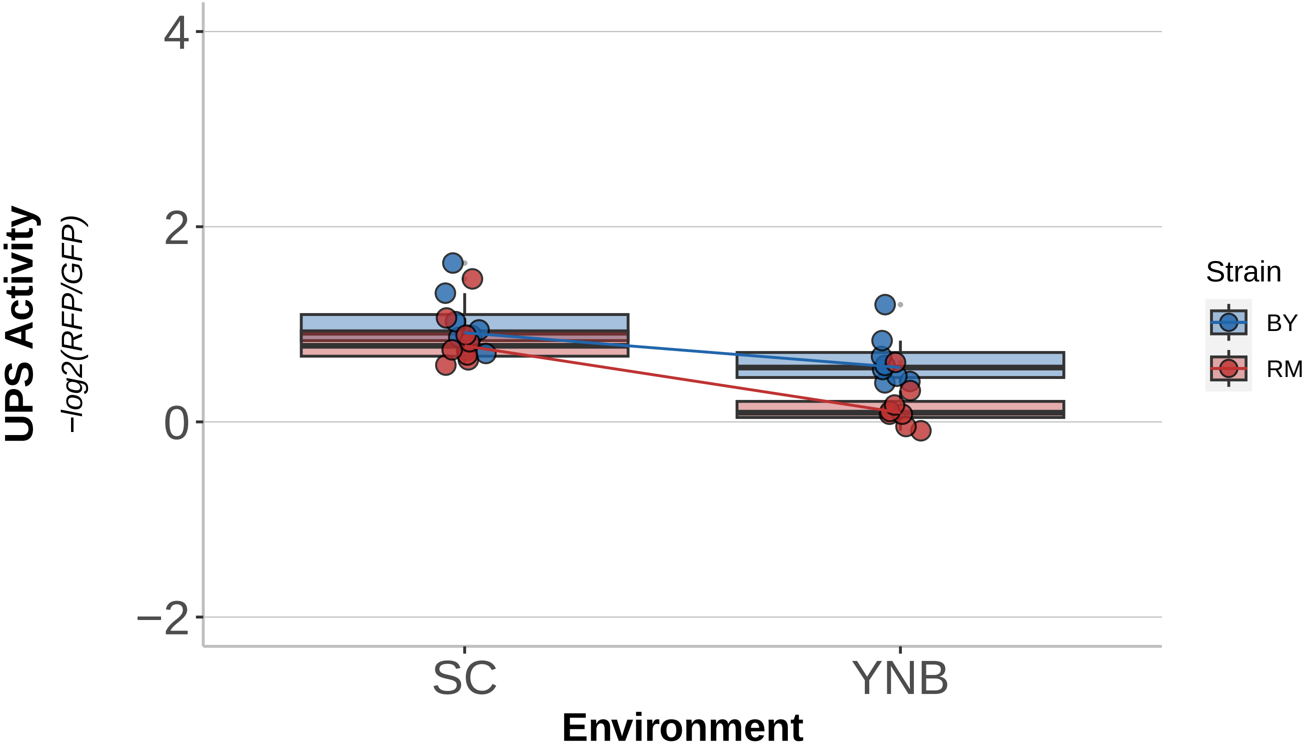

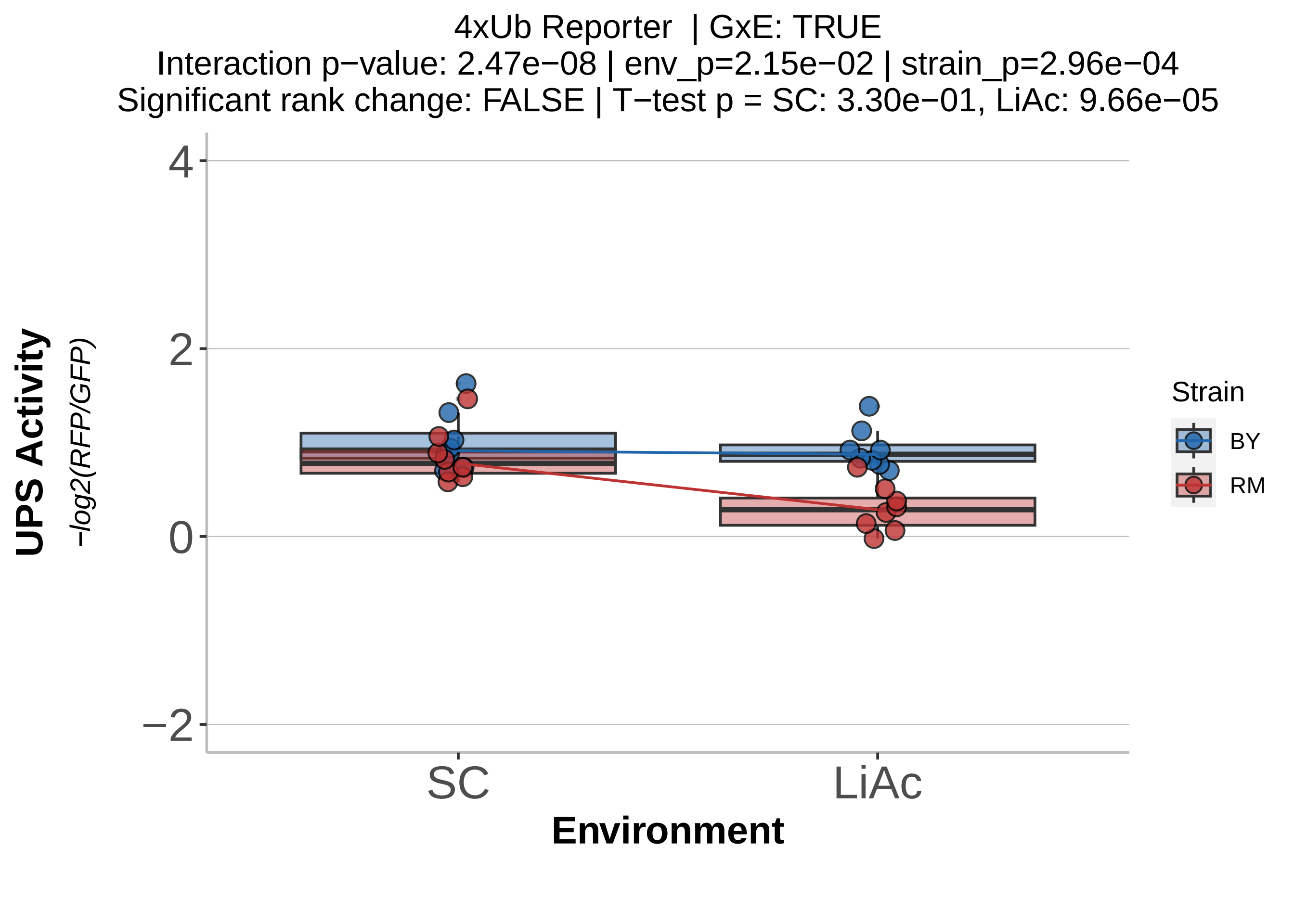
4xUb Reporter | GxE: TRUE
Interaction p−value: 2.47e−08 | env_p=2.15e−02 | strain_p=2.96e−04
Significant rank change: FALSE | T−test p = SC: 3.30e−01, LiAc: 9.66e−05
UPS Activity
−log2(RFP/GFP)
4
2
0
−2
SC
LiAc
Environment
Strain
BY
RM

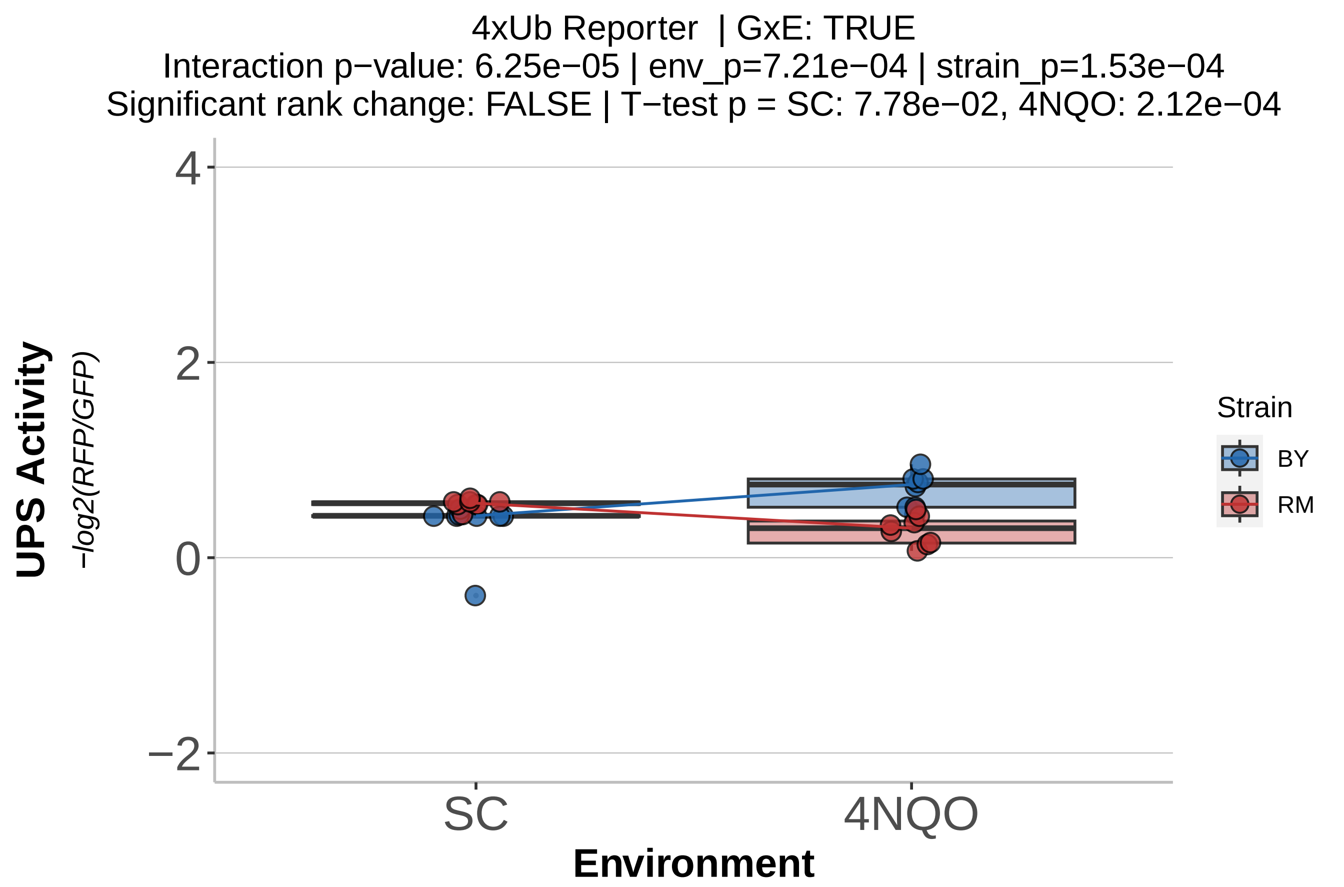

4xUb Reporter | GxE: TRUE
Interaction p−value: 6.25e−05 | env_p=7.21e−04 | strain_p=1.53e−04
Significant rank change: FALSE | T−test p = SC: 7.78e−02, 4NQO: 2.12e−04
UPS Activity
−log2(RFP/GFP)
4
2
0
−2
SC
4NQO
Environment
Strain
BY
RM

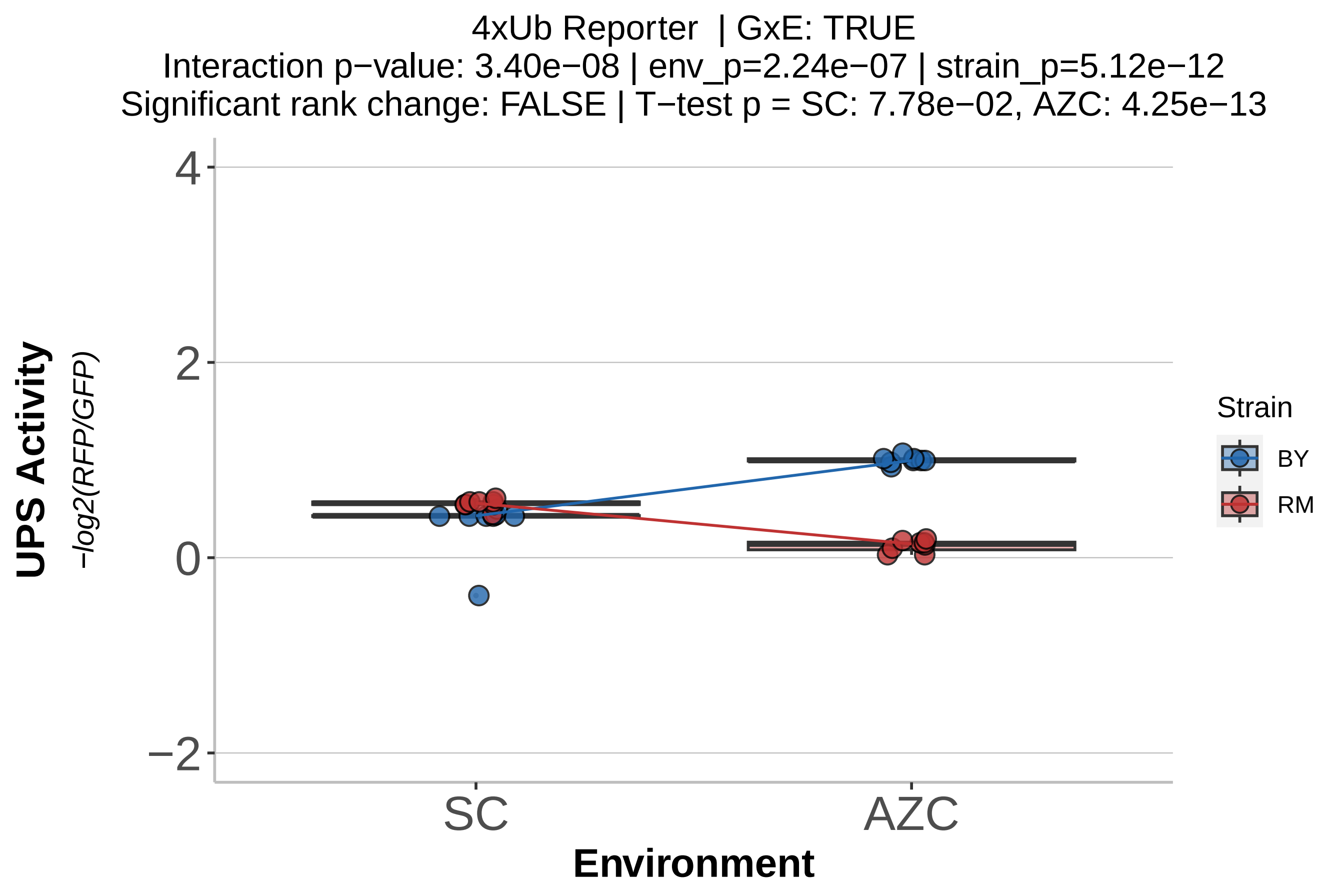
4xUb Reporter | GxE: TRUE
Interaction p−value: 3.40e−08 | env_p=2.24e−07 | strain_p=5.12e−12
Significant rank change: FALSE | T−test p = SC: 7.78e−02, AZC: 4.25e−13
UPS Activity
−log2(RFP/GFP)
4
2
0
−2
SC
AZC
Environment
Strain
BY
RM

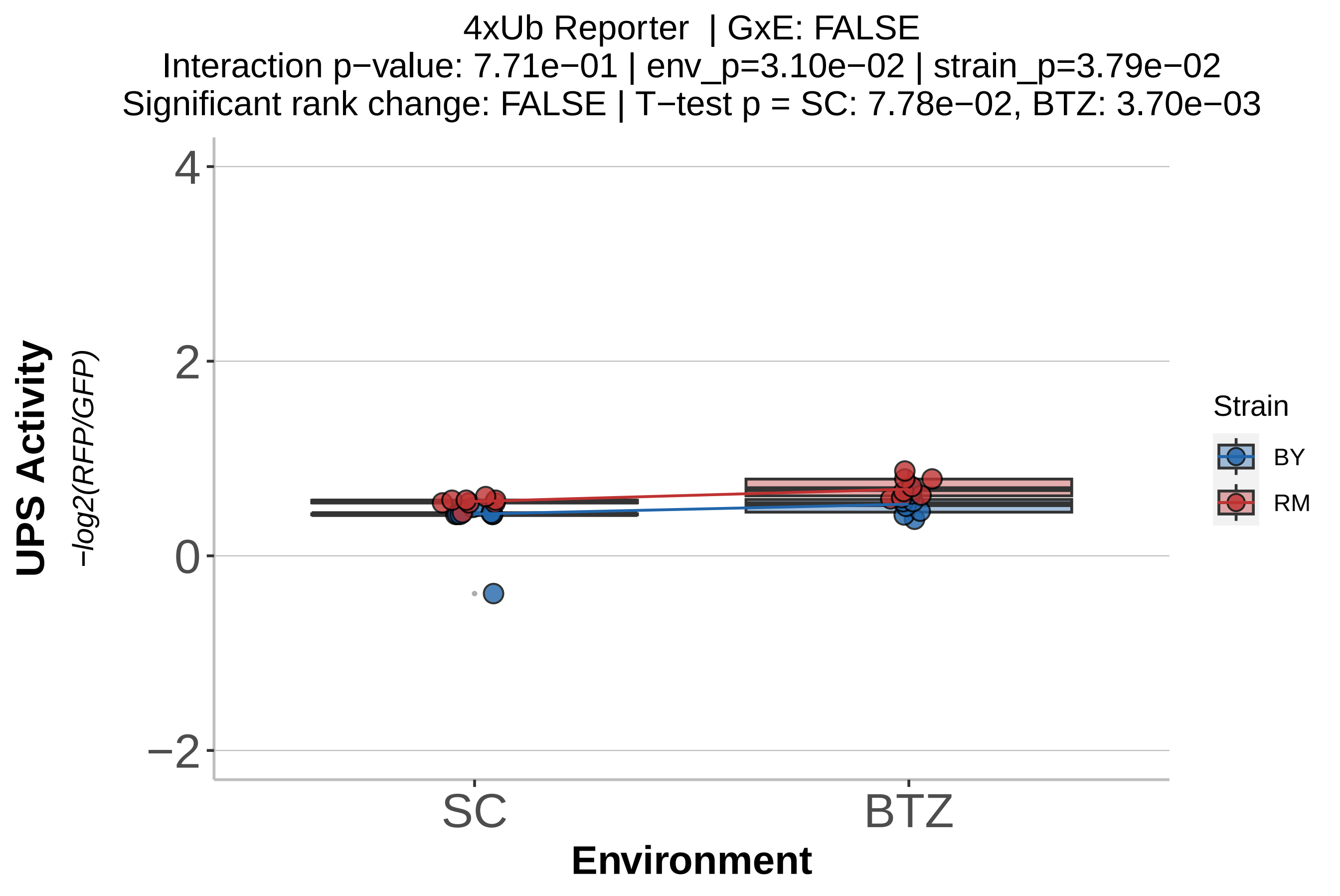
4xUb Reporter | GxE: FALSE
Interaction p−value: 7.71e−01 | env_p=3.10e−02 | strain_p=3.79e−02
Significant rank change: FALSE | T−test p = SC: 7.78e−02, BTZ: 3.70e−03
UPS Activity
−log2(RFP/GFP)
4
2
0
−2
SC
BTZ
Environment
Strain
BY
RM

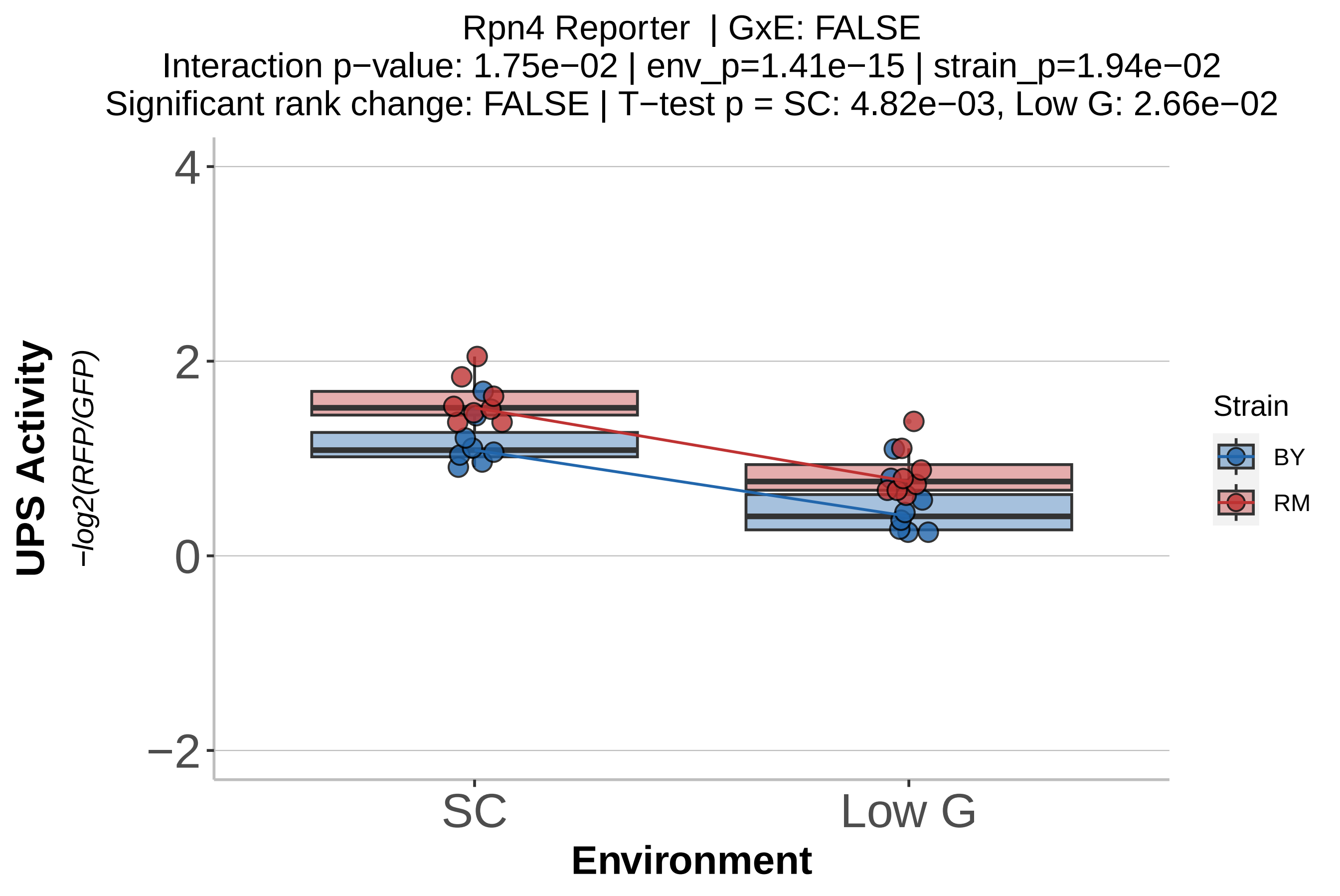

Rpn4 Reporter | GxE: FALSE
Interaction p−value: 1.75e−02 | env_p=1.41e−15 | strain_p=1.94e−02
Significant rank change: FALSE | T−test p = SC: 4.82e−03, Low G: 2.66e−02
UPS Activity
−log2(RFP/GFP)
4
2
0
−2
SC
Low G
Environment
Strain
BY
RM

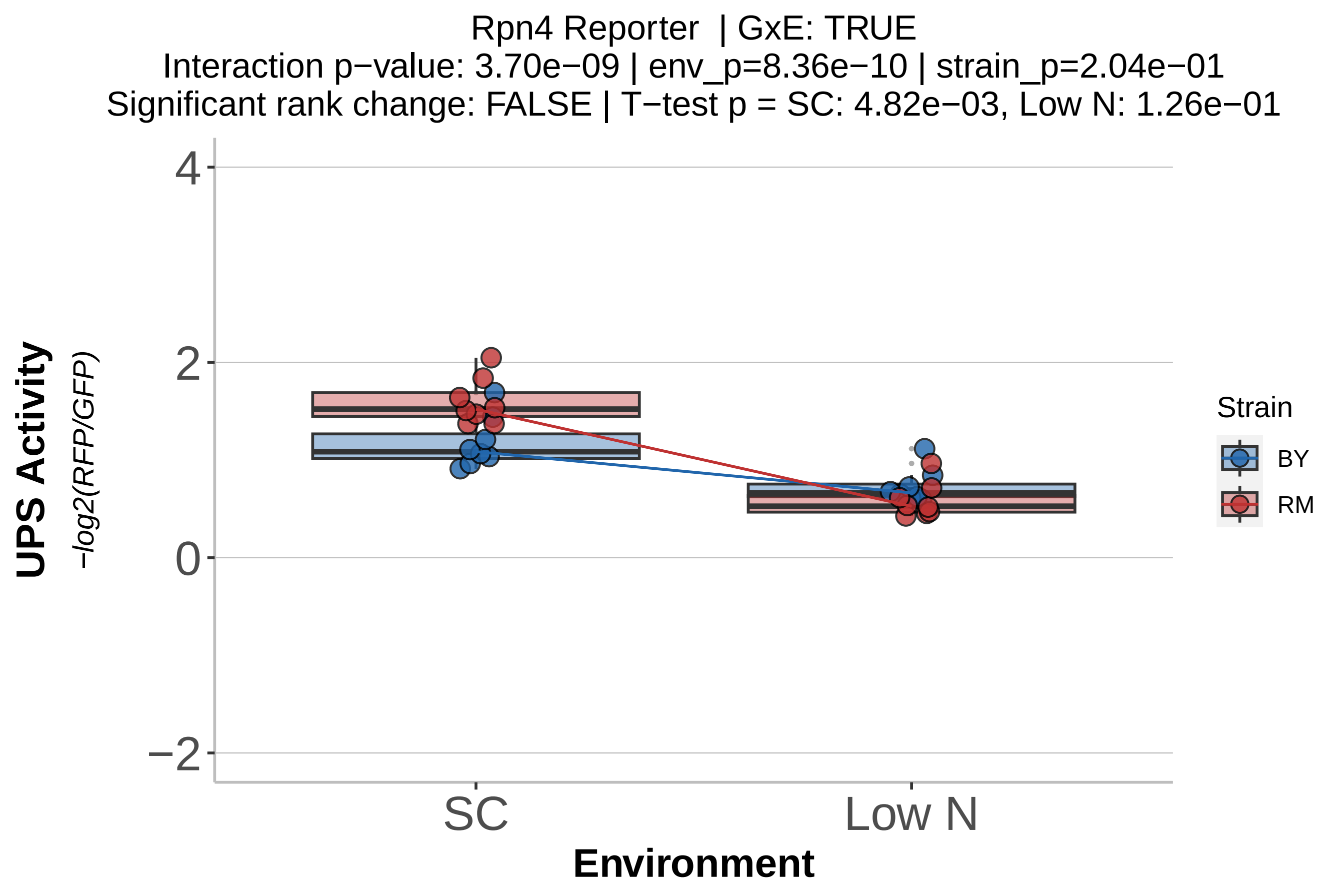
Rpn4 Reporter | GxE: TRUE
Interaction p−value: 3.70e−09 | env_p=8.36e−10 | strain_p=2.04e−01
Significant rank change: FALSE | T−test p = SC: 4.82e−03, Low N: 1.26e−01
UPS Activity
−log2(RFP/GFP)
4
2
0
−2
SC
Low N
Environment
Strain
BY
RM

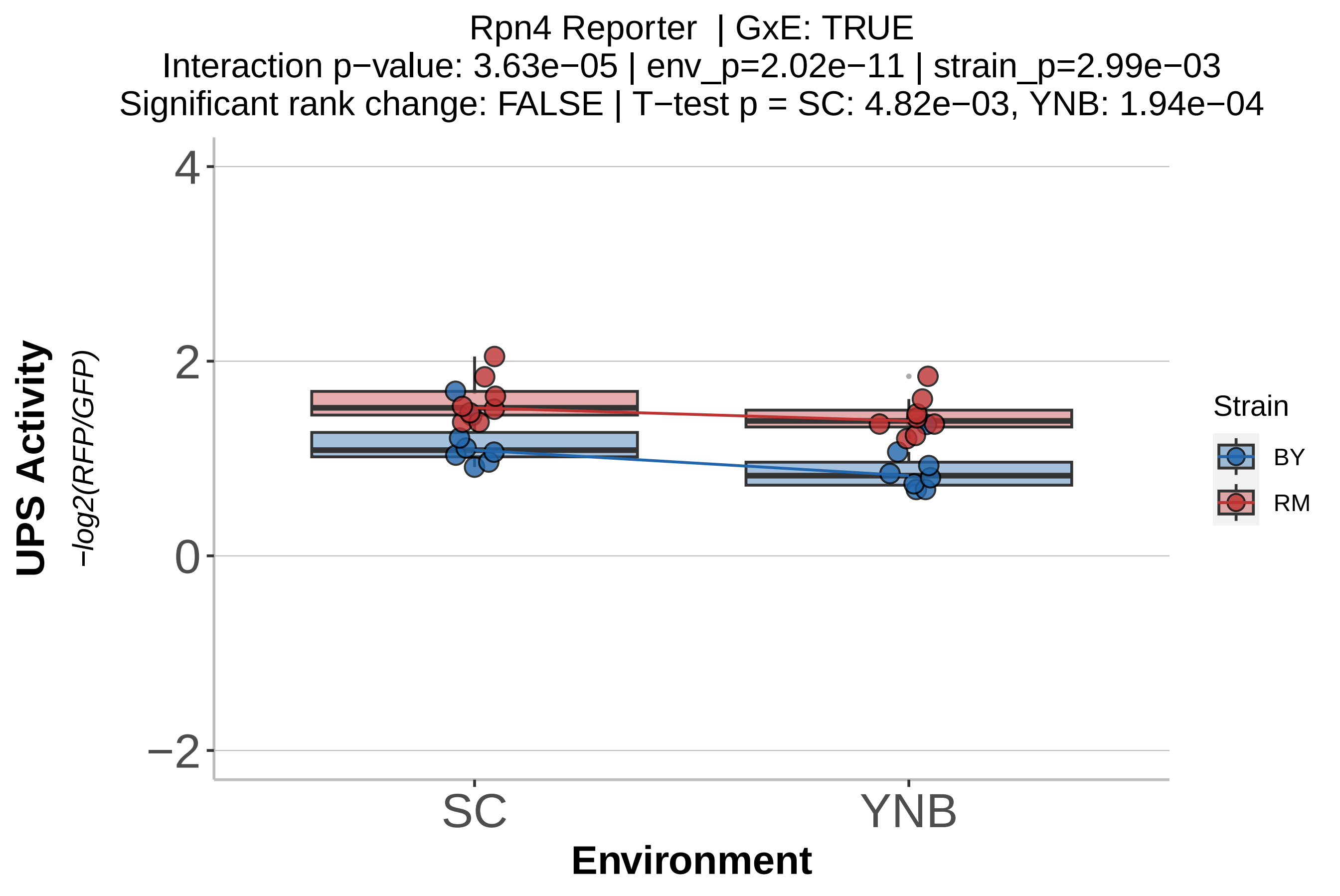
Rpn4 Reporter | GxE: TRUE
Interaction p−value: 3.63e−05 | env_p=2.02e−11 | strain_p=2.99e−03
Significant rank change: FALSE | T−test p = SC: 4.82e−03, YNB: 1.94e−04
UPS Activity
−log2(RFP/GFP)
4
2
0
−2
SC
YNB
Environment
Strain
BY
RM

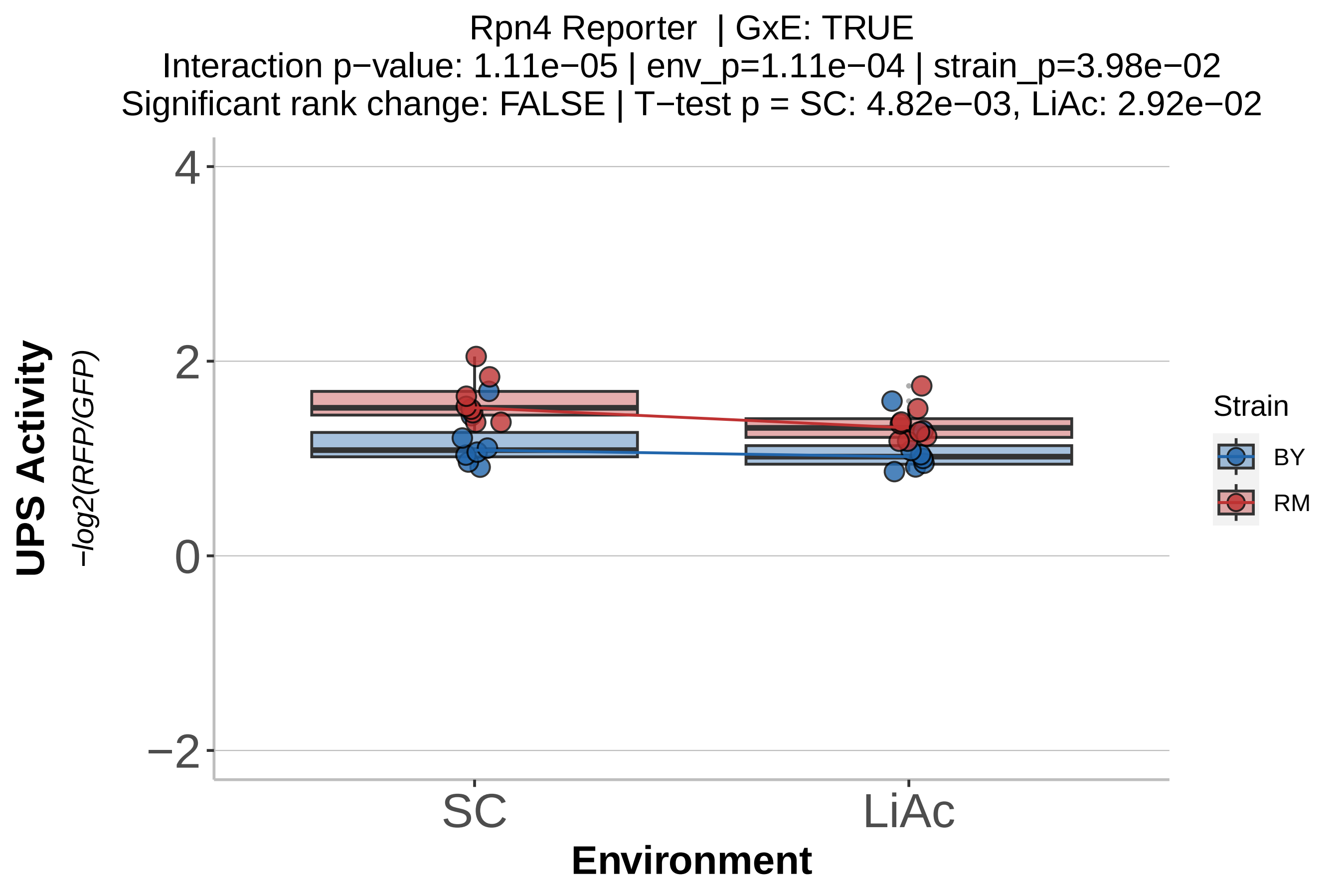

Rpn4 Reporter | GxE: TRUE
Interaction p−value: 1.11e−05 | env_p=1.11e−04 | strain_p=3.98e−02
Significant rank change: FALSE | T−test p = SC: 4.82e−03, LiAc: 2.92e−02
UPS Activity
−log2(RFP/GFP)
4
2
0
−2
SC
LiAc
Environment
Strain
BY
RM

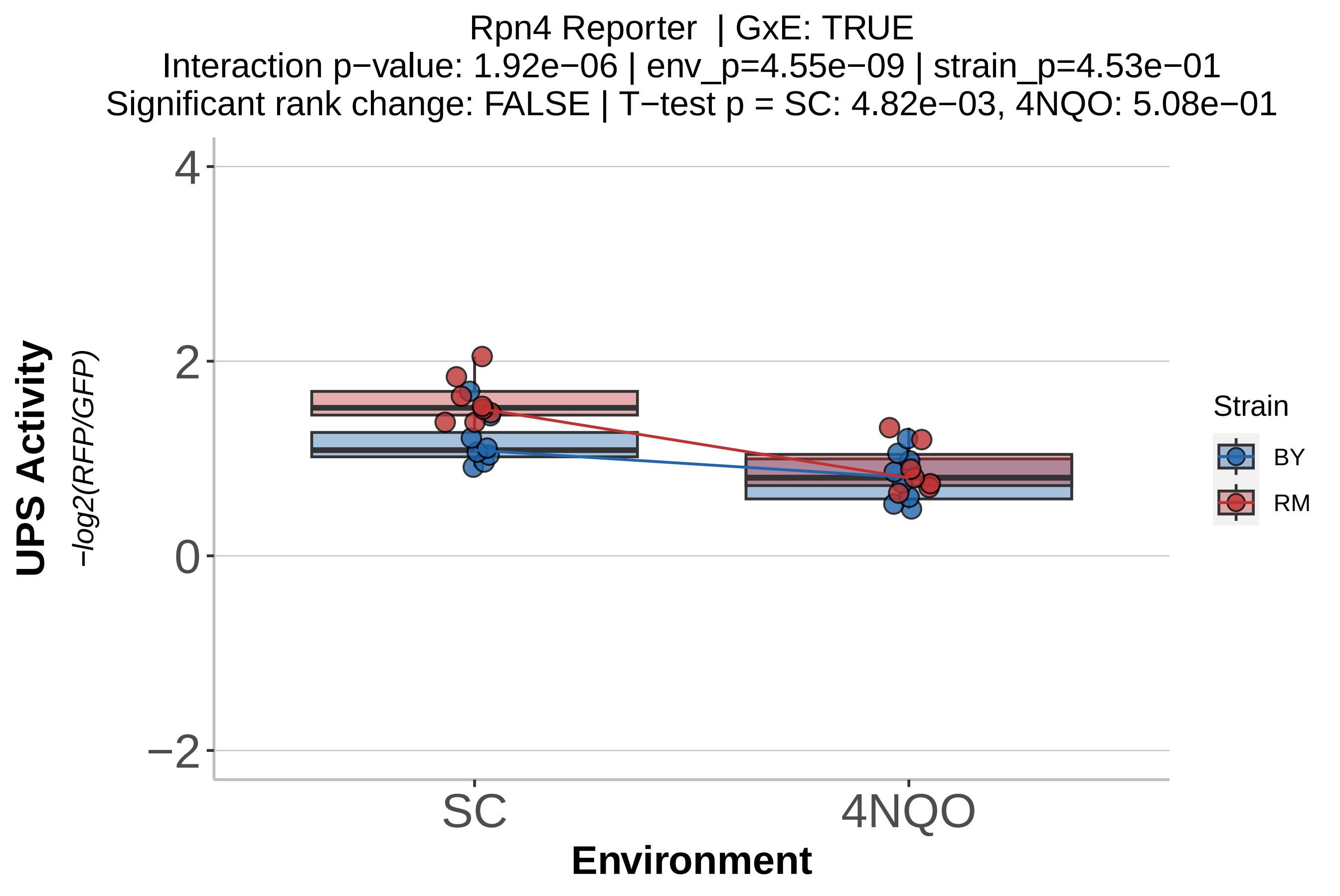
Rpn4 Reporter | GxE: TRUE
Interaction p−value: 1.92e−06 | env_p=4.55e−09 | strain_p=4.53e−01
Significant rank change: FALSE | T−test p = SC: 4.82e−03, 4NQO: 5.08e−01
UPS Activity
−log2(RFP/GFP)
4
2
0
−2
SC
4NQO
Environment
Strain
BY
RM

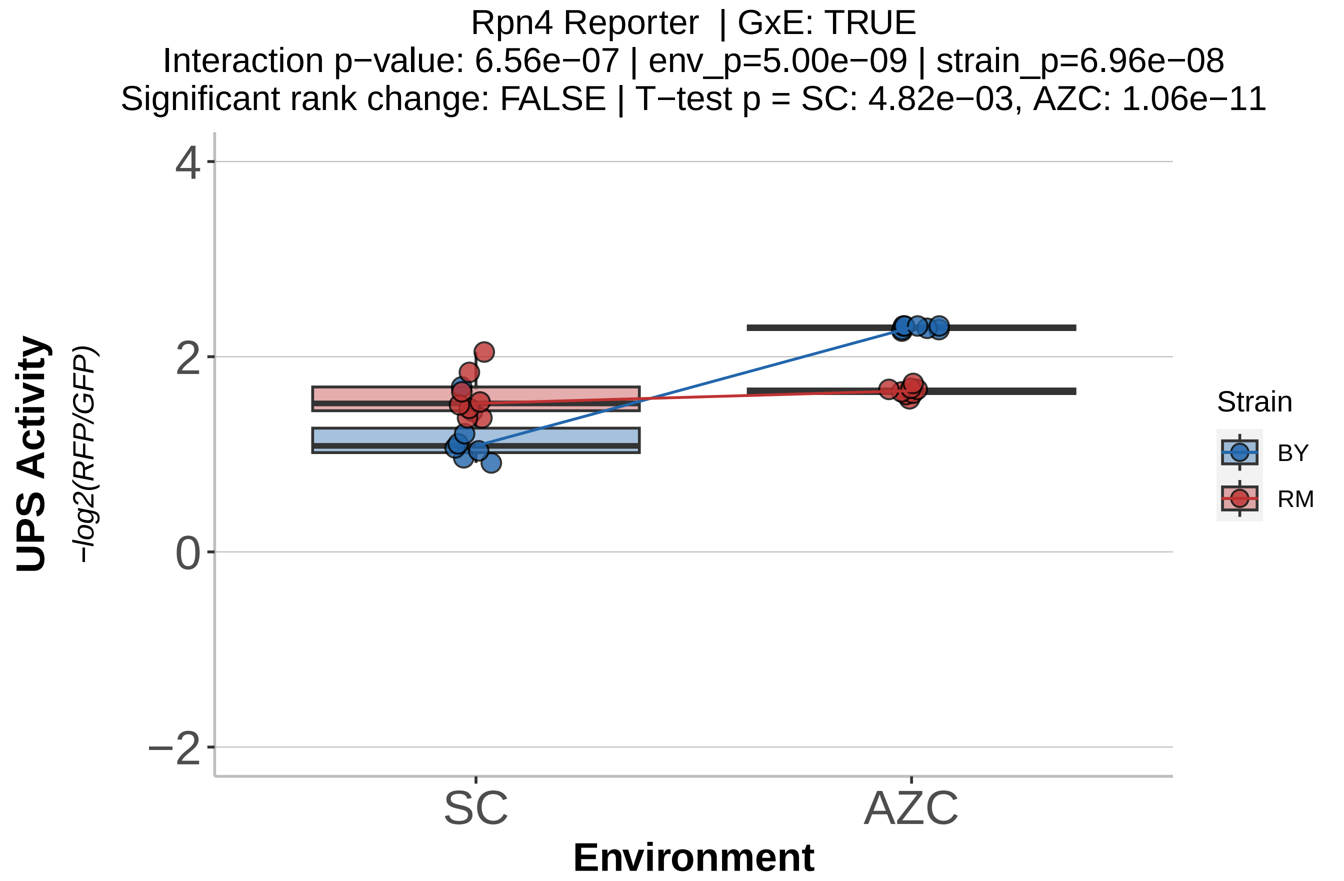
Rpn4 Reporter | GxE: TRUE
Interaction p−value: 6.56e−07 | env_p=5.00e−09 | strain_p=6.96e−08
Significant rank change: FALSE | T−test p = SC: 4.82e−03, AZC: 1.06e−11
UPS Activity
−log2(RFP/GFP)
4
2
0
−2
SC
AZC
Environment
Strain
BY
RM

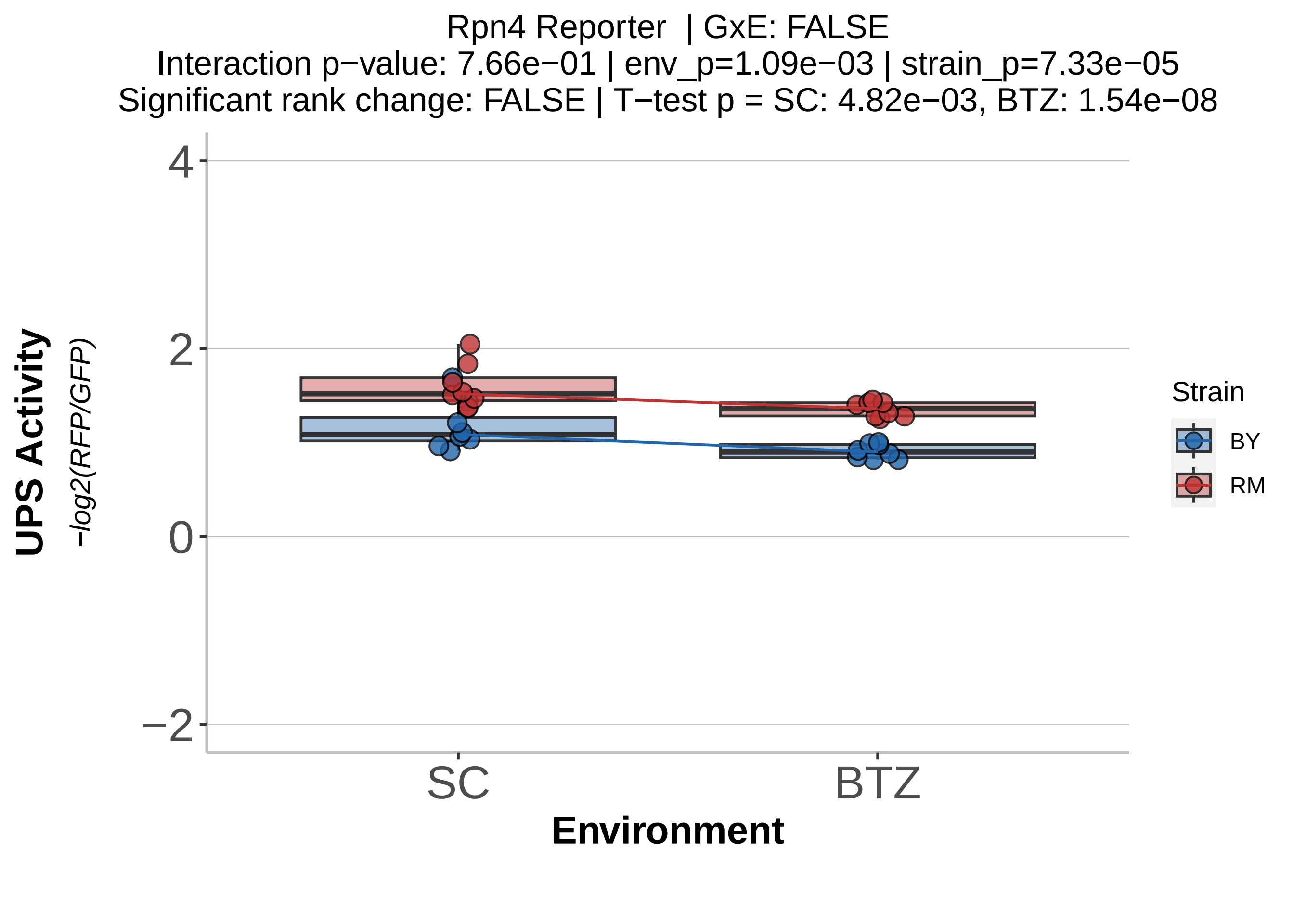

Rpn4 Reporter | GxE: FALSE
Interaction p−value: 7.66e−01 | env_p=1.09e−03 | strain_p=7.33e−05
Significant rank change: FALSE | T−test p = SC: 4.82e−03, BTZ: 1.54e−08
UPS Activity
−log2(RFP/GFP)
4
2
0
−2
SC
BTZ
Environment
Strain
BY
RM
