## Supplementary File 3 for "Genotype-by-environment interactions shape ubiquitin-proteasome system activity"

# 4x Ub in SC

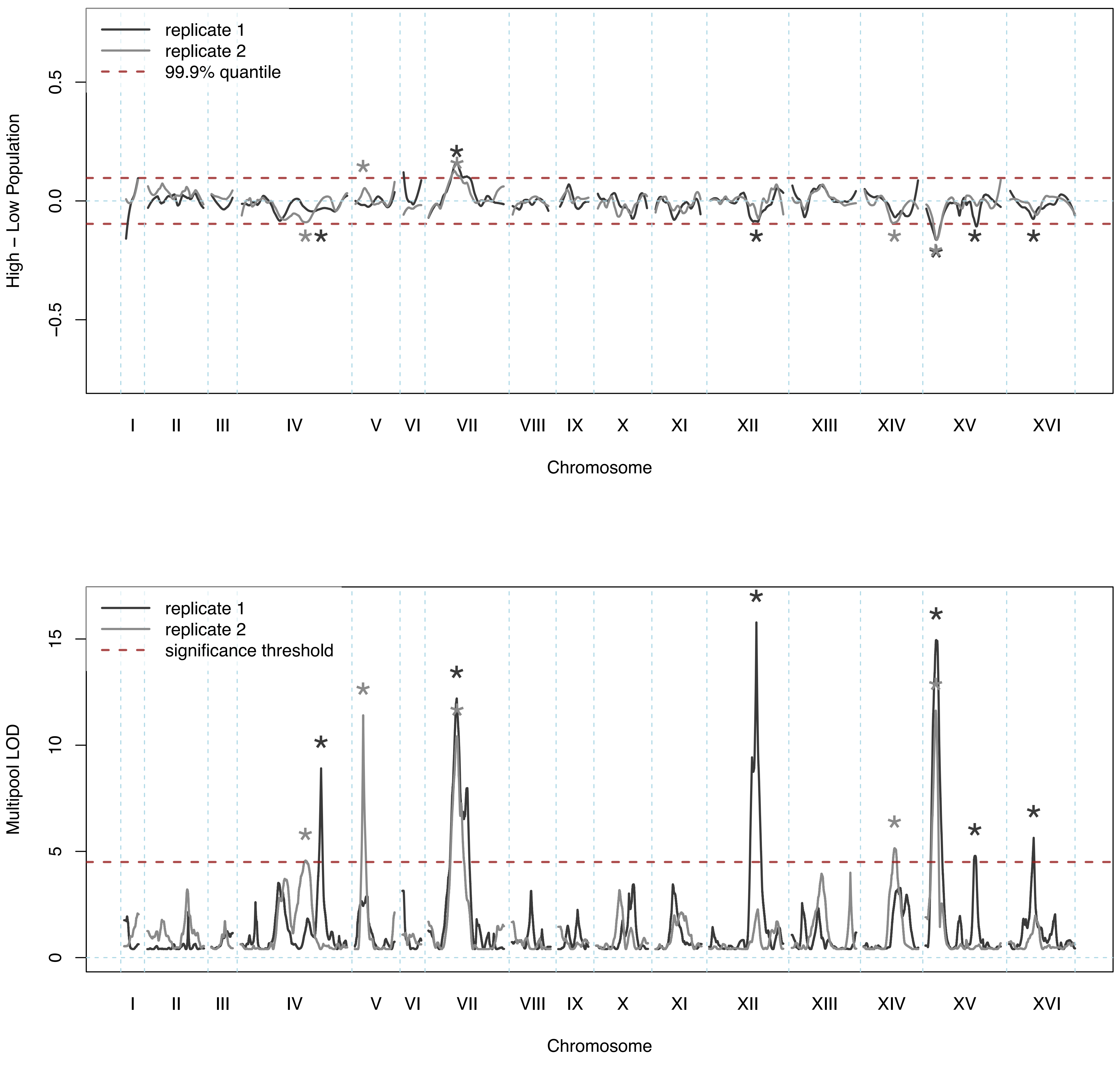
replicate 1
replicate 2
99.9% quantile
High – Low Population
0.5
0.0
−0.5
I II III IV V VI VII VIII IX X XI XII XIII XIV XV XVI
Chromosome
replicate 1
replicate 2
significance threshold
Multipool LOD
15
10
5
0
I II III IV V VI VII VIII IX X XI XII XIII XIV XV XVI
Chromosome

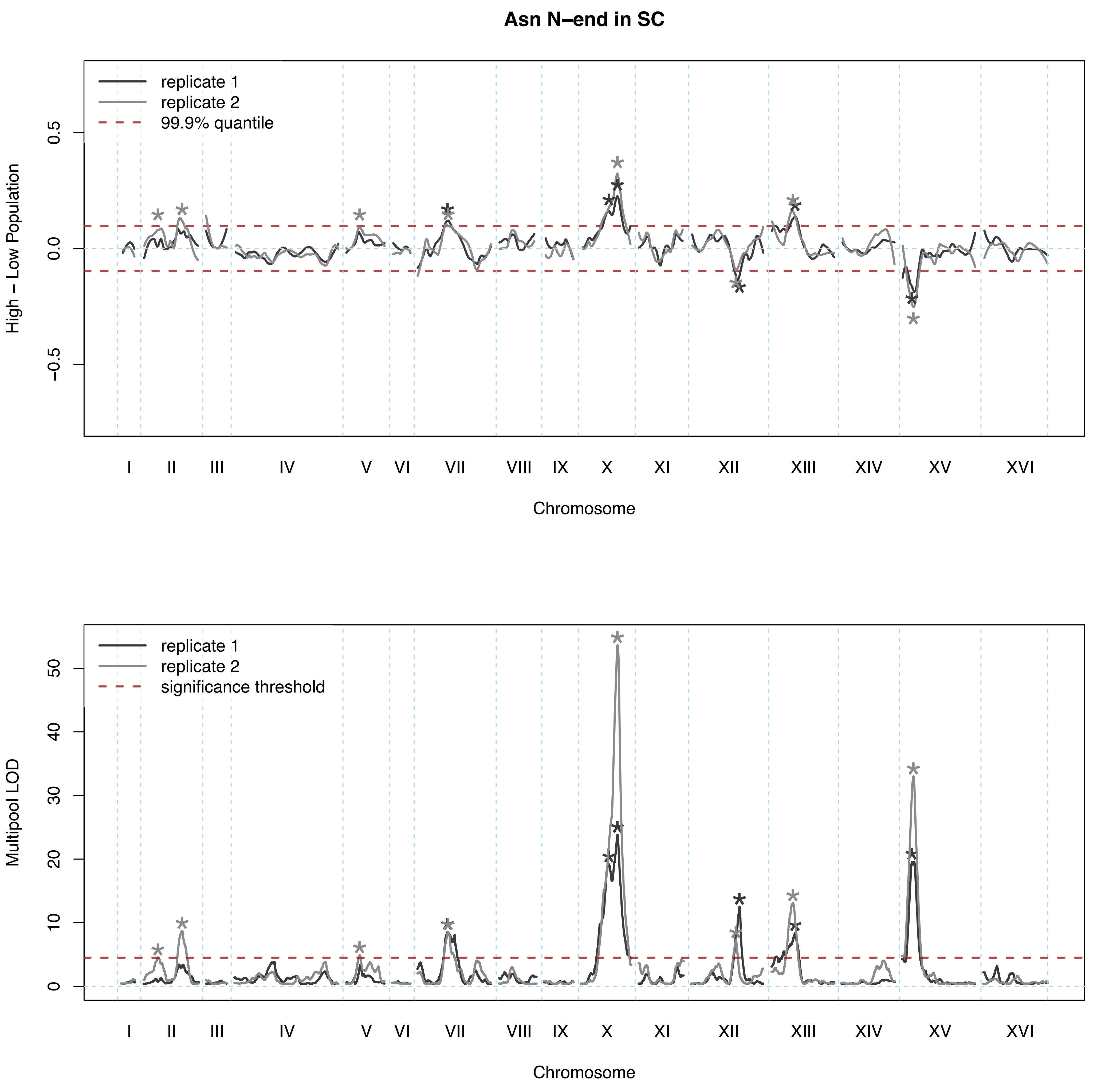

Asn N–end in SC
replicate 1
replicate 2
99.9% quantile
High – Low Population
0.5
0.0
−0.5
I
II
III
IV
V
VI
VII
VIII
IX
X
XI
XII
XIII
XIV
XV
XVI
Chromosome
replicate 1
replicate 2
significance threshold
Multipool LOD
50
40
30
20
10
0
I
II
III
IV
V
VI
VII
VIII
IX
X
XI
XII
XIII
XIV
XV
XVI
Chromosome

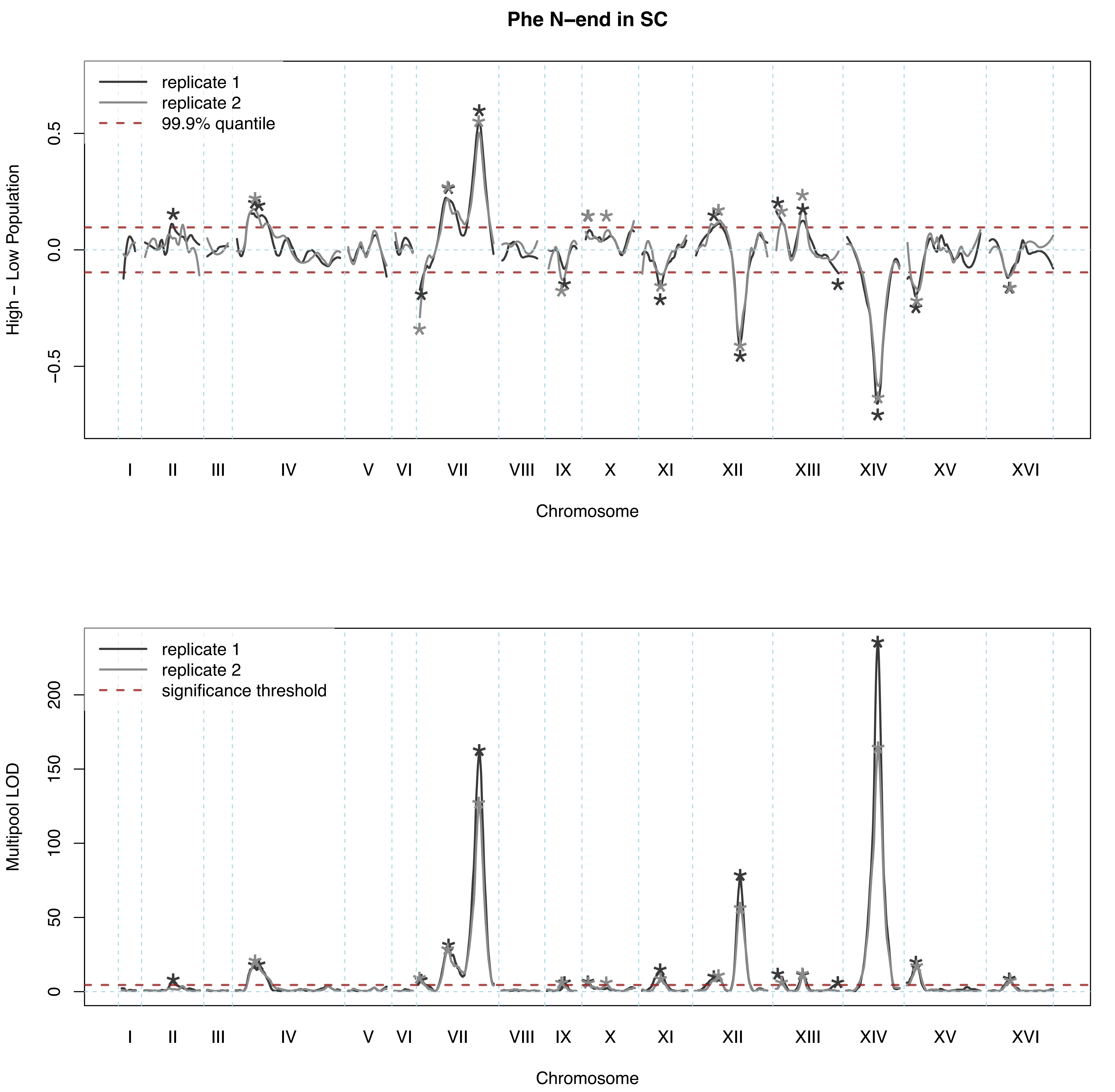
Phe N–end in SC
replicate 1
replicate 2
99.9% quantile
High – Low Population
0.5
0.0
−0.5
I
II
III
IV
V
VI
VII
VIII
IX
X
XI
XII
XIII
XIV
XV
XVI
Chromosome
replicate 1
replicate 2
significance threshold
Multipool LOD
0
50
100
150
200
Chromosome

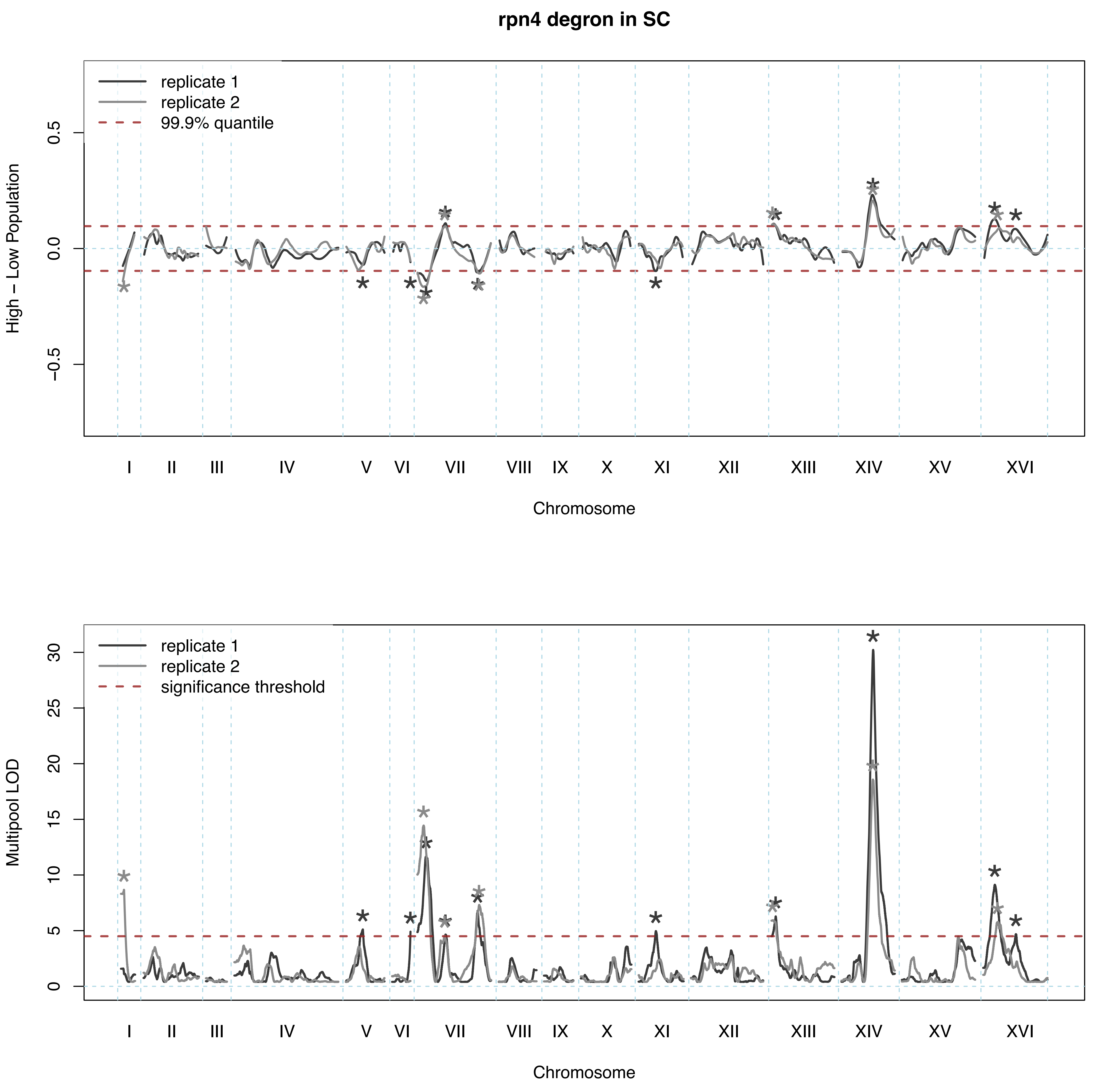

rpn4 degron in SC
replicate 1
replicate 2
99.9% quantile
High – Low Population
0.5
0.0
−0.5
I
II
III
IV
V
VI
VII
VIII
IX
X
XI
XII
XIII
XIV
XV
XVI
Chromosome
replicate 1
replicate 2
significance threshold
Multipool LOD
30
25
20
15
10
5
0
I
II
III
IV
V
VI
VII
VIII
IX
X
XI
XII
XIII
XIV
XV
XVI
Chromosome

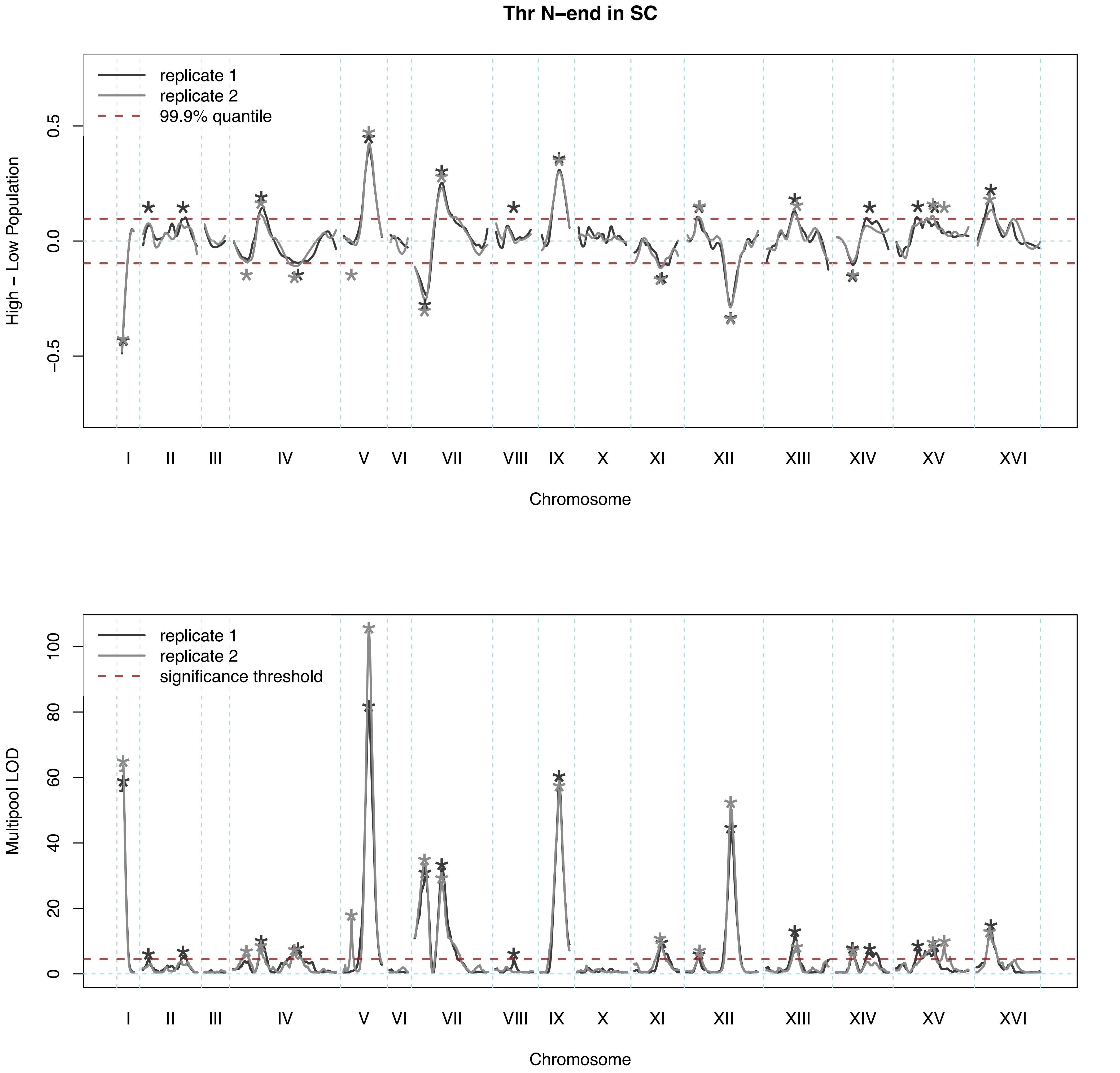

Thr N–end in SC
replicate 1
replicate 2
99.9% quantile
High – Low Population
0.5
0.0
−0.5
I
II
III
IV
V
VI
VII
VIII
IX
X
XI
XII
XIII
XIV
XV
XVI
Chromosome
replicate 1
replicate 2
significance threshold
Multipool LOD
100
80
60
40
20
0
Chromosome

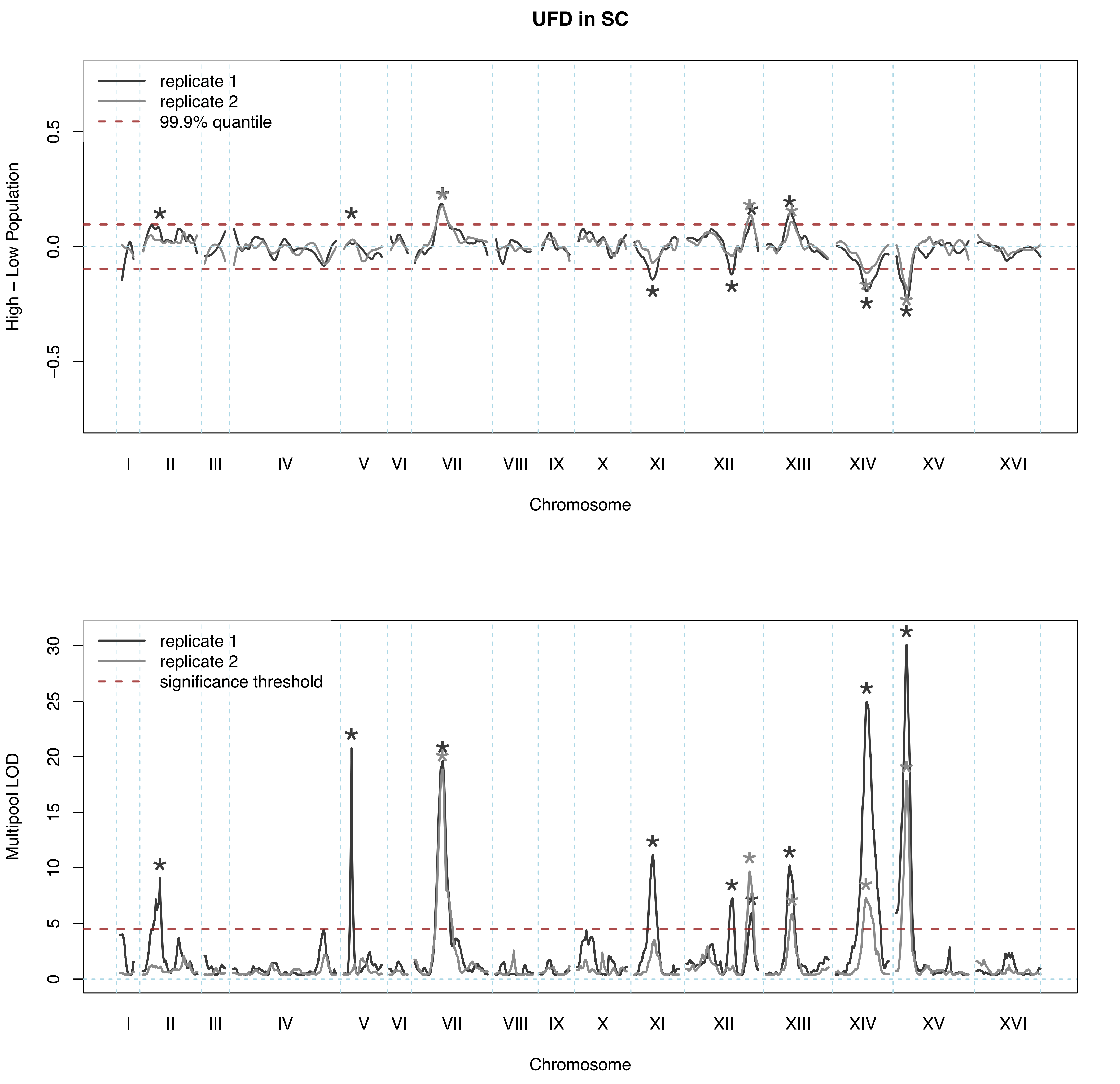

UFD in SC
replicate 1
replicate 2
99.9% quantile
High − Low Population
0.5
0.0
−0.5
I
II
III
IV
V
VI
VII
VIII
IX
X
XI
XII
XIII
XIV
XV
XVI
Chromosome
replicate 1
replicate 2
significance threshold
Multipool LOD
30
25
20
15
10
5
0
I
II
III
IV
V
VI
VII
VIII
IX
X
XI
XII
XIII
XIV
XV
XVI
Chromosome

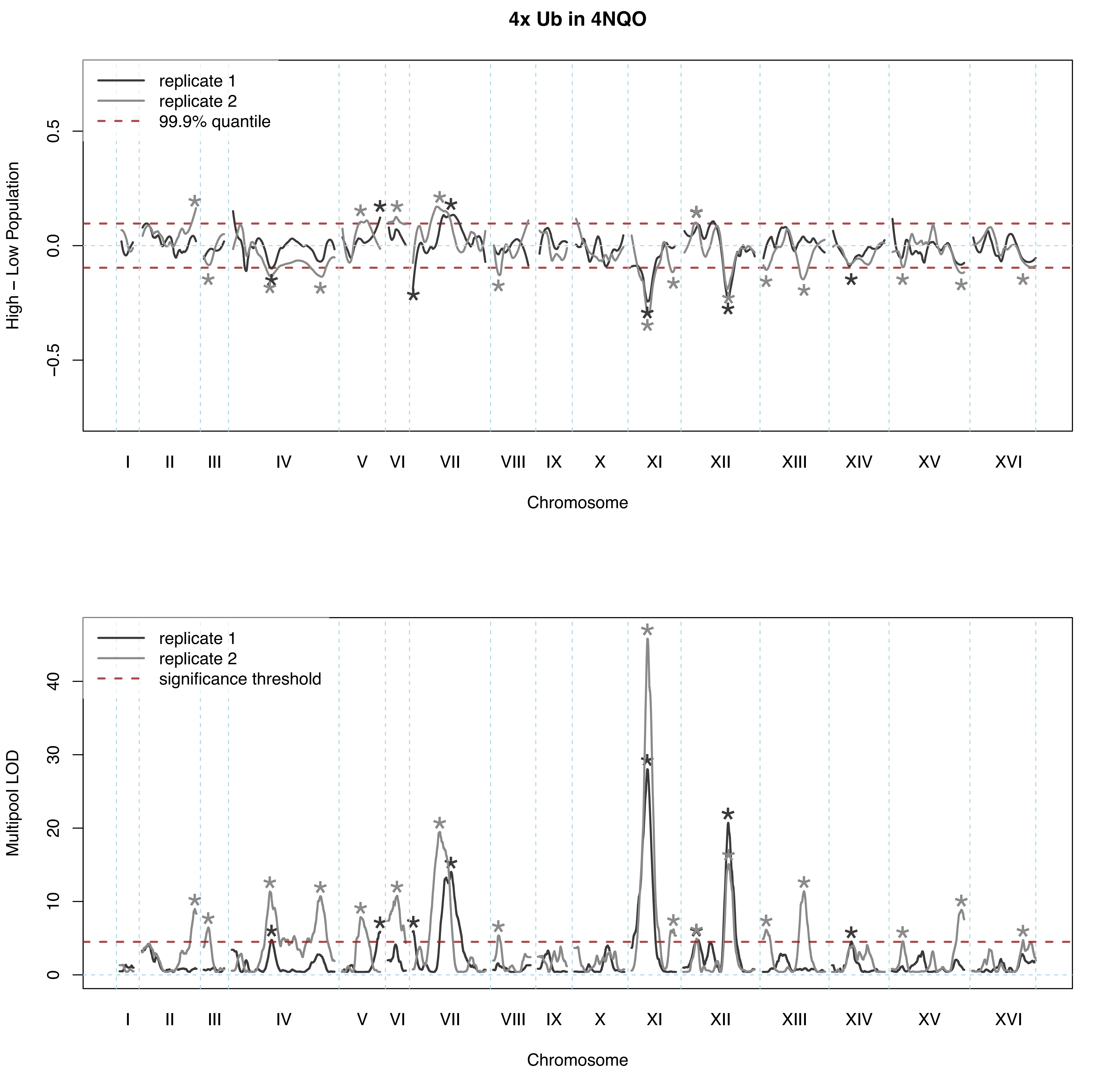

4x Ub in 4NQO
replicate 1
replicate 2
99.9% quantile
High – Low Population
0.5
0.0
−0.5
I
II
III
IV
V
VI
VII
VIII
IX
X
XI
XII
XIII
XIV
XV
XVI
Chromosome
replicate 1
replicate 2
significance threshold
Multipool LOD
0
10
20
30
40
I
II
III
IV
V
VI
VII
VIII
IX
X
XI
XII
XIII
XIV
XV
XVI
Chromosome

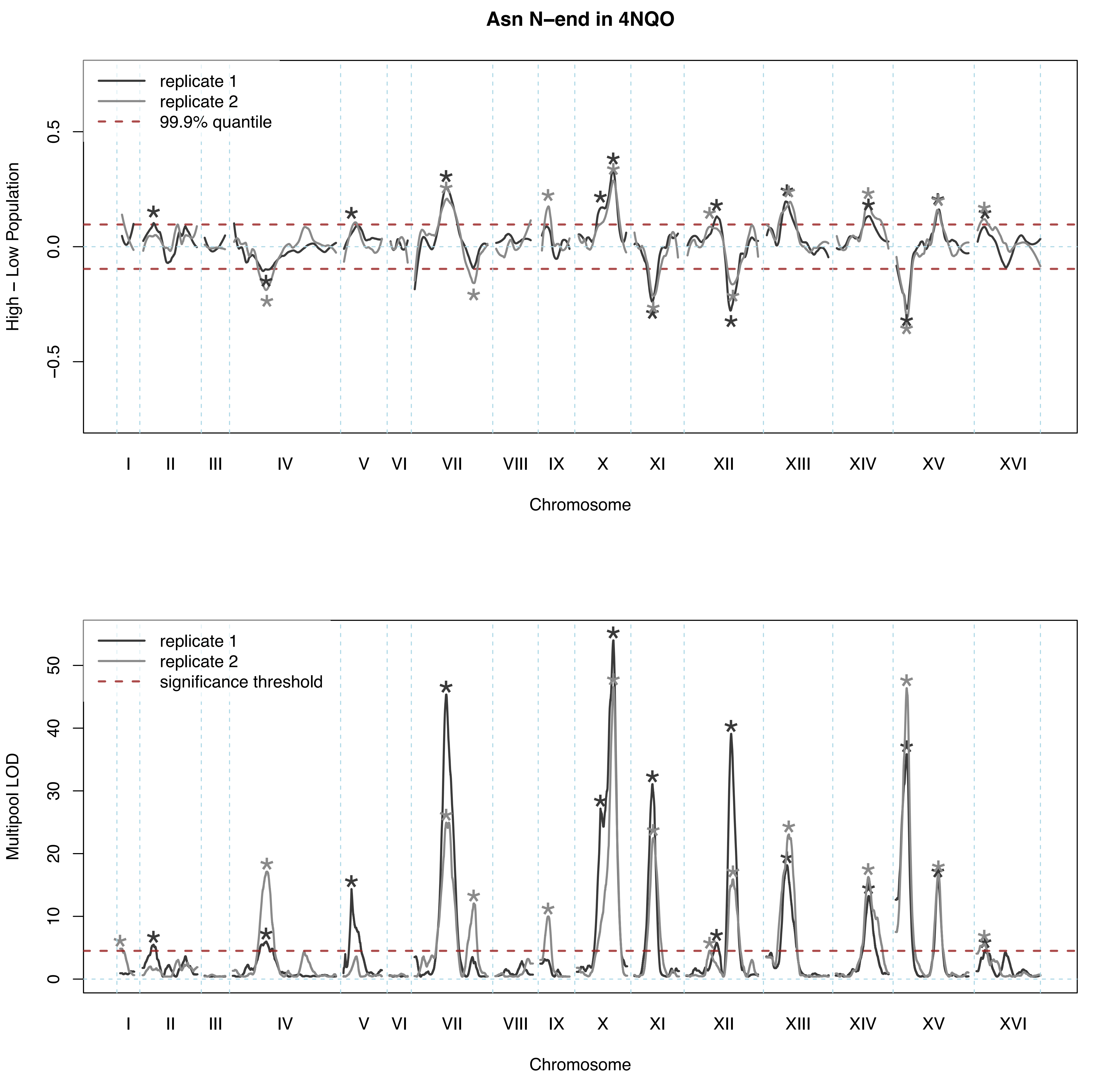

Asn N–end in 4NQO
replicate 1
replicate 2
99.9% quantile
High – Low Population
0.5
0.0
−0.5
I
II
III
IV
V
VI
VII
VIII
IX
X
XI
XII
XIII
XIV
XV
XVI
Chromosome
replicate 1
replicate 2
significance threshold
Multipool LOD
50
40
30
20
10
0
I
II
III
IV
V
VI
VII
VIII
IX
X
XI
XII
XIII
XIV
XV
XVI
Chromosome

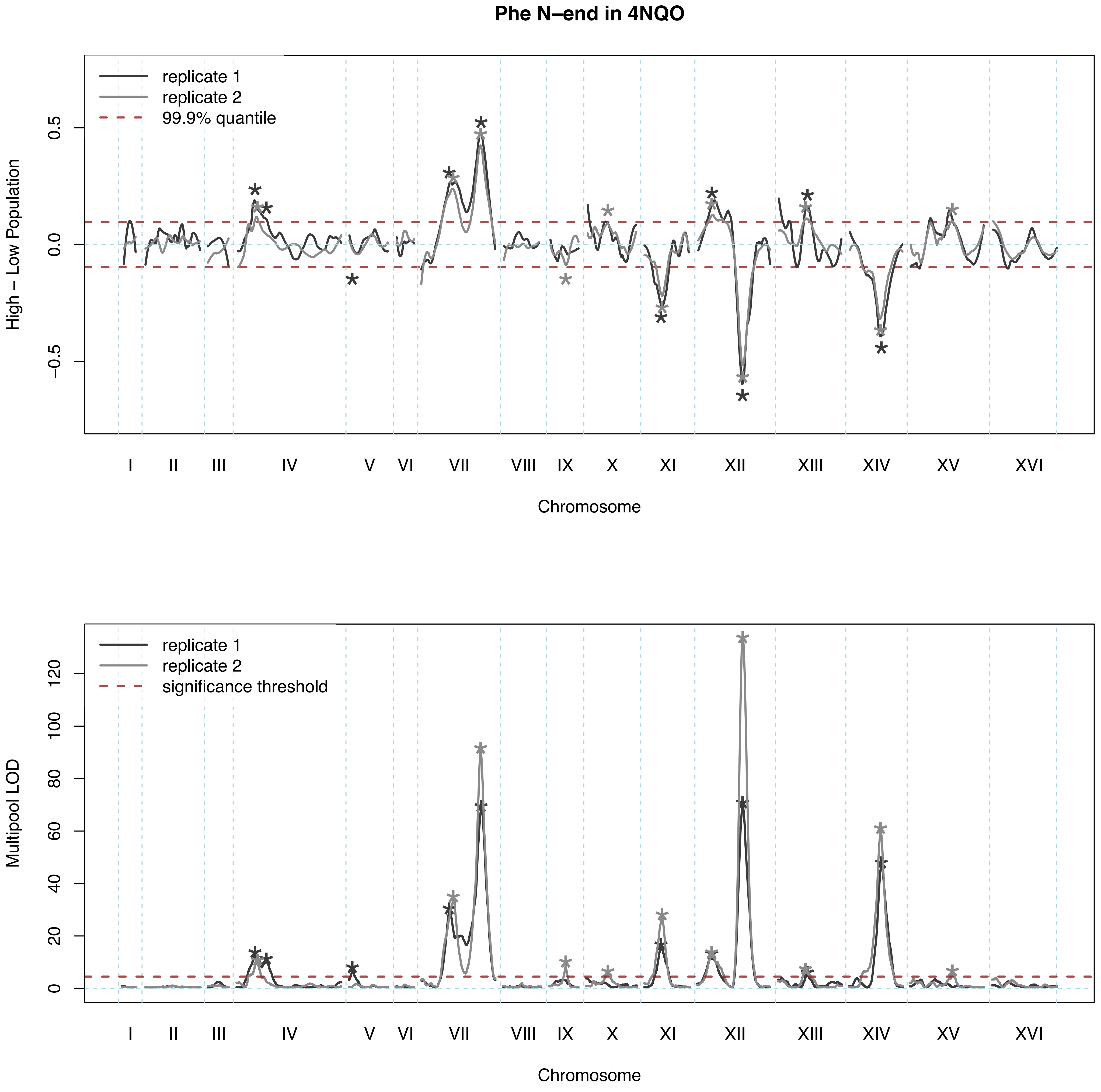

Phe N–end in 4NQO
replicate 1
replicate 2
99.9% quantile
High – Low Population
0.5
0.0
−0.5
I
II
III
IV
V
VI
VII
VIII
IX
X
XI
XII
XIII
XIV
XV
XVI
Chromosome
replicate 1
replicate 2
significance threshold
Multipool LOD
120
100
80
60
40
20
0
I
II
III
IV
V
VI
VII
VIII
IX
X
XI
XII
XIII
XIV
XV
XVI
Chromosome

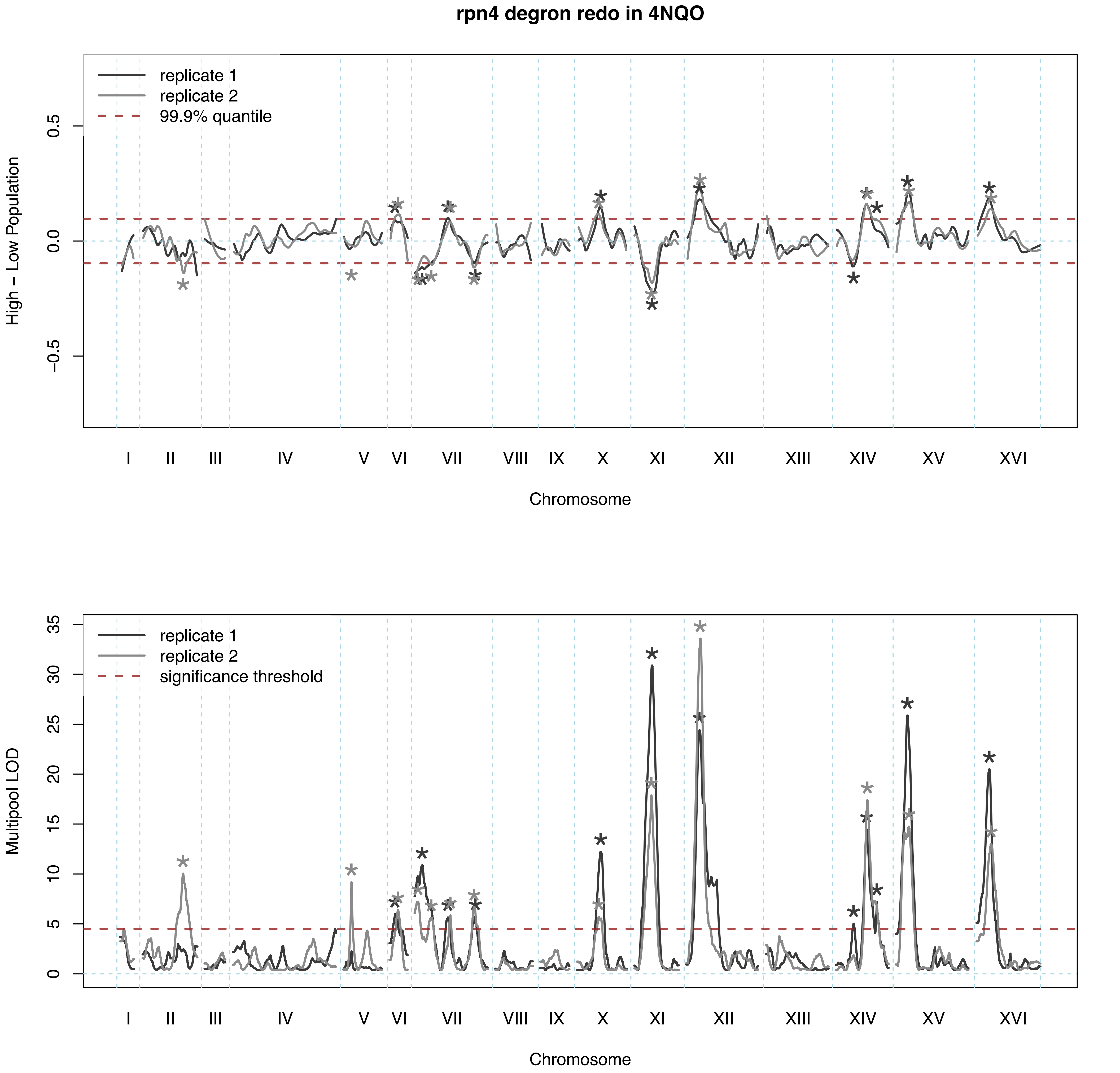

rpn4 degron redo in 4NQO
replicate 1
replicate 2
99.9% quantile
High – Low Population
0.5
0.0
−0.5
I
II
III
IV
V
VI
VII
VIII
IX
X
XI
XII
XIII
XIV
XV
XVI
Chromosome
replicate 1
replicate 2
significance threshold
Multipool LOD
35
30
25
20
15
10
5
0
I
II
III
IV
V
VI
VII
VIII
IX
X
XI
XII
XIII
XIV
XV
XVI
Chromosome

Thr N–end in 4NQO

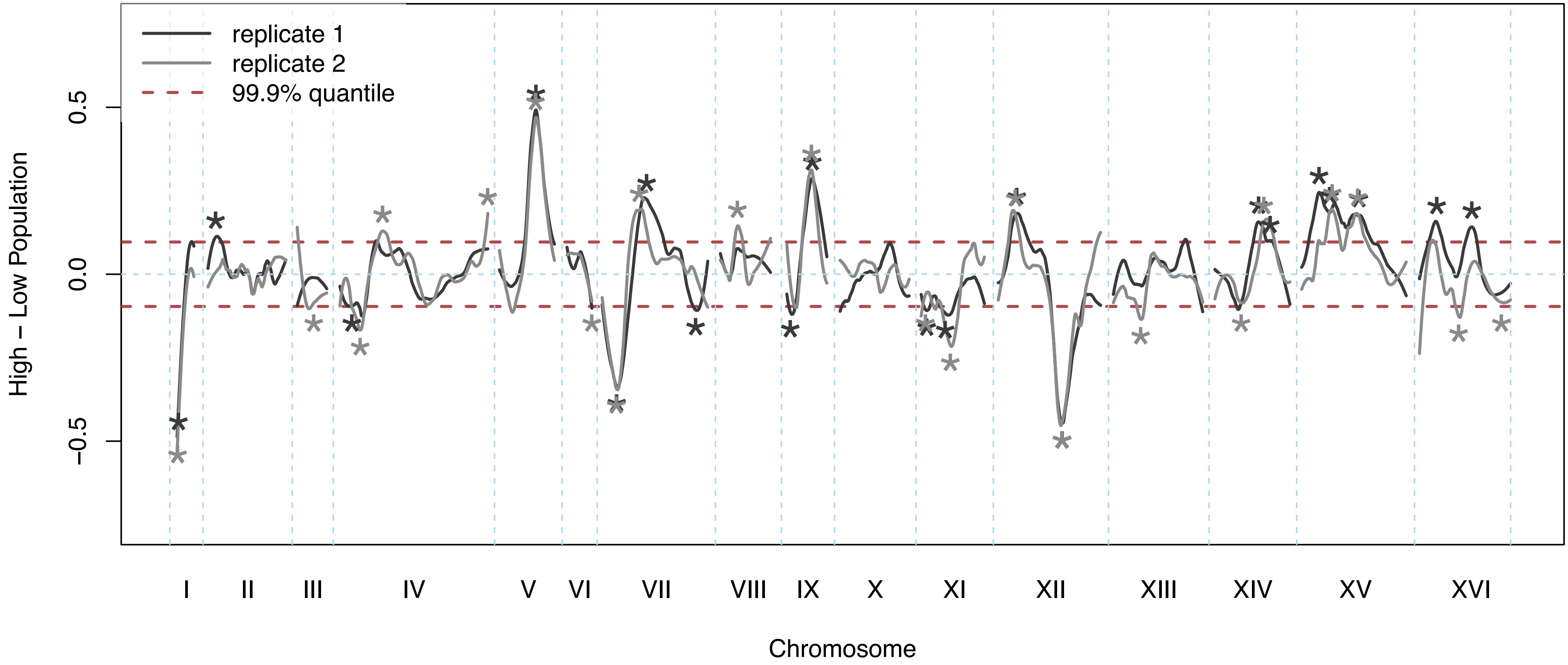


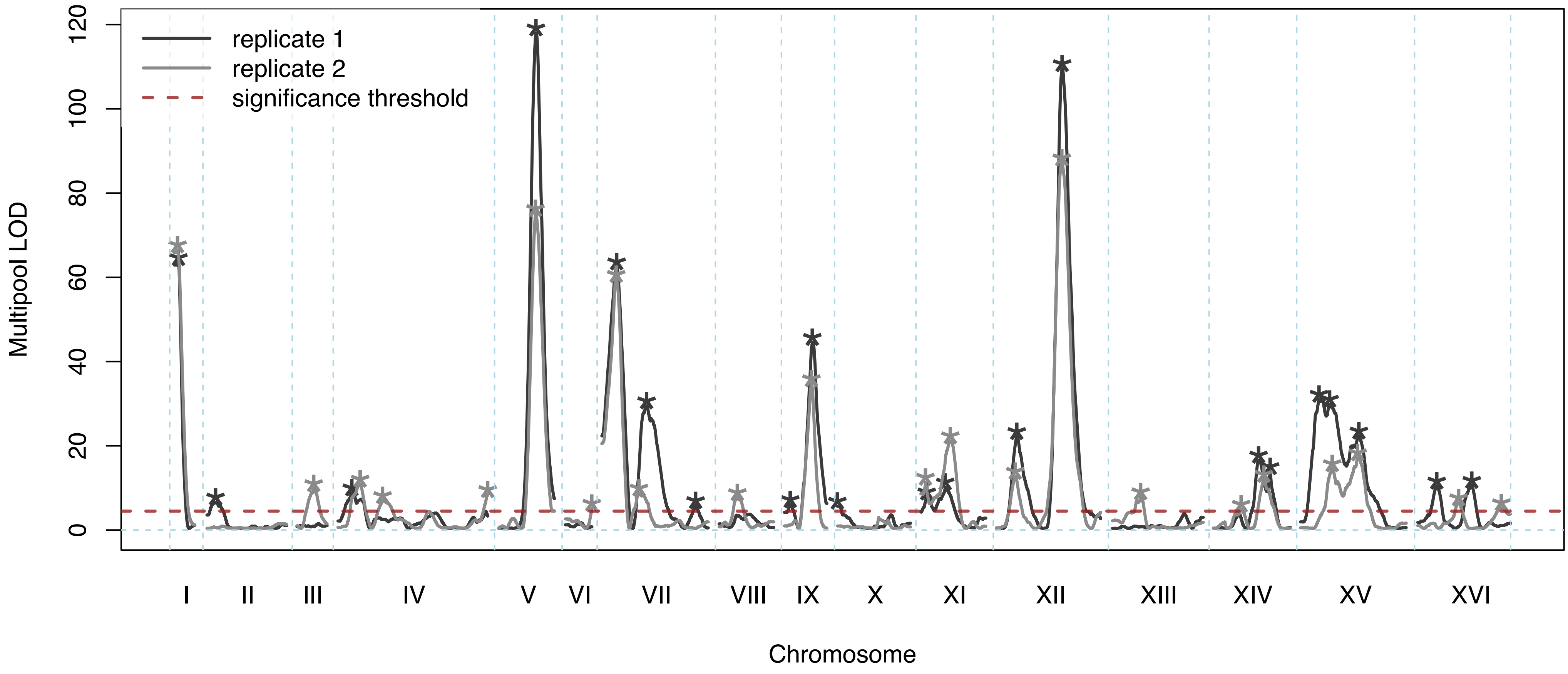

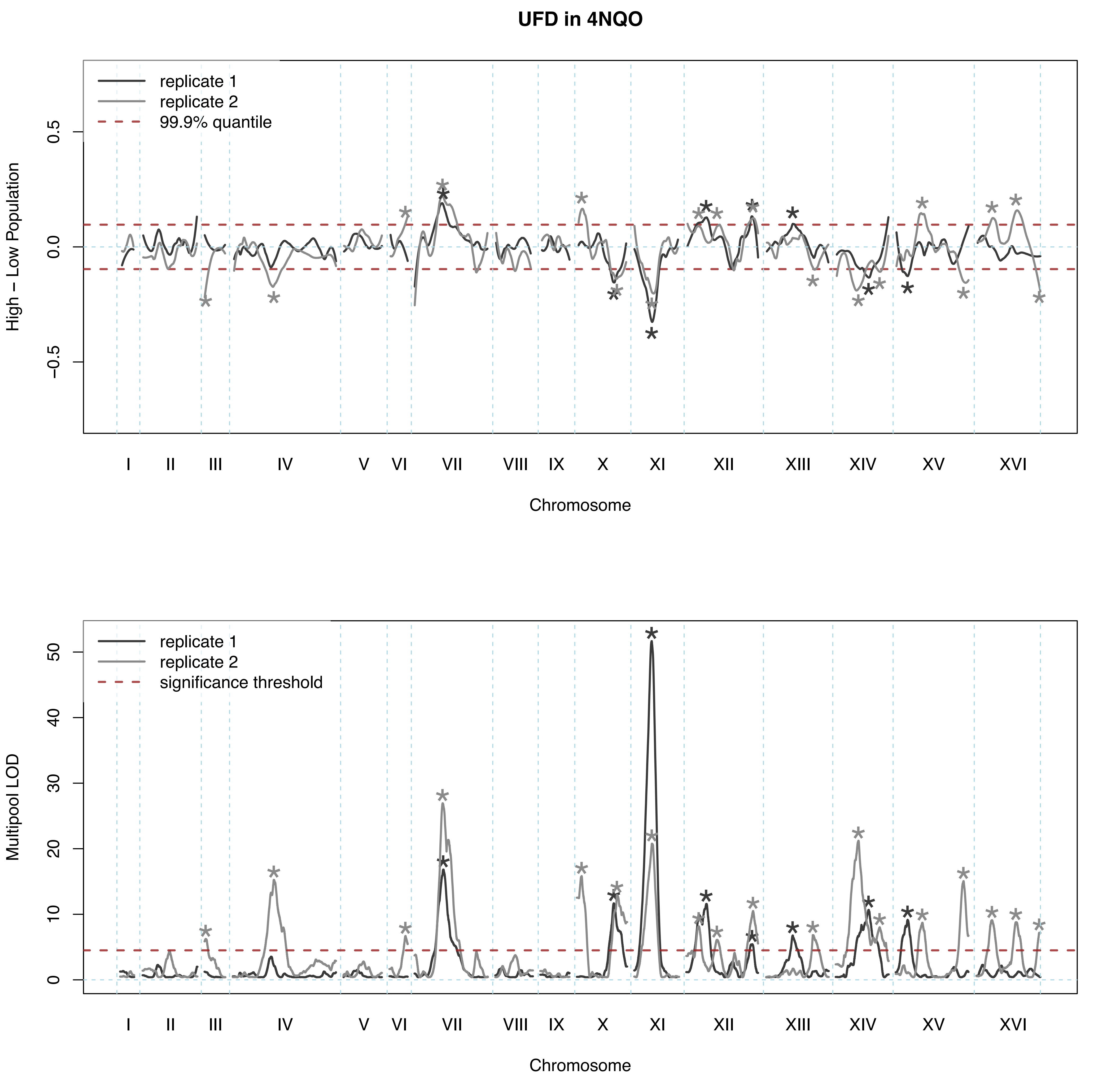

UFD in 4NQO
replicate 1
replicate 2
99.9% quantile
High – Low Population
0.5
0.0
−0.5
I
II
III
IV
V
VI
VII
VIII
IX
X
XI
XII
XIII
XIV
XV
XVI
Chromosome
replicate 1
replicate 2
significance threshold
Multipool LOD
50
40
30
20
10
0
I
II
III
IV
V
VI
VII
VIII
IX
X
XI
XII
XIII
XIV
XV
XVI
Chromosome

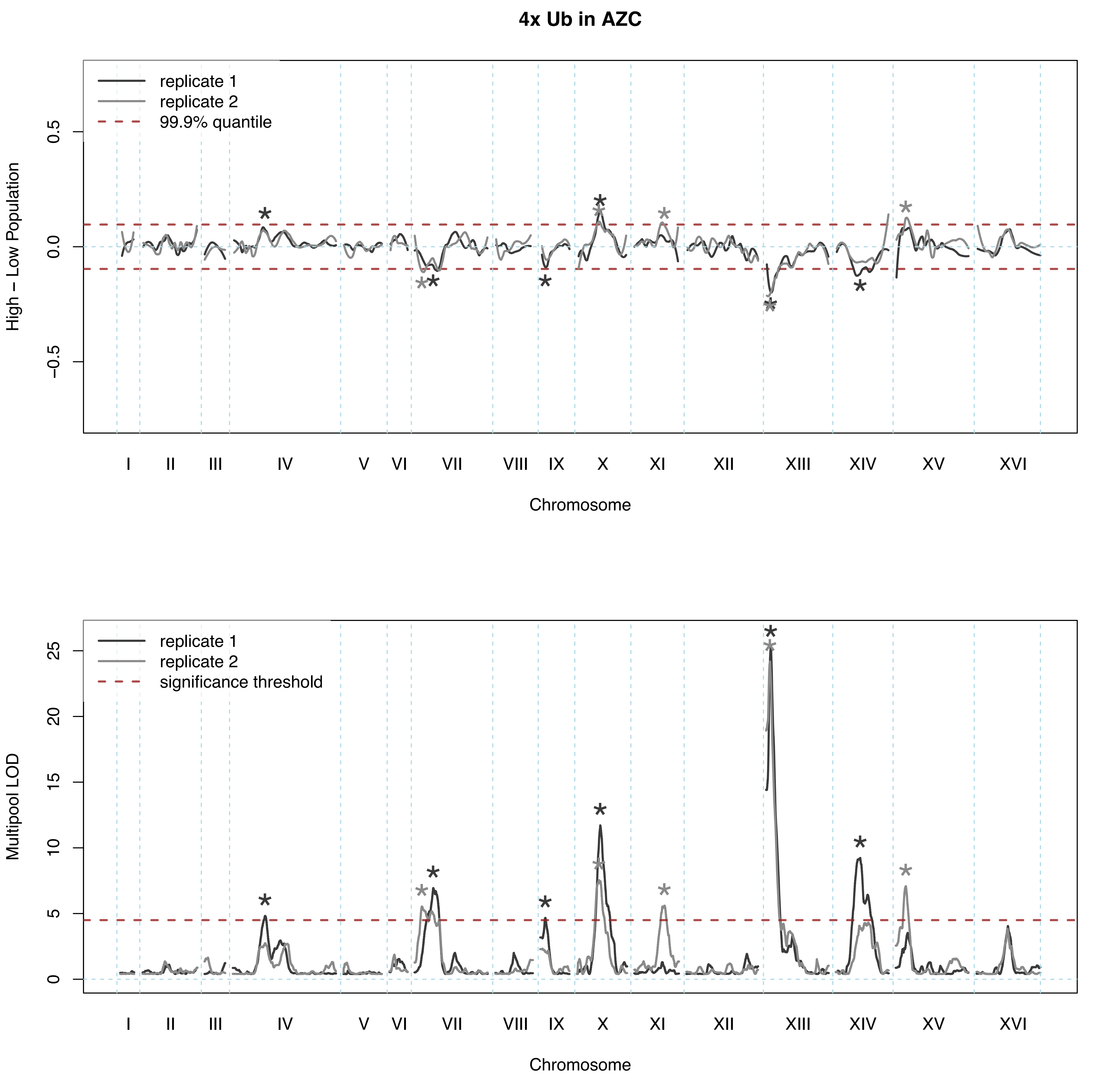
4x Ub in AZC
replicate 1
replicate 2
99.9% quantile
High – Low Population
0.5
0.0
−0.5
I
II
III
IV
V
VI
VII
VIII
IX
X
XI
XII
XIII
XIV
XV
XVI
Chromosome
replicate 1
replicate 2
significance threshold
Multipool LOD
25
20
15
10
5
0
Chromosome

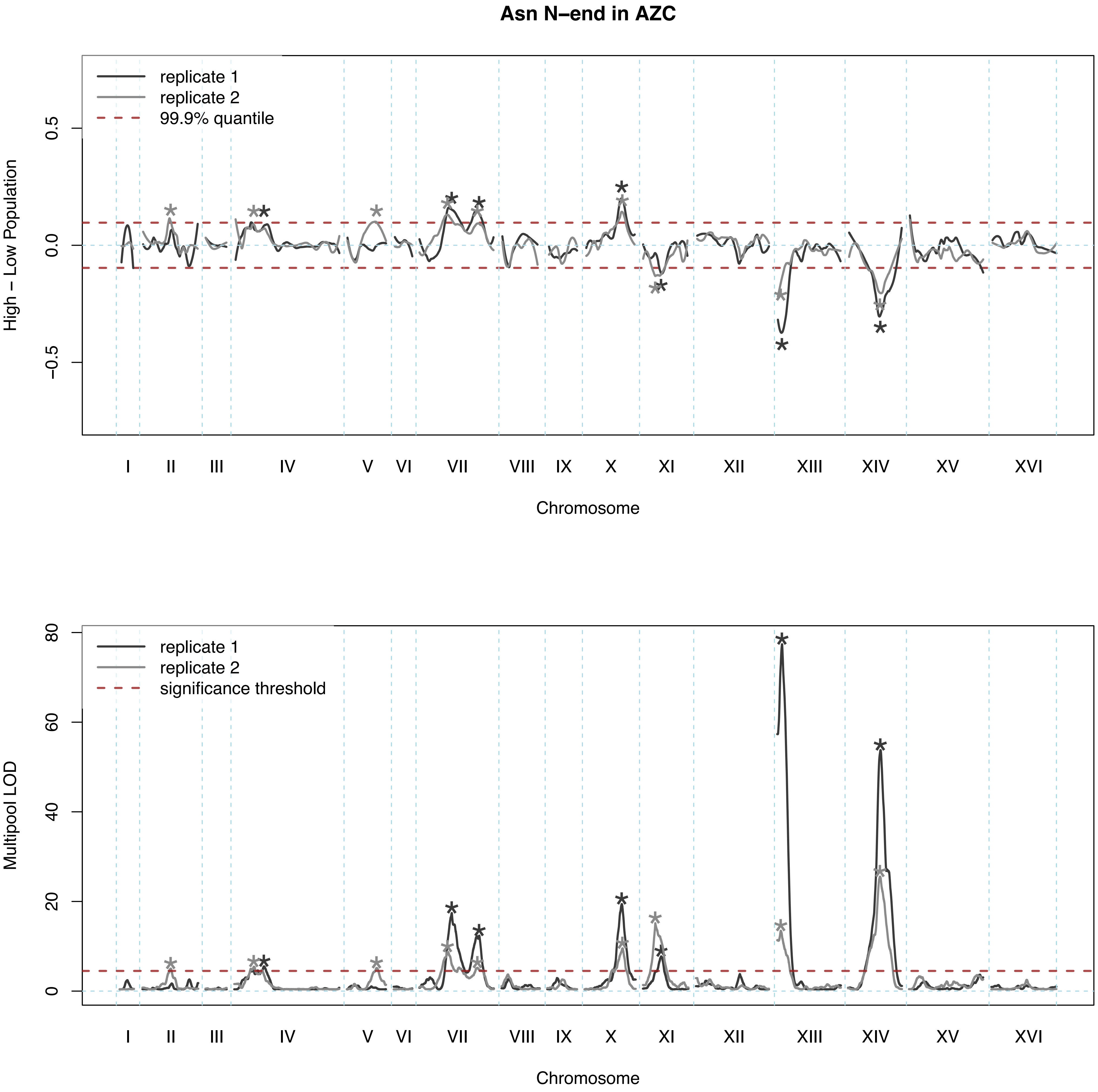

Asn N–end in AZC
replicate 1
replicate 2
99.9% quantile
High – Low Population
0.5
0.0
−0.5
I
II
III
IV
V
VI
VII
VIII
IX
X
XI
XII
XIII
XIV
XV
XVI
Chromosome
replicate 1
replicate 2
significance threshold
Multipool LOD
80
60
40
20
0
I
II
III
IV
V
VI
VII
VIII
IX
X
XI
XII
XIII
XIV
XV
XVI
Chromosome

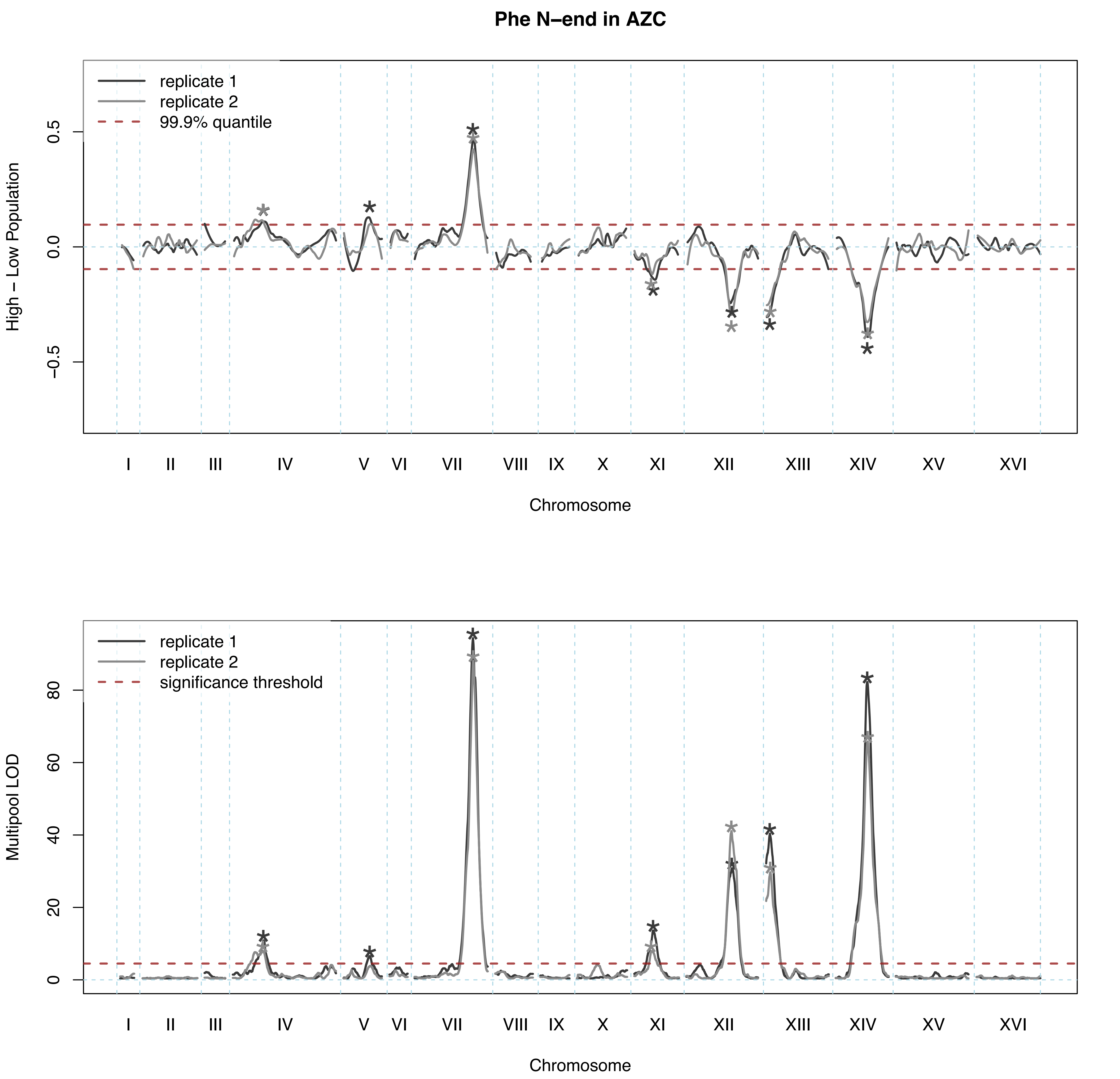

Phe N–end in AZC
replicate 1
replicate 2
99.9% quantile
High – Low Population
0.5
0.0
−0.5
I
II
III
IV
V
VI
VII
VIII
IX
X
XI
XII
XIII
XIV
XV
XVI
Chromosome
replicate 1
replicate 2
significance threshold
Multipool LOD
0
20
40
60
80
Chromosome

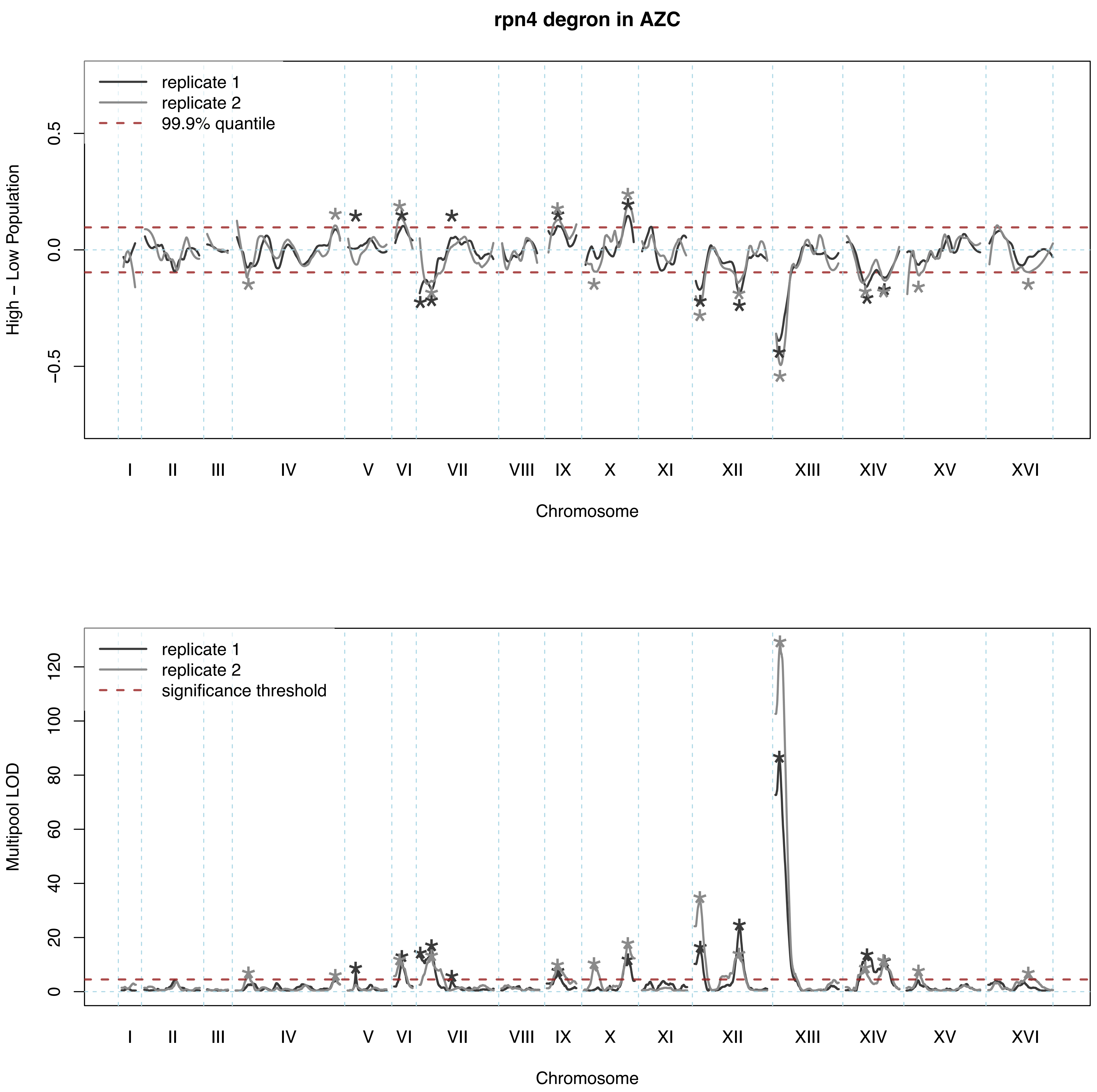
rpn4 degron in AZC
replicate 1
replicate 2
99.9% quantile
High – Low Population
0.5
0.0
−0.5
I II III IV V VI VII VIII IX X XI XII XIII XIV XV XVI
Chromosome
replicate 1
replicate 2
significance threshold
Multipool LOD
0 20 40 60 80 100 120
I II III IV V VI VII VIII IX X XI XII XIII XIV XV XVI
Chromosome

### Thr N–end in AZC

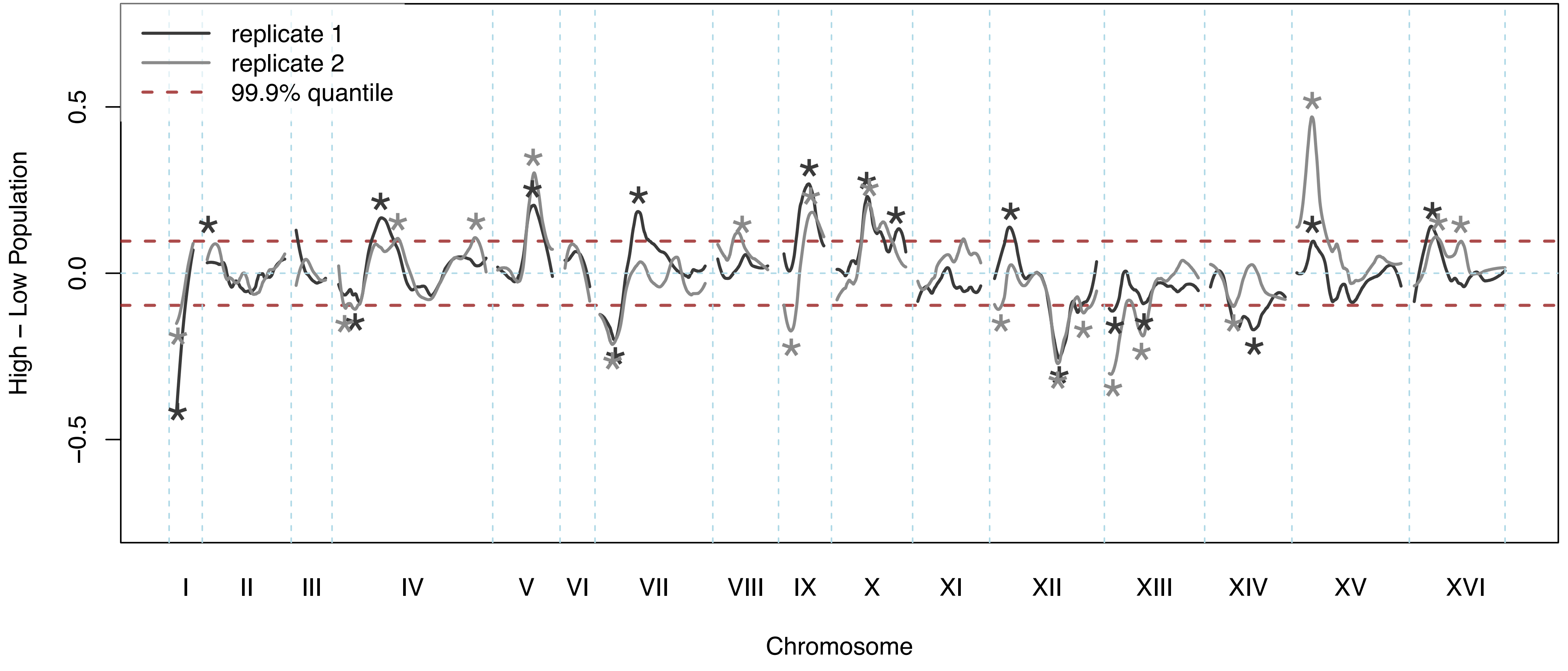


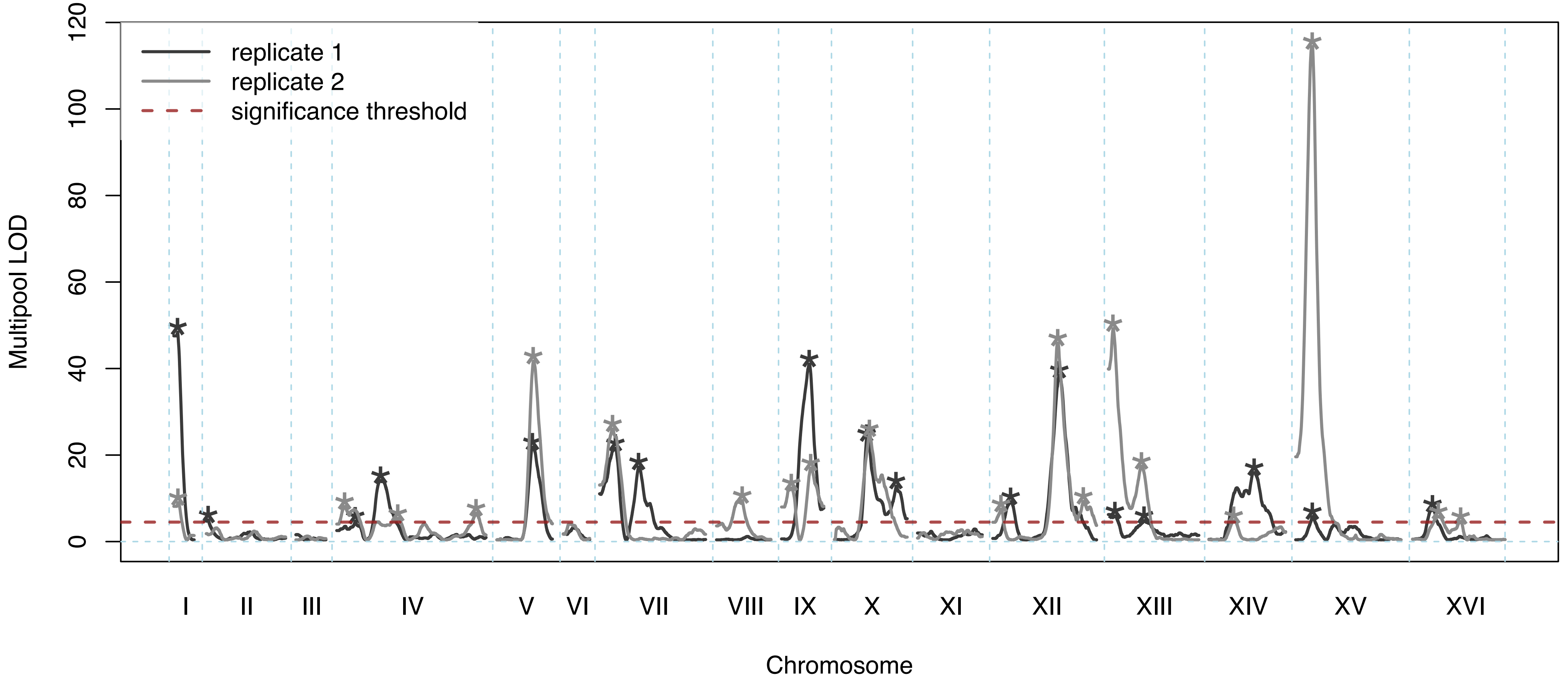

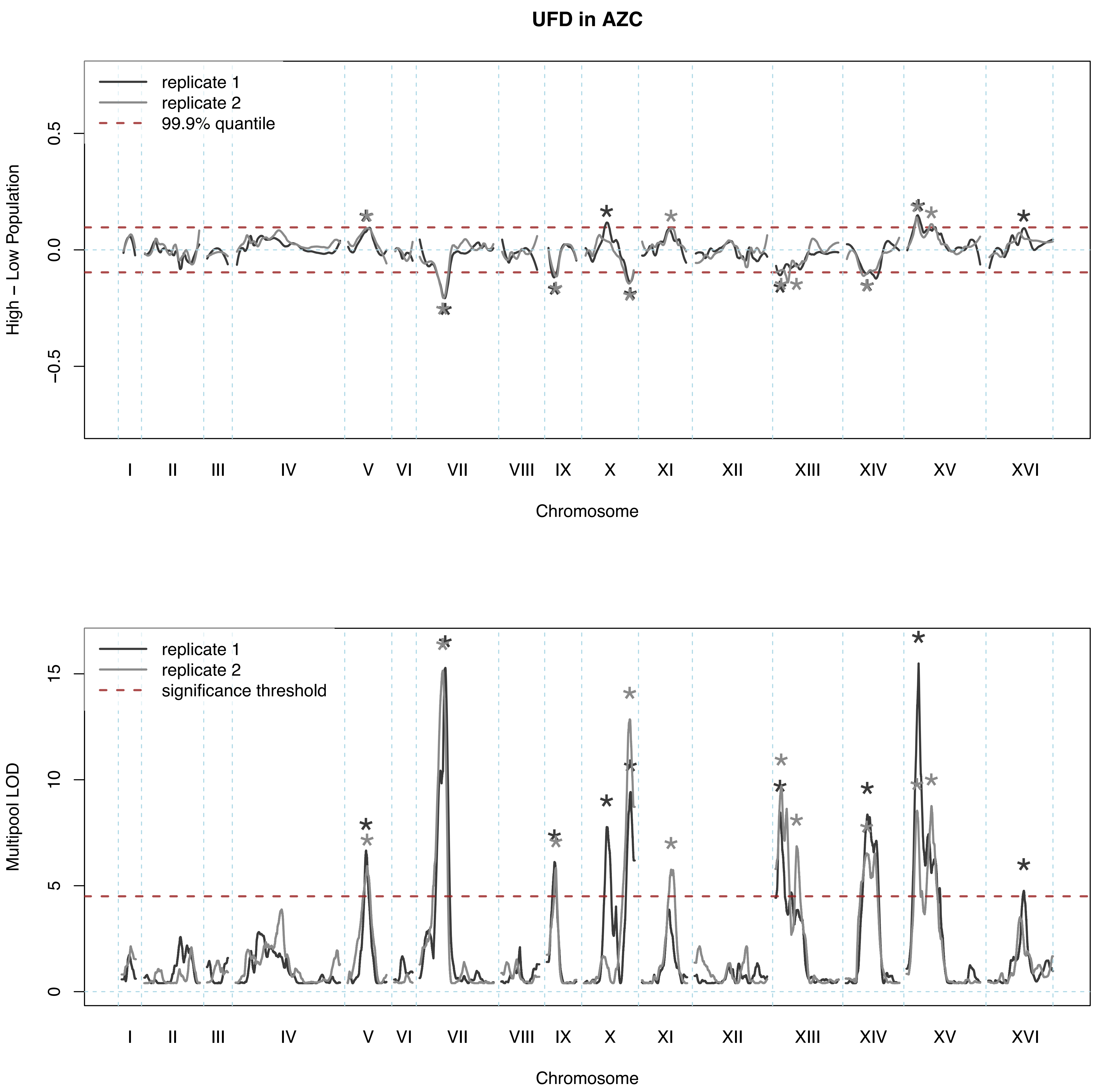

UFD in AZC
replicate 1
replicate 2
99.9% quantile
High – Low Population
0.5
0.0
−0.5
I II III IV V VI VII VIII IX X XI XII XIII XIV XV XVI
Chromosome
replicate 1
replicate 2
significance threshold
Multipool LOD
15
10
5
0
I II III IV V VI VII VIII IX X XI XII XIII XIV XV XVI
Chromosome

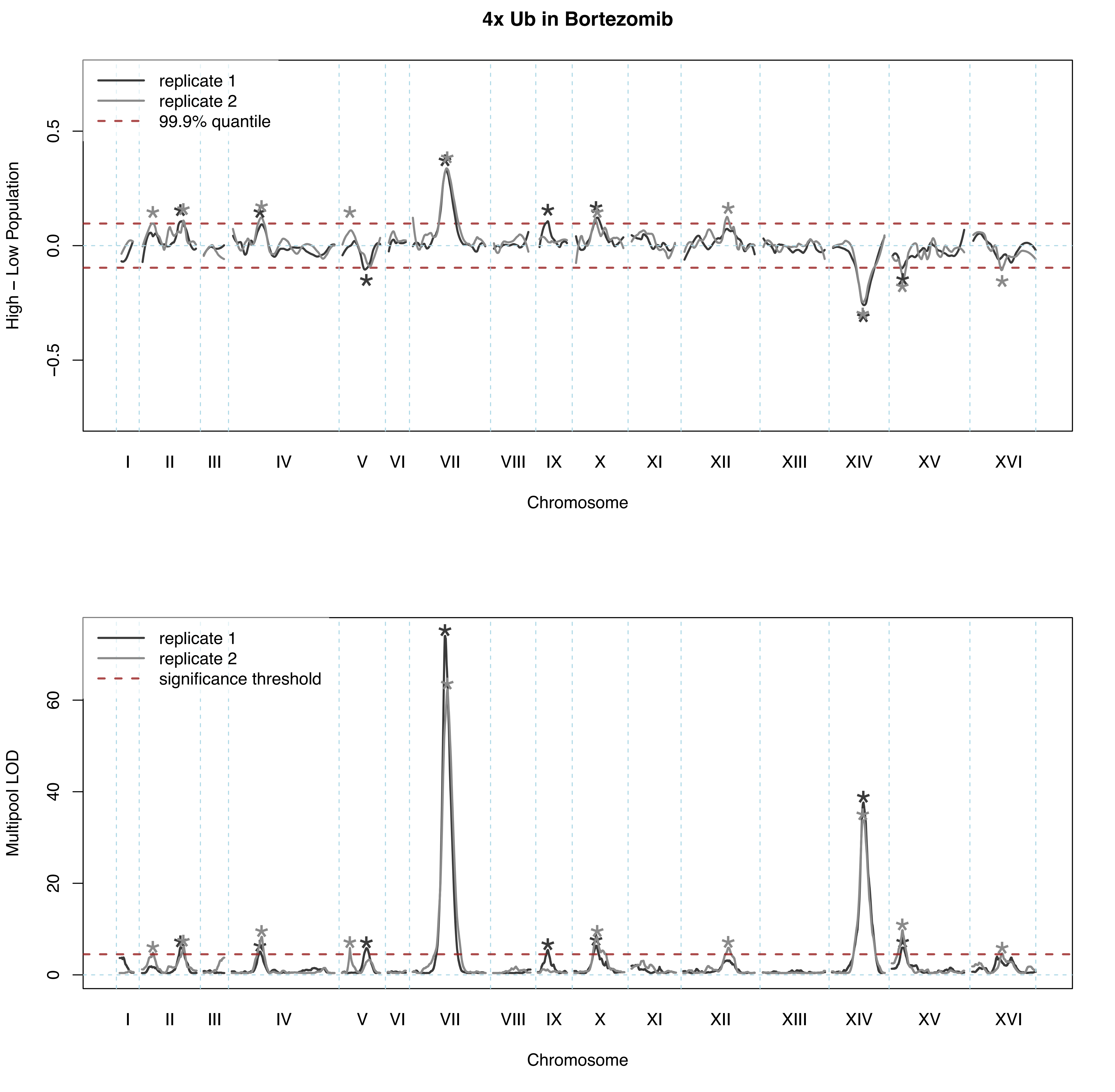

4x Ub in Bortezomib
replicate 1
replicate 2
99.9% quantile
High – Low Population
0.5
0.0
−0.5
I
II
III
IV
V
VI
VII
VIII
IX
X
XI
XII
XIII
XIV
XV
XVI
Chromosome
replicate 1
replicate 2
significance threshold
Multipool LOD
60
40
20
0
I
II
III
IV
V
VI
VII
VIII
IX
X
XI
XII
XIII
XIV
XV
XVI
Chromosome

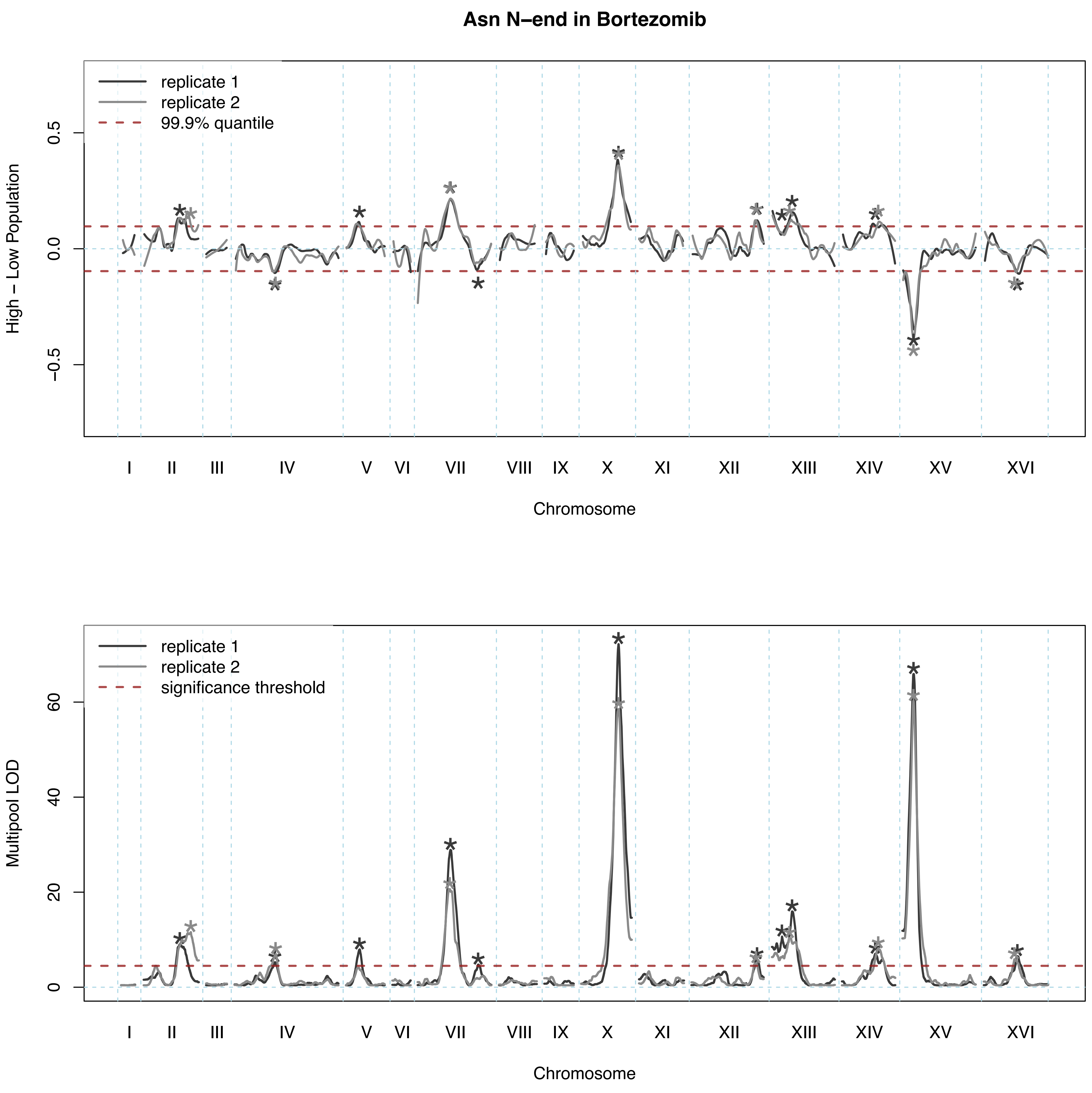

Asn N–end in Bortezomib
replicate 1
replicate 2
99.9% quantile
High – Low Population
0.5
0.0
−0.5
I
II
III
IV
V
VI
VII
VIII
IX
X
XI
XII
XIII
XIV
XV
XVI
Chromosome
replicate 1
replicate 2
significance threshold
Multipool LOD
60
40
20
0
I
II
III
IV
V
VI
VII
VIII
IX
X
XI
XII
XIII
XIV
XV
XVI
Chromosome

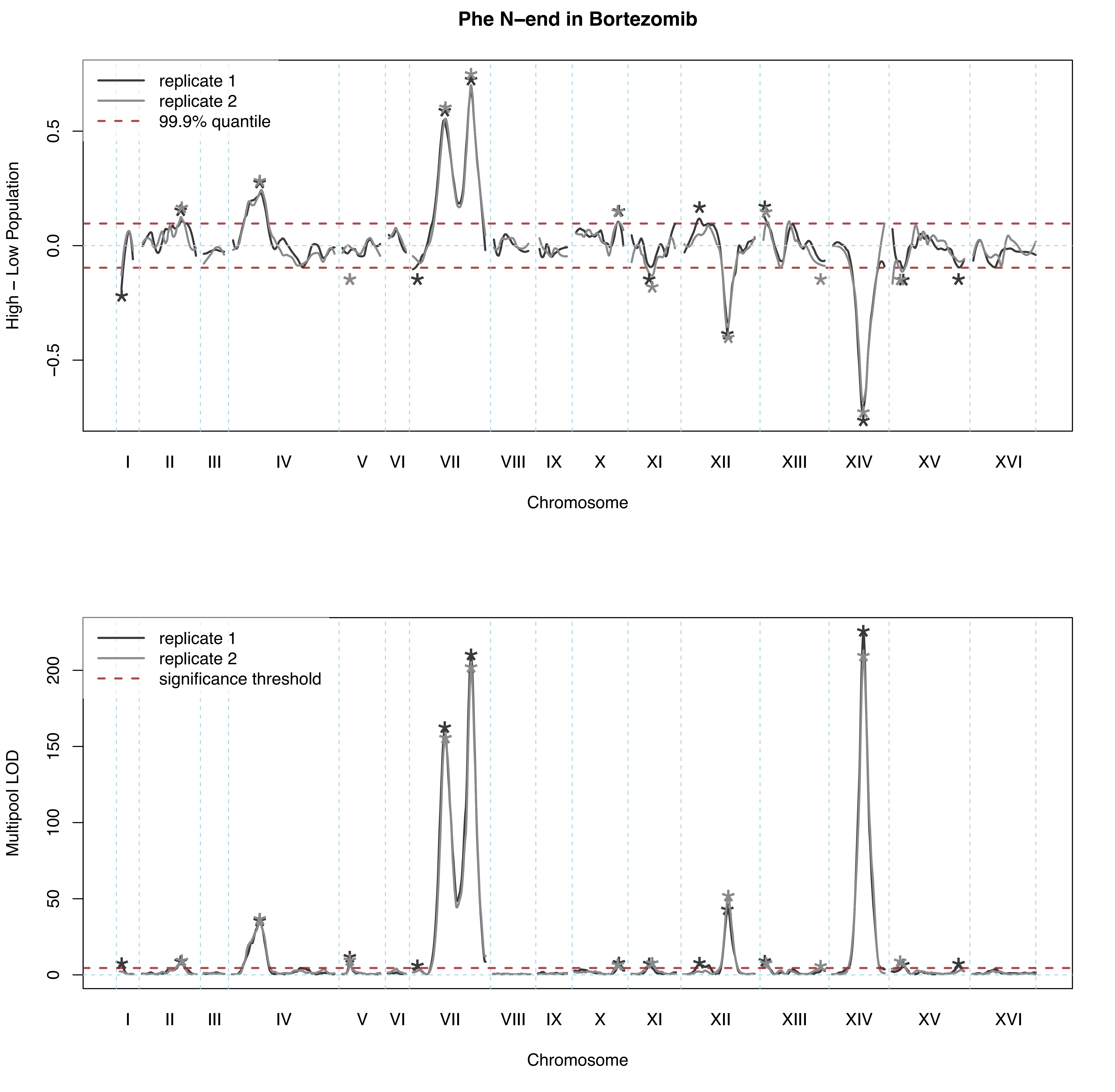
Phe N–end in Bortezomib
replicate 1
replicate 2
99.9% quantile
High – Low Population
0.5
0.0
−0.5
I II III IV V VI VII VIII IX X XI XII XIII XIV XV XVI
Chromosome
replicate 1
replicate 2
significance threshold
Multipool LOD
0
50
100
150
200
I II III IV V VI VII VIII IX X XI XII XIII XIV XV XVI
Chromosome

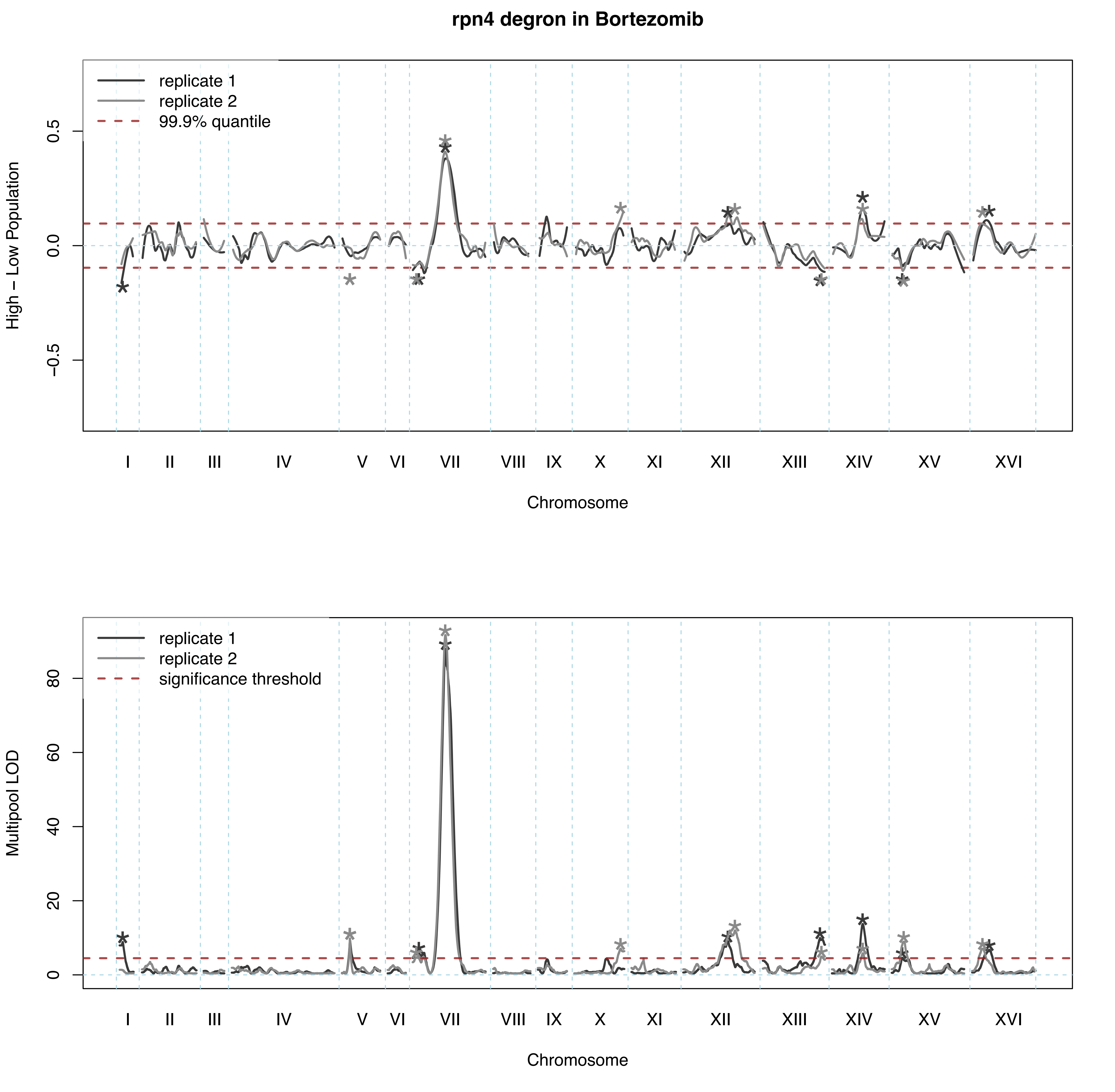
rpn4 degron in Bortezomib
replicate 1
replicate 2
99.9% quantile
High – Low Population
0.5
0.0
−0.5
I
II
III
IV
V
VI
VII
VIII
IX
X
XI
XII
XIII
XIV
XV
XVI
Chromosome
replicate 1
replicate 2
significance threshold
Multipool LOD
80
60
40
20
0
I
II
III
IV
V
VI
VII
VIII
IX
X
XI
XII
XIII
XIV
XV
XVI
Chromosome

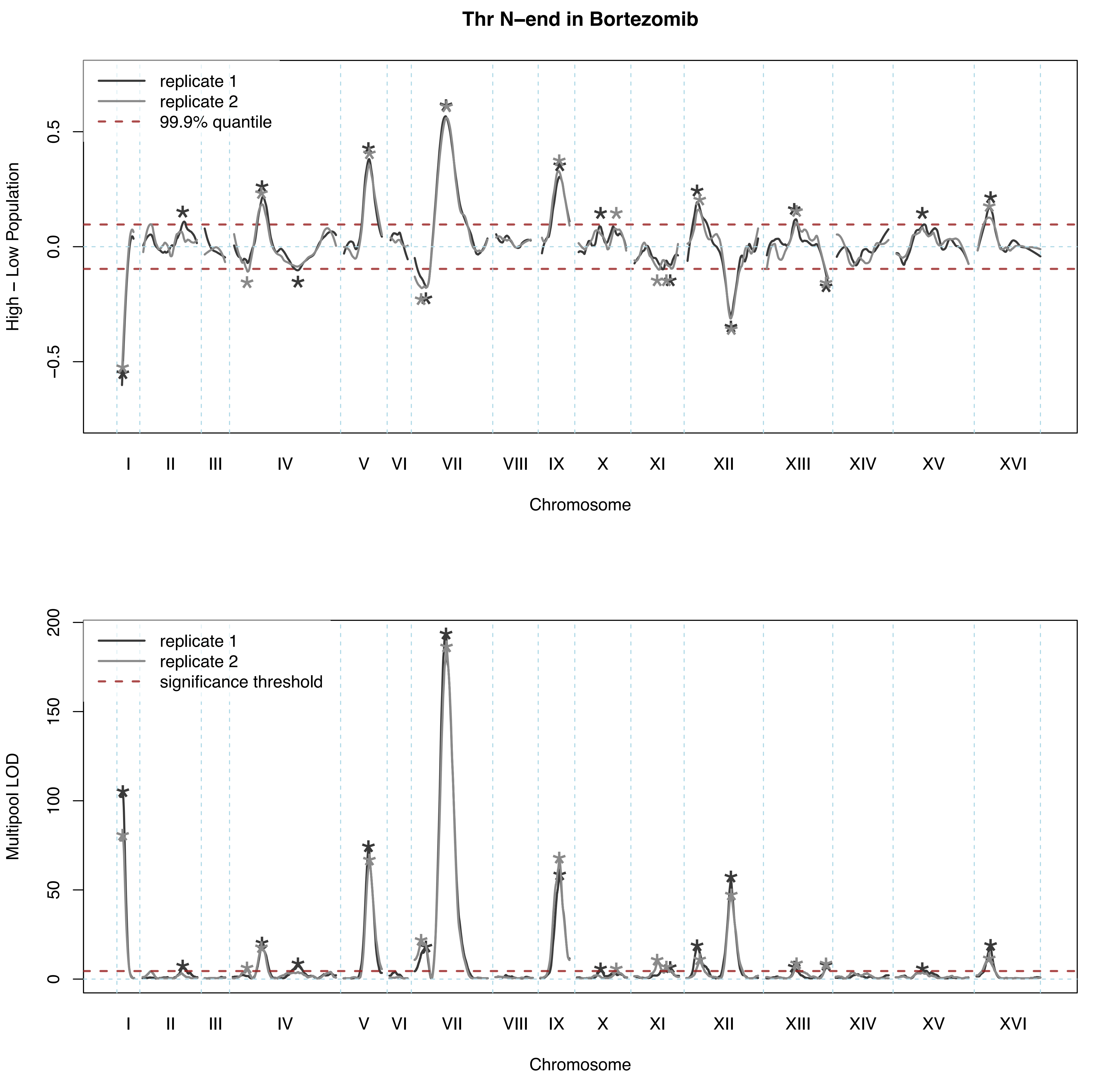

Thr N–end in Bortezomib
replicate 1
replicate 2
99.9% quantile
High – Low Population
0.5
0.0
−0.5
I
II
III
IV
V
VI
VII
VIII
IX
X
XI
XII
XIII
XIV
XV
XVI
Chromosome
replicate 1
replicate 2
significance threshold
Multipool LOD
200
150
100
50
0
I
II
III
IV
V
VI
VII
VIII
IX
X
XI
XII
XIII
XIV
XV
XVI
Chromosome

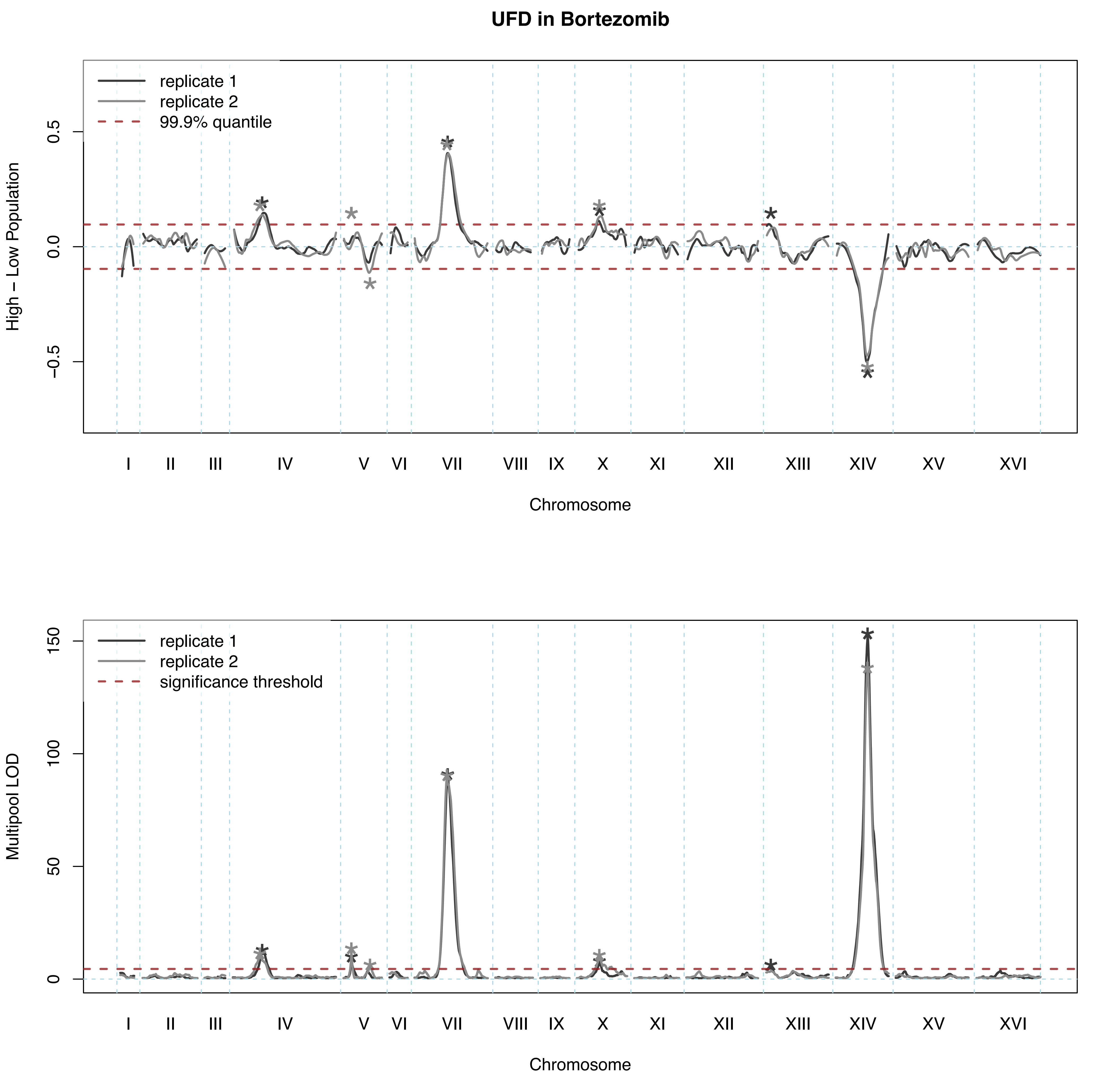

UFD in Bortezomib
replicate 1
replicate 2
99.9% quantile
High – Low Population
0.5
0.0
−0.5
I
II
III
IV
V
VI
VII
VIII
IX
X
XI
XII
XIII
XIV
XV
XVI
Chromosome
replicate 1
replicate 2
significance threshold
Multipool LOD
150
100
50
0
I
II
III
IV
V
VI
VII
VIII
IX
X
XI
XII
XIII
XIV
XV
XVI
Chromosome

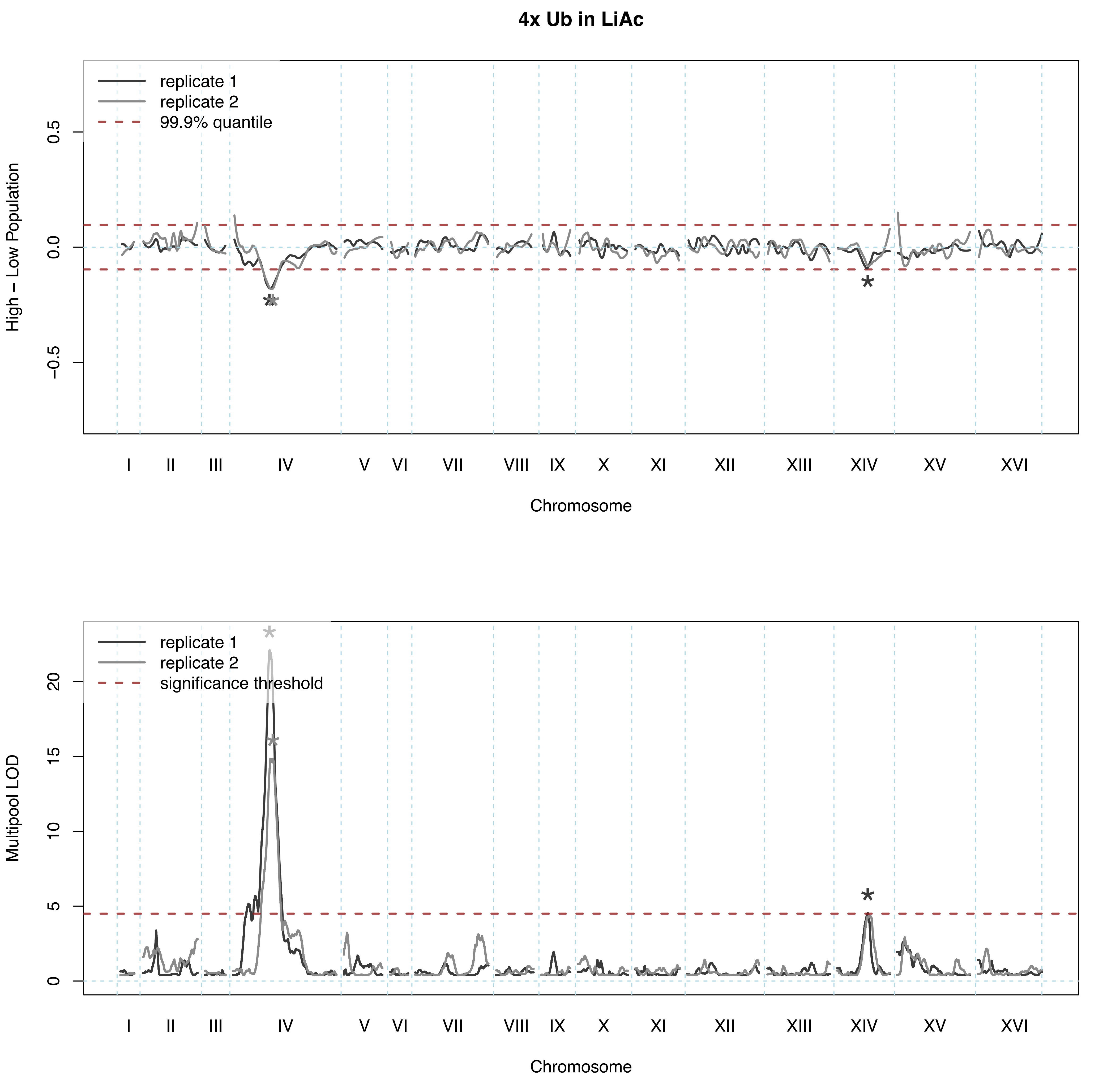

4x Ub in LiAc
replicate 1
replicate 2
99.9% quantile
High – Low Population
0.5
0.0
−0.5
I
II
III
IV
V
VI
VII
VIII
IX
X
XI
XII
XIII
XIV
XV
XVI
Chromosome
replicate 1
replicate 2
significance threshold
Multipool LOD
20
15
10
5
0
I
II
III
IV
V
VI
VII
VIII
IX
X
XI
XII
XIII
XIV
XV
XVI
Chromosome

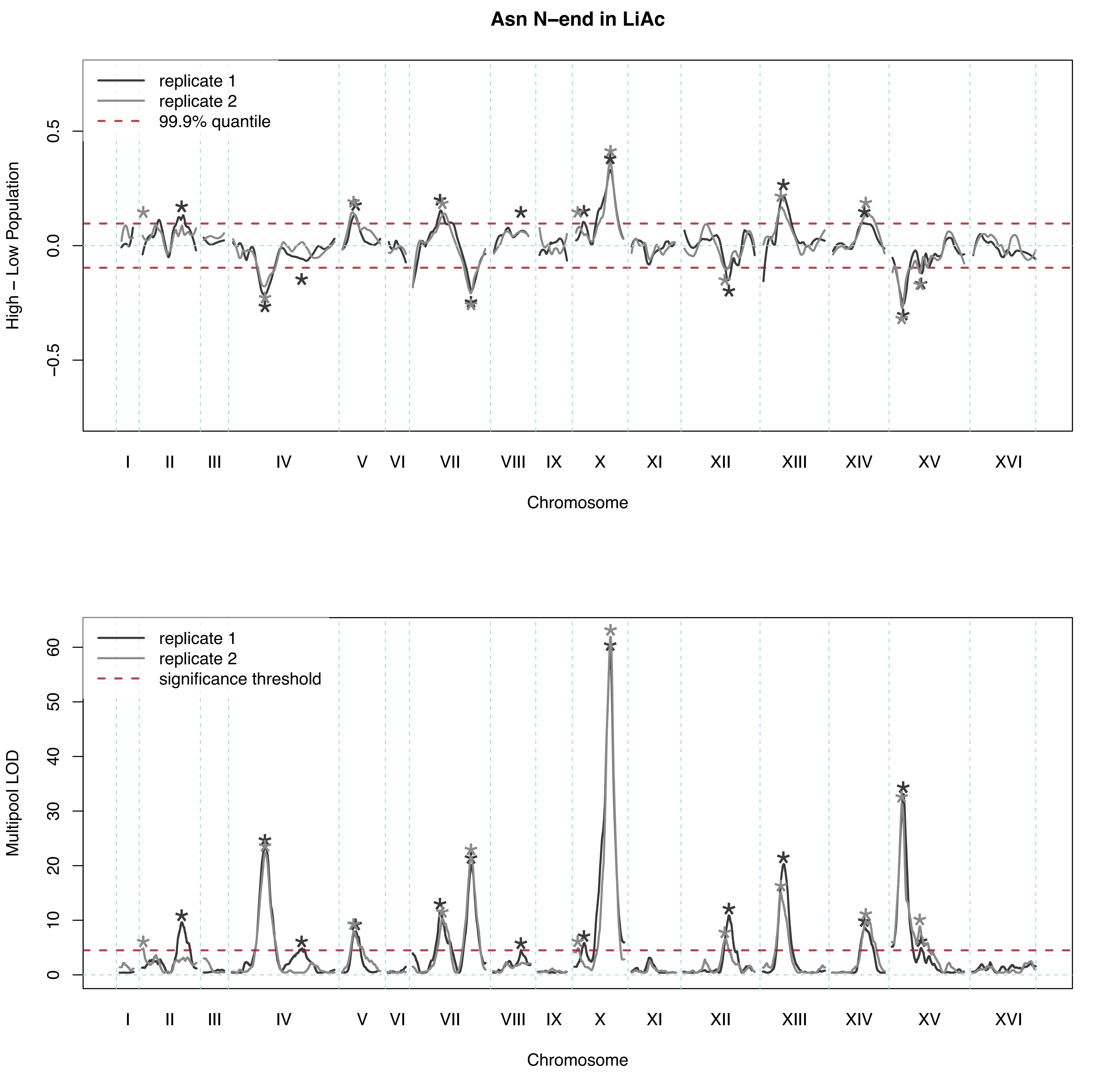

Asn N–end in LiAc
replicate 1
replicate 2
99.9% quantile
High – Low Population
0.5
0.0
−0.5
I
II
III
IV
V
VI
VII
VIII
IX
X
XI
XII
XIII
XIV
XV
XVI
Chromosome
replicate 1
replicate 2
significance threshold
Multipool LOD
0
10
20
30
40
50
60
I
II
III
IV
V
VI
VII
VIII
IX
X
XI
XII
XIII
XIV
XV
XVI
Chromosome

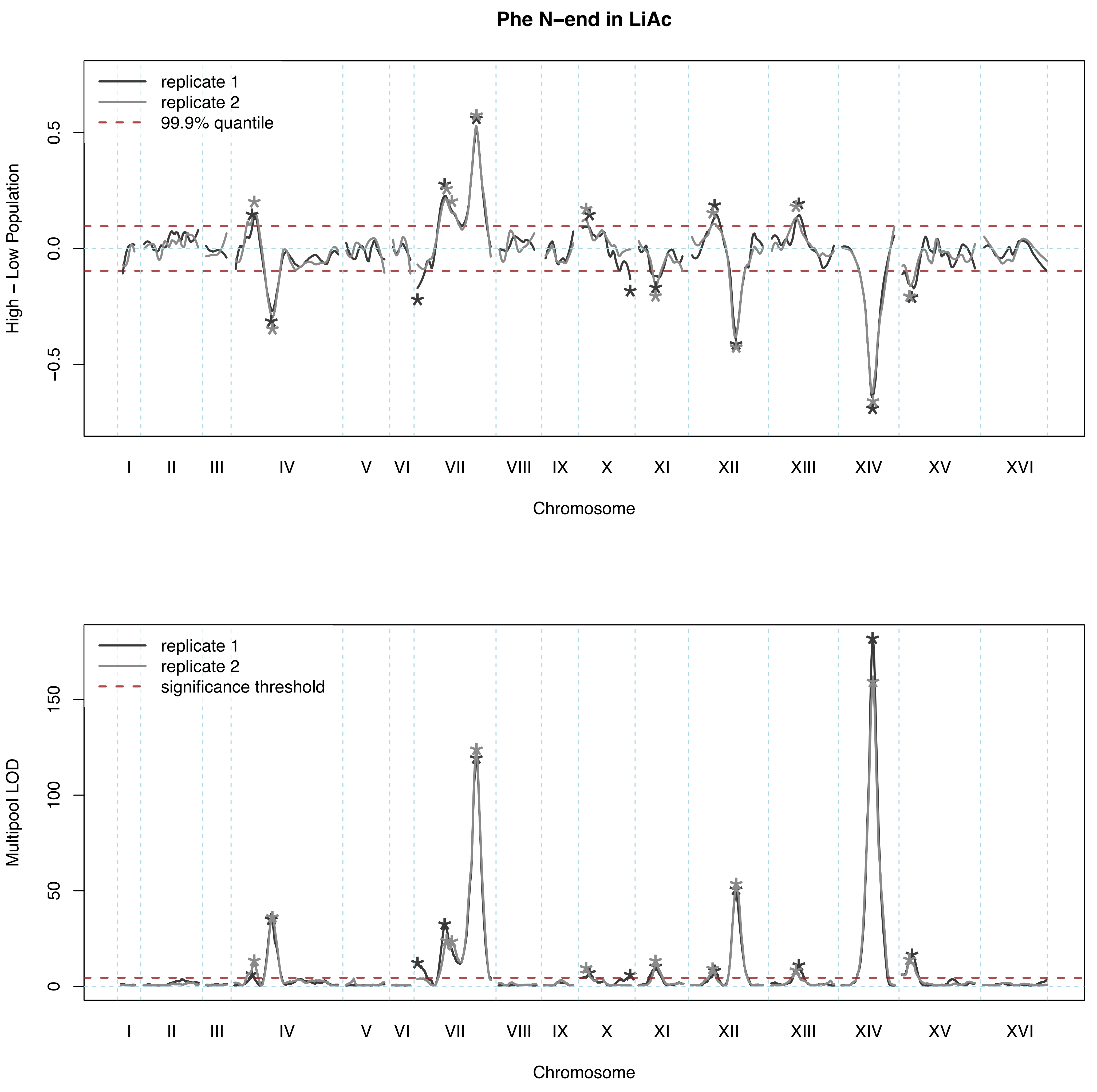

Phe N–end in LiAc
replicate 1
replicate 2
99.9% quantile
High – Low Population
0.5
0.0
−0.5
I
II
III
IV
V
VI
VII
VIII
IX
X
XI
XII
XIII
XIV
XV
XVI
Chromosome
replicate 1
replicate 2
significance threshold
Multipool LOD
150
100
50
0
I
II
III
IV
V
VI
VII
VIII
IX
X
XI
XII
XIII
XIV
XV
XVI
Chromosome

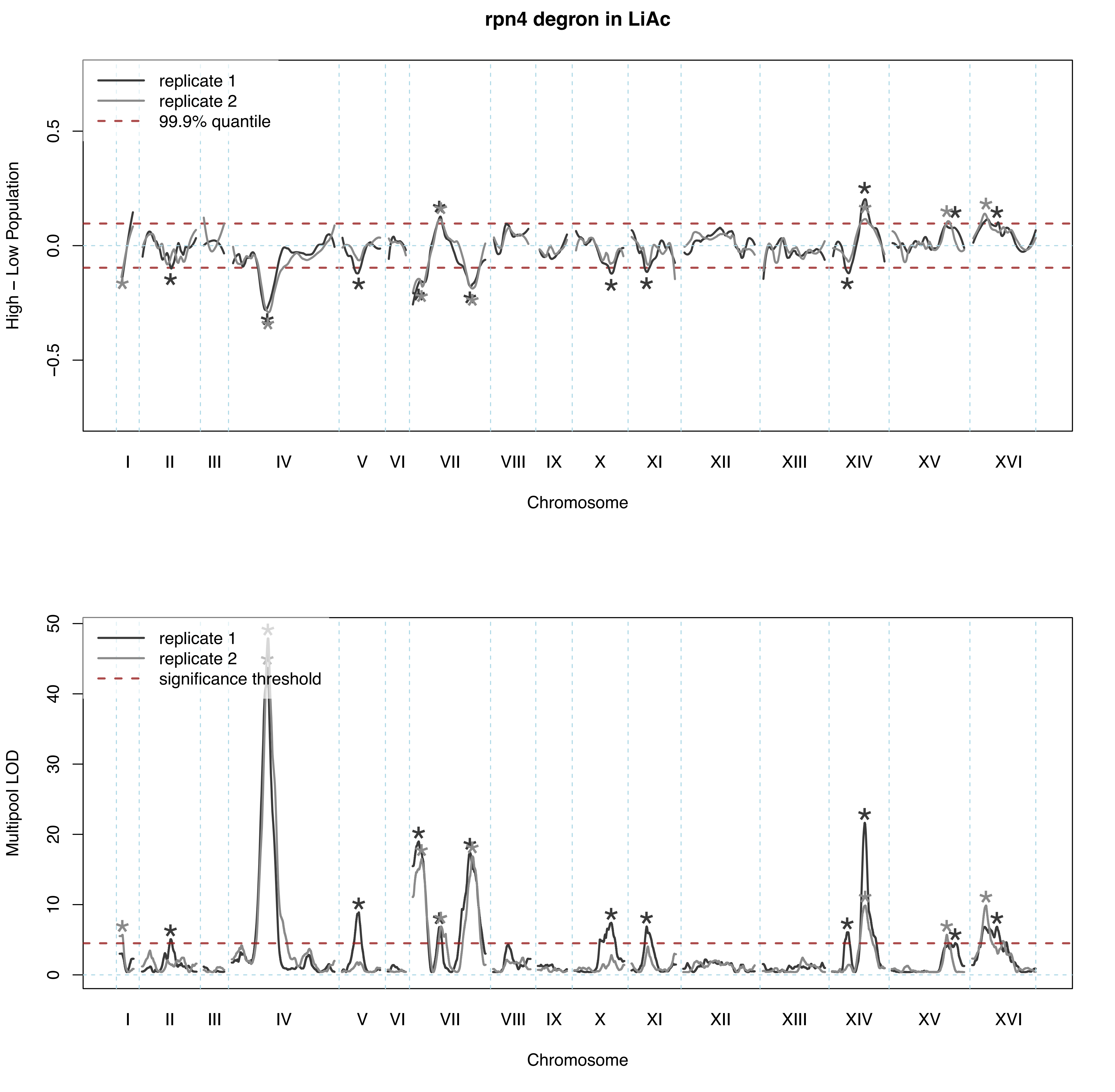
rpn4 degron in LiAc
replicate 1
replicate 2
99.9% quantile
High – Low Population
0.5
0.0
−0.5
I
II
III
IV
V
VI
VII
VIII
IX
X
XI
XII
XIII
XIV
XV
XVI
Chromosome
replicate 1
replicate 2
significance threshold
Multipool LOD
50
40
30
20
10
0
I
II
III
IV
V
VI
VII
VIII
IX
X
XI
XII
XIII
XIV
XV
XVI
Chromosome

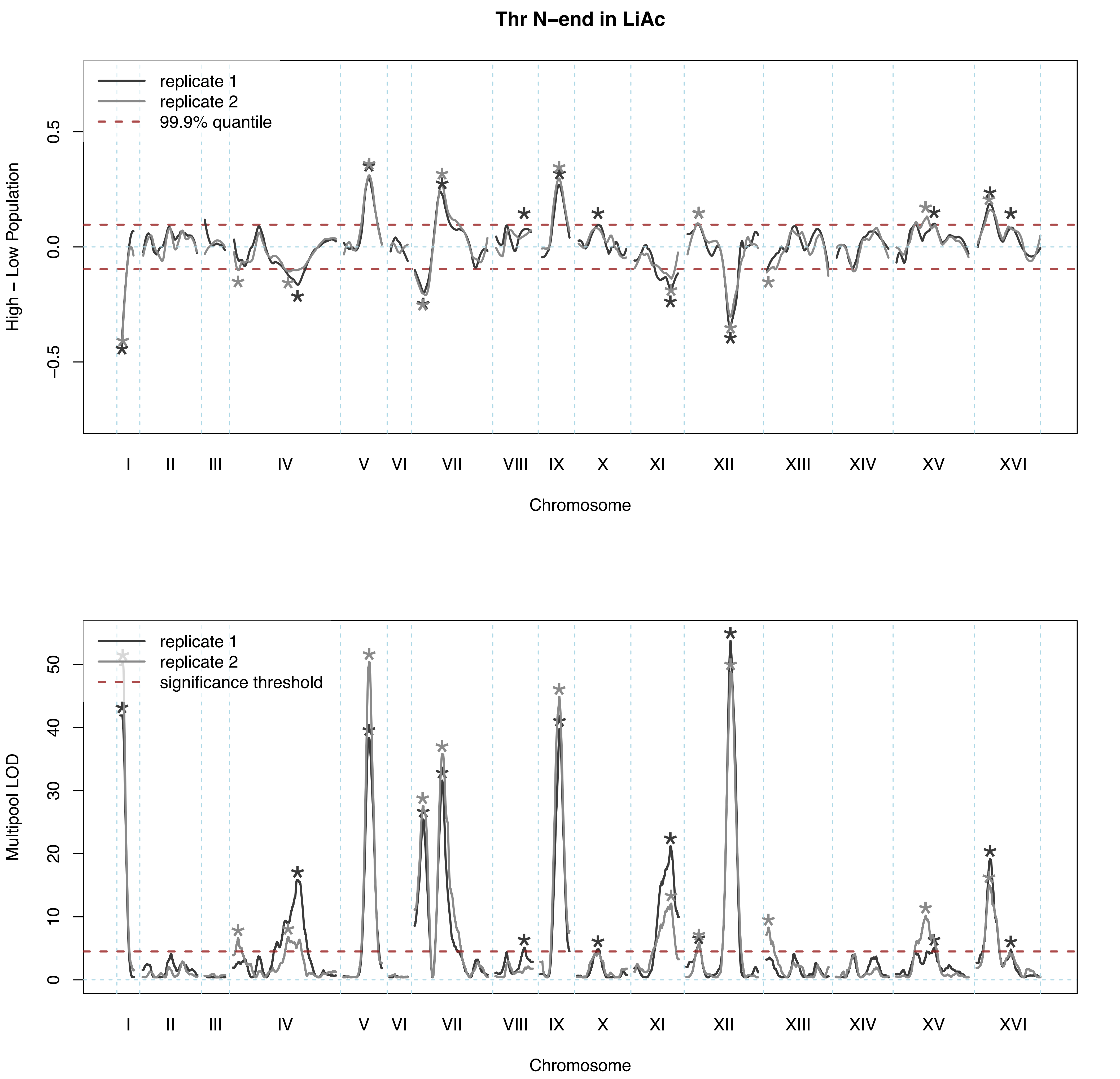

Thr N–end in LiAc
replicate 1
replicate 2
99.9% quantile
High – Low Population
0.5
0.0
−0.5
I
II
III
IV
V
VI
VII
VIII
IX
X
XI
XII
XIII
XIV
XV
XVI
Chromosome
replicate 1
replicate 2
significance threshold
Multipool LOD
0
10
20
30
40
50
Chromosome

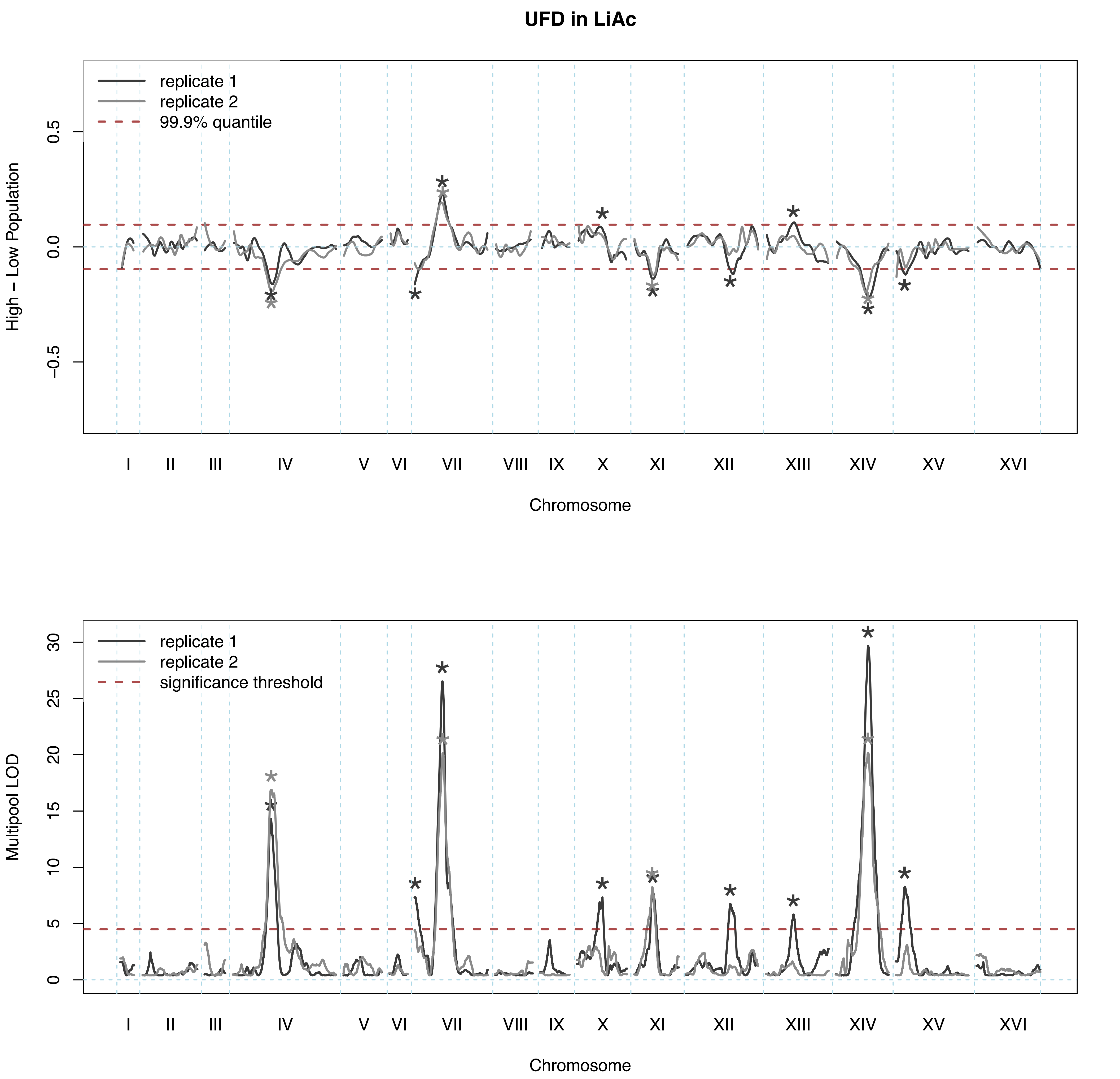
UFD in LiAc
replicate 1
replicate 2
99.9% quantile
High – Low Population
0.5
0.0
−0.5
I II III IV V VI VII VIII IX X XI XII XIII XIV XV XVI
Chromosome
replicate 1
replicate 2
significance threshold
Multipool LOD
30
25
20
15
10
5
0
I II III IV V VI VII VIII IX X XI XII XIII XIV XV XVI
Chromosome

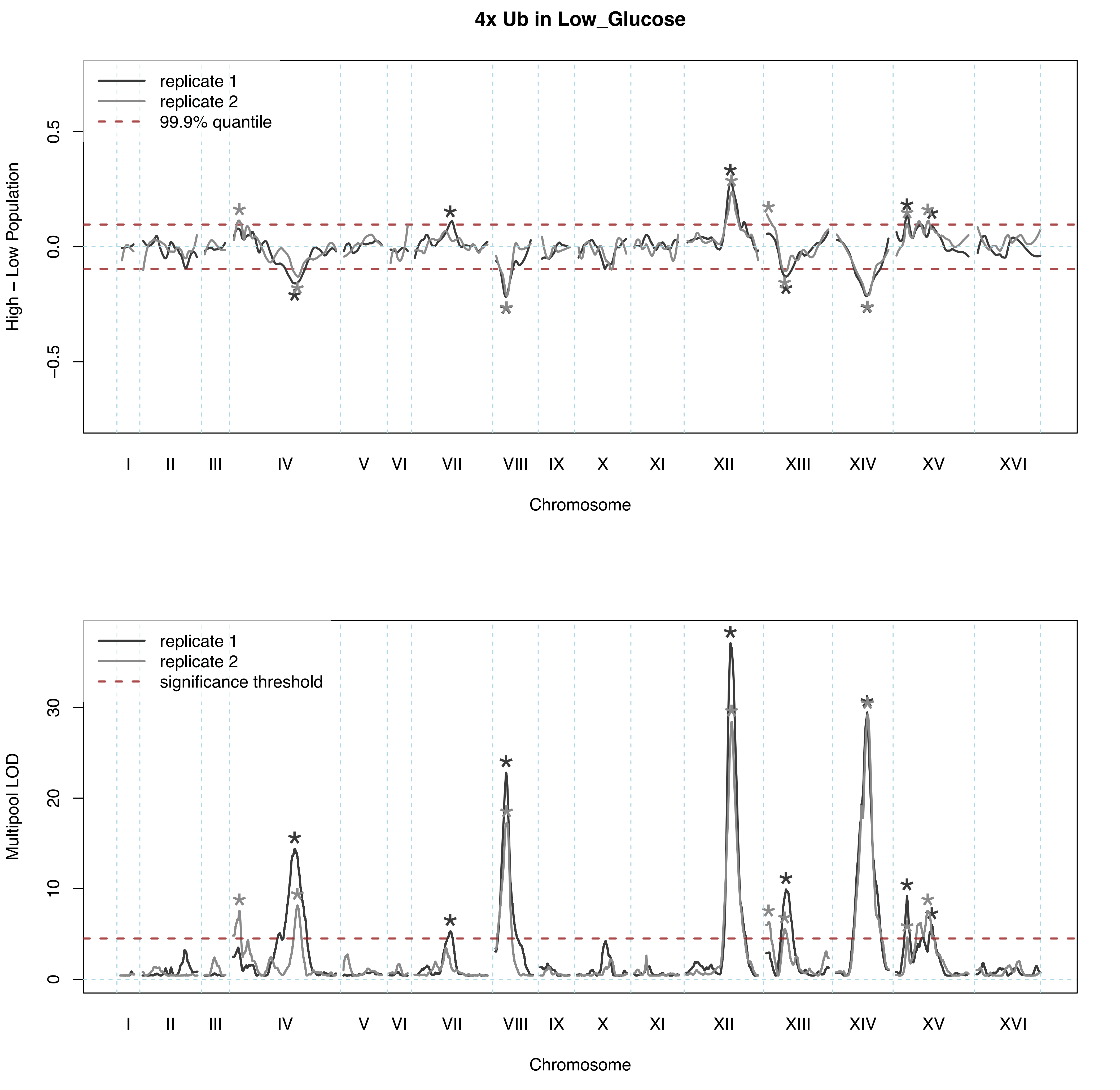

4x Ub in Low_Glucose
replicate 1
replicate 2
99.9% quantile
High – Low Population
0.5
0.0
−0.5
I
II
III
IV
V
VI
VII
VIII
IX
X
XI
XII
XIII
XIV
XV
XVI
Chromosome
replicate 1
replicate 2
significance threshold
Multipool LOD
30
20
10
0
Chromosome

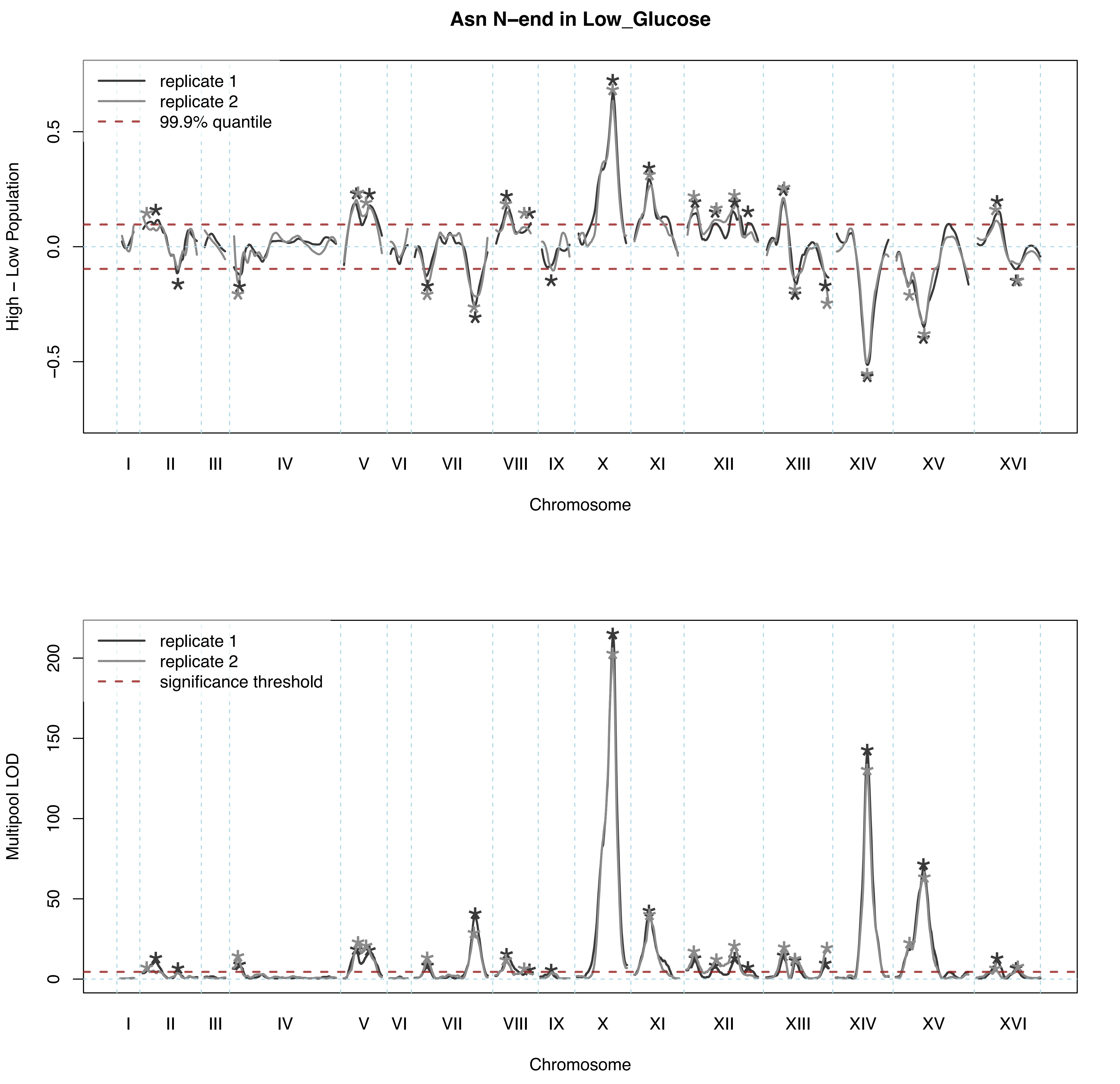

Asn N–end in Low_Glucose
replicate 1
replicate 2
99.9% quantile
High – Low Population
0.5
0.0
−0.5
I
II
III
IV
V
VI
VII
VIII
IX
X
XI
XII
XIII
XIV
XV
XVI
Chromosome
replicate 1
replicate 2
significance threshold
Multipool LOD
200
150
100
50
0
I
II
III
IV
V
VI
VII
VIII
IX
X
XI
XII
XIII
XIV
XV
XVI
Chromosome

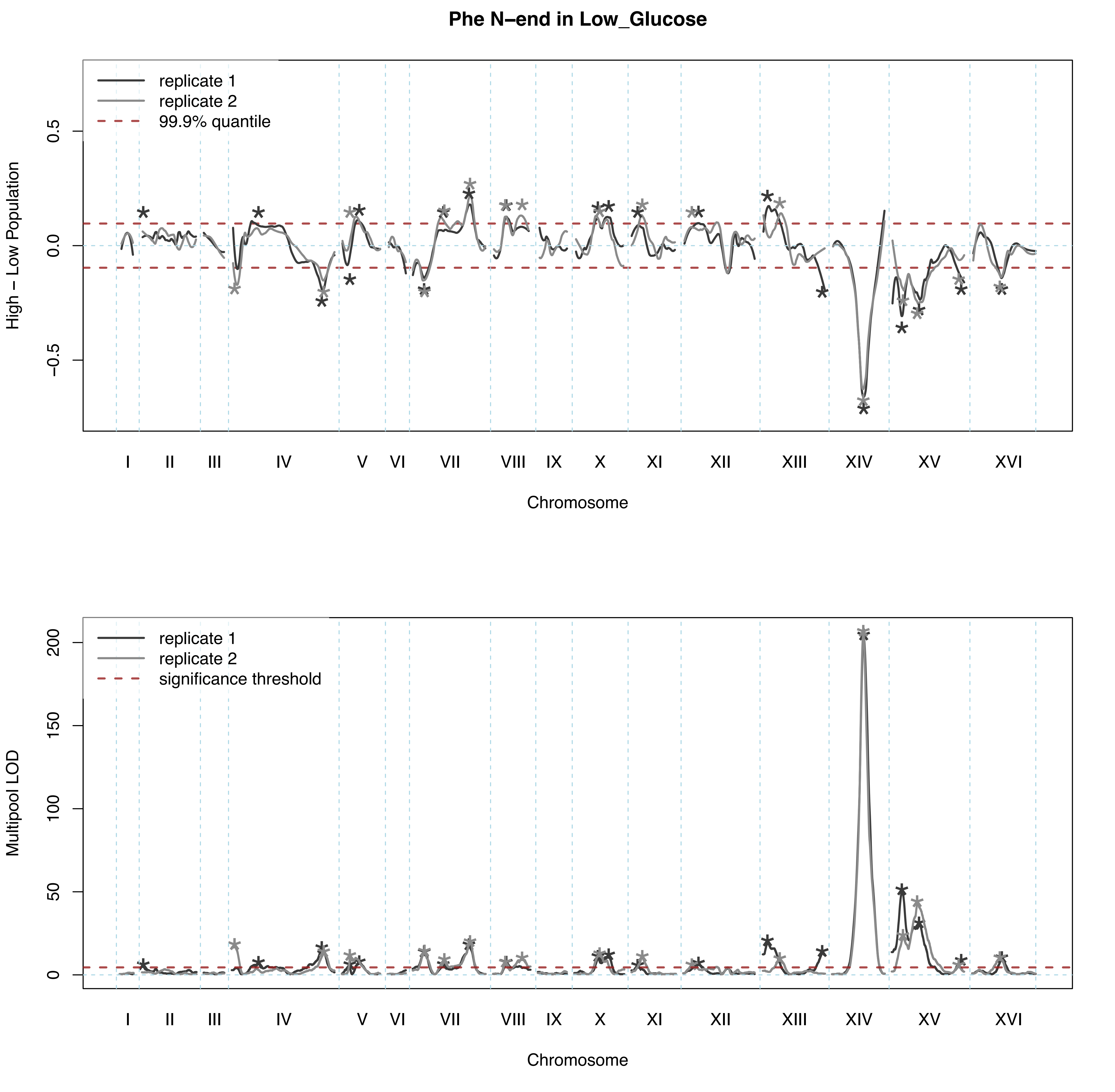

Phe N−end in Low_Glucose
replicate 1
replicate 2
99.9% quantile
High − Low Population
0.5
0.0
−0.5
I
II
III
IV
V
VI
VII
VIII
IX
X
XI
XII
XIII
XIV
XV
XVI
Chromosome
replicate 1
replicate 2
significance threshold
Multipool LOD
200
150
100
50
0
I
II
III
IV
V
VI
VII
VIII
IX
X
XI
XII
XIII
XIV
XV
XVI
Chromosome

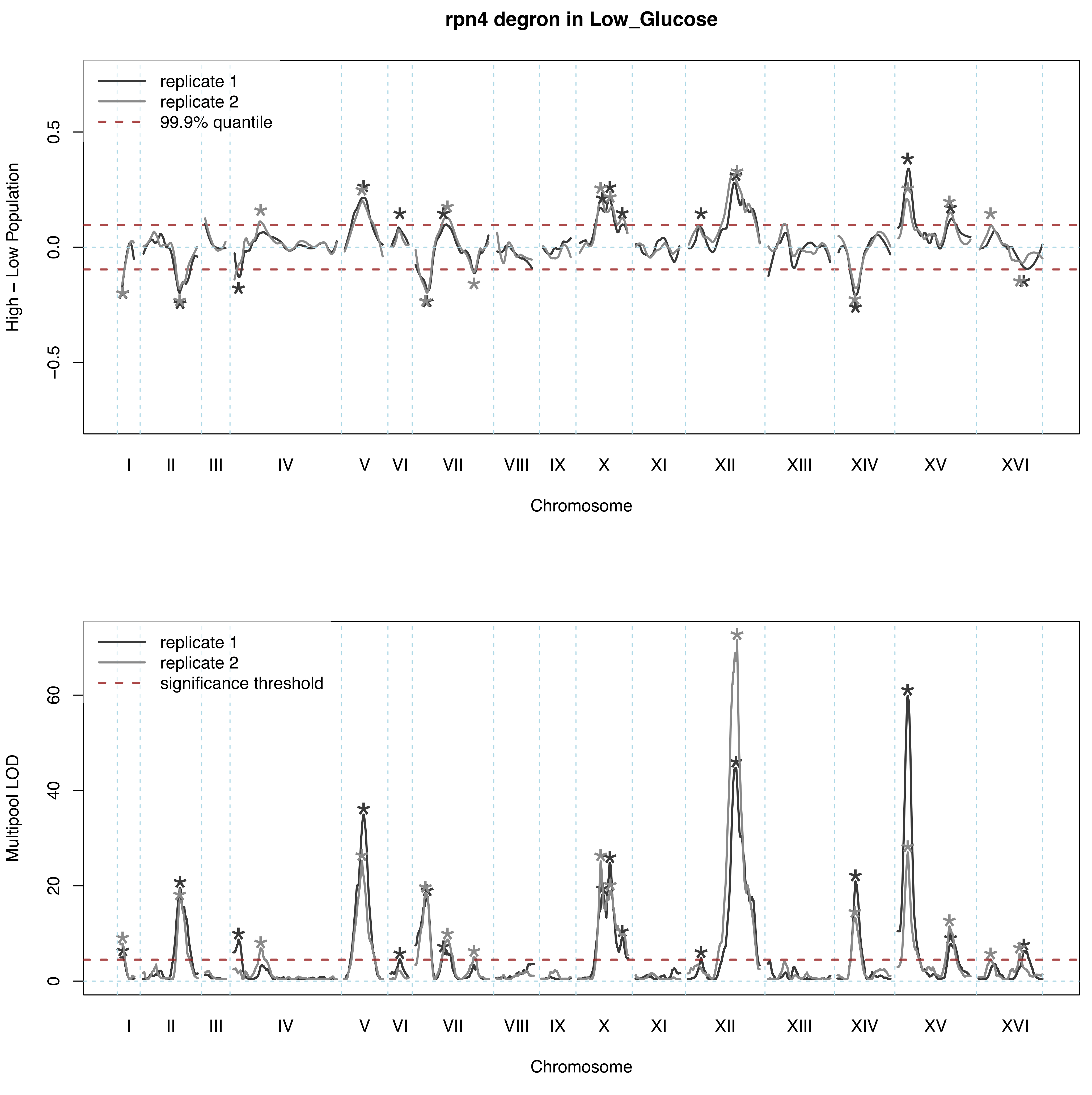
rpn4 degron in Low_Glucose
replicate 1
replicate 2
99.9% quantile
High − Low Population
0.5
0.0
−0.5
I II III IV V VI VII VIII IX X XI XII XIII XIV XV XVI
Chromosome
replicate 1
replicate 2
significance threshold
Multipool LOD
60
40
20
0
I II III IV V VI VII VIII IX X XI XII XIII XIV XV XVI
Chromosome

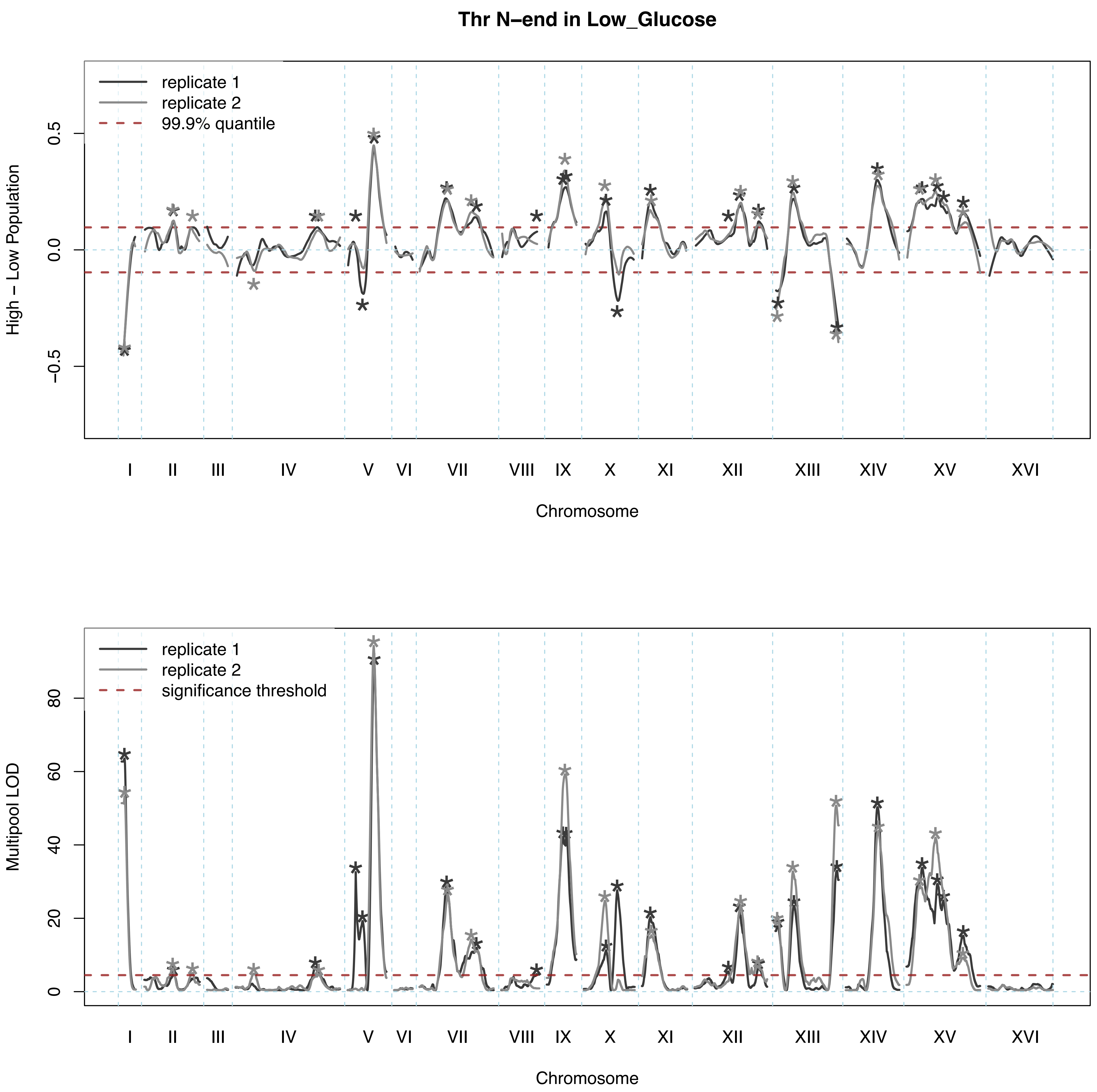
Thr N–end in Low_Glucose
replicate 1
replicate 2
99.9% quantile
High – Low Population
0.5
0.0
−0.5
I II III IV V VI VII VIII IX X XI XII XIII XIV XV XVI
Chromosome
replicate 1
replicate 2
significance threshold
Multipool LOD
0
20
40
60
80
I II III IV V VI VII VIII IX X XI XII XIII XIV XV XVI
Chromosome

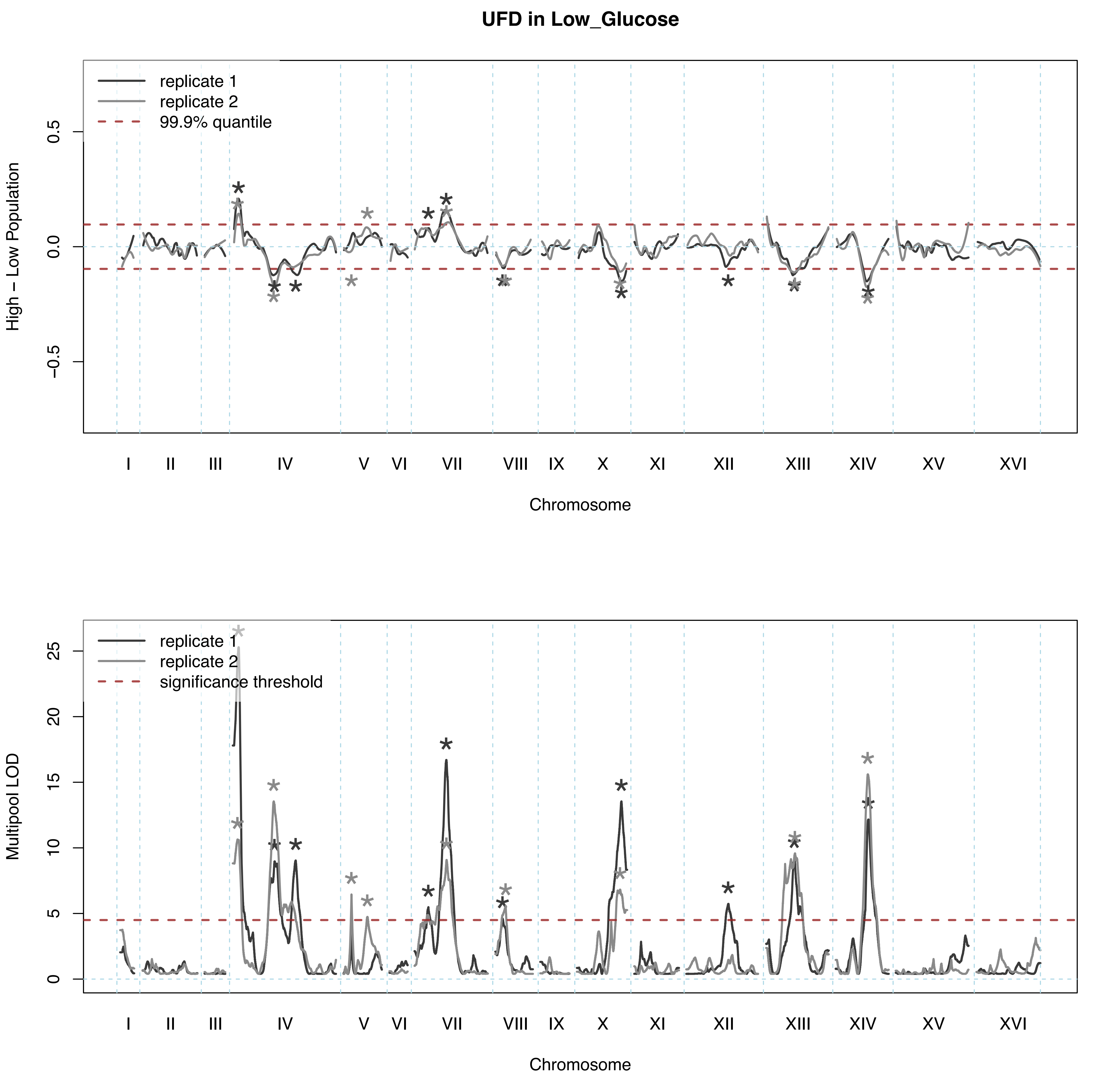

UFD in Low_Glucose
replicate 1
replicate 2
99.9% quantile
High − Low Population
0.5
0.0
−0.5
I
II
III
IV
V
VI
VII
VIII
IX
X
XI
XII
XIII
XIV
XV
XVI
Chromosome
replicate 1
replicate 2
significance threshold
Multipool LOD
25
20
15
10
5
0
I
II
III
IV
V
VI
VII
VIII
IX
X
XI
XII
XIII
XIV
XV
XVI
Chromosome

Asn N–end in Low_Nitrogen

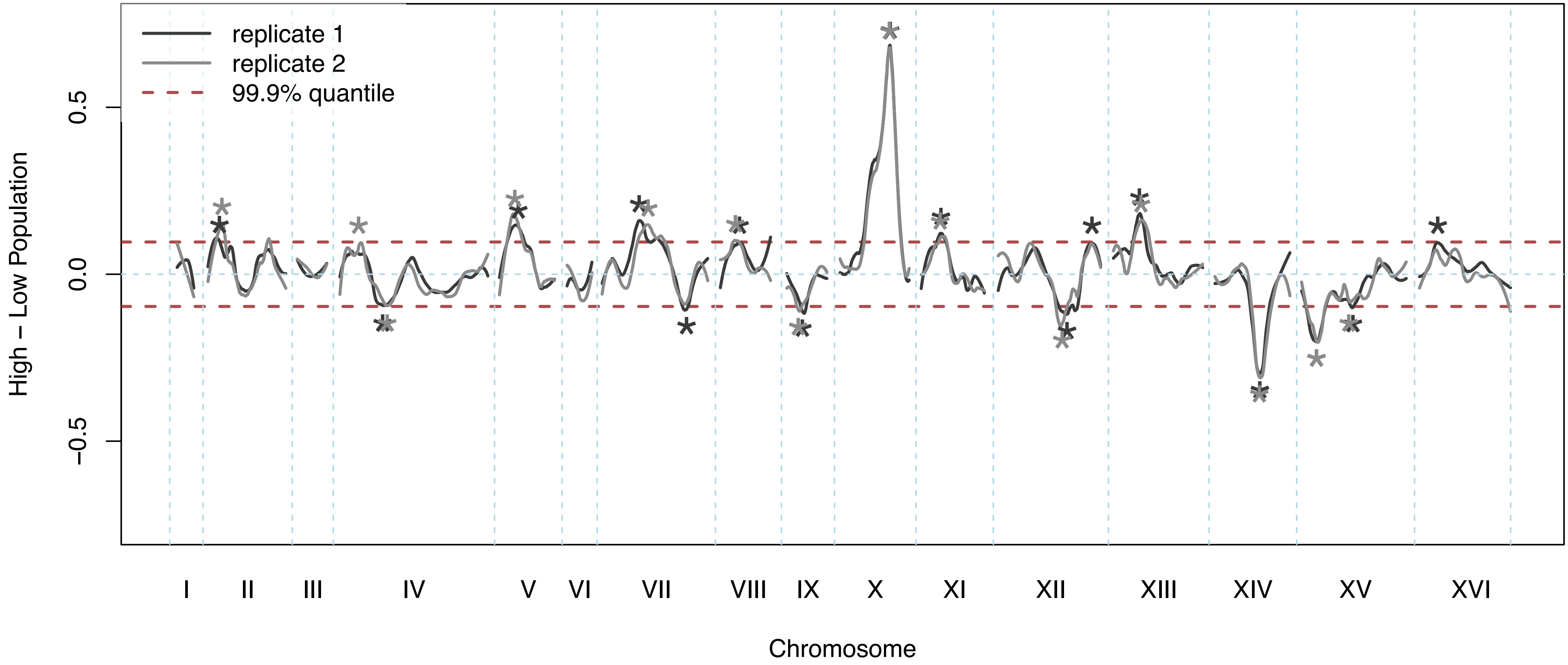


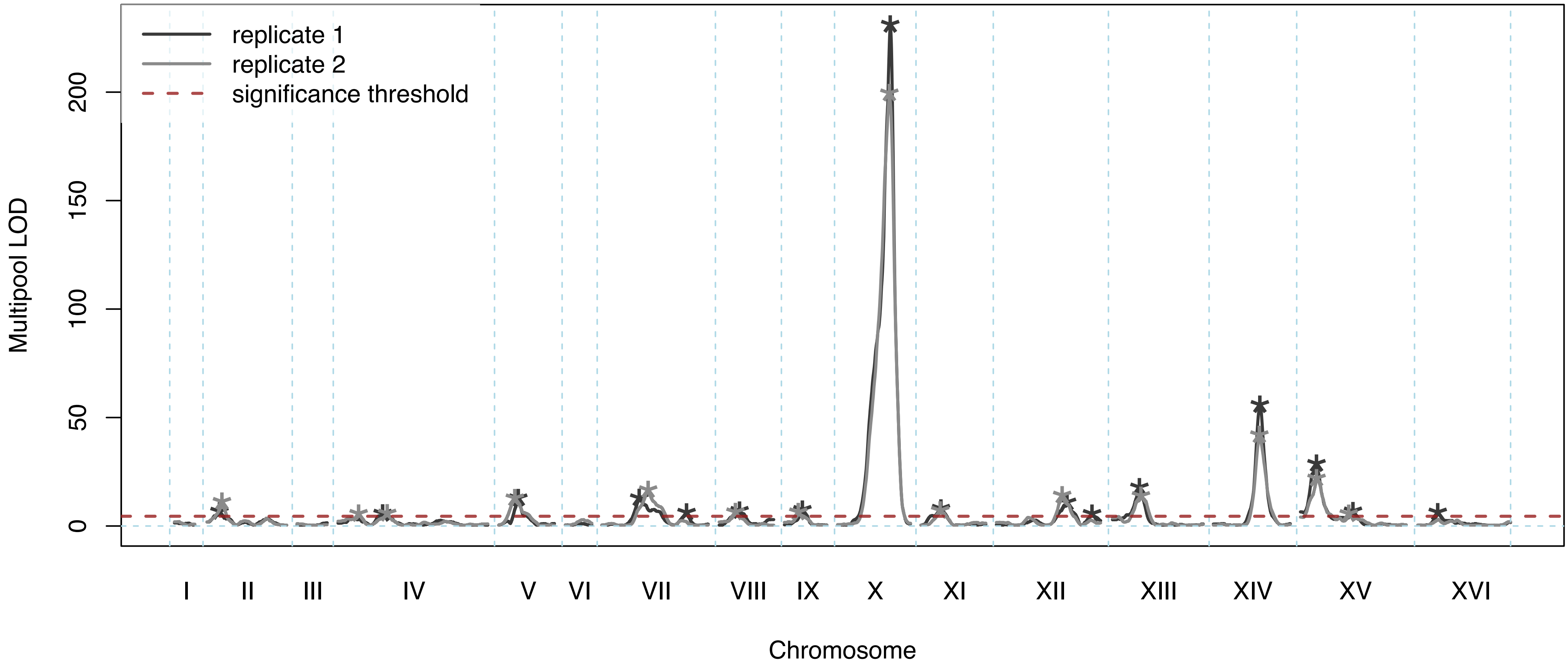

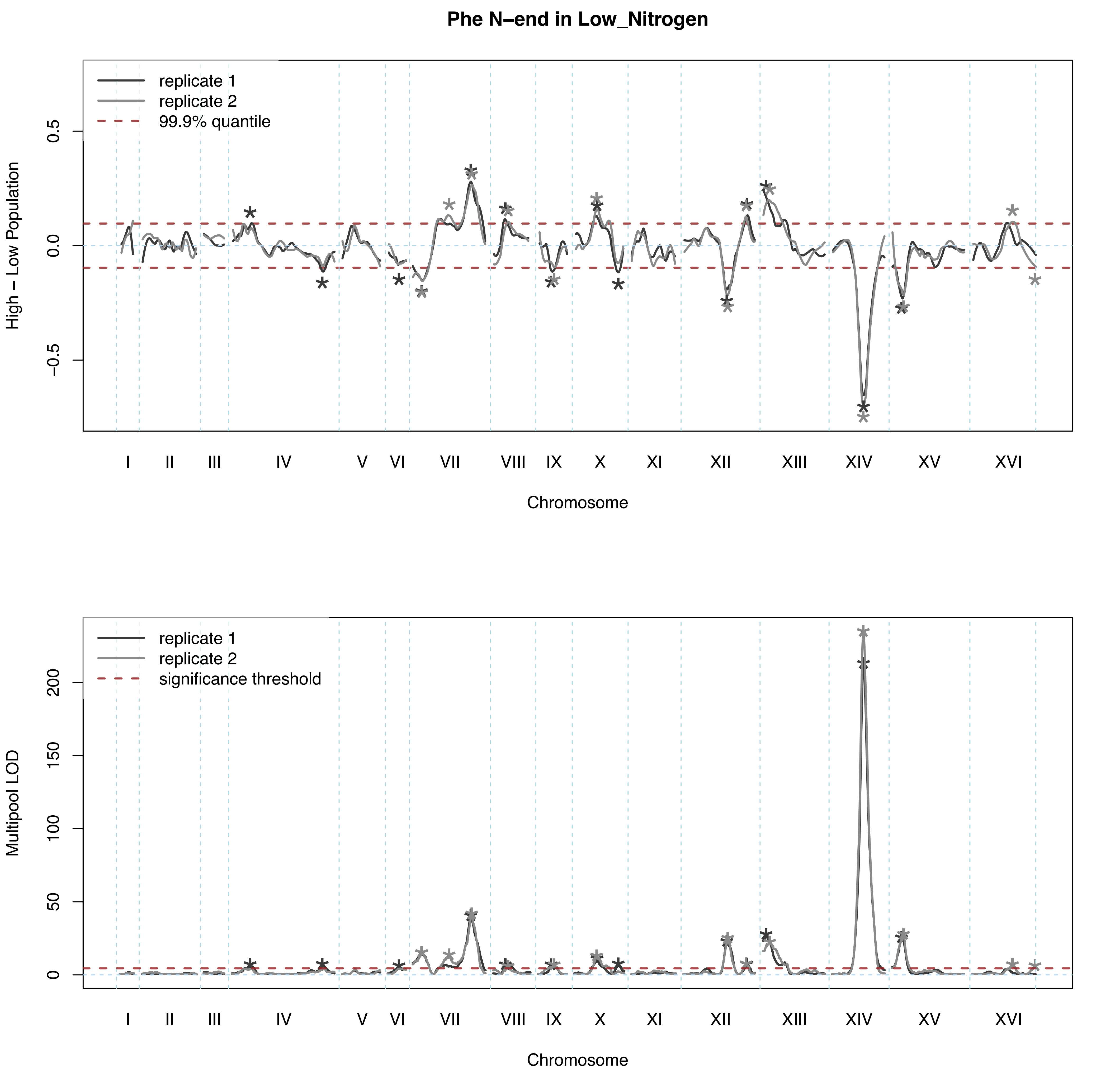

Phe N–end in Low_Nitrogen
replicate 1
replicate 2
99.9% quantile
High – Low Population
0.5
0.0
−0.5
I
II
III
IV
V
VI
VII
VIII
IX
X
XI
XII
XIII
XIV
XV
XVI
Chromosome
replicate 1
replicate 2
significance threshold
Multipool LOD
200
150
100
50
0
I
II
III
IV
V
VI
VII
VIII
IX
X
XI
XII
XIII
XIV
XV
XVI
Chromosome

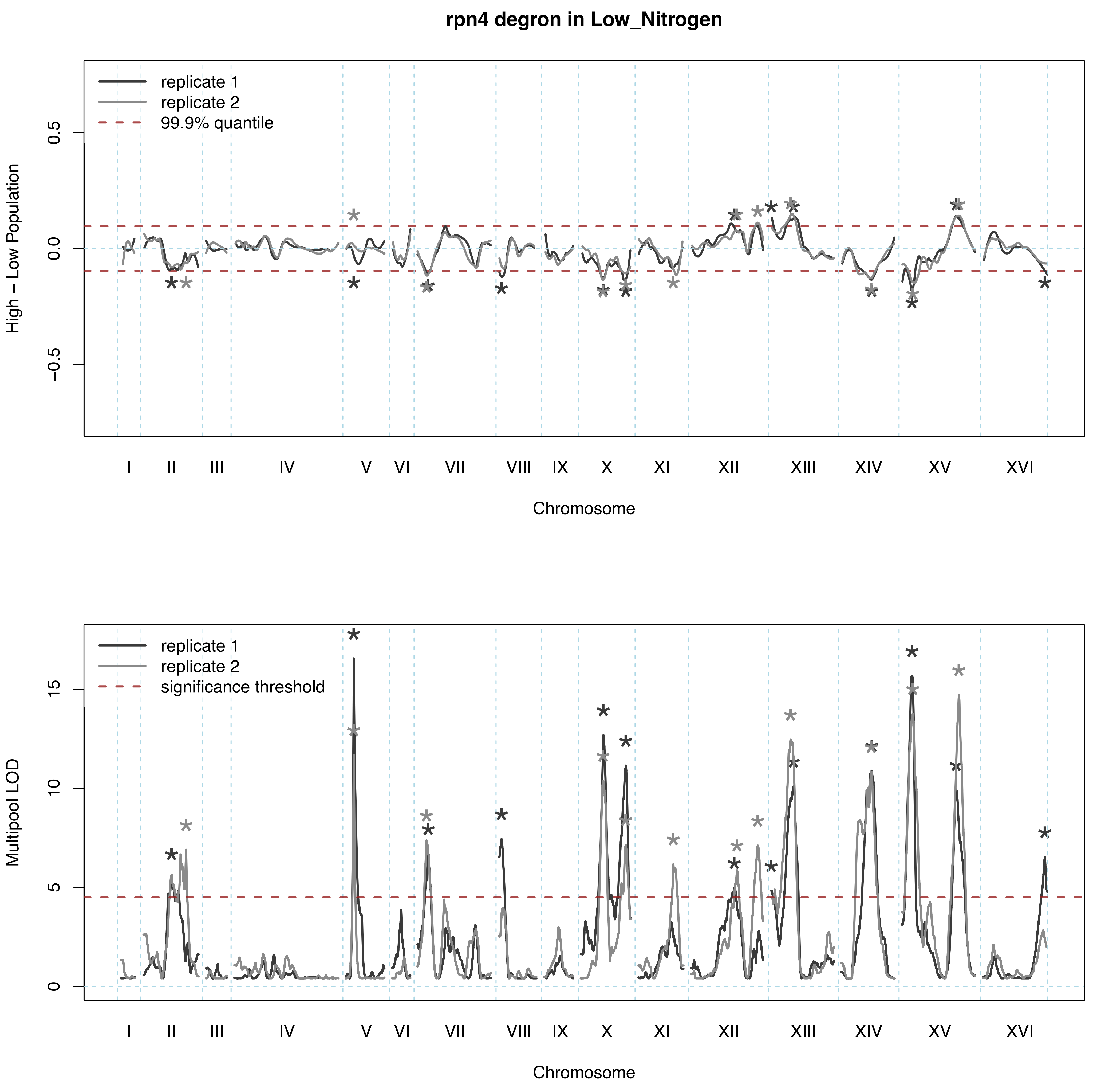
rpn4 degron in Low_Nitrogen
replicate 1
replicate 2
99.9% quantile
High − Low Population
0.5
0.0
−0.5
I
II
III
IV
V
VI
VII
VIII
IX
X
XI
XII
XIII
XIV
XV
XVI
Chromosome
replicate 1
replicate 2
significance threshold
Multipool LOD
15
10
5
0
I
II
III
IV
V
VI
VII
VIII
IX
X
XI
XII
XIII
XIV
XV
XVI
Chromosome

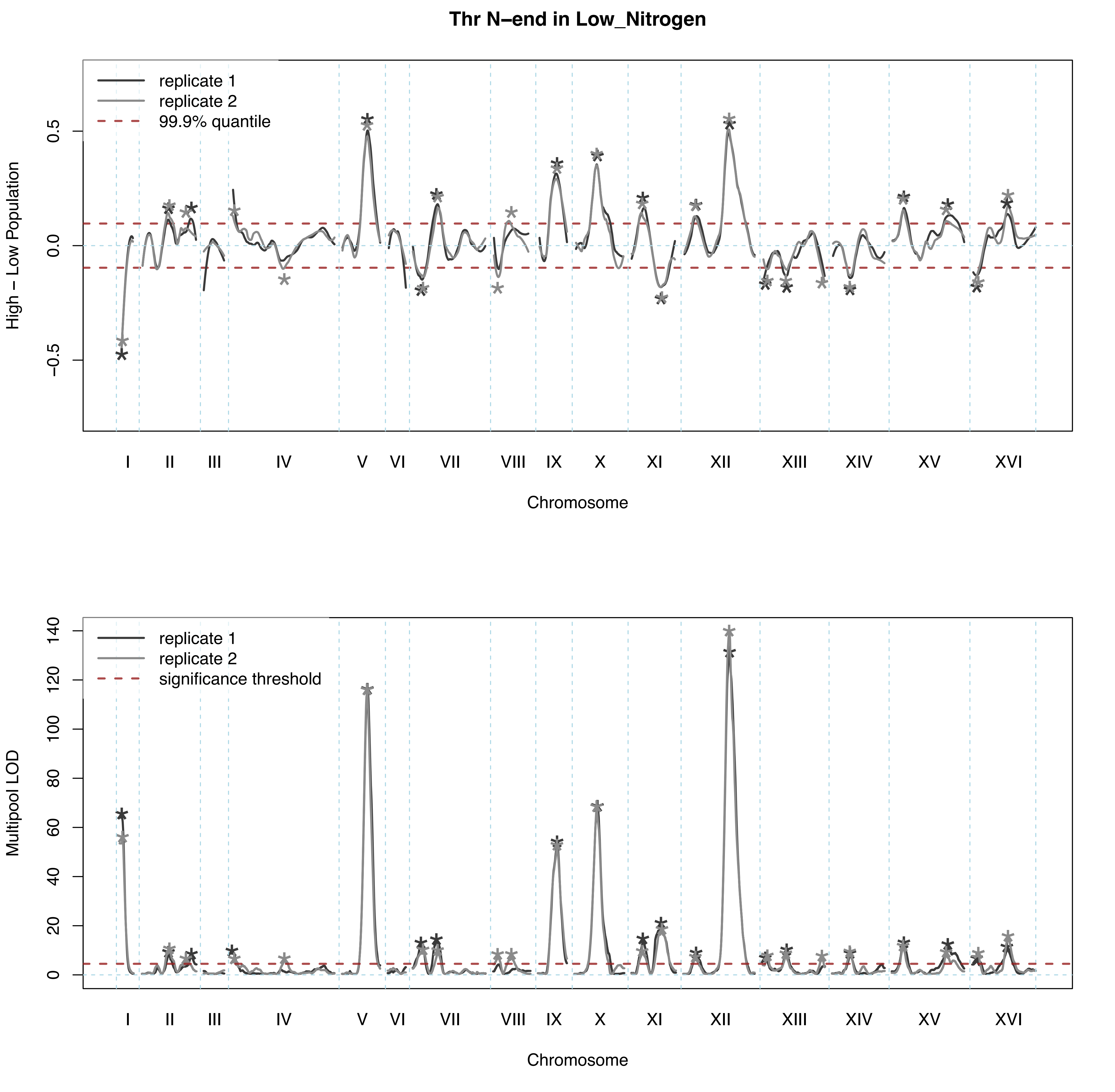

Thr N–end in Low_Nitrogen
replicate 1
replicate 2
99.9% quantile
High – Low Population
0.5
0.0
−0.5
I
II
III
IV
V
VI
VII
VIII
IX
X
XI
XII
XIII
XIV
XV
XVI
Chromosome
replicate 1
replicate 2
significance threshold
Multipool LOD
140
120
100
80
60
40
20
0
I
II
III
IV
V
VI
VII
VIII
IX
X
XI
XII
XIII
XIV
XV
XVI
Chromosome

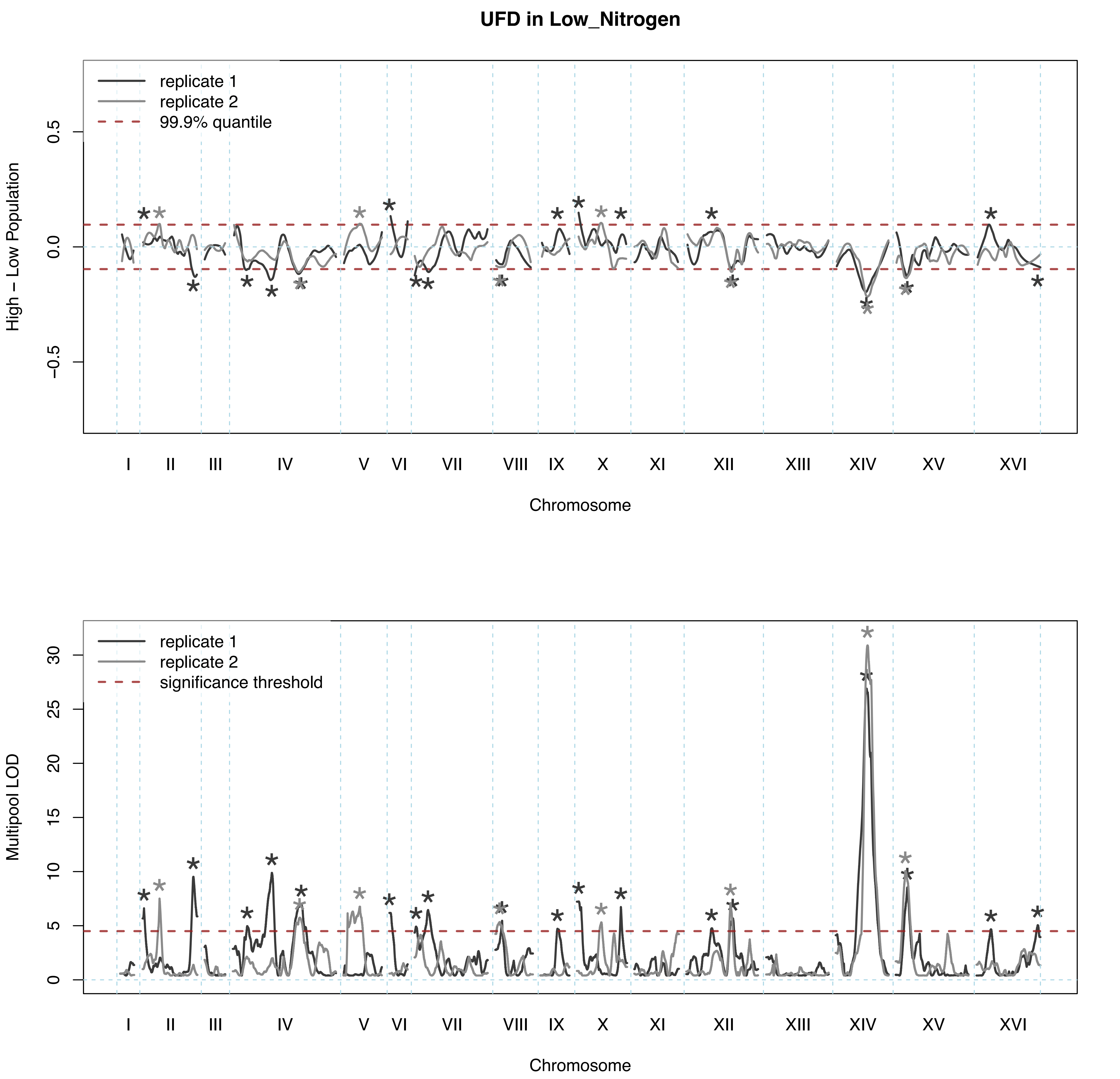
UFD in Low_Nitrogen
replicate 1
replicate 2
99.9% quantile
High – Low Population
0.5
0.0
−0.5
I
II
III
IV
V
VI
VII
VIII
IX
X
XI
XII
XIII
XIV
XV
XVI
Chromosome
replicate 1
replicate 2
significance threshold
Multipool LOD
0
5
10
15
20
25
30
I
II
III
IV
V
VI
VII
VIII
IX
X
XI
XII
XIII
XIV
XV
XVI
Chromosome

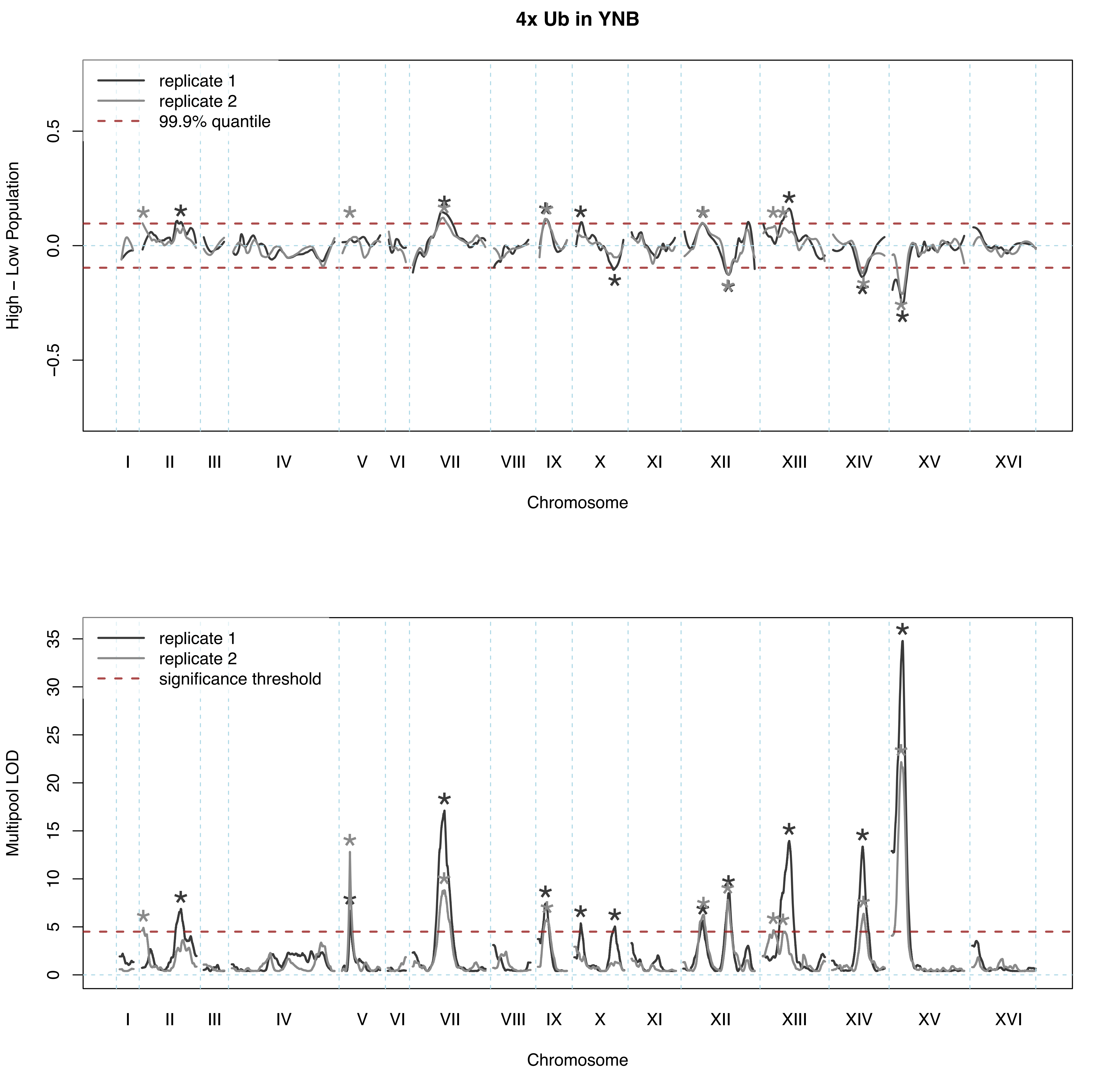

4x Ub in YNB
replicate 1
replicate 2
99.9% quantile
High – Low Population
0.5
0.0
−0.5
I II III IV V VI VII VIII IX X XI XII XIII XIV XV XVI
Chromosome
replicate 1
replicate 2
significance threshold
Multipool LOD
0 5 10 15 20 25 30 35
I II III IV V VI VII VIII IX X XI XII XIII XIV XV XVI
Chromosome

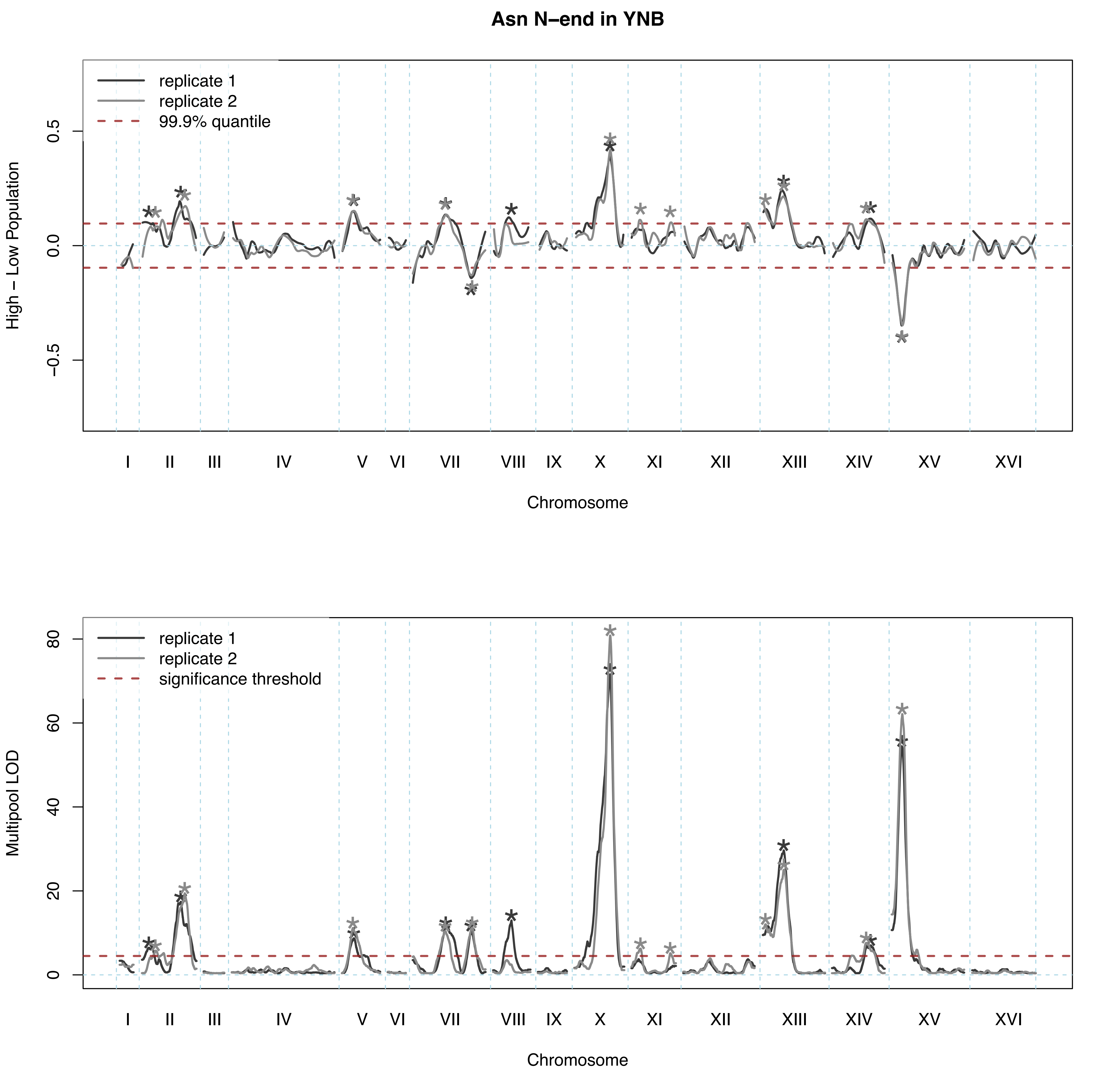

Asn N–end in YNB
replicate 1
replicate 2
99.9% quantile
High – Low Population
0.5
0.0
−0.5
I
II
III
IV
V
VI
VII
VIII
IX
X
XI
XII
XIII
XIV
XV
XVI
Chromosome
replicate 1
replicate 2
significance threshold
Multipool LOD
80
60
40
20
0
I
II
III
IV
V
VI
VII
VIII
IX
X
XI
XII
XIII
XIV
XV
XVI
Chromosome

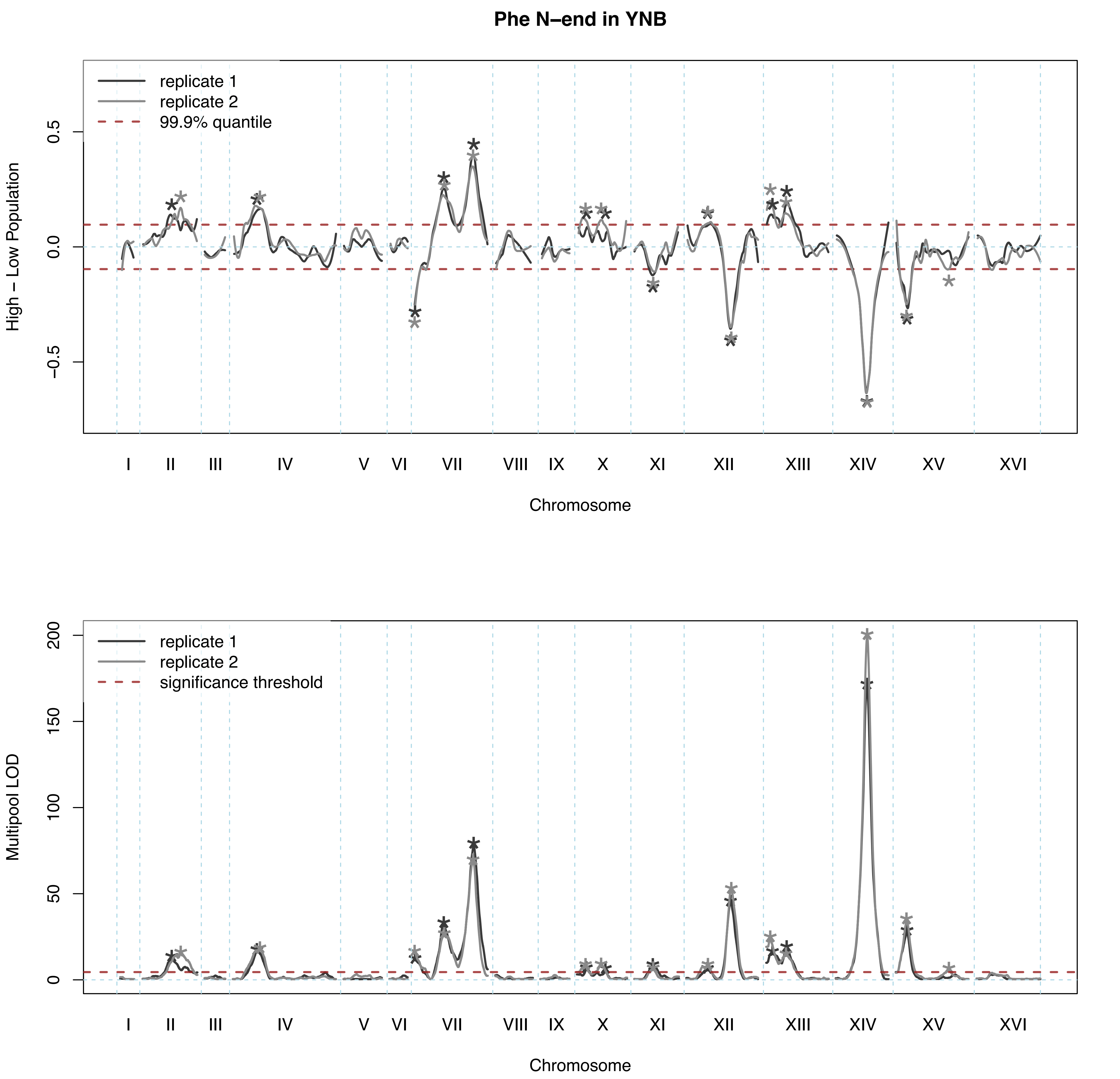

Phe N–end in YNB
replicate 1
replicate 2
99.9% quantile
High – Low Population
0.5
0.0
−0.5
I
II
III
IV
V
VI
VII
VIII
IX
X
XI
XII
XIII
XIV
XV
XVI
Chromosome
replicate 1
replicate 2
significance threshold
Multipool LOD
200
150
100
50
0
I
II
III
IV
V
VI
VII
VIII
IX
X
XI
XII
XIII
XIV
XV
XVI
Chromosome

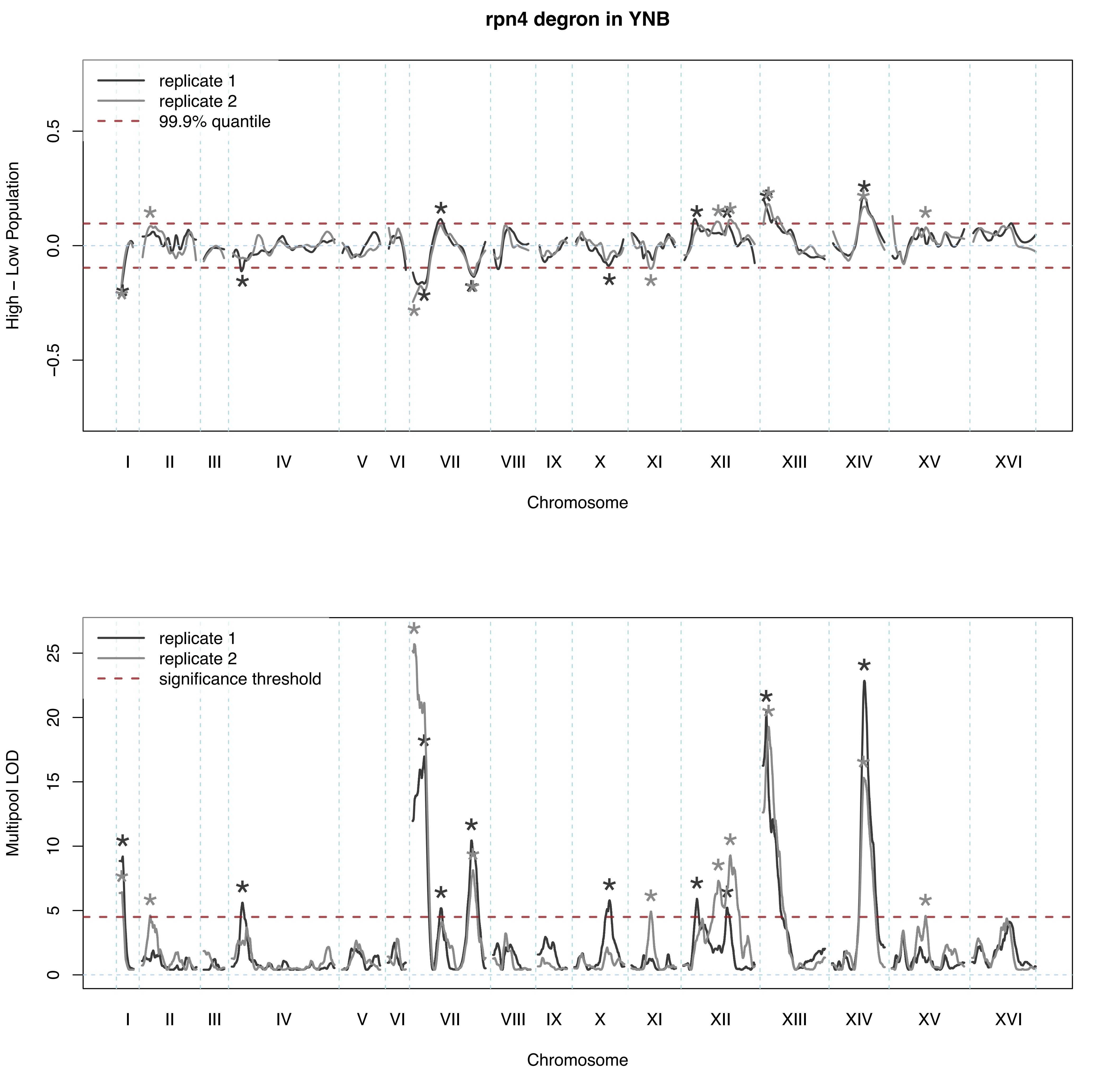
rpn4 degron in YNB
replicate 1
replicate 2
99.9% quantile
High − Low Population
0.5
0.0
−0.5
I II III IV V VI VII VIII IX X XI XII XIII XIV XV XVI
Chromosome
replicate 1
replicate 2
significance threshold
Multipool LOD
0
5
10
15
20
25
I II III IV V VI VII VIII IX X XI XII XIII XIV XV XVI
Chromosome

**Thr N–end in YNB**

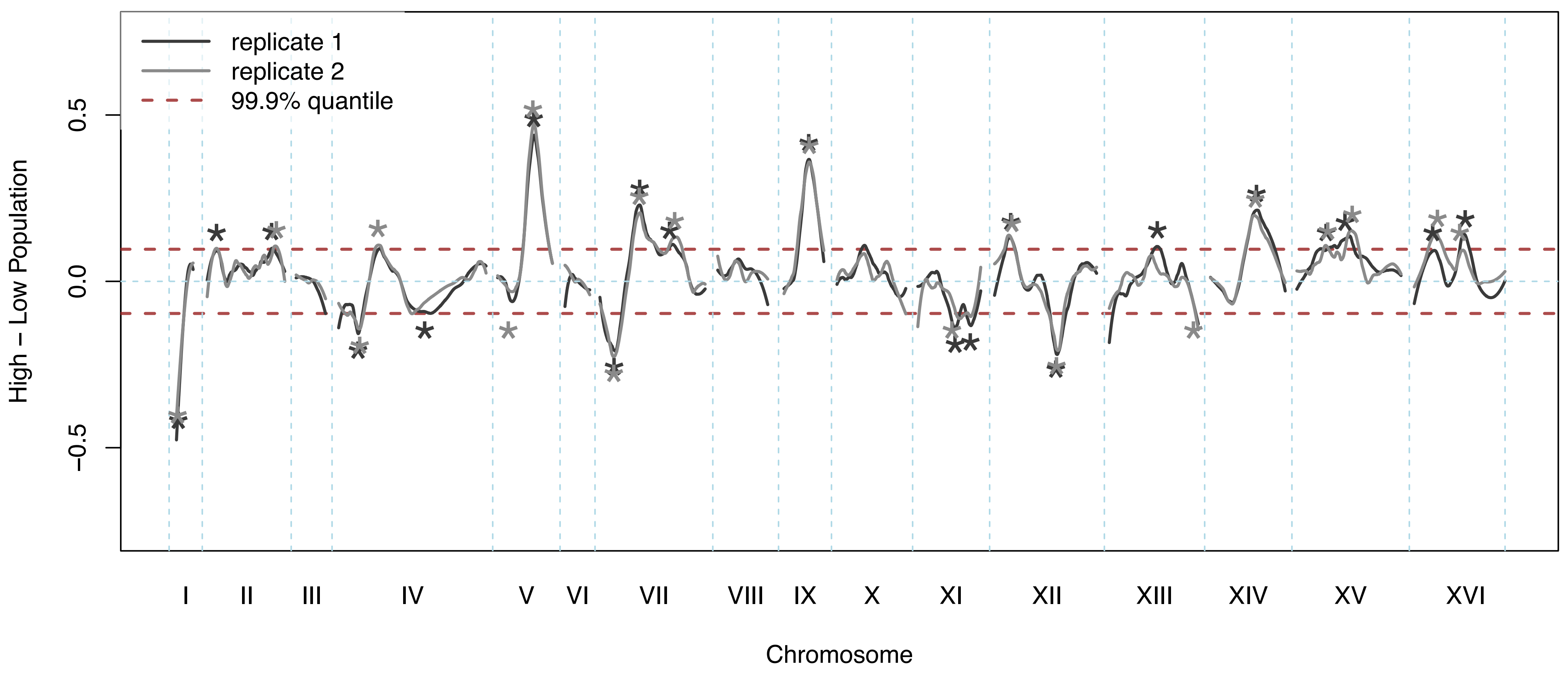


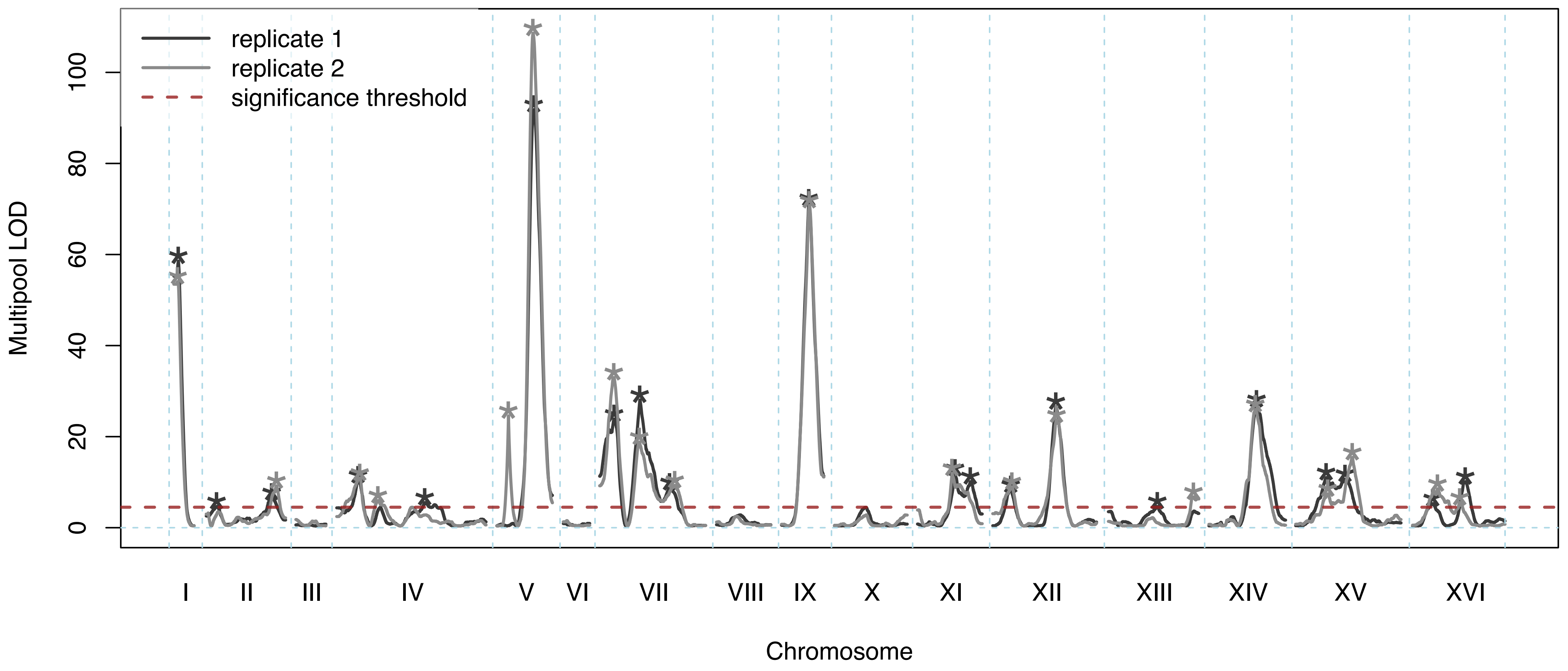

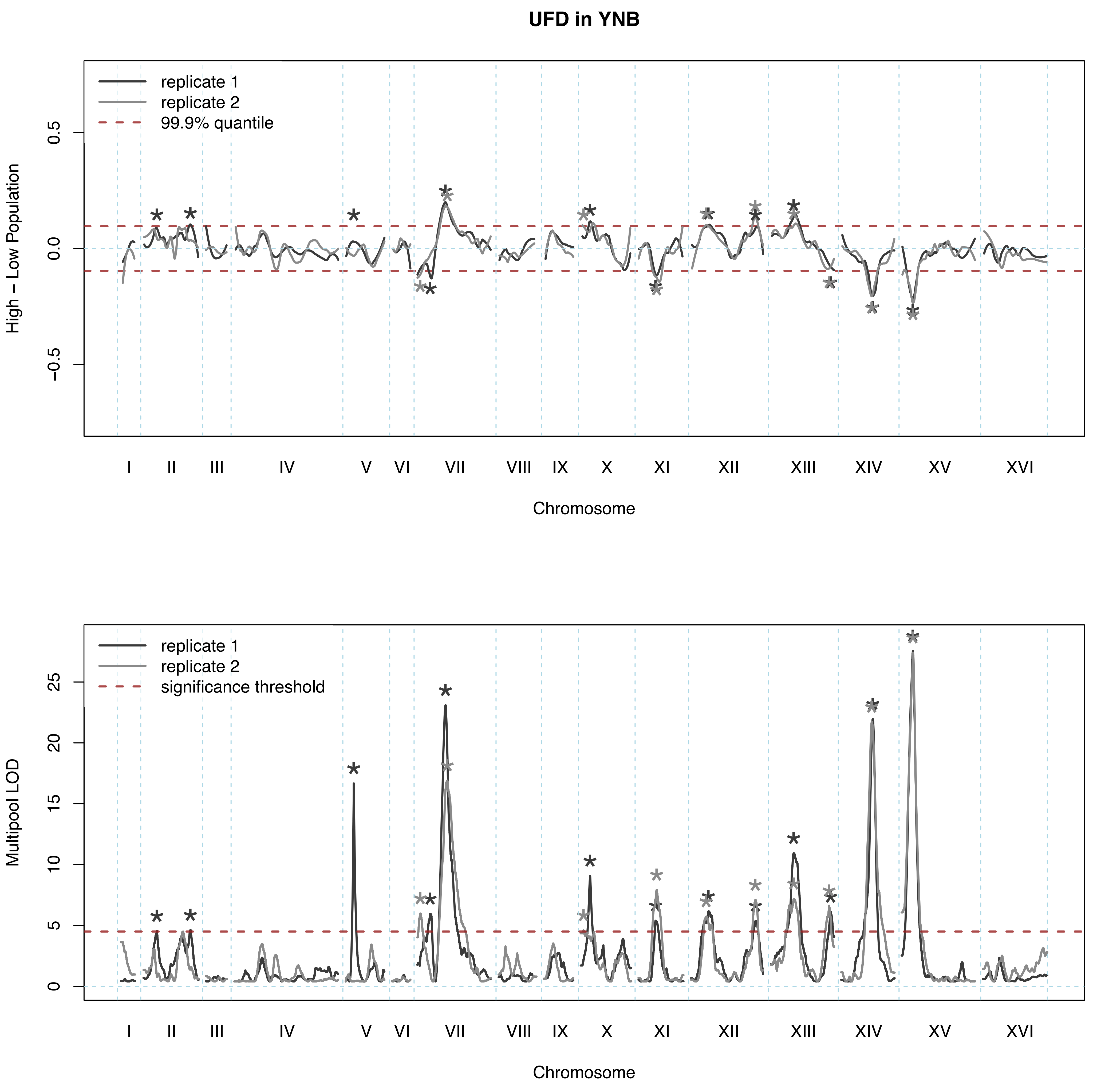
UFD in YNB
replicate 1
replicate 2
99.9% quantile
High − Low Population
0.5
0.0
−0.5
I
II
III
IV
V
VI
VII
VIII
IX
X
XI
XII
XIII
XIV
XV
XVI
Chromosome
replicate 1
replicate 2
significance threshold
Multipool LOD
25
20
15
10
5
0
I
II
III
IV
V
VI
VII
VIII
IX
X
XI
XII
XIII
XIV
XV
XVI
Chromosome
